# Biologically meaningful regulatory logic enhances the convergence rate in Boolean networks and bushiness of their state transition graph

**DOI:** 10.1101/2023.07.17.549398

**Authors:** Priyotosh Sil, Ajay Subbaroyan, Saumitra Kulkarni, Olivier C. Martin, Areejit Samal

## Abstract

Boolean network (BN) models of gene regulatory networks (GRNs) have gained widespread traction as they can easily recapitulate cellular phenotypes via their attractor states. The overall dynamics of such models are embodied in the system’s *state transition graph* (STG) which is highly informative. Indeed, even if two BN models have the same network structure and recover the same attractors, their STGs can be drastically different depending on the type of regulatory logic rules or Boolean functions (BFs) employed. A key objective of the present work is to systematically delineate the effects of different classes of regulatory logic rules on the structural features of the STG of reconstructed Boolean GRNs, while keeping BN structure and biological attractors fixed. Furthermore, we ask how such global features might be driven by characteristics of the underlying BFs. For that, we draw from ideas and concepts proposed in cellular automata for both the structural features and their associated proxies. We use the network of 10 reconstructed Boolean GRNs to generate ensembles that differ in the type of logic used while keeping their structure fixed and recovering their biological attractors, and compute quantities associated with the structural features of the STG: ‘bushiness’ and ‘convergence’, that are based on the number of garden-of-Eden (*GoE*) states and transient times to reach attractor states when originating at them. We find that ensembles employing *biologically meaningful* BFs have higher ‘bushiness’ and ‘convergence’ than those employing random ones. Computing these ‘global’ measures gets expensive with larger network sizes, stressing the need for more feasible proxies. We thus adapt Wuensche’s *Z*-parameter to BFs in BNs and provide 4 natural variants, which along with the network sensitivity, comprise our descriptors of *local* dynamics. One variant of the network *Z*-parameter as well as the network sensitivity correlate particularly very well with the bushiness, serving as a good proxy for the same. Finally, we provide an excellent proxy for the ‘convergence’ based on computing transient lengths originating at random states rather than *GoE* states.

## 1. INTRODUCTION

For several decades now, complex systems tools have been the cornerstone of quantitative analyses of biological systems, in particular for understanding principles of self-organization [1–8]. Some associated foundations had previously been laid in the field of cellular automata (CA) in attempts to apply automaton and computational theories to living systems [9–12]. In those frameworks, multiple types of automata were studied such as 2D CA [9], 1D elementary CA [10, 13, 14], non-uniform CA [15] and sequential dynamical systems [16] to name a few. In a shift towards a focus on *intra-cellular* regulatory processes, Stuart Kauffman proposed that gene networks can be aptly modeled by generalizations of automata, namely via Boolean networks (BN) [1, 17, 18], in which genes can assume 2 states - ‘on’ or ‘off’. BNs are comprised of nodes (corresponding to genes, whereas they correspond to spatially distributed cells in CA) and directed edges between nodes are associated with the regulation of each target gene by its controlling genes (the drivers). Each gene’s expression is updated based on a regulatory logic rule or Boolean function (BF) that has as input the expression levels of the regulators of that gene. Notably, types of regulatory logic rules have been shown to preponderant in reconstructed BNs [19] BNs have been extensively used to model the dynamics of gene regulatory networks (GRNs) to explain a wide range of biological processes including differentiation, metabolism, apoptosis and proliferation, among many others [20–28]. The full dynamics of a BN can be described by a *state transition graph* (STG) in which a fixed point attractor typically corresponds to a biological steady state expression pattern specific to a cell type while its basin of attraction corresponds to all state expression patterns that under the BN dynamics will converge to that steady state [17, 29, 30].

In a framework that is still widely used today, Stephen Wolfram showed how to systematically classify all elementary CA rules into 4 classes based on the dynamical behavior exhibited when simulated from random initial conditions [31]. To date, an analogous framework has not been proposed in the context of BNs though there have been extensive studies on how network structure and BFs affect the dynamics of BNs [32–36]. Thus we ask here how different classes of regulatory logical rules in BNs can affect systemic dynamics through the structural features of the associated STGs. For that, we provide mathematical measures of STG structure by building on previous work studying STGs in CAs, especially by Andrew Wuensche [37–39]. Wuensche studied different features of the STGs of CAs some of which include: (1) ‘bushiness’ which he associated with the fraction of garden-of-Eden (*GoE*) states in the STG (also called the *G*-density); (2) the in-degree distribution of the states of the STG, and (3) the lengths of trajectories going from random states to attractors. This last quantity informs on the transient times of the dynamics, short times corresponding to highly convergent dynamics and long times corresponding to weakly convergent dynamics [40]. In CA, Wuensche observed that highly bushy STGs with shorter transients have more ordered dynamics while the less bushy STGs with longer transients have more chaotic dynamics [40]. Though there has been a lot of work on determining the dynamical regimes of either random BNs [21, 41–44] or specific reconstructed Boolean GRNs [19, 45], such studies never looked at these regimes through the lens of the structural features of the STG. Clearly, in automata having synchronous updates, the dynamics can be represented by a STG in which each “state” of the system has one outgoing edge. The number of incoming edges can be 0, 1 or more for each state. The associated properties of the STGs have been measured in CAs but could also be measured in BNs. To the best of our knowledge, studies of such structural features of STGs have been primarily restricted to elementary CAs and Random BNs [14, 37, 46, 47] and have never been studied in reconstructed models of BNs. It is worth mentioning here that determining the STG and its features can be computationally very expensive, especially with increasing network sizes [48], making it is appropriate to search for proxies of those features. Although this was not done explicitly by Wuensche, in his studies of CAs he introduced the *Z*-parameter, a quantity that can be computed directly from the rule table which measures the probability that the next unknown cell in a partial pre-image is uniquely determined [38, 49, 50]. He showed that this *Z*-parameter shows a marked correlation with the *G*-density, and as a result the *Z*-parameter provides a useful prediction for the bushiness of the STG requiring computations only at the level of the CA rule [38]. In extending such an approach to BNs, one is confronted with the fact that in contrast to the situation in elementary CAs, each node in a BN is allowed to have a different ‘rule’ and there is no natural ordering of its inputs, making it non-trivial to adapt the *Z*-parameter to BNs. This challenge of finding rule-based measures in BNs has not been addressed so far, so we will fill that gap.

The primary objective of this work is to delineate how different classes of logic rules, namely effective functions (EFs), effective and unate functions (EUFs), read once functions (RoFs) and nested canalyzing functions (NCFs), affect the structural features of the STG of a Boolean GRN, and whether there exist proxies that can act as good predictors for those features. To do so, we first select 10 reconstructed Boolean GRNs and for each one generate 4 ensembles of models where each ensemble has the same network structure and reproduces the same set of biological fixed points but differs in the type of logic rule employed. We consider 2 features of the STG characterizing its structure: (1) bushiness and (2) degree of convergence. Given a STG, we quantify its ‘bushiness’ with the measures *G-density* and the *average in-degree of non* − *GoE states* (⟨*d*_*in*_⟩_*non−GoE*_) whereas we quantify its degree of ‘convergence’ with the measure *average convergence rate of trajectories originating at GoE states* (*λ*_*GoE*_). Given these *global* measures on the STG, we ask for each of the 10 *GRN* network structures how our ‘bushiness’ and ‘convergence’ measures vary across the 4 biologically plausible ensembles and particularly how biologically meaningful types of BFs (EUFs, RoFs and NCFs) perform in comparison to the EFs (which are random BFs for all practical purposes). Next we explore the correlation between bushiness and convergence. Following this, we address the challenge posed in the previous paragraph by proposing a scheme based on the permutation of inputs to a BF, to adapt the *Z*-parameter to BFs in BNs. Subsequently we posit 4 variants of the *Z*-parameter at the level of the BF and their respective counterparts at the level of the network which we collectively refer to as the network *Z*-parameters. We then study the distributions of the network *Z*-parameters for the biologically plausible ensembles for each of the 10 reconstructed GRNs. We also look at the distribution of the network sensitivity in the manner described above. Given these *local* descriptors of dynamics, namely, the 4 variants of the network *Z*-parameter and the network sensitivity, we evaluate the correlation between these different measures. Lastly, we inquire whether these descriptors of local dynamics can serve as ‘proxies’ for our two global measures of a STG, namely bushiness and convergence, because determining those two measures become computationally intractable when the networks have more than a few dozen nodes. Interestingly, we find that our descriptors provide useful predictors of bushiness but *not* of convergence. Nevertheless, we show how this difficulty can be circumvented so that both characteristics of the STG can be well predicted without computing the STG.

## 2. METHODS

### 2.1 Boolean network models of gene regulatory networks

A Boolean model of a gene regulatory network (GRN) consists of a set of nodes and directed edges that correspond to genes and the interactions between them respectively [1, 17, 18]. The nodes of a BN can assume one of two states, either ‘on’ or ‘off’, and are thus represented by a Boolean variable that can take values 1 or 0, respectively. We denote the state of node *i* at time *t* by *x*_*i*_(*t*), where *i* ∈ {*i* 1, 2, …, *N*} (*N* is the number of nodes in the network) and *x*_*i*_ ∈ {0, 1}. We denote the state of the network by the vector **X**(*t*) whose *i*^*th*^ entry is given by *x*_*i*_(*t*). At each node of the network, a Boolean function (BF) (or *logical update rule*) is assigned which governs its dynamics, thereby specifying the rule to go from the values of its inputs to the value of its output (see SI text, section 1). More succinctly, 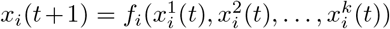, where *f*_*i*_ is the BF that acts on the *k* inputs to node *i*, 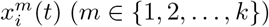, to return the value *x*_*i*_(*t* + 1). If all BFs **F** = (*f*_1_, *f*_2_, …, *f*_*N*_) act on all nodes simultaneously, the update is said to be synchronous [1], otherwise it is asynchronous [51]. In this work, all nodes are updated synchronously as expressed by the following equation:

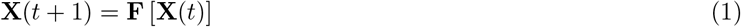

Biologically, the vector **X**(*t*) captures the gene expression pattern of the network at time *t*. Since each node in this network can assume 2 states, **X**(*t*) can take 2^*N*^ possible values, each one corresponding to a gene expression pattern. The set of all gene expression patterns defines the state space of the BN. Trajectories in the state space can be traced out by the recursive application of Eq.(1) to any given initial condition **X**(0). The *state transition graph* (STG) of the BN is just the directed graph in state space where each edge connects a state **X** to its image under Eq.(1). Under synchronous dynamics, a trajectory can meet with 2 possible fates: (a) it reaches a *fixed point attractor, i.e*., a state which on further update remains unaltered, or (b) it ends up in a *cyclic attractor* (or *cycle*), in which case the network keeps cycling through a fixed set of states indefinitely. All the states which converge to an attractor (including the attractor) constitute its basin of attraction. The above described dynamics generates a STG with multiple disconnected components, each one corresponding to one basin of attraction. Typically, fixed points are of biological significance as they represent steady state expression patterns that specify cell types. Furthermore, the set of states in the STG that cannot be obtained by the recursive application of Eq.(1) to any state of the network constitute its *garden-of-Eden* (*GoE*) states. In other words, the *GoE* states do not have any predecessors (or pre-images) in the STG (see Fig. 1 and SI Fig. S1). Another important feature of the STG are ‘transient’ states. All states that are not members of the attractor are called ‘transient’.

**FIG. 1.**
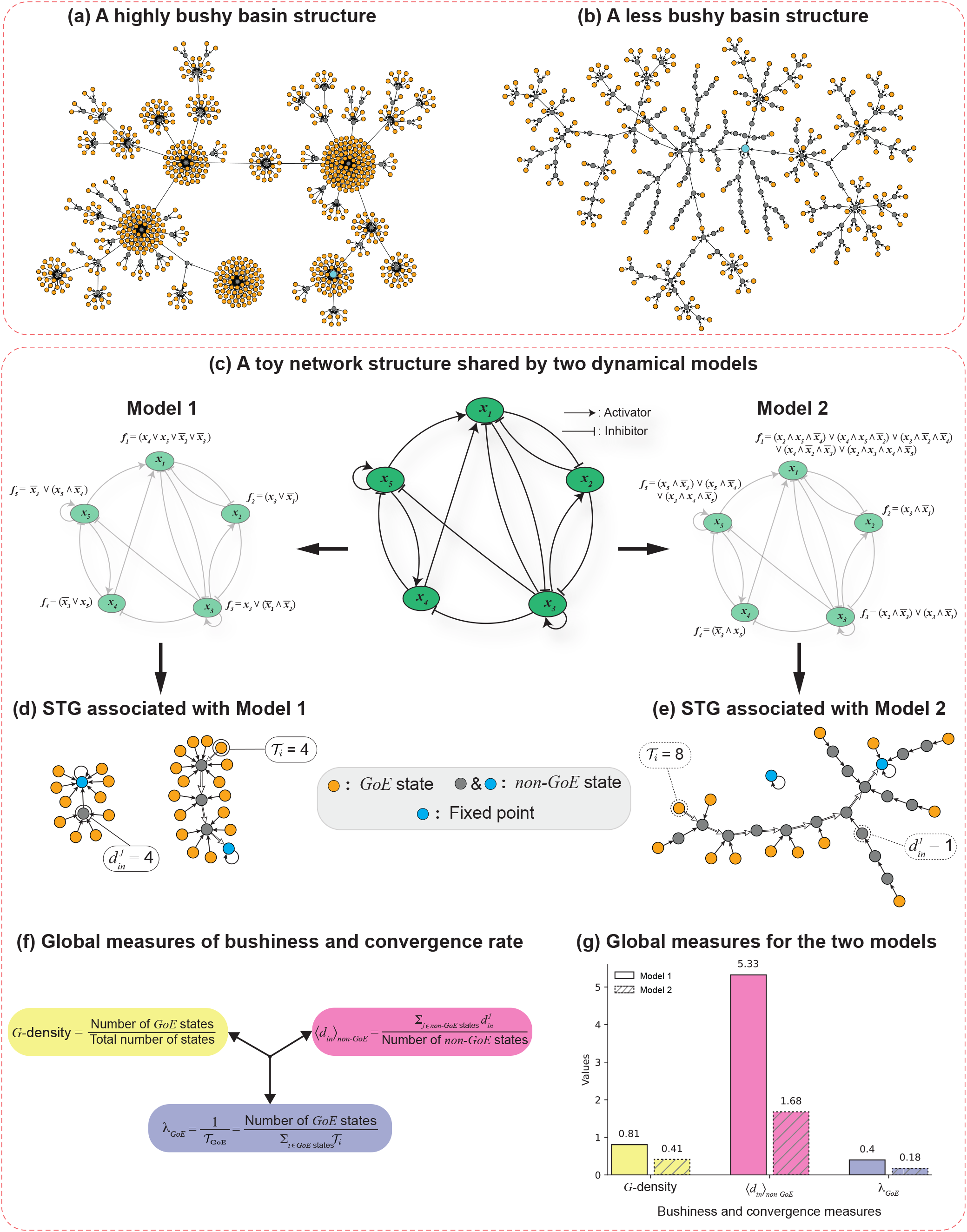
Measures of ‘bushiness’ and ‘convergence’ of STGs. **(a)** and **(b)** correspond to two basins of attraction with fixed point attractors for two different choices of Boolean rules for the biological network MD-GRN. Subplot (a) depicts a highly bushy basin structure whereas subplot (b) a relatively less bushy one. The structure of the biological network MD-GRN, the two dynamical models considered and the complete STGs obtained from the dynamics in these two models are displayed in the supplementary information (see SI Fig. S1). The ‘orange’, ‘blue’ and ‘grey’ nodes indicate the *GoE* states, fixed points and *non-GoE* states that are not fixed points respectively. **(c)** In the center is a toy network structure with 5 nodes and 14 edges. On its two sides are Boolean models (Model 1 and Model 2) with identical network structures but differing regulatory logic rules (as shown at each node). **(d)** and **(e)** display the complete STGs containing 32 states for both Model 1 and Model 2. Visual inspection shows that the STG for Model 1 appears significantly more bushy than the STG for Model 2. The transient path from a chosen *GoE* (encircled) to the attractor is traced via the unfilled arrow heads. 𝒯_*i*_ denotes the transient lengths (steps to reach the attractor) for the *i*^*th*^ *GoE* state (encircled *GoE* states in the subplot) in the STG. 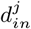 denotes the in-degree of the *j*^*th*^ *non* − *GoE* state (encircled grey *non* − *GoE* states in the subplot) in the STG. **(f)** The global measures that quantify the ‘bushiness’ of the STG are *G*-density and average in-degree of *non−GoE* states (⟨*d*_*in*_⟩_*non−GoE*_), whereas the measure that quantifies the ‘convergence’ is the average convergence rate of trajectories originating from *GoE* states (***λ***_***GoE***_). **(g)** Barplots representing the values of the bushiness and convergence measures described in (f) for Model 1 and Model 2. The *x* and *y* axes correspond to the three measures and their values respectively. The bars for the two models for the same quantities have been shown together to illustrate quantitatively the visible differences between (d) and (e).

### 2.2 Sampling the space of biologically plausible models of 10 published Boolean GRNs

This work provides a quantitative study of how the type of BFs can affect the bushiness and convergence of the STG. To achieve this, we first choose 10 reconstructed Boolean GRNs that have been published in the literature. The list of the 10 models, their abbreviations and references are provided in Table 1. The BFs used in each model are used to obtain the network structure of the GRN. These BFs are provided in the BoolNet format in SI Tables S1-S10. The type of BF used at each of the nodes in all 10 models is provided in SI Tables S11-S20. Note that all the BFs employed in all the models are unate [52], that is each input is signed (is either activatory or inhibitory, regardless of the values of the other inputs). Next, for each of the GRNs we generate 4 biologically plausible ensembles, each one constrained to use only one of the 4 types of BFs, namely, EF, EUF, RoF and NCF (see SI text, section 2 for description of types of BFs) while keeping the network structure of the GRN fixed and satisfying the biological fixed points [53, 54]. When using BFs of the EUF, RoF and NCF types, we also respect the signs of the regulatory interactions, but do not do so when using BFs of the EF type since such functions do not always allow one to define the sign of an interaction. The biological fixed points for the 10 models are provided in SI Tables S21-S30. After applying the above procedure, we determined, for each of the four ensembles of each of the 10 published models, the number of allowed functions for each node; these numbers are given in SI Tables S31-S40. These tables show that in spite of the multiple constraints applied, the space of biologically plausible models is still extremely large. Hence we sampled 10^6^ models from those ensembles in which the number of models exceeded 10^6^ (except for FOS-GRN and GSD-GRN where we sampled 10^5^ models), and used the exhaustive set of models otherwise. Our procedure to sample these ensembles is provided in SI text, section 3. We shall refer to these ensembles (sampled or exhaustively enumerated) of models for the different types of BF as EF-ensemble, EUF-ensemble, RoF-ensemble and NCF-ensemble.

**TABLE 1.**
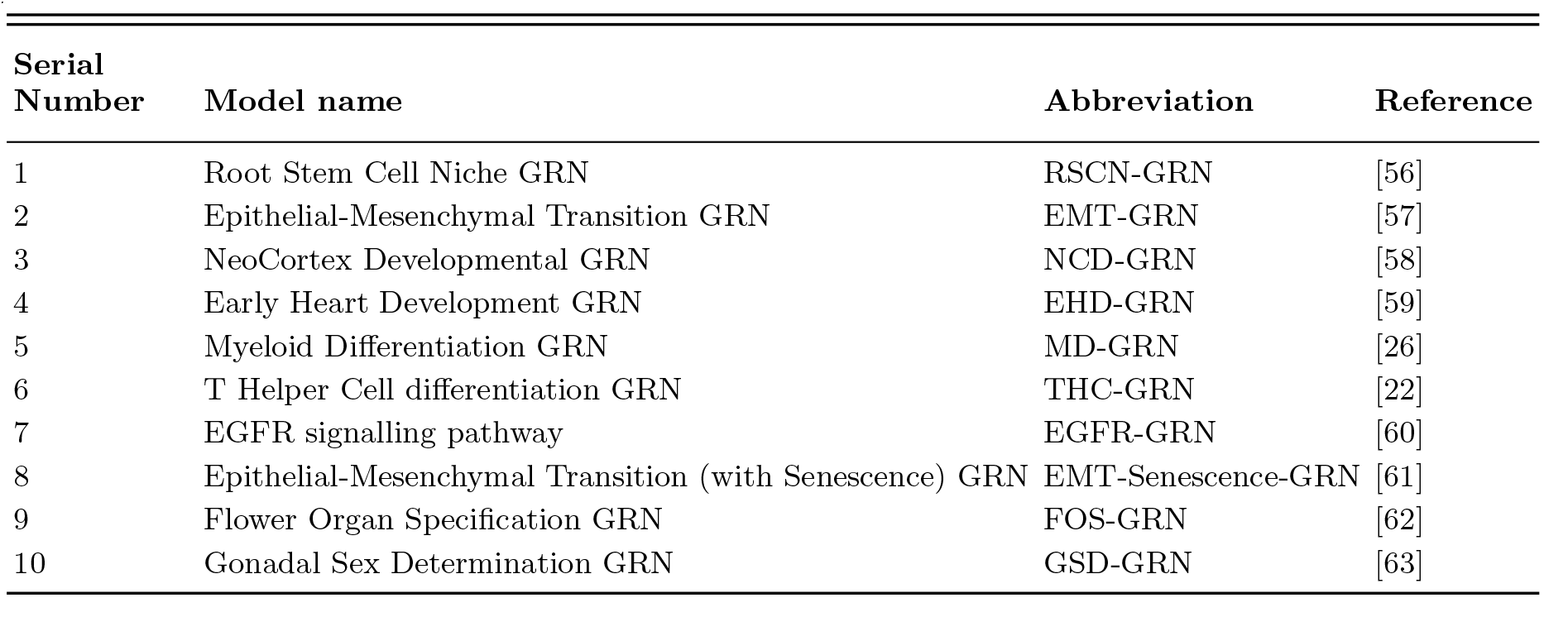
10 published reconstructed Boolean GRN models. ‘Model name’ column provides the names of each of the Boolean GRN network structures that are used in this work. The ‘Abbreviation’ column provides the abbreviated form we designate for each of the 10 models. The ‘Reference’ column provides the research articles associated with each of these 10 models.

| Serial Number | Model name | Abbreviation | Reference |
| --- | --- | --- | --- |
| 1 | Root Stem Cell Niche GRN | RSCN-GRN | [56] |
| 2 | Epithelial-Mesenchymal Transition GRN | EMT-GRN | [57] |
| 3 | NeoCortex Developmental GRN | NCD-GRN | [58] |
| 4 | Early Heart Development GRN | EHD-GRN | [59] |
| 5 | Myeloid Differentiation GRN | MD-GRN | [26] |
| 6 | T Helper Cell differentiation GRN | THC-GRN | [22] |
| 7 | EGFR signalling pathway | EGFR-GRN | [60] |
| 8 | Epithelial-Mesenchymal Transition (with Senescence) GRN | EMT-Senescence-GRN | [61] |
| 9 | Flower Organ Specification GRN | FOS-GRN | [62] |
| 10 | Gonadal Sex Determination GRN | GSD-GRN | [63] |

For 7 and 8 input nodes, it is not possible to obtain an exhaustive list of UFs, hence we proposed a method to sample UFs (and EUFs) that obey a given sign conformation and satisfy given biological fixed points. Our algorithm is based on an iterative color propagation scheme on the vertices of the Boolean hypercube (see SI text, section 3) that is induced by the monotonicity of the output as a function of inputs. SI Fig. S2 and SI Fig. S3 provide a detailed visual explanation of the algorithm. We note that though the distribution of UFs sampled using our iterative color propagation algorithm is not perfectly uniform, it does not deviate much from the uniform distribution (see SI Fig. S4).

In sum, for each of the 10 published GRNs (see SI text, section 4 for a brief overview of the 10 GRNs), we generate 4 ensembles, each constrained by a given type of BF that satisfies the biological fixed point constraints. We generate the STG for all the models in these ensembles using the R package BoolNet [55] and compute its bushiness and convergence using the different measures described below.

### 2.3 Global measures of bushiness and convergence of the state transition graph

To illustrate the notion of ‘bushiness’ visually, we first consider the STGs of 2 Boolean models (model A and model B) derived from a myeloid differentiation GRN (MD-GRN) that differ in their regulatory logic rules but recover the same biological attractors. Figs. 1(a) and 1(b) display the largest basins of attraction with fixed point attractors for these models (see SI Fig. S1 for full STG). It is visually clear that the STG in Fig. 1(a) is much more ‘bushy’ than that in Fig. 1(b). The complete STGs of these 2 MD-GRN Boolean models (SI Fig. S1) reveal that all the basins of attraction of model *A* are more bushy compared to those of model *B*. In this section, we quantify this intuitive notion of ‘bushiness’ of STGs using 2 measures, namely, *G*-density and average in-degree of non-*GoE* states. We also study ‘convergence’ using the average transient length of *GoE* states from which we extract a measure of the convergence rate of the dynamics in state space. To define these measures, we show how they are computed in the simple cases displayed in Fig. 1(c) that are based on toy models (model 1 and model 2) which share the same network structure but differ in the BFs used. Our three measures quantitatively capture the ‘bushiness’ and degree of ‘convergence’ of STGs as illustrated for models 1 and 2 in Figs. 1(d) and 1(e) respectively. Qualitatively, the STG in Fig. 1(d) seems more bushy than the STG in Fig. 1(e). To make this quantitative, we require that a bushy STG have high values for its *G*-density and average in-degree of non-*GoE* states. Similarly, if the dynamics is to be considered as highly convergent, we require that trajectories originating at *GoE* states have low average transient lengths. The reader may expect these two properties (bushiness and high convergence) to go hand in hand but in fact the situation is more subtle.

#### 2.3.1 G-density

In CA and RBN, *G*-density (or leaf density [64]) is considered as the simplest measure of ‘bushiness’ of a STG [38, 47, 49]. *G*-density is defined as the fraction of states that are *GoE* (Fig. 1(f)). Fig. 1(g) shows that the *G*-density of the STG of model 1 (Fig. 1(d)) is higher than that of the STG of model 2 (Fig. 1(e)). Wuensche [49] showed that in CA, high *G*-density is associated with ordered dynamics, whereas low *G*-density is associated with chaotic dynamics.

#### 2.3.2 Average in-degree of non-GoE states

Another intuitive characterization of the structure of the STG that has been studied in CA and RBN is based on its in-degree distribution [46, 49]. The in-degree of a state in the STG corresponds to the number of predecessors or pre-images under Eq.(1) of that state [49]. In this work, we propose the average of the in-degrees over all the *non-GoE* states as a measure of bushiness and denote it with the symbol ⟨*d*_*in*_⟩_*non−GoE*_. We exclude the *GoE* states from the definition since the average of the in-degrees over all the states of the STG is always exactly 1. Indeed, under synchronous update, each state in the STG has an out-degree of 1 and so the number of states and of edges in the STG are equal to 2^*N*^. It is expected that a highly bushy STG will have a large ⟨*d*_*in*_⟩_*non−GoE*_ compared to a less bushy STG. Fig. 1(g) shows that the ⟨*d*_*in*_⟩_*non−GoE*_ of the STG of model 1 (Fig. 1(d)) is higher than that of the STG of model 2 (Fig. 1(e)). Furthermore, we show that ⟨*d*_*in*_⟩_*non−GoE*_ can be expressed as a function of the *G*-density as follows:

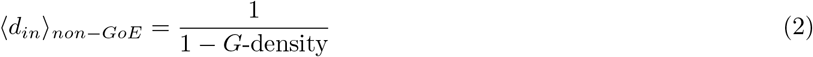

We provide the derivation of the above expression (Eq.(2)) in SI text, section 5.

#### 2.3.3 Average convergence rate of trajectories originating at GoE states

Another feature that can distinguish different STGs is the length of ‘transients’ of trajectories in its state space [40]. Transients are states in the STG that lie on trajectories which lie outside the attractors. For any state, the number of steps required to reach the attractor is the ‘transient length’ associated with that state. A highly convergent STG is expected to have shorter transient lengths on average compared to less convergent STGs [40, 49].

In this work, we first define the average transient length of *GoE* states (𝒯_*GoE*_) as the average over the lengths of transients originating at all the *GoE* states. We then define the *average convergence rate* of trajectories originating at *GoE* states (*λ*_*GoE*_) as the reciprocal of the average transient length of *GoE* states (1/ 𝒯_*GoE*_). Fig. 1(g) shows that the *λ*_*GoE*_ of the STG of model 1 (Fig. 1(d)) is higher than that of the STG of model 2 (Fig. 1(e)). As a variant to *λ*_*GoE*_, we also consider the reciprocal of the average of the transient lengths of trajectories originating at *all* states of the STG, which we denote as *λ*_*all*_(= 1/𝒯_*all*_). This quantity has the advantage that it can be estimated numerically via sampling, i.e., rather than considering all states, one can use a sample of random states; that (stochastic) estimator will be denoted as *λ*_*random*_(= 1/𝒯_*random*_).

### 2.4 Adaptation of a local descriptor of dynamics in CA to Boolean networks

In CA, the *Z*-parameter as defined by Wuensche is derived from an algorithm of pre-image calculation in the state space [49] (see SI text, section 6 for details). His *Z*-parameter is supposed to reflect the degree to which pre-images are identifiable within the considered automaton update rule [49]. That measure was introduced to indicate whether a random state is likely to have a high number of pre-images (low *Z*) or a low number of pre-images (high *Z*). To date, a parameter that is analogous to the *Z*-parameter defined on rules in CA is lacking for BFs in BNs, even though both CA and BNs share similar discrete dynamics.

For any one-dimensional CA rule, Wuensche introduces a computational procedure to define two intermediate values, *Z*_*left*_ and *Z*_*right*_, associated with reading the inputs from left to right and from right to left respectively, and then he defines the *Z*-parameter as the maximum of *Z*_*left*_ and *Z*_*right*_ [38] (see SI text, section 7 for details). The difficulty of adapting this definition to BNs lies in the fact that in 1D CAs, each cell has a well defined spatial relation to its neighbors that is then used to define *Z*_*left*_ and *Z*_*right*_ (see SI text, section 7), whereas in BNs, there is no notion of ‘left’ or ‘right’ of a node. We thus propose a procedure to define the *Z*-parameter for BFs in BNs as follows.

#### 2.4.1 Calculation of Z_left_

Suppose we are given a *k*-input BF *f*. We create its truth table as a succession of rows using the standard lexicographic ordering of the values of the *k* inputs *x*_1_, *x*_2_, …, *x*_*k*_, each of those Boolean variables being used to label the table’s successive columns. We can then use Wuensche’s prescription for a computational definition of *Z*_*left*_ [49] as follows.

For any index *i* ∈ {1, 2, …, *k* − 1}, one first defines a “block” of rows in the truth table as a set of rows sharing the same values of *x*_1_, *x*_2_, …, *x*_*k−i*_. For *i* = *k*, there is a unique block and it consists of all the rows of the truth table. Because of the lexicographic order imposed on the truth table, a block consists of 2^*i*^ consecutive rows and will therefore be referred to as a 2^*i*^-block (see SI text, section 1). Note that at given *i*, all 2^*i*^-blocks are disjoint while their union consists of all the rows of the truth table. Then, for any 2^*i*^-block, Wuensche determines whether it satisfies a switch-like property:

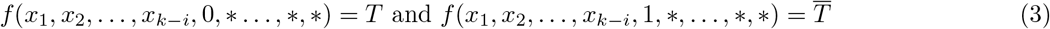

where *T* ∈ {0, 1} and 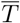 is its complement, while the ∗s mean that the two equalities are to be satisfied regardless of the corresponding input values of *x*_*k−i*+2_, …, *x*_*k*_ (for *i* = *k*, all but the first of the arguments are ‘*’ in Eq.(3)). If a 2^*i*^-block satisfies Eq.(3), we say it is “complement canalyzing” because, in the context (*x*_1_, *x*_2_, …, *x*_*k−i*_), the values 0 and 1 for *x*_*k−i*+1_ are both canalyzing and lead to complementary outputs. To be concise, a complement canalyzing 2^*i*^-block will be abbreviated as a CC-2^*i*^-block. It is worthwhile to note that all complement canalyzing blocks are disjoint, so that a CC-2^*i*^-block and a CC-2^*j*^-block will have no rows in common. We then consider the vector **n** = (*n*_1_, *n*_2_, …, *n*_*k−*1_, *n*_*k*_) where *n*_*i*_ is the sum of the cardinalities of the CC-2^*k−i*+1^-blocks, that is the total number of rows in the truth table belonging to such blocks. Given the vector **n**, Wuensche defines the fractions *R*_1_, *R*_2_, …, *R*_*k*_:

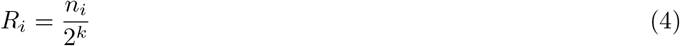

Since the complement canalyzing blocks are all disjoint, the sum of the *R*_*i*_ is at most 1. Finally, the parameter *Z*_*left*_ for this BF *f* (with the given ordering of inputs) is given by:

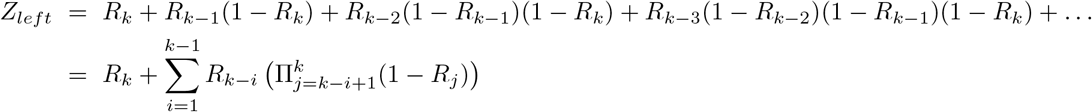

The computation of the *Z*_*left*_ parameter (which is always between 0 and 1) is illustrated on an example in Figs. 2(a), (b) and (c). Fig. 2(a) shows Wuensche’s approach to compute the elements of **n** by iterating over all candidate 2^*i*^-blocks, beginning with *i* = 1. Fig. 2(b) illustrates our alternate approach that is based on recursively visiting (*k* − *i* + 1) dimensional hyperplanes (these have a direct correspondence with the 2^*k−i*+1^-blocks), going from the largest blocks to the smallest ones and checking at each step of the recursion whether the hyperplanes contribute to **n**. Having obtained **n**, Fig. 2(c) shows how to compute *Z*_*left*_. The pseudocode for this algorithm is provided in SI text, Algorithm 1. Some interesting properties of *Z*_*left*_ for any BF are: (1) *Z*_*left*_ of *f* is invariant under the negation of any of its inputs (see SI text, section 8, Property 8.1). (2) *Z*_*left*_ of *f* is invariant under complementation (see SI text, section 8, Property 8.2).

**FIG. 2.**
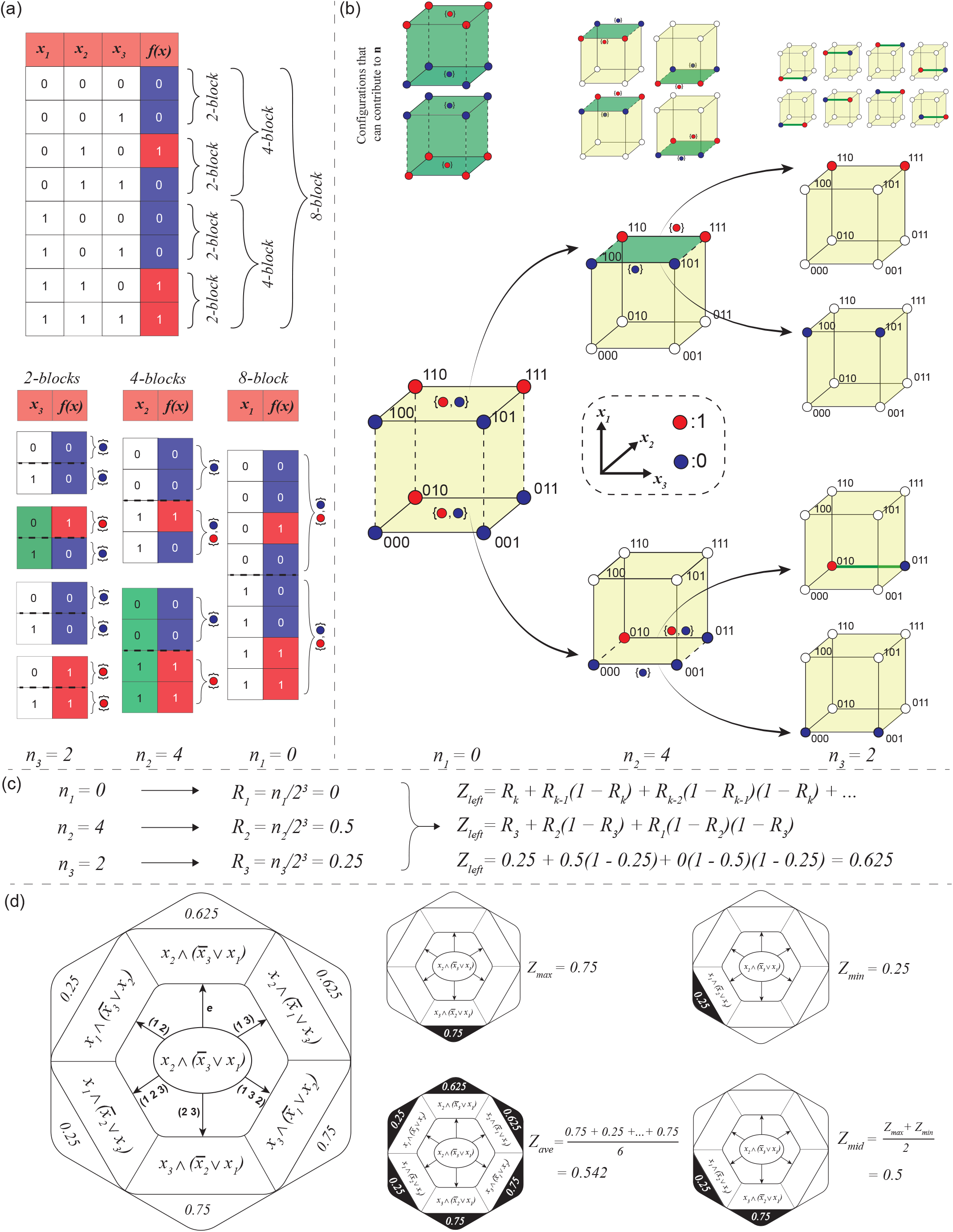
Calculation of *Z*-parameters of Boolean functions in Boolean network models. **(a)** Calculation of the vector **n** from the truth table as proposed by Wuensche. Top panel: The truth table of a particular BF for which the *Z*-parameters will be computed. The colors red and blue correspond to the output values 1 and 0 respectively. The list of 2^*i*^-blocks are shown via braces, where *i* ∈ {1, 2, 3}. Bottom panel: For every 2^*i*^-block, get the values spanned by the output in its top (respectively bottom) half (these halves are separated by the dashed lines). The corresponding values are shown to the right of the braces. If there is a single value for both top and bottom braces and if these are different, then we have a CC-2^*i*^-block and it contributes 2^*i*^ to **n**. The corresponding cells in the column of the (*k* − *i* + 1)^*th*^ variable are colored in green. *n*_1_, *n*_2_ and *n*_3_ are the contributions from the CC-2^3^-, CC-2^2^- and CC-2^1^-blocks (of values 0, 4 and 2 here) defining the vector **n. (b)** An equivalent formalism for calculating **n** on the Boolean hypercube. This figure illustrates how elements of **n** can be computed by recursively splitting the Boolean hypercube into 2 hyperplanes, obtaining at each step an element of **n** starting from *n*_1_. Top panel, from left to right: the cubes show which configurations (vertex colorings) can contribute to *n*_1_, *n*_2_ and *n*_3_ respectively. For instance, a 2^3^-block contributes to *n*_1_ if and only if all the vertices in the plane *x*_1_ = 0 are colored red (or dark blue) *and* all those in plane *x*_1_ = 1 are colored dark blue (or red), as shown at the top left of this subfigure. Bottom panel: By comparing the actual node coloring to the templates contributing in the top panel, we see that there is no contribution to **n** from the 2^3^-block, so *n*_1_ = 0. The next step of the recursion (just to the right) concerns contributions of 2^2^-blocks (associated with two faces of the cube). Here, the upper face (colored dark green) contributes to *n*_2_ whereas the lower face does not (based again on comparing to the templates just above them). Finally, still on the right of that, we have the case of the 2^1^-blocks that may contribute to *n*_3_. We have colored in green the ones that do contribute. Reaching the 2^1^-blocks marks the end of the recursion. Note that if a grouping (cube, face or in higher dimensions any hyperplane) contributes to **n**, then none of its sub-parts will contribute to **n. (c)** This subfigure summarizes the computation of *Z*_*left*_ of a BF once the vector **n** is obtained. **(d)** Computation of the four *Z*-parameter variants: *Z*_*max*_, *Z*_*min*_, *Z*_*ave*_ and *Z*_*mid*_. Left panel: The Boolean expression of the BF in the truth table in sub-figure (a) is *x*_2_ (*x*_3_ *x*_1_), as displayed in the center of the hexagon. We then apply the *k*! permutations (cf. arrows) to produce the (distinct) permuted BFs as shown at the end of these arrows. The values shown in the outer hexagon are the *Z*_*left*_ values for each of the permuted Boolean expressions. Right panel: a hexagon is produced for each of the 4 variants of the *Z*-parameter: *Z*_*max*_ is the maximum *Z*_*left*_ value over all permutations. *Z*_*min*_ is the minimum *Z*_*left*_ value over all permutations. *Z*_*ave*_ is the average of the *Z*_*left*_ over all permutations. *Z*_*mid*_ is the average of *Z*_*max*_ and *Z*_*min*_.

#### 2.4.2 Defining other Z-parameters

In this section we propose a number of other extensions to BFs of Wuensche’s *Z*-parameter. Our definitions are based on the fact that since there is no specific spatial order or arrangement of the inputs to a node in a BN, all possible orderings or permutations of input variables (see SI text, section 1) should be accounted for when defining the *Z*-parameter. Mathematically, this is equivalent to considering the permutations in the symmetric group *S*_*k*_ to the order of input variables when constructing the truth table of the Boolean function *f*. Each permutation ***σ*** ∈ *S*_*K*_ leads to a unique rearrangement of the input variables and a new rearranged truth table, and for each of these we compute its corresponding *Z*_*left*_ value. With this collection of *Z*_*left*_ values, we propose the following inequivalent definitions to extend Wuensche’s *Z*-parameter to BFs in BNs.

***Z***_***max***_: The first such variant, that we denote by *Z*_*max*_, is constructed in the same spirit as Wuensche by considering a maximum. In our case, we take the maximum of all the *Z*_*left*_ values when considering all permutations of the *k* inputs to that BF. Interestingly, we find that NCFs have the maximum *Z*_*max*_ value for any *k*-input BF with odd bias *P* and is given by:

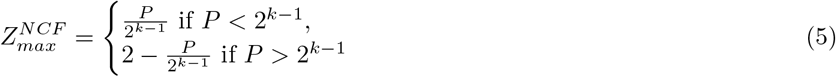

The associated proof is provided in SI as Properties 8.3 and 8.4.

***Z***_***min***_: The second possibility we consider is the *minimum* of this collection of *Z*_*left*_ values of the BF *f* and we denote it by *Z*_*min*_. In this case, we find that NCFs have the minimum *Z*_*min*_ for any *k*-input BF with odd bias *P* and is given by:

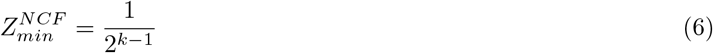

Thus *Z*_*min*_ for NCFs is solely dependent on the number of inputs *k* and independent of the bias. The associated proof is provided in SI as Properties 8.5 and 8.6.

***Z***_***ave***_: The third possibility we consider is the average of the collection of the *k*! values of *Z*_*left*_ which we denote by *Z*_*ave*_. For a BF with *k*-inputs, different permutations on the input variables may or may not result in the same BF. We have been able to show that computing *Z*_*ave*_ as mentioned above is equivalent to computing the *Z*_*left*_ values only for the non-equivalent permutations of the BF and averaging over them. That observation led us to the following 2 conjectures for any *k*-input BF. (1) The number of non-equivalent permutations divides *k*!. (2) Each of the *m non-equivalent* permutations occurs an equal number of times.

Using a group-theoretic approach we prove these conjectures in the SI text, section 9. This result speeds up the computation of *Z*_*ave*_ since now *Z*_*left*_ need not be computed for all *k*! permutations of a BF. Additionally, we find that NCFs and RoFs have a much lower fraction of non-equivalent permutations compared to EFs and EUFs (see SI Fig. S5). In fact, we find that BFs used in the reconstructed BNs have a much lower number of non-equivalent permutations of BFs than when drawing the BFs at random within each of the corresponding type of BF (see SI Fig. S5). These observations highlight that our method to compute *Z*_*ave*_ can significantly reduce computation time in practice.

***Z***_***mid***_: The final possibility we consider is the average of *Z*_*max*_ and *Z*_*min*_, ((*Z*_*max*_ + *Z*_*min*_)/2) and we denote it by *Z*_*mid*_. Using formulas of *Z*_*max*_ (Eq.(5)) and *Z*_*min*_ (Eq.(6)) for any NCF, the formula for *Z*_*mid*_ for any *k*-input NCF with bias *P* is (see SI text, section 8, Property 8.7):

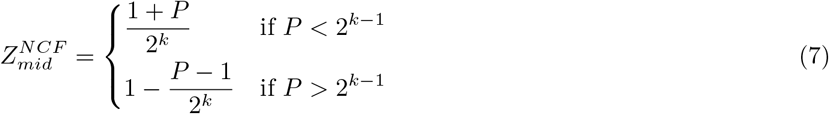

Note that the *Z*_*max*_, *Z*_*min*_ and *Z*_*mid*_ of a NCF are dependent only on the number of inputs and the bias and hence we do not generate its permutations, making its calculation simple and efficient.

#### 2.4.3 Z-parameter of a Boolean network

In CAs, cells receive inputs from their nearest neighbors and all use the same rule (the logic is thus homogeneous across all cells). In such a situation, there is no distinction between a *Z*-parameter at the level of a single cell (or node) and at the level of the network. But in BNs, there is heterogeneity in both the number of inputs and in the logic rules at different nodes. Consequently, we define the *network Z*-parameter via the average of the considered *Z*-parameter over all the nodes of the network. Specifically,

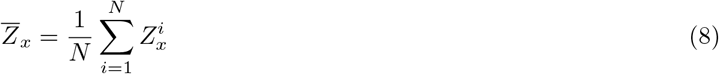

Here, *i* ∈ {1, 2, 3, …, *N*} and is the index of a node in the network, and 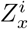 is the *Z*-parameter of the BF at node where *x* ∈ {*max, min, ave, mid*}. In this work the network *Z*-parameters are denoted by 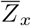.

### 2.5 Average sensitivity of a BF and network sensitivity

The average sensitivity of a BF indicates how sensitive it is to one-bit flips of its inputs [65]. For a *k*-input BF *f*, the sensitivity for a specific assignment of input variables *x* = (*x*_1_ = *a*_1_, *x*_2_ = *a*_2_, …, *x*_*k*_ = *a*_*k*_) is determined by counting the number of neighboring assignments *y* of *x* that yield a different output *f* (*y*) compared to *f* (*x*). Assignments *y* and *x* are called “neighbors” if they differ in exactly one of their *k* variables. The average sensitivity of a BF is obtained by calculating the average of the sensitivities over all possible input combinations and is expressed as:

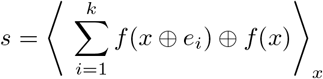

where ⊕ is the logical XOR operator and *e*_*i*_ ∈ {0, 1} ^*k*^ is the unit vector with *x*_*i*_ = 1 and *x*_*j*_ = 0 ∀ *j ≠ i*. We denote the average sensitivity of a BF by *s*.

The *network* sensitivity is then defined as the mean over all nodes of the network, of the average sensitivities of the BFs at each node [45, 65]. We denote the network sensitivity of a BN by 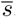.

## 3. RESULTS

To date, studies of the bushiness and convergence of STGs have primarily been restricted to CA and Random BNs (RBNs) [40, 47]. Here, we not only quantify the bushiness and convergence of STGs in numerous reconstructed Boolean GRNs but we also study which properties of the underlying rules drive that high bushiness and convergence. Specifically, we provide a connection between the values of different measures of STG bushiness and the values of particular descriptors of local dynamics. This connection is analyzed for 4 classes of Boolean rules by working in each case with an associated ensemble of models (EF-ensemble, EUF-ensemble, RoF-ensemble and NCF-ensemble). To ensure representative results, we perform these studies in the context of 10 reconstructed Boolean GRNs (see section 2.2 and SI text, section 3). For brevity, the main part of the manuscript presents results only for 4 of the 10 reconstructed GRNs (RSCN-GRN, EMT-GRN, NCD-GRN and EHD-GRN), and those pertaining to the remaining 6 GRNs can be found in the SI. Also, in what follows and unless stated otherwise, we use the term ‘distribution’ of a quantity for the distribution of that quantity in the EF, EUF, RoF and NCF ensembles, for each of the 10 reconstructed GRNs. To provide additional context to our results, it is appropriate to note that EFs are practically ‘random’ BFs since their fraction in the space of all BFs tends to 1 with an increasing number of input regulators [19]. This is not true for the other types BFs, namely EUFs, RoFs and NCFs which represent only a tiny fraction of all BFs in that limit.

### 3.1 Biologically meaningful functions lead to highly bushy and convergent state transition graphs

We first obtain the distribution of the *G*-density as shown by the box plots in Fig. 3 and SI Fig. S6. On comparing the mean values of the distribution of *G*-density across the different ensembles, we observe two striking and consistent patterns across all 10 GRNs (see Fig. 3 and SI Fig. S6). First, STGs in ensembles employing biologically meaningful regulatory logic (EUF-ensemble, RoF-ensemble and NCF-ensemble) are more bushy than STGs in the EF-ensemble. Second, among ensembles generated with biologically meaningful regulatory logic, the STGs in the NCF-ensemble are typically more bushy than those in the EUF-ensemble or RoF-ensemble. Note that higher *G*-densities are indicative of greater order in the dynamics [40].

**FIG. 3.**
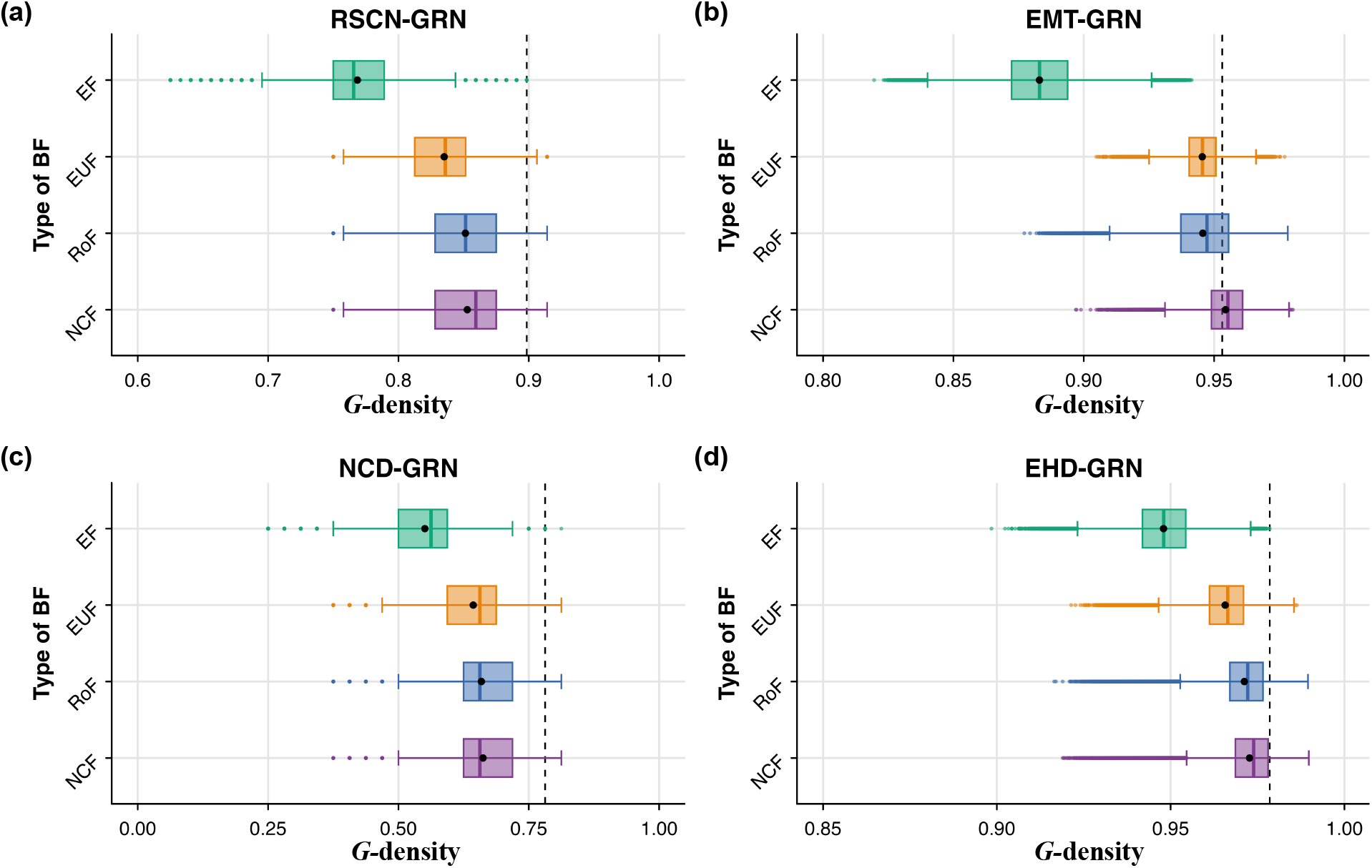
Distribution of *G*-density values for different ensembles generated using network structures from 4 published Boolean GRNs. The subplots **(a), (b), (c)** and **(d)** correspond to the RSCN-GRN, EMT-GRN, NCD-GRN and EHD-GRN network structures respectively. The *x* and *y* axes of each subplot correspond respectively to *G*-density and to the type of BF imposed on the nodes of the networks. The box plots display the distribution of *G*-density in the 4 ensembles that each use a given type of regulatory logic. The ensembles generated using biologically meaningful BFs (EUFs, RoFs, NCFs) have higher *G*-density compared to the ensemble generated using EFs. Note that we impose that ensembles recover the biological fixed points of the associated published model. The mean and median of the distribution are indicated by the black dot and the vertical line within the box respectively. The vertical dashed lines in each subplot correspond to the *G*-density of the Boolean model provided by the modelers in the published article.

Similarly, we consider the distribution of the average in-degree of *non* − *GoE* states (⟨*d*_*in*_⟩_*non−GoE*_). Since ⟨*d*_*in*_⟩_*non−GoE*_ is a non-linear monotone increasing function of *G*-density as derived in Eq.(2), we expect the trends observed in the distribution of *G*-density to be replicated here. That is indeed the case as shown in SI Figs. S7 and S8. More explicitly, the mean values of the distribution of ⟨*d*_*in*_⟩_*non−GoE*_ is larger in ensembles constrained with biologically meaningful logics compared to ensembles constrained with EFs. Again, among ensembles generated with biologically meaningful logic, the STGs in the NCF-ensemble are typically more bushy than those in the EUF-ensemble or RoF-ensemble.

Next, we consider the distribution of the *average convergence rate* of trajectories originating at *GoE* states (*λ*_*GoE*_). From Fig. 4 and SI Fig. S9, we observe that on average, the models in the EF-ensemble have longer transient times (smaller values of *λ*_*GoE*_) in comparison to ensembles that employ biologically meaningful logic rules. We also find that the 3 ensembles constrained with biologically meaningful logic rules have similar values of *λ*_*GoE*_. These trends are consistent across the 10 GRNs as can be seen in Fig. 4 and SI Fig. S9. Wuensche [40] observed that highly bushy and convergent structures imply more ordered dynamics, implying that NCFs lead to more ordered dynamics compared to the other types of BFs. This is indeed consistent with previous studies on the effect of logic rules on the dynamical regimes of the network [19, 45]. Lastly, we have computed the Spearman correlation between *G*-density and *λ*_*GoE*_ for ensembles across the 10 GRNs. The corresponding scatter plots are provided in the SI (see SI Figs. S10 and S11). Although we find that those two measures are positively correlated, the correlation coefficient is surprisingly small, indicating that the two measures are capturing different aspects of the STG.

**FIG. 4.**
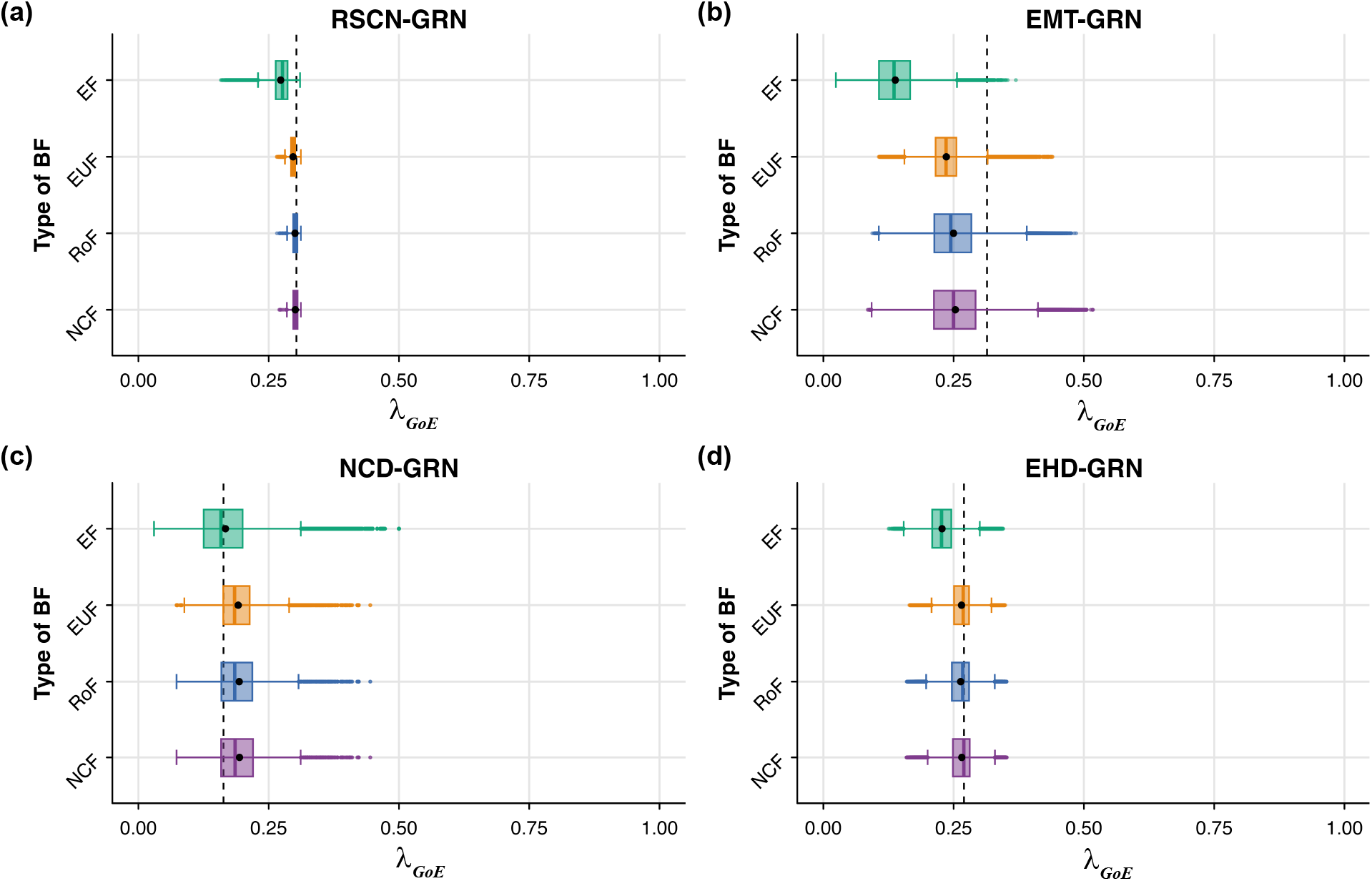
Distribution of average convergence rate of trajectories originating at *GoE* states (*λ*_*GoE*_) for different ensembles generated using network structures from 4 published Boolean GRNs. The subplots **(a), (b), (c)** and **(d)** correspond to the RSCN-GRN, EMT-GRN, NCD-GRN and EHD-GRN network structures respectively. The *x* and *y* axes of each subplot correspond respectively to *λ*_*GoE*_ and to the type of BF imposed on the nodes of the networks. The box plots display the distribution of *λ*_*GoE*_ in the 4 ensembles that each use a given type of regulatory logic. The ensembles generated using biologically meaningful BFs (EUFs, RoFs, NCFs) have higher convergence rates compared to the ensemble generated using EFs. Note that we impose that ensembles recover the biological fixed points of the associated published model. The mean and median of the distribution are indicated by the black dot and the vertical line within the box respectively. The vertical dashed lines in each subplot correspond to the *λ*_*GoE*_ of the Boolean model provided by the modelers in the published article.

### 3.2 Biologically meaningful functions have lower local dynamics descriptor values

In section 2.4, we presented our approach to adapt the *Z*-parameter defined in CA to regulatory logic rules (BFs) in BNs. This is illustrated via the schematic provided in Fig. 2. Briefly, given a BF, we first generate all of its permutations (see left panel, Fig. 2(d)) and compute their *Z*_*left*_ values using the method given in Fig. 2(b-c) (see section 2.4.1 for details). From this collection of *Z*_*left*_ values, we define 4 potential variants for the *Z*-parameter, namely, *Z*_*max*_, *Z*_*min*_, *Z*_*ave*_, *Z*_*mid*_ as explained in section 2.4.2. To obtain the network-level *Z*-parameters of a given BN (hereafter referred to as the network *Z*-parameters), we average each of these different *Z*-parameters over all nodes of the network to get 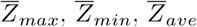 and 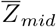.

We begin with the distributions of 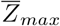 (see SI Figs. S12 and S13) and 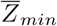 (see SI Figs. S14 and S15). We find that the ensembles generated with biologically meaningful functions have lower 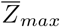 and 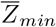 compared to the ensemble generated with EFs. This trend is consistent across all 10 reconstructed GRNs. Suppose we now restrict to the ensembles generated with biologically meaningful regulatory logic. First, we find that 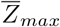 does not show any particular trend across the 10 GRNs. Second, 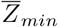 is minimum for NCFs, followed by RoFs and then EUFs across the 10 GRNs. In particular, the 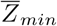 for the NCF-ensembles across all GRNs have zero variance. This is because *Z*_*min*_ for NCFs depends only on the number of inputs *k* of a BF (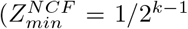, see the SI text, section 8 for a proof). Since all models in a given NCF-ensemble have the same network structure and thus in-degrees, their 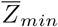 values are necessarily identical.

Next, we consider the distribution of 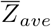 (see SI Figs. S16 and S17) and 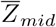 (see Fig. 5 and SI Fig. 18). We find that the ensembles generated with biologically meaningful functions have lower 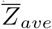 and 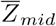 values compared to the ensemble generated with EFs. Among ensembles generated with biologically meaningful regulatory logic, 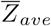 does not show a consistent trend across the 10 GRNs, whereas 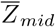 consistently achieves its lowest values in the NCF-ensemble, across all 10 GRNs.

**FIG. 5.**
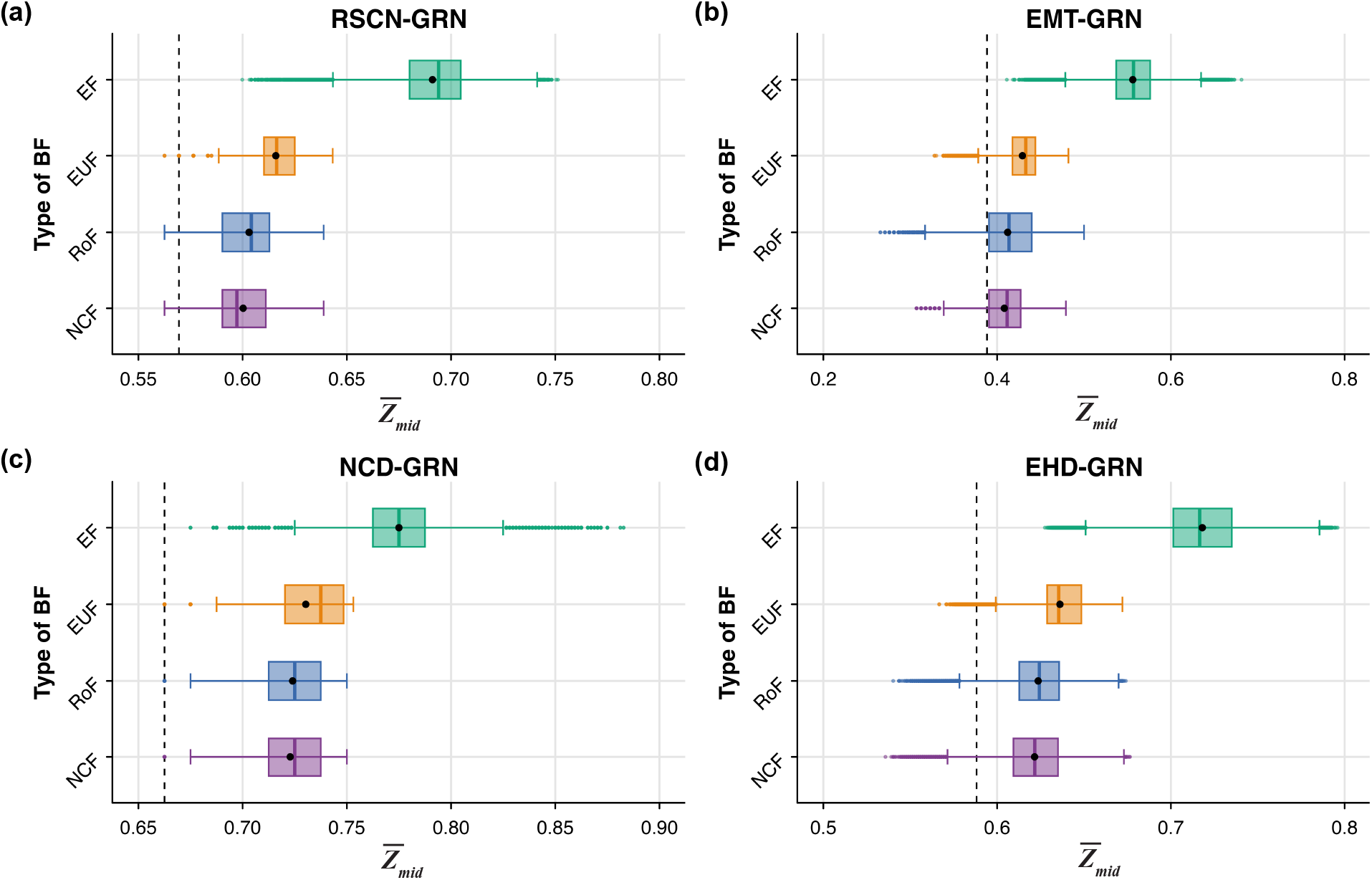
Distribution of 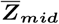 values for different ensembles generated using network structures from 4 published Boolean GRNs. The subplots **(a), (b), (c)** and **(d)** correspond to the RSCN-GRN, EMT-GRN, NCD-GRN and EHD-GRN network structures respectively. The *x* and *y* axes of each subplot correspond respectively to 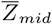 and to the type of BF imposed on the nodes of the networks. The box plots display the distribution of 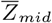 in the 4 ensembles that each use a given type of regulatory logic. The ensembles generated using biologically meaningful BFs (EUFs, RoFs, NCFs) have lower 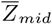 compared to the ensemble generated using EFs. Note that we impose that ensembles recover the biological fixed points of the associated published model. The mean and median of the distribution are indicated by the black dot and the vertical line within the box respectively. The vertical dashed lines in each subplot correspond to the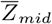 of the Boolean model provided by the modelers in the published article.

Our last descriptor of local dynamics is the network sensitivity, 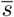 (see Fig. 6 and SI Fig. S19). Some of us have previously shown that NCFs possess the minimum average sensitivity across all odd values of bias for any given number of inputs [19] and hence it is expected that the NCFs should have the lowest values of 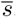. That is indeed the case as can be seen from the box plots in Fig. 6 and SI Fig. S19. Lastly, we investigated the correlations between the different descriptors of local dynamics namely, 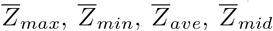 and 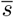. The heat maps displayed in SI Figs. S20 and S21 also provide the Spearman correlation coefficients between all pairs of descriptors. The general trend observed is that most pairs of descriptors show moderate to very strong positive correlation across the various GRNs. This is illustrated in particular via our scatter plots between 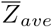 and 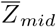 (see SI Figs. S22 and S23), and *s* and *Z*_*mid*_ (see SI Figs. S24 and S25). In the following sub-section, we explore which of these 5 descriptors of local dynamics can serve as good predictors of bushiness or convergence.

**FIG. 6.**
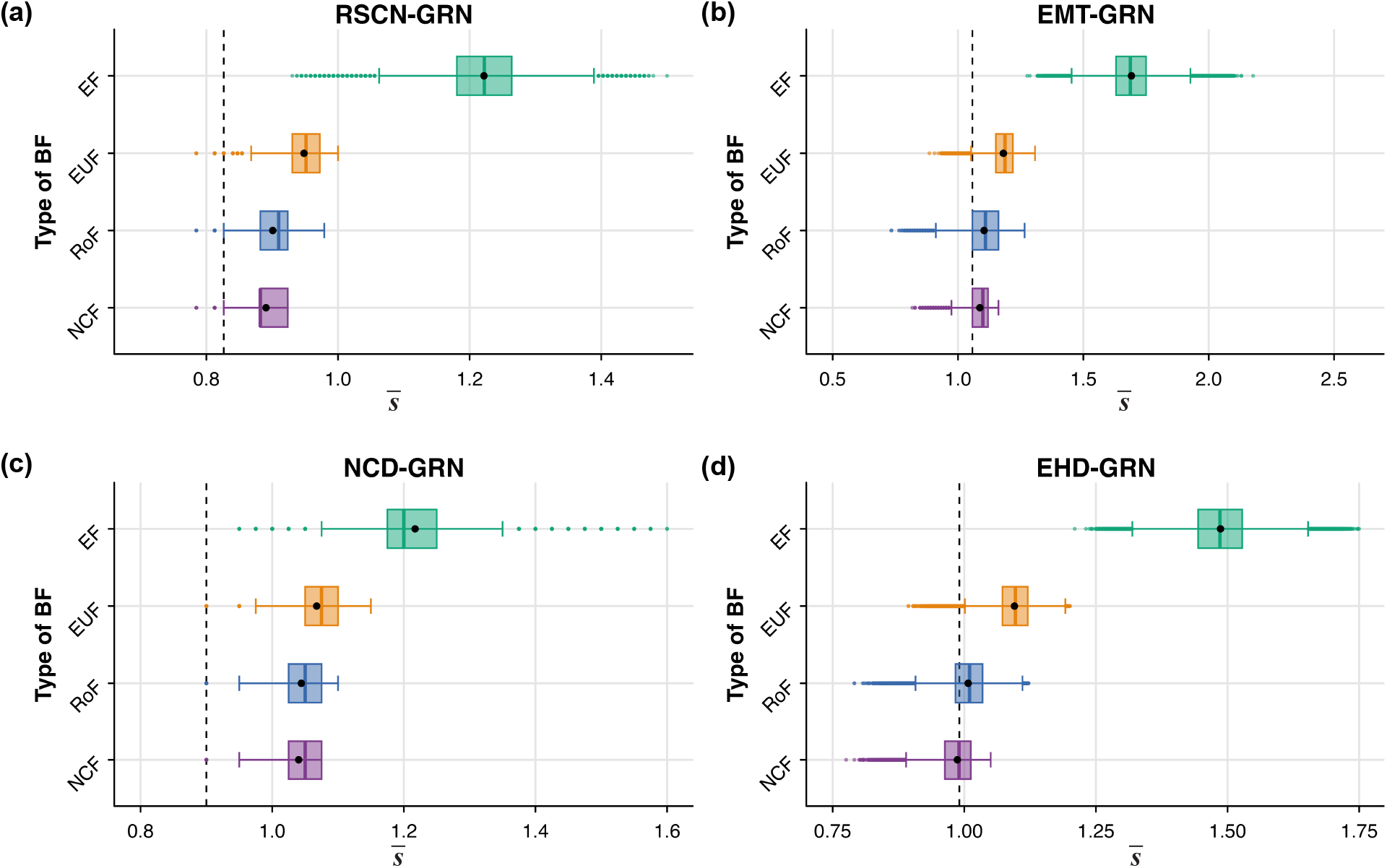
Distribution of network sensitivity (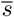) values for different ensembles generated using network structures from 4 published Boolean GRNs. The subplots **(a), (b), (c)** and **(d)** correspond to the RSCN-GRN, EMT-GRN, NCD-GRN and EHD-GRN network structures respectively. The *x* and *y* axes of each subplot correspond respectively to 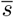 and to the type of BF imposed on the nodes of the networks. The box plots display the distribution of 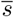 in the 4 ensembles that each use a given type of regulatory logic. The ensembles generated using biologically meaningful BFs (EUFs, RoFs, NCFs) have lower 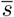 compared to the ensemble generated using EFs. Note that we impose that ensembles recover the biological fixed points of the associated published model. The mean and median of the distribution are indicated by the black dot and the vertical line within the box respectively. The vertical dashed lines in each subplot correspond to the 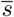 of the Boolean model provided by the modelers in the published article.

### 3.3 Descriptors of local dynamics can serve as good predictors of bushiness of state transition graphs

Wuensche showed that, in CA, the *Z*-parameter exhibits a ‘marked’ correlation with the *G*-density [38]. That suggests that it may be possible to infer at least qualitative information about the bushiness of the STG of a CA from properties of its rule table. Extending this question to the BN framework, we ask whether descriptors of local dynamics (in fact, characteristics of the rule tables for each node) can serve as good predictors of bushiness of such a model’s STG. The distributions obtained in the previous two sub-sections for the measures of bushiness and descriptors of local dynamics provide a hint that some of these descriptors could serve as proxies for the global measures of bushiness.

To test if this is indeed the case, we computed the Spearman correlation coefficient of the bushiness measure, *G*-density, with the different descriptors of local dynamics. These correlation coefficients are given in the heatmaps in Fig. 7 and SI Fig. S26. From these figures it is clear that for most GRNs, the 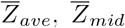 and *s* are more strongly correlated with the bushiness measure, *G*-density, than 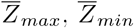 are. Since those 3 descriptors of local dynamics show a moderate to strong negative correlation with the *G*-density, they can be useful predictors of the bushiness of a model’s STG. Note that since ⟨*d*_*in*_⟩_*non−GoE*_ is a monotone increasing function of the *G*-density, the Spearman correlation coefficients of ⟨*d*_*in*_⟩_*non−GoE*_ with the 5 descriptors will be the same as that of *G*-density. Hence ⟨*d*_*in*_⟩_*non−GoE*_ is not shown in the heat maps. The scatter plots of *G*-density with each of the 5 descriptors of local dynamics are provided in the 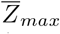 (SI Figs. S27 and S28), 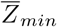 (SI Figs. S29 and S30), 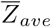 (SI Figs. S31 and S32), 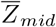 (SI Figs. S33 and S34) and *s* (SI Figs. S35 and S36).

An implication of these correlations is that for ensembles of larger networks (where generating the STG is computationally infeasible), we may simply use the descriptors of local dynamics such as 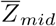 and 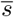 to predict which models will have a more bushy STG.

**FIG. 7.**
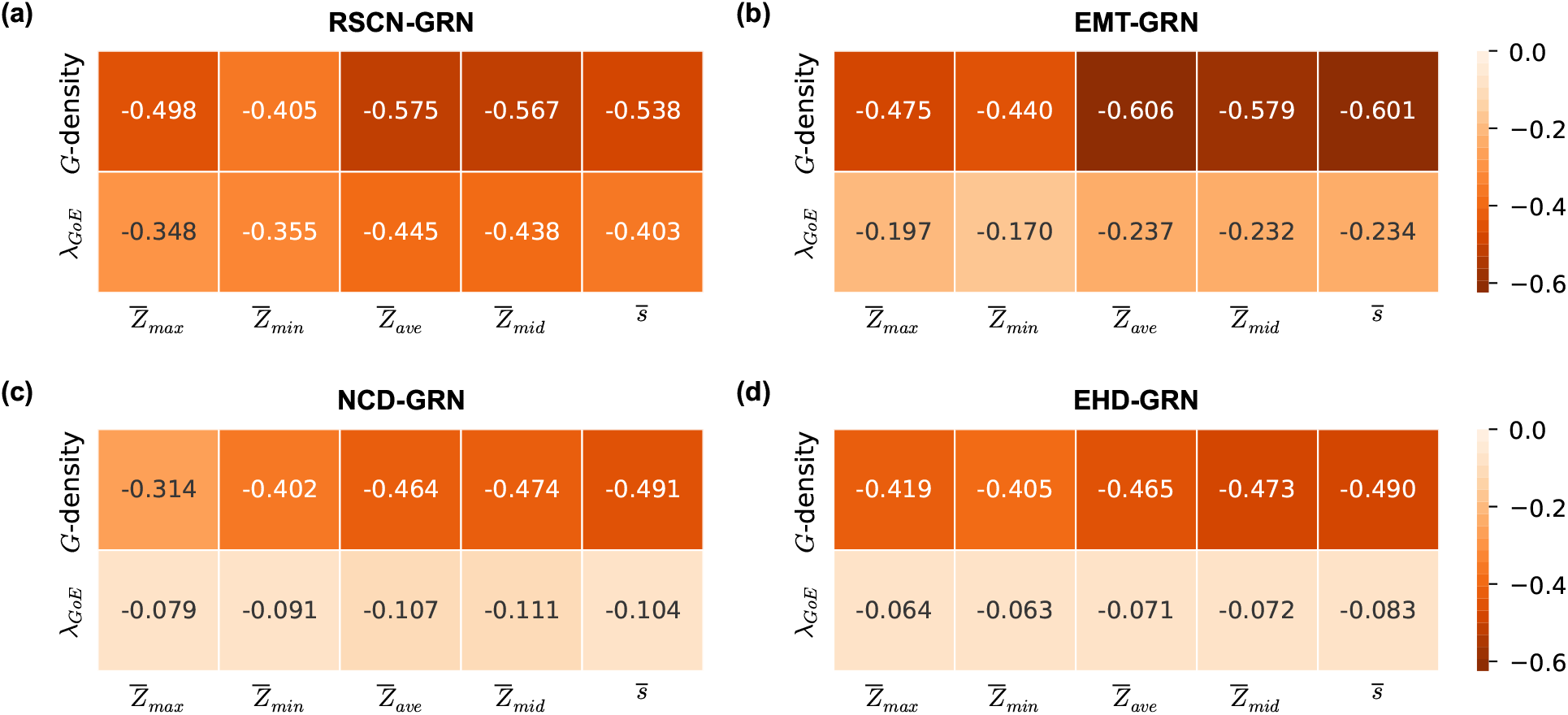
Correlation heat map between descriptors of local dynamics and global measures (bushiness and convergence rate) for ensembles generated using network structures from 4 published Boolean GRNs. The subplots **(a), (b), (c)** and **(d)** correspond to the RSCN-GRN, EMT-GRN, NCD-GRN and EHD-GRN network structures respectively. The EF-ensemble for each network structure was used to generate the heat maps. The rows of each subplot correspond to quantities computed on the state transition graph, namely *G*-density and average convergence rate of trajectories originating at *GoE* states (*λ*_*GoE*_). The columns of each subplot correspond to descriptors of local dynamics, namely, the network *Z*-parameters 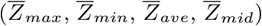 and network sensitivity 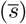. The heat maps show the pair-wise Spearman correlations between descriptors of local dynamics and global measures of bushiness and convergence rate defined on the state transition graph. The descriptors of local dynamics show a moderate to strong negative correlations with *G*-density and very weak to moderate negative correlations with *λ*_*GoE*_.

### 3.4 Descriptors of local dynamics are not good predictors of convergence rate in state transition graphs

In direct analogy with the case of *G*-density, we consider here the correlation coefficients of average convergence rate (*λ*_*GoE*_) with the descriptors of local dynamics. The results are given as before in the heatmaps of Fig. 7 and SI Fig. S26 which allows one to compare the two cases. Though both lead to negative correlation coefficients, for the majority of the ensembles the correlations are very much weaker when using *λ*_*GoE*_ than when using *G*-density. As a result, it is not appropriate to use any of these descriptors of local dynamics for predicting *λ*_*GoE*_, even qualitatively. Nevertheless, one trend that is in common between the current situation and that when considering *G*-density is that for most GRNs, the 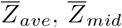 and 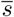 show stronger correlations than 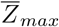 and 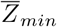. As was done for *G*-density, we provide in the SI the scatter plots between the *λ*_*GoE*_ and each of the 5 descriptors of local dynamics: 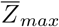 (SI Figs. S37 and S38), 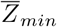 (SI Figs. S39 and S40), 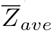 (SI Figs. S41 and S42), 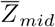 (SI Figs. S43 and S44) and 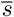 (SI Figs. S45 and S46).

### 3.5 The average convergence rate of trajectories originating at random states is a good proxy for the average convergence rate of trajectories originating at *GoE* states

Given the fairly poor correlation between the descriptors of local dynamics and *λ*_*GoE*_, we propose *λ*_*random*_, the average convergence rate of trajectories originating at random states, as an alternative to *λ*_*GoE*_. First we find that *λ*_*GoE*_ is very strongly correlated with *λ*_*all*_ which is the average convergence rate over all states in the STG (SI Figs. S47 and S48) with a Spearman correlation coefficient that approaches 1 in the limit of large sample sizes. This provides an impetus to ask whether trajectories originating at randomly sampled states may be a good proxy for *λ*_*GoE*_, because if so, one could use it as a proxy and no longer have to compute the STG or find the *GoE* states. SI Figs. S49 and S50 reveal that *λ*_*random*_ can indeed serve as an excellent predictor of *λ*_*GoE*_.

## 4. DISCUSSION AND CONCLUSIONS

In this work, we systematically studied how the choice of a type of BF, given a network structure, influences two features of the STG, namely bushiness and convergence, and how certain descriptors of local dynamics might serve as predictors or proxies of those features.

First, we find that biologically meaningful regulatory logic, namely, EUFs, RoFs and NCFs lead to more bushy and convergent STGs, as measured via the *G*-density and the average convergence rate of trajectories originating at *GoE* states (*λ*_*GoE*_), compared to EFs (essentially equivalent to random BFs) across 10 reconstructed GRNs. Furthermore, among the biologically meaningful regulatory logic rules, the NCFs typically lead to a more bushy and convergent STG compared to other types of BFs. Second, we observe that *G*-density correlates only very weakly with *λ*_*GoE*_, a counter-intuitive result that also goes against previous expectations [40]. Third, we generalized Wuensche’s *Z*-parameter to BNs leading to the 4 network *Z*-parameters 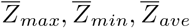 and 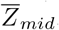. Examining their distributions for various ensembles reveals that ensembles employing biologically meaningful regulatory logic rules have lower values of the 4 proposed parameters than ensembles employing just EFs, across all 10 reconstructed GRNs. In addition, among the biologically meaningful regulatory logic rules, the NCFs consistently led to the lowest values for *Z*_*mid*_ across all 10 reconstructed GRNs. The network sensitivity *s* showed exactly the same trend, which is expected given previous observations [19]. Fourth, among the descriptors of local dynamics, we find that 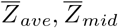 and 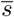 show very similar correlations with the global measures and furthermore these correlations are significantly stronger than those of 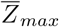 and 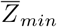.

The ‘bushiness’ of STGs has been well studied and characterized in CA. Wuensche showed that the signature of highly ordered or convergent dynamics is the presence of short, highly branching transient trees, with a high proportion of leaves in the STG, whereas the signature of chaotic dynamics is long transients, low branching in the STG and a low proportion of leaves therein [38, 40, 49]. These observations and results are limited primarily to 1D CA in which both network architecture and update rules are regular and homogeneous respectively. Such studies are absent in generalized automata that mimic biological systems, namely, BNs - which generally lack a regular network architecture and employ heterogenous rules across their nodes. Though dynamical regimes of biological BNs based on the type of BF employed have been explored [19, 45], a study that characterizes those regimes directly based on the features of the STG has been lacking. In particular, the effect of different types of rules on the bushiness or convergence of the STG under several constraints that are biologically meaningful has not been studied so far. This work is a first such study of the bushiness and convergence of STGs for different reconstructed GRNs using *G*-density and *λ*_*GoE*_. We have found that biologically meaningful BFs lead to more bushy and convergent STGs compared to random EFs.

To the best of our knowledge, the relationship between the 2 measures, namely, *G*-density and *λ*_*GoE*_, has not been explored previously. Contrary to our initial expectations that more bushy STGs result in higher convergence, we were surprised to find that these measures are only weakly correlated. A posteriori, it is easy to convince oneself that the two measures are indeed capturing different features. For example, consider a rooted tree of constant connectivity *K* for which all the leaves are at the same distance *H* from the root. The bushiness, that is *G*-density, is close to *K/*(*K* − 1) while the transient times are all equal to *H*, leading to a convergence rate of 1*/H*. Thus bushiness and convergence rate can be varied essentially independently. Another example closer to the STGs we have examined so far consists of a bush that is tethered to a line of nodes; in such a situation, the measure of bushiness varies little when the node associated with the fixed point is changed whereas the average transient length will be quite sensitive to that choice.

In our study we were able to use exhaustive enumeration to identify the garden-of-Eden states (GoEs) in all of our models. Computationally, such an approach requires on the order of 2^*N*^ operations for *N* genes and thus it is not appropriate for larger networks. Unfortunately the computational complexity of identifying the garden-of-Eden states in BNs is a #**P**-complete problem, meaning that it is at least as hard as a NP-complete problem to compute the *G*-density of a general BN [48]. Thus, for large networks it is necessary to introduce proxies that can bypass this computational complexity to give us information about the bushiness of the STG. Wuensche introduced his *Z*-parameter in CA as the probability of pre-image determination (identifying predecessors to the CA dynamics in a sequential procedure) [49, 50] and found that it showed a ‘marked’ correlation with the *G*-density [50]. We thus extended his *Z*-parameter to generalizations applicable to BNs and found good correlations with *G*-density, allowing them to be used as proxies for *G*-density. Among all the network *Z*-parameters we explored, the choice 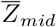 was typically the one with the highest correlation with *G*-density. Furthermore, its value was consistently at its minimum across all BFs when focusing on NCFs, across all 10 GRNs. Of particular interest is the fact that for NCFs, 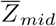 can be computed extremely quickly.

We found that the network sensitivity 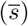 is also a good predictor of *G*-density. Previous observations had shown that NCFs lead to more ordered dynamics compared to other types of BFs such as EUFs and EFs [19]. This can be justified by the fact that NCFs achieve the minimum average sensitivity over all possible BFs [19], thereby leading to lower values of 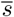 which is associated with a greater bushiness of the STG. Note that all these descriptors are defined at the level of individual nodes and are ‘unaware’ of the network architecture (which is not an issue in CA). As discussed in the results, since the average transient length correlates only very weakly with the descriptors of local dynamics, we required a good estimator of the same without having to generate the entire STG or finding its *GoE* states. So we tested whether the average convergence rate of the trajectories of randomly sampled states from the STG (*λ*_*random*_) might serve as a proxy for *λ*_*GoE*_. We found that it is in fact an excellent proxy. In sum, we provide 3 proxies: 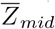 and 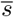 which inform us about the ‘bushiness’, and *λ*_*random*_ which informs us about the ‘convergence’.

Notwithstanding the developments of new methods and results, there are some limitations that arise in our work which we would like to address in the future. (i) Our proposed sampling algorithm for EUFs for sampling large inputs (*k* = 7 and beyond) is not perfectly uniform. Although the associated bias is small and thus does not affect our conclusions (most of the networks studied have nodes with at most 6 inputs), it certainly will strengthen the conclusions for networks having nodes with more than 6 inputs. Furthermore, it will be a useful tool for the community at large to be able to generate random EUFs that satisfy predefined constraints on interaction signs and fixed points. (ii) It is computationally expensive to compute the *Z*-parameter for nodes with a large number of inputs such as 7 or 8 because of the *k*! permutations of the inputs that have to be studied. Hence faster algorithms to accomplish this are necessary if we are to speed up the computation of *Z*-parameters. (iii) The structure of STG generated under asynchronous update is different from the STG generated under synchronous update; in particular, under asynchronous updates, states may have more than one out-degree. Thus, it can be expected that the number of GoE states under asynchronous update is almost always less than what is obtained in the synchronous case. Can one define a *G*-density that is computationally tractable in that situation? Of course this question also applies to our measure *λ*_*GoE*_. (iv) Finally, although we provide an excellent proxy for *λ*_*GoE*_, that proxy depends not only on the BFs, the actual dynamics in the network are required. Hence the question of whether a good proxy of *λ*_*GoE*_ exists solely at the level of the BFs remains open.

In conclusion, we have systematically characterized how the structural features of the STG of Boolean GRNs vary depending on the type of logic rule employed by drawing inspiration from the rich theory of CA which likely can provide even further insights into understanding dynamics of BNs.

## DATA AND CODE AVAILABILITY

All data and codes needed to reproduce the results in this manuscript are deposited in GitHub and are available at: https://github.com/asamallab/BushySTG.

## ACKNOWLEDGMENTS

The authors acknowledge the use of the supercomputing machine Nandadevi at The Institute of Mathematical Sciences, Chennai. Priyotosh Sil would like to thank Velmurugan S for discussions. Ajay Subbaroyan would like to thank Ajaya Kumar Sahoo for discussions. Areejit Samal acknowledges support from the Max Planck Society, Germany, through the award of a Max Planck Partner Group in Mathematical Biology, and the Department of Atomic Energy, Government of India. IPS2 benefits from the support of Saclay Plant Sciences-SPS (ANR-17-EUR-0007).

## AUTHORS’ CONTRIBUTIONS

Designed the research: P.S., Aj.S., O.C.M, Ar.S.; Performed the research: P.S., Aj.S., S.K., O.C.M, Ar.S.; Performed the computations: P.S., Aj.S., S.K.; Wrote the paper: P.S., Aj.S., O.C.M, Ar.S.

## References

[1] S. A. Kauffman. The origins of order: self-organization and selection in evolution. Oxford University Press, New York, 1993.

[2] U. Alon. An Introduction to Systems Biology: Design Principles of Biological Circuits. Chapman and Hall/CRC., 2006.

[3] A. L. Barabási and Z. N. Oltvai. Network biology: understanding the cell’s functional organization. Nature Reviews Genetics, 5(2):101–113, 2004.

[4] B. ø. Palsson. Systems Biology: Properties of Reconstructed Networks. Cambridge University Press, 2006.

[5] S. Camazine, J.-L. Deneubourg, N. R. Franks, J. Sneyd, G. Theraula, and E. Bonabeau. Self-Organization in Biological Systems. Princeton University Press, Princeton, 2001.

[6] H. Kitano. Computational systems biology. Nature, 420(6912):206–210, 2002.

[7] H. de Jong. Modeling and simulation of genetic regulatory systems: a literature review. Journal of Computational Biology, 9(1):67–103, 2002.

[8] K. Kaneko. Life: An Introduction to Complex Systems Biology. Understanding Complex Systems. Springer Berlin Heidelberg, 2006.

[9] J. Von Neumann and A. W. Burks. Theory of self-reproducing automata. IEEE Transactions on Neural Networks, 5(1):3–14, 1966.

[10] S. Wolfram. Statistical mechanics of cellular automata. Reviews of Modern Physics, 55(3):601–644, 1983.

[11] C. G. Langton. Studying artificial life with cellular automata. Physica D: Nonlinear Phenomena, 22(1-3):120–149, 1986.

[12] G. B. Ermentrout and L. Edelstein-Keshet. Cellular Automata Approaches to Biological Modeling. Journal of Theoretical Biology, 160(1):97–133, 1993.

[13] O. Martin, A. M. Odlyzko, and S. Wolfram. Algebraic properties of cellular automata. Communications in Mathematical Physics, 93(2):219–258, 1984.

[14] A. Wuensche and M. Lesser. Global dynamics of cellular automata: an atlas of basin of attraction fields of one-dimensional cellular automata. Addison-Wesley, Reading, MA, 1992.

[15] R. A. Brooks and P. Maes. Artificial Life IV: Proceedings of the Fourth International Workshop on the Synthesis and Simulation of Living Systems. The MIT Press, 1994.

[16] H. S. Mortveit and C. M. Reidys. An Introduction to Sequential Dynamical Systems. Springer-Verlag, Berlin, Heidelberg, 2007.

[17] S. A. Kauffman. Metabolic stability and epigenesis in randomly constructed genetic nets. Journal of Theoretical Biology, 22(3):437–467, 1969.

[18] S. A. Kauffman. Homeostasis and differentiation in random genetic control networks. Nature, 224(5215):177–178, 1969.

[19] A. Subbaroyan, O. C. Martin, and A. Samal. Minimum complexity drives regulatory logic in Boolean models of living systems. PNAS Nexus, 1(1):pgac017, 2022.

[20] R. Albert and H. G. Othmer. The topology of the regulatory interactions predicts the expression pattern of the segment polarity genes in Drosophila melanogaster. Journal of Theoretical Biology, 223(1):1–18, 2003.

[21] S. A. Kauffman, C. Peterson, B. Samuelsson, and C. Troein. Random Boolean network models and the yeast transcriptional network. Proceedings of the National Academy of Sciences, 100(25):14796–14799, 2003.

[22] L. Mendoza and I. Xenarios. A method for the generation of standardized qualitative dynamical systems of regulatory networks. Theoretical Biology and Medical Modelling, 3(1):13, 2006.

[23] A. Samal and S. Jain. The regulatory network of E. coli metabolism as a Boolean dynamical system exhibits both homeostasis and flexibility of response. BMC Systems Biology, 2(21):1–18, 2008.

[24] M. I. Davidich and S. Bornholdt. Boolean Network Model Predicts Cell Cycle Sequence of Fission Yeast. PLoS ONE, 3(2):1–8, 2008.

[25] L. Calzone, L. Tournier, S. Fourquet, D. Thieffry, B. Zhivotovsky, E. Barillot, and A. Zinovyev. Mathematical modelling of cell-fate decision in response to death receptor engagement. PLoS Computational Biology, 6(3):e1000702, 2010.

[26] J. Krumsiek, C. Marr, T. Schroeder, and F. J. Theis. Hierarchical Differentiation of Myeloid Progenitors Is Encoded in the Transcription Factor Network. PLoS ONE, 6(8):e22649, 2011.

[27] A. Saadatpour, R. S. Wang, A. Liao, X. Liu, T. P. Loughran, I. Albert, and R. Albert. Dynamical and structural analysis of a T cell survival network identifies novel candidate therapeutic targets for large granular lymphocyte leukemia. PLoS Computational Biology, 7(11):e1002267, 2011.

[28] M. L. García-Gomez, E. Azpeitia, and E. R. Álvarez Buylla. A dynamic genetic-hormonal regulatory network model explains multiple cellular behaviors of the root apical meristem of Arabidopsis thaliana. PLoS Computational Biology, 13(4):1–36, 2017.

[29] S. Huang and D. E. Ingber. Shape-dependent control of Cell Growth, Differentiation, and Apoptosis: Switching between Attractors in Cell Regulatory Networks. Experimental Cell Research, 261(1):91–103, 2000.

[30] S. Huang, G. Eichler, Y. Bar-Yam, and D. E. Ingber. Cell fates as high-dimensional attractor states of a complex gene regulatory network. Physical Review Letters, 94(12):128701, 2005.

[31] S. Wolfram. A New Kind of Science. Wolfram Media, 2002.

[32] Maximino Aldana. Boolean dynamics of networks with scale-free topology. Physica D: Nonlinear Phenomena, 185(1):45–66, 2003.

[33] R. Serra, M. Villani, A. Barbieri, S.A. Kauffman, and A. Colacci. On the dynamics of random Boolean networks subject to noise: Attractors, ergodic sets and cell types. Journal of Theoretical Biology, 265(2):185–193, 2010.

[34] A. Henry, F. Monéger, A. Samal, and O. C. Martin. Network function shapes network structure: the case of the Arabidopsis flower organ specification genetic network. Molecular BioSystems, 9(7):1726–1735, 2013.

[35] J. G. T. Zañudo and R. Albert. An effective network reduction approach to find the dynamical repertoire of discrete dynamic networks. Chaos: An Interdisciplinary Journal of Nonlinear Science, 23(2):025111, 2013.

[36] A. J. Gates, R. B. Correia, X. Wang, and L. M. Rocha. The effective graph reveals redundancy, canalization, and control pathways in biochemical regulation and signaling. Proceedings of the National Academy of Sciences, 118(12), 2021.

[37] A. Wuensche. The Ghost in the Machine: Basins of Attraction of Random Boolean Networks. volume 17, pages 465–501. Addison-Wesley, 1994.

[38] A. Wuensche. Complexity in one-D cellular automata: Gliders, basins of attraction and the Z parameter. Working Paper 94-04-025, Santa Fe Institute, 1994.

[39] A. Wuensche. Attractor basins of discrete networks. D.Phil, The University of Sussex, 1997.

[40] A. Wuensche. Basins of attraction in network dynamics: A conceptual framework for biomolecular networks. In Modularity in Development and Evolution, pages 288–311. Chicago University Press, 2004.

[41] B. Derrida and Y. Pomeau. Random networks of automata: a simple annealed approximation. Europhysics Letters, 1(2):45, 1986.

[42] B. Drossel, T. Mihaljev, and F. Greil. Number and length of attractors in a critical Kauffman model with connectivity one. Physical Review Letters, 94(8):088701, 2005.

[43] K. Klemm and S. Bornholdt. Stable and unstable attractors in Boolean networks. Physical Review E, 72(5):055101, 2005.

[44] A. Roli, M. Villani, A. Filisetti, and R. Serra. Dynamical Criticality: Overview and Open Questions. Journal of Systems Science and Complexity, 31(3):647–663, 2018.

[45] B. C. Daniels, H. Kim, D. Moore, S. Zhou, H. B. Smith, B. Karas, S. A. Kauffman, and S. I. Walker. Criticality Distinguishes the Ensemble of Biological Regulatory Networks. Physical Review Letters, 121(13):138102, 2018.

[46] A. Wuensche. Discrete dynamical networks and their attractor basins. Complex systems, 98:3–21, 1998.

[47] C. Gershenson. Guiding the self-organization of random Boolean networks. Theory in Biosciences, 131(3):181–191, 2012.

[48] P. T. Tošić and C. Ordonez. Boolean Network Models of Collective Dynamics of Open and Closed Large-Scale Multi-agent Systems. In V. Mařík, W. Wahlster, T. Strasser, and P. Kadera, editors, Industrial Applications of Holonic and Multi-Agent Systems, pages 95–110, Cham, 2017. Springer International Publishing.

[49] A. Wuensche. Classifying cellular automata automatically: Finding gliders, filtering, and relating space-time patterns, attractor basins, and the Z parameter. Complexity, 4(3):47–66, 1999.

[50] A. Wuensche. CELLULAR AUTOMATA ENCRYPTION: the reverse algorithm, Z-parameter and chain-rules. Parallel Processing Letters, 19(2):283–297, 2009.

[51] R. Thomas. Kinetic logic: a Boolean approach to the analysis of complex regulatory systems, Proceedings of the EMBO course “Formal analysis of genetic regulation”, held in Brussels, September 6–16, 1977, Lecture notes in Biomathematics. Springer-Verlag, 1979.

[52] J. Aracena. Maximum Number of Fixed Points in Regulatory Boolean Networks. Bulletin of Mathematical Biology, 70(5):1398, 2008.

[53] J. X. Zhou, A. Samal, A. F. d’Hérouël, N. D. Price, and S. Huang. Relative stability of network states in Boolean network models of gene regulation in development. Biosystems, 142-143:15–24, 2016.

[54] A. Subbaroyan, P. Sil, O. C. Martin, and A. Samal. Leveraging developmental landscapes for model selection in Boolean gene regulatory networks. Briefings in Bioinformatics, 24(3):bbad160, 2023.

[55] C. Müssel, M. Hopfensitz, and H. A. Kestler. BoolNet—an R package for generation, reconstruction and analysis of Boolean networks. Bioinformatics, 26(10):1378–1380, 2010.

[56] E. Azpeitia, M. Benítez, I. Vega, C. Villarreal, and E. R. Alvarez-Buylla. Single-cell and coupled GRN models of cell patterning in the Arabidopsis thaliana root stem cell niche. BMC Systems Biology, 4(1):134, 2010.

[57] E. Sullivan, M. Harris, A. Bhatnagar, E. Guberman, I. Zonfa, and E. Ravasz Regan. Boolean modeling of mechanosensitive epithelial to mesenchymal transition and its reversal. iScience, 26(4):106321, 2023.

[58] C. E. Giacomantonio and G. J. Goodhill. A Boolean Model of the Gene Regulatory Network Underlying Mammalian Cortical Area Development. PLoS Computational Biology, 6(9):e1000936, 2010.

[59] F. Herrmann, A. Groß, D. Zhou, H. A. Kestler, and M. Kühl. A Boolean Model of the Cardiac Gene Regulatory Network Determining First and Second Heart Field Identity. PLoS ONE, 7(10):e46798, 2012.

[60] C. Biane and F. Delaplace. Causal reasoning on boolean control networks based on abduction: theory and application to cancer drug discovery. IEEE/ACM Transactions on Computational Biology and Bioinformatics, 16(5):1574–1585, 2019.

[61] J. E. Narv áez-Chavez, E. R. Álvarez Buylla, and J. C. Martínez-García. Uncovering the role of mutations in Epithelial-to-Mesenchymal transition through computational analysis of the underlying gene regulatory network. In Hisham Al-Mubaid, Tamer Aldwairi, and Oliver Eulenstein, editors, Proceedings of International Conference on Bioinformatics and Computational Biology (BICOB-2023), volume 92, pages 92–101. EasyChair, 2023.

[62] Y. S ánchez-Corrales, E. R. Aĺvarez Buylla, and L. Mendoza. The Arabidopsis thaliana flower organ specification gene regulatory network determines a robust differentiation process. Journal of Theoretical Biology, 264(3):971–983, 2010.

[63] O. R íos, S. Frias, A. Rodr íguez, S. Kofman, H. Merchant, L. Torres, and L. Mendoza. A Boolean network model of human gonadal sex determination. Theoretical Biology and Medical Modelling, 12(1):26, 2015.

[64] A. Wuensche. Complex and chaotic dynamics, basins of attraction, and memory in discrete networks. In Acta Physica Polonica. Series B, Proceedings Supplement, volume 3, pages 463–478, 2009.

[65] I. Shmulevich and S. A. Kauffman. Activities and Sensitivities in Boolean Network Models. Physical Review Letters, 93(4):48701, 2004.

