## Supplementary material for "Biologically meaningful regulatory logic enhances the convergence rate in Boolean networks and bushiness of their state transition graph": SI

### 1. BACKGROUND ON BOOLEAN FUNCTIONS

Mathematically, a Boolean function (BF) can be represented as a mapping:

$$f : \{0, 1\}^k \rightarrow \{0, 1\}$$

where $\{0, 1\}$ is known as the Boolean domain and $k$ is the number of inputs of the function $f$. Every $k$ input BF can be represented by a propositional expression of its $k$ variables $x_1, x_2, \ldots$ and $x_k$. BFs can also be specified via a truth table having $k$ input columns and one output column. There, we use the following convention for ordering the truth table throughout this manuscript. If $x_1, x_2, ..., x_k$ are the inputs to the BF $f$, then that same order (from left to right) specifies the variables appearing in the successive columns of the truth table. We also use an integer representation for each rule. To do so, we read the output column of the truth table as a string ordered from left (top) to right (bottom). Thus, the leftmost (most significant) bit corresponds to the output of the first row (all input bits equal to 0) and the rightmost (least significant) bit corresponds to the output of the last row (all input bits equal to 1). The integer equivalent of the binary string is then the (integer) encoding of the BF.

#### Permutations of BFs

Given a $k$-input BF, one can apply a permutation of the $k$ input variables in its propositional formula to produce a modified BF. There are $k!$ such permutations. Consider for instance the 3-input BF $f = x_1 \wedge (\overline{x}_2 \vee x_3)$. The $3! = 6$ permutations on its inputs correspond to $(x_1, x_2, x_3), (x_1, x_3, x_2), (x_2, x_1, x_3), (x_2, x_3, x_1), (x_3, x_1, x_2)$ and $(x_3, x_2, x_1)$. The associated permutations on the BFs are $f_1 = x_1 \wedge (\overline{x}_2 \vee x_3)$, $f_2 = x_1 \wedge (\overline{x}_3 \vee x_2)$, $f_3 = x_2 \wedge (\overline{x}_1 \vee x_3)$, $f_4 = x_2 \wedge (\overline{x}_3 \vee x_1)$, $f_5 = x_3 \wedge (\overline{x}_1 \vee x_2)$ and $f_6 = x_3 \wedge (\overline{x}_2 \vee x_1)$. Note that in this particular example, the 6 permutations lead to 6 distinct BFs, hereafter referred to as non-equivalent BFs. But that is not always the case. As an example, consider the permutations of the BF $f = x_1 \wedge (x_2 \vee x_3)$: $f_1 = x_1 \wedge (x_2 \vee x_3)$, $f_2 = x_1 \wedge (x_3 \vee x_2)$, $f_3 = x_2 \wedge (x_1 \vee x_3)$, $f_4 = x_2 \wedge (x_3 \vee x_1)$, $f_5 = x_3 \wedge (x_1 \vee x_2)$ and $f_6 = x_3 \wedge (x_2 \vee x_1)$. Here, there are only 3 non-equivalent permuted BFs because $f_1 = f_2, f_3 = f_4$ and $f_5 = f_6$. The operation of permuting variables in the Boolean expression of a BF is equivalent to permuting the corresponding columns of the truth table and then reordering the rows (which include the output of the BF) in lexicographical order to bring the table into its canonical form. In the case of a 3-input BF, one is to apply a permutation $\sigma \in S_3$ (the symmetric group on 3 elements) and that will generate different orderings of the variables. The identity element of $S_3$ (absence of a permutation), $e$, does not change the order of the inputs so it corresponds to the ordering $(x_1, x_2, x_3)$. The transposition (1 2) interchanges the $1^{st}$ and $2^{nd}$ variables keeping the $3^{rd}$ variable fixed and generates the ordering: $(x_2, x_1, x_3)$. The transposition (1 3) interchanges the $1^{st}$ and $3^{rd}$ variables keeping the $2^{nd}$ variable fixed and generates the ordering: $(x_3, x_2, x_1)$. The transposition (2 3) interchanges the $2^{nd}$ and $3^{rd}$ variables keeping the $1^{st}$ variable fixed and generates to the ordering: $(x_1, x_3, x_2)$. The three-cycle (1 2 3) replaces the $1^{st}$ variable by the $2^{nd}$, the $2^{nd}$ by the $3^{rd}$ and the $3^{rd}$ by the $1^{st}$ and generates the ordering: $(x_2, x_3, x_1)$. The three-cycle (1 3 2) replaces the $1^{st}$ variable by the $3^{rd}$, the $2^{nd}$ by the $1^{st}$ and the $3^{rd}$ by the $2^{nd}$ and generates the ordering: $(x_3, x_1, x_2)$.

* These authors contributed equally: Priyotosh Sil and Ajay Subbaroyan
† To whom correspondence should be addressed:


#### Negation of BFs

An input $x_i$ of a BF $f$ is referred to as negated if the roles of 0 and 1 for that variable are exchanged. This operation generally changes the BF. It is not difficult to see that the table of this new BF can be obtained by swapping the values of the initial table's *output* column in all pairs of rows that differ only by the value of $x_i$. Equivalently, on the hypercube, this operation corresponds to swapping the hyperplanes $x_i = 0$ with $x_i = 1$ where $i$ is the index of the input to be negated. For example, consider a $k$-input BF $f$ where the order of the input variables in the associated truth table is $x_1, ..., x_i, ..., x_k$ and say

$$f(x_1, ..., x_i, ..., x_k) = T_1 \text{ and } f(x_1, ..., \overline{x_i}, ..., x_k) = T_2 \text{ where, } T_1, T_2 \in \{0, 1\}$$

Note that for 2 different sets of values for the inputs $x_1, x_2, \ldots, x_{i-1}, x_{i+1}, \ldots, x_{k-1}, x_k$, the 2 corresponding output values for $f(x_1, ..., x_i, ..., x_k) = T_1$ and $f(x_1, ..., \overline{x_i}, ..., x_k) = T_2$ may be different.

Then, the negation of the $i^{th}$ input leads to the new BF $f'$:

$$f'(x_1, ..., \overline{x_i}, ..., x_k) = T_1 \text{ and } f'(x_1, ..., x_i, ..., x_k) = T_2$$

As an example, consider $f = x_1 \wedge (\overline{x}_2 \vee x_3)$. Negation of the variable $x_1$ leads to the BF $f = \overline{x}_1 \wedge (\overline{x}_2 \vee x_3)$. Note that for any BF, there are $2^k$ negations possible, 2 per input, if we allow each input to be negated or not.

Using both permutations and negations, there are a total of $k!\ 2^k$ possible combinations of permutations and negations for a $k$-input BF. We can represent these permutations with negations as a matrix with rows corresponding to permutations and columns to negations. For illustration, consider the example provided above where $f_1 = x_1 \wedge (\overline{x}_2 \vee x_3)$:

$$\begin{array}{cccccccc} +\,+\,+ & +\,+\,- & +\,-\,+ & -\,+\,+ & +\,-\,- & -\,+\,- & +\,-\,- & -\,-\,- \end{array}$$

$$\mathbf{M} = \begin{bmatrix}
x_1 \wedge (x_2 \vee x_3) & x_1 \wedge (x_2 \vee \overline{x}_3) & x_1 \wedge (\overline{x}_2 \vee x_3) & \overline{x}_1 \wedge (x_2 \vee x_3) & x_1 \wedge (\overline{x}_2 \vee \overline{x}_3) & \overline{x}_1 \wedge (x_2 \vee \overline{x}_3) & \overline{x}_1 \wedge (\overline{x}_2 \vee x_3) & \overline{x}_1 \wedge (\overline{x}_2 \vee \overline{x}_3) \\
x_1 \wedge (x_3 \vee x_2) & x_1 \wedge (x_3 \vee \overline{x}_2) & x_1 \wedge (\overline{x}_3 \vee x_2) & \overline{x}_1 \wedge (x_3 \vee x_2) & x_1 \wedge (\overline{x}_3 \vee \overline{x}_2) & \overline{x}_1 \wedge (x_3 \vee \overline{x}_2) & \overline{x}_1 \wedge (\overline{x}_3 \vee x_2) & \overline{x}_1 \wedge (\overline{x}_3 \vee \overline{x}_2) \\
x_2 \wedge (x_1 \vee x_3) & x_2 \wedge (x_1 \vee \overline{x}_3) & x_2 \wedge (\overline{x}_1 \vee x_3) & \overline{x}_2 \wedge (x_1 \vee x_3) & x_2 \wedge (\overline{x}_1 \vee \overline{x}_3) & \overline{x}_2 \wedge (x_1 \vee \overline{x}_3) & \overline{x}_2 \wedge (\overline{x}_1 \vee x_3) & \overline{x}_2 \wedge (\overline{x}_1 \vee \overline{x}_3) \\
x_2 \wedge (x_3 \vee x_1) & x_2 \wedge (x_3 \vee \overline{x}_1) & x_2 \wedge (\overline{x}_3 \vee x_1) & \overline{x}_2 \wedge (x_3 \vee x_1) & x_2 \wedge (\overline{x}_3 \vee \overline{x}_1) & \overline{x}_2 \wedge (x_3 \vee \overline{x}_1) & \overline{x}_2 \wedge (\overline{x}_3 \vee x_1) & \overline{x}_2 \wedge (\overline{x}_3 \vee \overline{x}_1) \\
x_3 \wedge (x_1 \vee x_2) & x_3 \wedge (x_1 \vee \overline{x}_2) & x_3 \wedge (\overline{x}_1 \vee x_2) & \overline{x}_3 \wedge (x_1 \vee x_2) & x_3 \wedge (\overline{x}_1 \vee \overline{x}_2) & \overline{x}_3 \wedge (x_1 \vee \overline{x}_2) & \overline{x}_3 \wedge (\overline{x}_1 \vee x_2) & \overline{x}_3 \wedge (\overline{x}_1 \vee \overline{x}_2) \\
x_3 \wedge (x_2 \vee x_1) & x_3 \wedge (x_2 \vee \overline{x}_1) & x_3 \wedge (\overline{x}_2 \vee x_1) & \overline{x}_3 \wedge (x_2 \vee x_1) & x_3 \wedge (\overline{x}_2 \vee \overline{x}_1) & \overline{x}_3 \wedge (x_2 \vee \overline{x}_1) & \overline{x}_3 \wedge (\overline{x}_2 \vee x_1) & \overline{x}_3 \wedge (\overline{x}_2 \vee \overline{x}_1)
\end{bmatrix}$$

If one starts with an NCF, these operations generate new BFs that are themselves NCFs, and in fact NCFs with the same bias as the original NCF. If one denotes by $\odot$ the different operations involved (OR and AND), the previously described matrix is of the following form:

$$\begin{array}{cccc} ++...++ & ++...+- & \cdots & --...-- \end{array}$$

$$\mathbf{M} = \begin{bmatrix}
x_1 \odot (x_2 \odot ...(x_{k-1} \odot x_k)...) & x_1 \odot (x_2 \odot ...(x_{k-1} \odot \overline{x}_k)...) & \cdots & \overline{x}_1 \odot (\overline{x}_2 \odot ...(\overline{x}_{k-1} \odot \overline{x}_k)...) \\
x_2 \odot (x_1 \odot ...(x_{k-1} \odot x_k)...) & x_2 \odot (x_1 \odot ...(x_{k-1} \odot \overline{x}_k)...) & \cdots & \overline{x}_2 \odot (\overline{x}_1 \odot ...(\overline{x}_{k-1} \odot \overline{x}_k)...) \\
\vdots & \vdots & \ddots & \vdots \\
x_k \odot (x_{k-1} \odot ...(x_2 \odot x_1)...) & x_k \odot (x_{k-1} \odot ...(x_2 \odot \overline{x}_1)...) & \cdots & \overline{x}_k \odot (\overline{x}_{k-1} \odot ...(\overline{x}_2 \odot \overline{x}_1)...)
\end{bmatrix} \tag{1}$$

#### $2^i$-blocks of a BF

In the truth table for a $k$-input BF, there may arise pairs of rows ($R$,$R'$) such that:

$$R = (x_1 = a_1, x_2 = a_2, \ldots, x_{k-1} = a_{k-1}, x_k = 0) \text{ and } R' = (x_1 = a_1, x_2 = a_2, \ldots, x_{k-1} = a_{k-1}, x_k = 1)$$

In the canonical ordering of truth tables, these rows are consecutive and we refer to them as a $2^1$-block. We can similarly define $2^2$-blocks by considering the rows

$$(x_1 = a_1, x_2 = a_2, \ldots, x_{k-2} = a_{k-2}, x_{k-1} = 0, x_k = *) \text{ and } x' = (x_1 = a_1, x_2 = a_2, \ldots, x_{k-2} = a_{k-2}, x_{k-1} = 1, x_k = *)$$

where $* \in \{0, 1\}$ means that any values are allowed. Such blocks are now formed by $2^2$ consecutive rows. More generally, we will call a set of rows a $2^i$-block if it corresponds to combining two adjacent $2^{i-1}$-blocks corresponding to:

$$(x_1 = a_1, \ldots, x_{k-i} = a_{k-i}, x_{k-i+1} = 0, x_{k-i+2} = *, \ldots, x_k = *) \text{ and } (x_1 = a_1, \ldots, x_{k-i} = a_{k-i}, x_{k-i+1} = 1, x_{k-i+2} = *, \ldots, x_k = *)$$

In this way, one can go up to a $2^k$-block that in fact corresponds to all rows in the truth table.

### 2. TYPES OF BOOLEAN FUNCTIONS

#### Effective functions (EFs)

A BF $f$ with $k$ inputs is said to be *effective* if each input $i$ produces a change in the output under at least one specific context formed by all other inputs. Mathematically,

$$\forall\ i \in \{1,2,\ldots,k\},\ \exists\ \mathbf{x} \in \{0,1\}^k \text{ with } x_i = 0,\ f(\mathbf{x}) \neq f(\mathbf{x}+\mathbf{e}_i). \tag{2}$$

Here $\mathbf{e}_i \in \{0,1\}^k$ denotes the unit vector associated to the component of index $i$ having all entries set to 0 except for entry $i$.

The effectiveness of a BF in all its inputs is a necessary criterion for being biologically meaningful but not a sufficient one. Indeed, as $k$ increases, the fraction of BFs that are not effective goes to 0 [1]. Hence in practice EF BFs can be considered to be 'random', as we do in this work.

#### Biologically meaningful Boolean functions

Not surprisingly, there are a number of properties that random BFs do not often possess but are typical of biological BFs (as arising in reconstructed Boolean networks of biological systems). We now consider these types of BFs that are biologically meaningful.

##### *Unate functions (UFs)*

A BF $f$ with $k$ inputs is *unate* if each input $i$ to the BF is either monotone increasing (activatory) or monotone decreasing (inhibitory). An input $i$ (or variable $x_i$) is increasing monotone (or activatory) if:

$$\forall\ \mathbf{x} \in \{0,1\}^k \text{ with } x_i = 0,\ f(\mathbf{x}) \leq f(\mathbf{x}+\mathbf{e}_i), \tag{3}$$

and decreasing monotone (or inhibitory) if:

$$\forall\ \mathbf{x} \in \{0,1\}^k \text{ with } x_i = 0,\ f(\mathbf{x}) \geq f(\mathbf{x}+\mathbf{e}_i). \tag{4}$$

If all inputs to a BF are positive monotone, the BF is said to be a positive monotone BF.

##### *Read-once functions (RoFs)*

A BF $f$ with $k$ inputs is said to be *read-once* if using the operations of conjunction, disjunction and negation it can be expressed in such a way that each variable appears exactly once in the expression. Mathematically, a BF $f = f(x_1, x_2, \ldots, x_k)$ is a RoF if and only if there is a permutation $\sigma$ on its inputs $\{1,2,\ldots,k\}$ such that

$$f = x_{\sigma(1)} \odot x_{\sigma(2)} \odot x_{\sigma(3)} \ldots \odot x_{\sigma(k)}, \tag{5}$$

where $\odot$ represents the AND ($\wedge$) or OR ($\vee$) operator and in this expression we have omitted all parentheses that may need to be placed to define the function.

##### *Nested canalyzing functions (NCFs)*

A BF $f$ with $k$ inputs is *nested canalyzing* if there exists a permutation $\sigma$ on its inputs $\{1,2,\ldots,k\}$ such that:

$$f(\mathbf{x}) = \begin{cases} b_1 & \text{if } x_{\sigma(1)} = a_1, \\ b_2 & \text{if } x_{\sigma(1)} \neq a_1, x_{\sigma(2)} = a_2, \\ b_3 & \text{if } x_{\sigma(1)} \neq a_1, x_{\sigma(2)} \neq a_2, x_{\sigma(3)} = a_3, \\ \vdots & \\ b_k & \text{if } x_{\sigma(1)} \neq a_1, x_{\sigma(2)} \neq a_2, \ldots, x_{\sigma(k)} = a_k, \\ \overline{b}_k & \text{if } x_{\sigma(1)} \neq a_1, x_{\sigma(2)} \neq a_2, \ldots, x_{\sigma(k)} = \overline{a}_k. \end{cases} \tag{6}$$

In the above equation, $a_1, a_2, \ldots, a_k$ are the canalyzing input values and $b_1, b_2, \ldots, b_k$ are the canalyzed output values for inputs $\sigma(1), \sigma(2), \ldots, \sigma(k)$ in the permutation $\sigma$ of the $k$ inputs. Here, $\overline{a}_k$ and $\overline{b}_k$ are the complements of the Boolean values $a_k$ and $b_k$, respectively.

### 3. SAMPLING DIFFERENT TYPES OF BOOLEAN FUNCTIONS

For a given network structure, the total number of Boolean models that satisfy the fixed point constraints – even after imposing that the BFs be biologically meaningful (biologically plausible models) – is usually astronomical. Therefore, it is usually computationally infeasible to carry out exhaustive analysis on all the models of a biologically plausible ensemble, barring a few cases. In view of this fact, we resort to *sampling* the space of different biologically plausible ensembles (each one constrained by a different type of BF) in order to study the effect of the type of BF on the bushiness of the STG. To sample models uniformly from the space of a biologically plausible ensemble, it is sufficient to uniformly sample BFs at all nodes of the network. In other words, combining BFs that are sampled uniformly from the space of a particular type of BF and satisfying the fixed point constraints (and signs of inputs when necessary) results in a model that is uniformly drawn from the space of all possible models. So, to sample the space of a given biologically plausible ensemble, we first need to generate at each node, the exhaustive set or a sampled set of allowed BFs. We now explain how we do so for the different types of BFs.

#### Effective functions (EFs)

For nodes with up to 4 inputs, it is possible to list all possible EFs. For any such list, we check which ones satisfy the fixed point constraints. The resulting list is then the set of allowed BFs in this ensemble. When confronted with genes having more than 4 inputs, it is infeasible to generate the exhaustive list of EFs so we sample them instead. To do so, we leverage the fact that most BFs are EFs for $k \geq 4$ [1]. Specifically, we generate a random $k$-input EF by randomly assigning the output of rows of the truth table to either 0 or 1, except for the rows associated with the biological attractors that are fixed by precisely those constraints. We then check whether the resulting BF is EF. If it is, we are done. If it is not, we try again until obtaining an effective BF. This rejection-based random generation of BFs samples the space of EFs uniformly, and all the more efficiently that the fraction of EFs in the space of all BFs is close to 1 (as occurs with increasing $k$).

#### Effective and unate functions (EUFs)

For nodes with up to 6 inputs, an exhaustive approach is possible. That is done by first generating the exhaustive list of all the UFs for which all inputs are activatory. (Such a list is far smaller than if we allow both signs for each input.) We then step through this list of UFs and for each we generate the corresponding UF with the desired sign for each input. Specifically,

(a) we flip the signs of the inputs of the UF so as to match the sign combination provided for that node in the original Boolean model.

(b) we check whether the UF obtained is effective in all of its inputs.

(c) if (b) is True, we check whether that function satisfies the biological fixed point constraints. If (b) is False, we skip (d) and go to (a) for the next BF in the list.

(d) if (c) is True, we store the resulting EUF. If (c) is False, we discard the EUF and go to (a) for the next BF in the list.

The obtained list of EUFs constitutes the allowed EUFs for the considered node.

Beyond 6 inputs, it is computationally infeasible to generate the exhaustive list of UFs, and consequently we require a different algorithm based on a sampling approach. The problem of sampling a UF with a given sign combination can easily be mapped to the problem of sampling a positive monotone BF following the same logic as in the enumeration case described previously. Thus, we will now describe how we sample positive monotone BFs.

##### *Sampling positive monotone functions*

Our algorithm to sample positive monotone BFs is based on the idea that on a $k$-dimensional hypercube, positive monotonicity forces the ‘propagation’ of the color of a red or blue vertex (corresponding to output values 1 or 0) to its 1-Hamming neighbours that are ‘ahead’ or ‘behind’ it respectively. We now describe such a propagation of vertex colors on the Boolean hypercube.

The propagation of the color of a vertex due to monotonicity is illustrated in SI Fig. S2. In SI Fig. S2(a), the vertex 4(0100) is colored red (assigned the value 1) while the other vertices are not yet colored. The 1-Hamming neighbours of vertex 4 are (0(0000), 5(0101), 6(0110), 12(1100)). If any of the vertices 5, 6 or 12 were colored blue, then the corresponding inputs $x_4$, $x_2$ or $x_1$ respectively would violate positive monotonicity. But vertex 0(0000) can be colored either red or blue without violating the positive monotonicity condition on input $x_3$. Thus, the vertices 5, 6 or 12 are forced to be red. Note that the 1-Hamming neighbours of vertex 4 labelled with the integers 5, 6 and 12 are “larger” than 4, that is can be reached from 4 using positive increases to the variables. Thus if a vertex is colored red, all “larger” nodes should also be colored red, and in particular the ones that are 1-Hamming neighbors. This ordering is easily identified provided that the vertices of Boolean hypercube are labeled by the integer equivalent of the binary string associated with the vertices. We then say that the vertices 5, 6 and 12 are ‘ahead’ of vertex 4 which then can propagate its color to them. This propagation is continued iteratively until no nodes are ahead of the ones colored (see bottom right panel of SI Fig. S2(a)).

A similar argument of propagation of color holds if, in the initial configuration of the BF, the vertex 4 was colored blue instead of red (see SI Fig. S2(b)). The only difference being that the propagation of the blue color of vertex 4 is now towards the 1-Hamming neighbors (vertex 0) that have an integer label smaller than that of node under consideration. In this case, that vertex is 0. Now suppose vertex 0 was colored red, then the input $x_3$ violates the condition of positive monotonicity. Thus, the vertex 0 is forced to be blue. We say that the vertex 0 is 'behind' the vertex 4. The vertex 0 does not have any vertex 'behind' it, hence the propagation of color is terminated here as shown in the bottom panel of SI Fig. S2(b). In sum, if we demand that a BF is positive monotone in all its inputs, then any vertex colored red (respectively blue) can iteratively propagate its color to 1-Hamming neighbors 'ahead' of it (respectively 'behind' it).

In order to sample positive monotone BFs, we begin with a hypercube in which the vertices are not colored. We then choose a random vertex from the set of uncolored vertices (all vertices are equally probable) and color it randomly (either red or blue). The color of this vertex is then iteratively propagated till no further propagation of color is possible. If any uncolored vertices remain on the hypercube, a vertex is randomly chosen from the set of uncolored vertices, colored randomly and iteratively propagated till no further propagation of color is possible. This procedure is repeated till all the vertices of the hypercube are colored. The final outcome is a positive monotone BF. Note that this BF may or may not be effective in all its inputs. To sample positive monotone BFs that are effective in all inputs, we simply sample a positive monotone BF and check if it is an EF. If it is not, we reject the BF and start again until obtaining an EF.

Notably, our sampling algorithm can generate all the positive monotone EUFs. An immediate question is whether the above algorithm samples positive monotone EUFs *uniformly*. To answer this question, we sampled $n = 10000$, $n = 20000$ and $n = 1000000$ positive monotone EUFs with $k = 3$, $k = 4$ and $k = 5$ inputs respectively (see SI Fig. S4). The sampling is not exactly uniform but it is close enough to uniform for our purposes.

#### *Sampling unate functions with a given sign combination that satisfies fixed point constraints*

The above mentioned algorithm can be easily extended to generate UFs that satisfy predefined constraints on the BF, namely, biological fixed point constraints. We now show how this sampling can be achieved using the logic provided in SI Fig. S3 on a BF for which we assume that the signs of the 4 inputs $x_1$, $x_2$, $x_3$ and $x_4$ are '$+--+$'. The next assumption is that the fixed points constrain the vertices 7 and 10 to have the output values 0 and 1 respectively. Note that the fixed point constraints have to be consistent with the given signs, otherwise there is no UF that will satisfy the fixed point constraints. Since the BF in SI Fig. S3(a) is consistent in the manner described above, we can now begin the sampling procedure. We first flip all the inhibitory inputs to make them activatory. This is achieved by swapping the hyperplanes $x_2 = 0$ with $x_2 = 1$, and $x_3 = 0$ with $x_3 = 1$ as shown in SI Fig. S3(b), thereby resulting in a positive monotone UF. We now use the method described in the previous section to sample positive monotone UFs as shown in SI Figs. S3(c)-(g). Once a positive monotone BF is obtained as shown in SI Fig. S3(g), the signs of the inputs are returned to the initial sign combination by re-swapping the hyperplanes $x_2 = 0$ with $x_2 = 1$, and $x_3 = 0$ with $x_3 = 1$. Note that on reswapping these hyperplanes, the fixed point constraints (SI Fig. S3(h)) return to their original configuration as shown in SI Fig. S3(a). The final BF is thus a random UF with the desired signs for the inputs and that satisfies the fixed point constraints. As before, the BF generated may be ineffective in some inputs. We thus again check if the sampled UF is an EF. If not, we keep sampling till we find an UF that is an EF.

### Read-once functions (RoFs)

The logic here is very similar to the one for sampling EUF BFs. For nodes with up to 8 inputs, it is possible to generate an exhaustive list of all the RoFs for which all inputs are activatory. We then step through this list of RoFs and for each function:

(a) we flip the signs of the inputs of the RoF so as to match the sign combination provided for that node in the original Boolean model.

(b) we check if the RoF obtained from (a) satisfies the biological fixed point constraints.

(c) if (b) is True, we store the resulting RoF. If (b) is False, we discard the RoF and go to the next iteration.

The obtained list of RoFs constitute the allowed RoFs for that node.

### Nested canalyzing functions (NCFs)

For nodes with up to 8 inputs, it is possible to generate an exhaustive list of all the NCFs with all inputs being activatory. Following the logic for EUF BFs, we step through this list of NCFs and for each function:

(a) we flip the signs of the inputs of the NCF so as to match the sign combination provided for that node in the original Boolean model.

(b) check if the NCF obtained from (a) satisfies the biological fixed point constraints.

(c) if (b) is True, we store the resulting NCF. If (b) is False, we discard the NCF and go to the next iteration.

The obtained list of NCFs constitutes the allowed NCFs for that node.

In sum, we have algorithms to sample different classes of BFs, namely, EFs, EUFs, RoFs and NCFs subject to fixed point constraints. We exhaustively list all EFs up to 4 inputs and sample them for larger $k$. We uniformly sample EUFs with given sign combination up to 6 inputs, and use our iterative vertex color propagation algorithm to sample them when there are more than 6 inputs. We uniformly sample RoFs and NCFs with a given sign combination up to 8 inputs.

### 4. A BRIEF OVERVIEW OF THE 10 PUBLISHED BOOLEAN NETWORK MODELS USED IN THIS STUDY

In this work, we study the bushiness of the STG using 10 published Boolean network models. These 10 Boolean models are listed in Table 1, in the main manuscript and are succinctly described below.

#### Root Stem Cell Niche GRN (RSCN-GRN)

The RSCN Boolean model (*model A* in [2]) is a reconstructed GRN that controls the differentiation of cells located in the root apex of the plant *Arabidopsis thaliana*, a region referred to as the root stem cell niche. The BFs of this model are provided in SI Table S1 and the type of BF for each gene is provided in SI Table S11. The nodes (genes), their inputs and the sign of the regulatory interactions in the network are all derived from the BoolNet file (SI Table S1). This Boolean model has 4 fixed points corresponding to these cell types: Quiescent center (QC), Vascular initials (VI), Cortex-Endodermis initials (CEI) and Columella epidermis initials (CEpI) (SI Table S21). On applying the various biologically meaningful constraints (see SI text, section 3) the number of BFs allowed at each node of the RSCN-GRN is given in SI Table S31. The BF for the AUX node for all models in all 4 ensembles were chosen to be the same as the one provided in [3].

#### Epithelial-Mesenchymal Transition GRN (EMT-GRN)

The EMT-GRN is a sub-network of a 150 node reconstructed GRN that controls EMT via mechanosensing–mitogen crosstalk [4]. The BFs of this model are provided in SI Table S2 and the type of BF for each gene is provided in SI Table S12. The nodes, their inputs and the sign of the regulatory interactions in the network are all derived from the BoolNet file (SI Table S2). This Boolean model has 3 fixed points corresponding to these cell types: Epithelial, Mesenchymal and Hybrid (see SI Table S22). On applying the various biologically meaningful constraints (see SI text, section 3), the number of BFs allowed at each node of the EMT-GRN is given in SI Table S32.

#### NeoCortex Developmental GRN (NCD-GRN)

The NCD-GRN is a network that models cellular differentiation in the earliest stages of cortical arealisation [5]. The BFs of this model are provided in SI Table S3 and the type of BF for each gene is provided in SI Table S13. The nodes, their inputs and the sign of the regulatory interactions in the network are all derived from the BoolNet file (SI Table S3). This Boolean model has 2 fixed points corresponding to Anterior and Posterior compartments of the developing neocortex (see SI Table S23). On applying the various biologically meaningful constraints (see SI text, section 3) the number of BFs allowed at each node of the NCD-GRN is given in SI Table S33.

#### EHD-GRN: Early Heart Development GRN

The EHD-GRN models two areas of differential gene expression that arise from a common cardiovascular progenitor cell population [6]. The BFs of this model are provided in SI Table S4 and the type of BF for each gene is provided in SI Table S14. The nodes, their inputs and the sign of the regulatory interactions in the network are all derived from the BoolNet file (SI Table S4). This Boolean model has 2 fixed points corresponding to 2 different areas with differential expression of genes, namely, First Heart Field (FHF) and Second Heart Field (SHF) (see SI Table S24). On applying the various biologically meaningful constraints (see SI text, section 3) the number of BFs allowed at each node of the EHD-GRN is given in SI Table S34. Exogen BMP2 (exogen_BMP2_II) is an external signal that is 'fixed' to an output value of '1'. Since the number of nodes in this model is 15 but there is one fixed node, the size of the state space is $2^{14}$.

#### Myeloid Differentiation GRN (MD-GRN)

The MD-GRN models hematopoietic stem cell differentiation. In particular, it models myeloid differentiation from common myeloid progenitors to various cell types [7]. The BFs of this model are provided in SI Table S5 and the type of BF for each gene is provided in SI Table S15. The nodes, their inputs and the sign of the regulatory interactions in the network are all derived from the BoolNet file (SI Table S5). This Boolean model has 4 fixed points corresponding to these cell types: granulocytes, erythrocytes, megakaryocytes and monocytes (see SI Table S25). On applying the various biologically meaningful constraints (see SI text, section 3) the number of BFs allowed at each node of the MD-GRN is given in SI Table S35.

#### T helper cell differentiation GRN (THC-GRN)

The THC-GRN models the T helper cell network that governs the differentiation of precursor T helper cells (Th0 cells) to effector T helper cells (Th1 and Th2 cells) [8]. The BFs of this model are provided in SI Table S6 and the type of BF for each gene is provided in SI Table S16. The nodes, their inputs and the sign of the regulatory interactions in the network are all derived from the BoolNet file (SI Table S6). This Boolean model has 3 fixed points corresponding to the cell types: Th0, Th1 and Th2 (see SI Table S26). On applying the various constraints (see SI text, section 3) the number of BFs allowed at each node of the THC-GRN is given in SI Table S36. Of the 23 nodes in this network, 4 do not have any inputs and are fixed to '0', namely IFN-$\beta$, IL-12, IL-18 and TCR. Thus the size of the state space is $2^{19}$.

#### EGFR signalling pathway (EGFR-GRN)

The EGFR-GRN models the regulation of cell division and apoptosis by the EGFR signalling pathway and a BRCA1/TP53 DNA damage response module [9]. The BFs of this model are provided in SI Table S7 and the type of BF for each gene is provided in SI Table S17. The nodes, their inputs and the sign of the regulatory interactions in the network are all derived from the BoolNet file (SI Table S7). This Boolean model has 2 fixed points corresponding to these cell states: apoptosis and cell division (see SI Table S27). On applying the various constraints (see SI text, section 3) the number of BFs allowed at each node of the EGFR-GRN is given in SI Table S37.

#### Epithelial-Mesenchymal Transition (with Senescence) (EMT-Senescence-GRN)

The EMT-Senescence-GRN models epithelial to mesenchymal transition in the context of epithelial cancer [10]. The BFs of this model are provided in SI Table S8 and the type of BF for each gene is provided in SI Table S18. The nodes, their inputs and the sign of the regulatory interactions in the network are all derived from the BoolNet file (SI Table S8). This Boolean model has 3 fixed points corresponding to these cell types: epithelial cells, senescent epithelial cells and stem-like mesenchymal cells with a tumorigenic potential (see SI Table S28). On applying the various constraints (see SI text, section 3) the number of BFs allowed at each node of the EMT-Senescence-GRN is given in SI Table S38.

#### Flower Organ Specification GRN (FOS-GRN)

The FOS-GRN controls early flower development in *Arabidposis thaliana*, in particular, the primordial cells of inflorescence, sepals, petals, stamens and carpels. The BFs of this model are provided in SI Table S9 and the type of BF for each gene is provided in SI Table S19. The nodes, their inputs and the sign of the regulatory interactions in the network are all derived from the BoolNet file (SI Table S9). This Boolean model has 10 fixed points corresponding to these cell types: inflorescence (4 cell types), sepal (1 cell type), petal (2 cell types), stamen (2 cell types) and carpel (1 cell type) (see SI Table S29). On applying the various constraints (see SI text, section 3) the number of BFs allowed at each node of the FOS-GRN is given in SI Table S39. Since the computation of the $Z$-parameters was computationally expensive for 7 and 8 input BFs, such as AP3 and AG respectively, we sampled 100,000 models for each of the 4 ensembles.

#### GSD-GRN: Gonadal Sex Determination GRN

The Gonadal sex determination involves the decision taken by bipotential gonadal primordium (BGP) to differentiate to testis or ovaries. In particular, the GSD-GRN governs cellular differentiation from Sertoli progenitor cells (SPC) and granulosa progenitor cells (GPC) towards Sertoli (in testis) and granulosa (in ovaries) cells respectively [11]. The BFs of this model are provided in SI Table S10 and the type of BF for each gene is provided in SI Table S20. The nodes, their inputs and the sign of the regulatory interactions in the network are all derived from the BoolNet file (SI Table S10). This Boolean model has 2 fixed points corresponding to these cell types: sertoli and granulosa (see SI Table S30). On applying the various constraints (see SI text, section 3) the number of BFs allowed at each node of the GSD-GRN is given in SI Table S40. Since the computation of the $Z$-parameters was computationally expensive for 7 and 8 input BFs, namely, SOX9 and SRY respectively, we sampled 100,000 models for each of the 4 ensembles.

### 5. AVERAGE IN-DEGREE OF NON-GOE STATES ($\langle d_{\text{in}} \rangle_{non\text{-}GoE}$) AS A FUNCTION OF $G$-DENSITY

Consider a GRN with $N$ nodes (genes) and associated update rules. Then, the associated STG will have $2^N$ states. Suppose, the number of $GoE$ states is $G$. Then the $G$-density associated with the STG is given by

$$G\text{-density} = \frac{G}{2^N}$$

In a STG, each state has exactly one successor and so the number of edges in the STG is equal to $2^N$. Since, the number of $GoE$ states is $G$, number of $non\text{-}GoE$ states must be $2^N - G$. Hence, the average in-degree of the $non\text{-}GoE$ states ($\langle d_{in} \rangle_{non-GoE}$) is given by the total number of edges divided by the total number of $non\text{-}GoE$ states:

$$\begin{aligned} \langle d_{in} \rangle_{non-GoE} &= \frac{2^N}{2^N - G} \\ &= \frac{1}{1 - \frac{G}{2^N}} \\ &= \frac{1}{1 - G\text{-density}} \end{aligned}$$

### 6. A BRIEF OVERVIEW OF 1-DIMENSIONAL CELLULAR AUTOMATA

A cellular automaton (CA) is a self-contained discrete dynamical system. It consists of cells (or sites) arranged on a regular lattice and that are assigned values from a finite alphabet. These values are synchronously updated at each time step according to a function that takes as inputs the values of cells in a fixed neighborhood of each cell.

Wuensche considered the simple case of 1-dimensional (1D) CAs consisting of a 1D ring of $N$ sites with each site carrying a value '0' or '1' [12]. The values $C_i$ at each site $i$ get updated synchronously in discrete time steps according to a Boolean rule whose inputs are the values of $i$'s neighboring sites but can include the site $i$ itself.

If we consider a 1D lattice with $N$ cells and a size $K$ neighbourhood then the time evolution of $i^{th}$ cell can be represented by [12],

$$C_i^{(t+1)} = f\left(C_{i-r_{left}}^{(t)}, \ldots, C_{i-1}^{(t)}, C_i^{(t)}, C_{i+1}^{(t)}, \ldots, C_{i+r_{right}}^{(t)}\right)$$

where to satisfy the periodic boundary conditions, for $x < 1$, $C_x = C_{N+x}$ while for $x > N$, $C_x = C_{x-N}$. Here, $r_{left}$ and $r_{right}$ are the neighborhood radii (number of neighbors) to the left and right respectively, and thus $K = r_{left} + 1 + r_{right}$.

A 1D CA architecture can be considered as a network having the cells (or sites) as its nodes and the interacting $K$-neighbors as the input edges to the nodes. At any instant, each cell (or site) holds a value either '0' or '1'. The arrangement of these 0s and 1s across the entire network forms the network's current configuration or *state*, *i.e.*, its list of values $C_i$ at all sites, which can be represented by a binary string. At each time step the synchronous update transforms the current state to the subsequent one, a process that can be iterated indefinitely. A time sequence of states is called trajectory. Having a finite state space of size $2^N$, a trajectory will either converge to a fixed point or to a limit cycle, both of which are termed 'attractors'. Each state has exactly one 'successor' but can have a variable number of immediate *pre-images* or 'predecessors', from 0 to $2^N$. The number of pre-images of a state is called its in-degree.

### 7. THE Z-PARAMETER IN CELLULAR AUTOMATA

Various quantities have been introduced in CA based on properties of the rule tables, in particular to characterize the global nature of CA dynamics [13–15]. One such parameter which is relevant to this work is Wuensche's $Z$-parameter. Wuensche proposed the $Z$-parameter as an indicator of irreversibility of the discrete dynamics [12, 16–18]. The starting point considers an arbitrary CA state and asks which could be its pre-images. Wuensche tries to determine these pre-images by exploiting the 1D architecture of the CA, attempting to infer the values of consecutive cells by shifting a window from left to right. He thus introduces the $Z_{left}$ parameter as the probability that the value of the next unknown cell to the right of the current partial pre-image is unique, corresponding to reversible dynamics. This probability is calculated from the CA rule table (see section 2.4.1) in main manuscript for the methodology of computation of $Z_{left}$) by determining the fraction of configurations where the uniqueness condition holds. In a similar fashion, one can compute the value of $Z_{right}$ giving the probability that the value of the next unknown cell to the left of the partial pre-image has a unique value. Wuensche then defines his $Z$-parameter as the maximum of $Z_{left}$ and $Z_{right}$ *i.e.*,

**Algorithm 1** Recursive algorithm to determine the vector **n** arising in the computation of $Z_{left}$.

**Input 1:** $f$ is the truth table of the BF represented as a string of bits.
**Input 2:** $k_{var}$ is the number of inputs of $f$.
**Input 3:** $k_{fixed}$ is the number of inputs of the BF for which the $Z_{left}$ is to be computed.
**Output:** The array of elements of the vector **n**.

```
function ZLEFTVECTOR(f, k_var, k_fixed)
    if k_var == 1 then                          ▷ If the contribution of 2^1-blocks have to be computed
        if (f ==‘01’) or (f ==‘10’) then
            return [2, 0, . . . , 0]            ▷ The n vector of length k_fixed with element n_k = 2 and other elements as 0
        else
            return [0, 0, . . . , 0]            ▷ The n vector of length k_fixed with all elements as 0
        end if
    else
        Split f into upper (f_upper) and lower (f_lower) halves.
        Uniquef_upper ← Values arising in f_upper
        Uniquef_lower ← Values arising in f_lower
        if (Length of Uniquef_upper is 1) & (Length of Uniquef_lower is 1) & (Uniquef_upper ≠ Uniquef_lower) then ▷ These
    conditions guarantee that there will be a contribution to n vector
            CountArray ← [0, 0, . . . , 0]      ▷ Count array is the n vector
            CountArray[k_inp − 1] ← 2^(k_inp − 1)
            return CountArray
        else
            return ZLEFTVECTOR(f_upper, k_var − 1, k_fixed) + ZLEFTVECTOR(f_lower, k_var − 1, k_fixed)
        end if
    end if
end function
```

$$Z = max(Z_{left}, Z_{right})$$

Being a probability, $Z$ varies between 0 and 1. Ideally a value of $Z$ close to 1 should indicate that there is a unique pre-image for many – if not most – states. Then many or most of the STG's nodes should have in-degrees equal to 1 (corresponding to reversible dynamics) in which case the STG will not be bushy at all. In contrast, a low $Z$ value should indicate strongly irreversible dynamics in which case the STG will have many states with a high in-degree. Because the average in-degree of the STG is mathematically constrained to be 1, this situation will also lead to having many *GoE* states, so the STG will be bushy. In the Main text we showed how to compute $Z_{left}$ of a general BF (whether in the context of a CA or not) by following Wuensche's prescriptions [18]. Algorithm 1 provides the associated logic in the form of pseudocode. In the next section we derive a number of associated properties.

### 8. PROPERTIES OF $Z$ PARAMETERS

Let $f$ be a $k$-input Boolean function $f$. We then have the following properties.

**Property 8.1.** $Z_{left}$ of $f$ is invariant under the negation of any of its inputs.
*Proof*: In the truth table representation of $f$, negation of the $i^{th}$ input $x_i$ will swap the outputs belonging to the lower and upper half sub-blocks of each $2^{k-i+1}$-block. Since $Z_{left}$ is completely determined by the $n_i$'s and $k$, the negation of the $i^{th}$ input will swap the output values of the associated sub-blocks, keeping the value of $n_i$ unchanged, and consequently leaving $Z_{left}$ unchanged. The negation of multiple inputs will also keep the value of $Z_{left}$ unchanged since the negation of multiple inputs can be achieved by successively negating single inputs.

**Property 8.2.** $Z_{left}$ of $f$ is invariant under complementation.
*Proof*: Let $g = \overline{f}$ be the complement of $f$. Then for any input combination, if $f$ is $T$, $g$ is $\overline{T}$. Consider

$$\begin{aligned} f(x_1, x_2, \ldots, x_{k-i}, 0, *, \ldots, *, *) &= T \\ f(x_1, x_2, \ldots, x_{k-i}, 1, *, \ldots, *, *) &= \overline{T} \end{aligned}$$

where $T \in \{0, 1\}$ and $*$ may be 0 or 1. Then,

$$\begin{aligned} g(x_1, x_2, \ldots, x_{k-i}, 0, *, \ldots, *, *) &= \overline{T} \\ g(x_1, x_2, \ldots, x_{k-i}, 1, *, \ldots, *, *) &= \overline{\overline{T}} = T \end{aligned}$$

As a result, $n_i$'s are the same for $f$ and $g$ for any $i \in \{1, 2, ..., k\}$ and consequently so are the $R_i$ values. Thus $Z_{left}$ values of $f$ and $g$ are equal.

**Property 8.3.** The maximum value of $Z_{left}$ over all $k$-input BFs with bias $P$ is:

$$Z_{left} = \begin{cases} \dfrac{P}{2^{k-1}} & \text{if } P \leq 2^{k-1} \\ 2 - \dfrac{P}{2^{k-1}} & \text{if } P > 2^{k-1} \end{cases} \tag{7}$$

*Proof*: For a $k$-input BF $f$ with bias $P \leq 2^{k-1}$, consider the vector $\mathbf{n} = (n_1, n_2, \ldots, n_{k-1}, n_k)$ as defined in section 2.4.1 in the main text. Recall that $n_i$ counts the number of lines in the truth table that arise in CC-$2^{k-i+1}$ blocks, each of which consists of lines where half of the output values are 0s and the other half are 1s. Then, using the fact that all CC-blocks (of all sizes) are non overlapping, the total number of these CC-block associated lines is twice the number of 1s in those same lines. This immediately leads to

$$\sum_{i=1}^{i=k} n_i \leq 2P$$

i.e., the sum of the elements of the vector $\mathbf{n}$ must be at most $2P$.

Consider now the expression of $Z_{left}$,

$$\begin{aligned} Z_{left} &= R_k + R_{k-1}(1 - R_k) + R_{k-2}(1 - R_{k-1})(1 - R_k) + R_{k-3}(1 - R_{k-2})(1 - R_{k-1})(1 - R_k) + ... \\ &\leq R_k + R_{k-1} + R_{k-2} + \ldots + R_1 \\ &= \frac{n_k}{2^k} + \frac{n_{k-1}}{2^k} + \ldots + \frac{n_1}{2^k} \\ &= \frac{n_k + n_{k-1} + \ldots + n_1}{2^k} \\ &\leq \frac{2P}{2^k} \\ &= \frac{P}{2^{k-1}} \end{aligned}$$

showing that $Z_{left}$ is bounded from above by $P/2^{k-1}$ (if $P \leq 2^{k-1}$). In addition, for each $P \leq 2^{k-1}$ there are functions $f$ that reach this maximum value. For instance for any $P$, consider a function of the type

$$f = 000\ldots0\overbrace{0101\ldots01}^{P \text{ `01' pairs}}$$

Clearly for such a choice, $n_k = 2P$ and $n_i = 0 \; \forall \; i \neq k$. This results in $Z_{left} = P/2^{k-1}$.

So, for all bias values $P \leq 2^{k-1}$, the upper bound of $Z_{left}$ is $P/2^{k-1}$. From property (8.2) it follows that for any function $f$ with bias value $P > 2^{k-1}$, it would have the same value of $Z_{left}$ as its complement function. Now, consider a BF $f$ with bias $P > 2^{k-1}$. Then the maximum possible value of $Z_{left}$ is the same for $f$ and $\overline{f}$ (complement of $f$). However, the bias of $\overline{f}$ is $2^k - P$ and since $P > 2^{k-1}$, $2^k - P < 2^{k-1}$. Hence, the maximum value of $Z_{left}$ for $\overline{f}$ is $\frac{2^k - P}{2^{k-1}} = 2 - \frac{P}{2^{k-1}}$. Finally, the maximum possible value of $Z_{left}$ for $f$ is then $2 - \frac{P}{2^{k-1}}$. This proves that the maximum value of $Z_{left}$ for any $k$-input BF with bias $P$ is:

$$Z_{left} = \begin{cases} \dfrac{P}{2^{k-1}} & \text{if } P \leq 2^{k-1} \\ 2 - \dfrac{P}{2^{k-1}} & \text{if } P > 2^{k-1} \end{cases} \tag{8}$$

**Property 8.4.** NCFs have the maximum $Z_{max}$ for any $k$-input BF with odd bias $P$.
*Proof*: Consider the output column of the truth table for the following $k$-input BF with all inputs activatory and having a bias $P$ such that $P < 2^{k-1}$:

$$f = 000\ldots0\overbrace{0101\ldots01}^{P \text{ `01' pairs}}$$

It is not difficult to see that this BF is an NCF with

$$\begin{aligned} & n_k = 2P \text{ and } n_i = 0 \; \forall \; i \in \{1, 2, \ldots, k-1\} \\ \Longrightarrow \quad & R_k = \frac{2P}{2^k} = \frac{P}{2^{k-1}} \text{ and } R_i = 0 \; \forall \; i \in \{1, 2, \ldots, k-1\} \\ \Longrightarrow \quad & Z_{left} = \frac{P}{2^{k-1}} + 0\left(1 - \frac{P}{2^{k-1}}\right) + 0\left(1 - \frac{P}{2^{k-1}}\right)(1 - 0) + ... \\ \Longrightarrow \quad & Z_{left} = \frac{P}{2^{k-1}} \end{aligned}$$

We thus see that this NCF has reached the maximum possible value (given the results above) and thus its $Z_{max}$ value is precisely $Z_{left}$. Interestingly, this same value is that of all NCFs with that same bias $P$ because the set of all NCFs of a given bias are obtained by permuting and negating variables of this particular NCF, a result that was shown in [1]. The desired result then follows from the fact that $Z_{left}$ is invariant under negations. Consider now the case of a function $g$ with bias $P > 2^{k-1}$. The bias of the complement of our function is then $2^k - P$; this complementation does not change $Z_{left}$. Hence, the value of $Z_{max}$ of our complemented function is $\frac{2^k-P}{2^{k-1}} = 2 - \frac{P}{2^{k-1}}$ (since $2^k - P < 2^{k-1}$). Consequently, the value of $Z_{max}$ of $g$ is $2 - \frac{P}{2^{k-1}}$. In summary, the $Z_{max}$ of a $k$-input NCF with a bias $P$ (which is necessarily odd) is given by:

$$Z_{max} = \begin{cases} \frac{P}{2^{k-1}} \text{ if } P < 2^{k-1}, \\ 2 - \frac{P}{2^{k-1}} \text{ if } P > 2^{k-1} \end{cases}$$

**Property 8.5.** The minimum value for $Z_{left}$ for any non-constant $k$-input BF with bias $P$ is $1/2^{k-1}$.
*Proof*: The minimum possible value of $Z_{left}$ across all $k$-input BFs is 0 (in particular for 'constant' BFs with bias 0 and bias $2^k$). For these BFs, $R_i = 0$ for all $i \in \{1, 2, \ldots k\}$. The minimum possible value of $Z_{left}$ across all non-constant $k$-input BFs are achieved by BFs with a single CC-$2^1$-block. For these BFs, $Z_{left}$ is $2/2^k = 1/2^{k-1}$ since $n_k = 2$ and $n_i = 0 \; \forall \; i \in \{1, 2, \ldots, k-1\}$. There are functions $f$ that reach this minimum value (for instance bias 1 and bias $2^k - 1$ functions).

**Property 8.6.** NCFs have the minimum $Z_{min}$ for any $k$-input BF with odd bias $P$.
*Proof*: Consider the output column of the truth table of a $k$-input NCF with bias $P$ where all inputs are 'activatory':

$$f = \underbrace{000\ldots0}_{(2^k-P)\text{ 0's}}\overbrace{11\ldots1}^{P\text{ 1's}}$$

Since $P$ is odd for NCFs [1, 19] we have,

$$\begin{aligned} & n_k = 2 \text{ and } n_i = 0 \; \forall \; i \in \{1, 2, \ldots, k-1\} \\ \implies \quad & R_k = \frac{2}{2^k} = \frac{1}{2^{k-1}} \text{ and } R_i = 0 \; \forall \; i \in \{1, 2, \ldots, k-1\} \\ \implies \quad & Z_{left} = \frac{1}{2^{k-1}} \end{aligned}$$

$Z_{min}$ is the minimum of the $Z_{left}$ values over all permutations of a BF and $Z_{left}$ is invariant under negation of any of its inputs. Again, there is only one equivalence class per bias for NCFs (equivalence under both permutations and negations), so $Z_{min}$ for any $k$-input NCF is given by $1/2^{k-1}$. Hence, NCFs have the minimum $Z_{min}$ for any $k$-input BF with odd bias $P$. The value of $Z_{min}$ for any $k$-input NCF is independent of bias ($P$) and its value is $1/2^{k-1}$.

**Property 8.7.** The value of $Z_{mid}$ for any $k$-input and $P$-bias NCF is:

$$Z_{mid} = \begin{cases} \dfrac{1+P}{2^k} & \text{if } P < 2^{k-1} \\ 1 - \dfrac{P-1}{2^k} & \text{if } P > 2^{k-1} \end{cases} \tag{9}$$

*Proof*: $Z_{mid}$ of a BF is defined as the average of its $Z_{max}$ and $Z_{min}$ values (see section 2.4.2 in the main text). $Z_{min}$ and $Z_{max}$ for any $k$-input NCF with bias $P$ (see Properties 8.4 and 8.6) is given by:

$$\begin{aligned} Z_{min} &= \frac{1}{2^{k-1}} \\ Z_{max} &= \begin{cases} \frac{P}{2^{k-1}} \text{ if } P < 2^{k-1}, \\ 2 - \frac{P}{2^{k-1}} \text{ if } P > 2^{k-1} \end{cases} \end{aligned}$$

Hence,

$$\begin{aligned} Z_{mid} &= \frac{Z_{max} + Z_{min}}{2} \\ &= \begin{cases} \frac{1}{2}\left(\frac{1}{2^{k-1}} + \frac{P}{2^{k-1}}\right) \text{ if } P < 2^{k-1}, \\ \frac{1}{2}\left(\frac{1}{2^{k-1}} + 2 - \frac{P}{2^{k-1}}\right) \text{ if } P > 2^{k-1} \end{cases} \\ &= \begin{cases} \frac{1+P}{2^k} \text{ if } P < 2^{k-1}, \\ 1 - \frac{P-1}{2^k} \text{ if } P > 2^{k-1} \end{cases} \end{aligned}$$

### 9. RESULTS ON NON-EQUIVALENT PERMUTATIONS OF THE $k!$ PERMUTATIONS OF BOOLEAN FUNCTIONS

#### Definitions and theorems

**Definition 1** (Left coset)**.** Let $G$ be a group and $H$ be a subgroup of $G$. Then for any element $a \in G$ the set $\{ah \mid h \in H\}$ is called the *left coset* of $H$ in $G$ containing $a$ and is denoted by $aH$. In this case, $a$ is called the coset representative of $aH$.

Note: The cardinality of any left coset of $H$ is equal to the cardinality of $H$.

**Definition 2** (Left group action)**.** Let $G$ be a group with $e$ as its identity element and $X$ be a set, then the (left) *group action* $\alpha$ of $G$ on $X$ is the function

$$\alpha : G \times X \to X$$

that obeys the two axioms:

- $\alpha(e, x) = e.x = x$ for all $x \in X$
- $\alpha(g_1, \alpha(g_2, x)) = \alpha(g_1 g_2, x)$ i.e. $g_1.(g_2.x) = (g_1 g_2).x$ for all $x \in X$ and $g_1, g_2 \in G$

**Definition 3** (Orbits)**.** Suppose that $G$ is a group that acts on a set $X$ on the left. Let $x$ be an element of $X$. Then, the *orbit* of $x$ under $G$ is defined by

$$Orb_G(x) = \{g.x \mid g \in G\} = \{y \in X \mid y = g.x \text{ for some } g \in G\}$$

**Definition 4** (Stabilizers)**.** Suppose that $G$ is a group that acts on a set $X$ on the left. Then, for an element $x \in X$, the *stabilizer* of $x$ is defined to be the set

$$Stab_G(x) = \{g \in G \mid g.x = x\}$$

Note: Stabilizers are subgroups of $G$.

**Theorem 9.1** (Orbit-Stabilizer theorem [20])**.** Suppose that $G$ is a finite group acting (on the left) on a set $X$. Then, for any $x \in X$,

$$|Orb_G(x)||Stab_G(x)| = |G|$$

#### Two theorems on non-equivalent permutations of the $k!$ permutations of BFs and their proofs

**Theorem 9.2.** For any $k$-input BF, the total number of non-equivalent permutations divides $k!$.

*Proof.* We approach the problem from a group theoretic perspective. First we define a map

$$\alpha : G \times T \to T$$

where $G$ is the symmetric group $S_k$, *i.e.*, the set of all permutations of $k$ elements ($k$ is the number of inputs to the BF).

We define $T$ as the set of sets by:

$$T = \{t \mid t = \{(X_i, b_i) \mid b_i = 0 \text{ or } 1 \text{ for all } i = 1, 2, ..., 2^k\}\}$$

where, $X_i = (x_1, x_2, ..., x_k)_i$ corresponds to the input values of the $i^{th}$ row of the truth table. Clearly, each element $t \in T$ can be associated with the truth table of a BF of $k$-inputs. Each element of a set $t$ is an ordered pair, the first component denoting some assignment of input variables of the truth table and second component denoting the associated output value. Now, the cardinality of $T$ is $|T| = 2^{2^k}$. For a better understanding of the set $T$ let us take one example. Consider the case of $k = 2$. Then, the set $T$ would be:

$$T = \{t \mid t = \{((x_1, x_2)_i, b_i) \mid b_i = 0 \text{ or } 1 \text{ for all } i = 1, 2, 3, 4\}\}$$

Let us take one element $t$ (for $b_1 = 0,\ b_2 = 1,\ b_3 = 0,\ b_4 = 0$) from the set $T$ as

$$t = \{((0, 0), 0), ((0, 1), 1), ((1, 0), 0), ((1, 1), 0)\}$$

Clearly, one can observe that $t$ is a set that corresponds to the truth table for a specific Boolean function. Each element of $t$ is 2-tuple whose first component is an input assignment and the second component is the associated output. The set $T$ consists of all such possible $t$'s. So, in this case the cardinality of the set $T$ is $2^{2^2} = 16$.

Now we define the map $\alpha : S_k \times T \to T$ by

$$\begin{aligned} \sigma.t &= t' \\ \text{or, } \sigma.\{((x_1, x_2, ..., x_k)_i, b_i) \text{ for } i = 1, 2, ..., 2^k\} &= \{((x_{\sigma(1)}, x_{\sigma(2)}, ..., x_{\sigma(k)})_i, b_i) \text{ for } i = 1, 2, ..., 2^k\} \end{aligned}$$

where, $\sigma \in S_k$ and $t = \{((x_1, x_2, ..., x_k)_i, b_i) \text{ for } i = 1, 2, ..., 2^k\}$, $t' = \{((x_{\sigma(1)}, x_{\sigma(2)}, ..., x_{\sigma(k)})_i, b_i) \text{ for } i = 1, 2, ..., 2^k\} \in T$

Now, we intend to show that the defined map $\alpha$ is a left group action of $S_k$ on $T$. For that, we need to show that the map $\alpha$ obeys the axioms stated in definition 2.

Let, $e \in S_k$ be the identity permutation and $t \in T$. Then,

$$\begin{aligned} & e.t \\ = \; & e.\{((x_1, x_2, ..., x_k)_i, b_i) \text{ for } i = 1, 2, ..., 2^k\} \\ = \; & \{((x_{e(1)}, x_{e(2)}, ..., x_{e(k)})_i, b_i) \text{ for } i = 1, 2, ..., 2^k\} \\ = \; & \{((x_1, x_2, ..., x_k)_i, b_i) \text{ for } i = 1, 2, ..., 2^k\} \\ = \; & t \end{aligned}$$

Again, for $\sigma_1, \sigma_2 \in S_k$ and $t \in T$,

$$\begin{aligned} & \sigma_1.(\sigma_2.t) \\ = \; & \sigma_1.\{((x_{\sigma_2(1)}, x_{\sigma_2(2)}, ..., x_{\sigma_2(k)})_i, b_i) \text{ for } i = 1, 2, ..., 2^k\} \\ = \; & \{((x_{\sigma_1(\sigma_2(1))}, x_{\sigma_1(\sigma_2(2))}, ..., x_{\sigma_1(\sigma_2(k))})_i, b_i) \text{ for } i = 1, 2, ..., 2^k\} \\ = \; & \{((x_{(\sigma_1\sigma_2)(1)}, x_{(\sigma_1\sigma_2)(2)}, ..., x_{(\sigma_1\sigma_2)(k)})_i, b_i) \text{ for } i = 1, 2, ..., 2^k\} \\ = \; & (\sigma_1\sigma_2).\{((x_1, x_2, ..., x_k)_i, b_i) \text{ for } i = 1, 2, ..., 2^k\} \\ = \; & (\sigma_1\sigma_2).t \end{aligned}$$

Hence, the defined map $\alpha$ satisfies both of the conditions for being a group action (left).

Now, the orbit of an element $t \in T$ under $S_k$ is given by (see definition 3)

$$Orb_{S_k}(t) = \{\sigma.t \mid \sigma \in S_k\}$$

So, the set $Orb_{S_k}(t)$ provides all possible elements that correspond to non-equivalent permutations of the BF associated with $t$. In other words, the cardinality of the set $Orb_{S_k}(t)$, $|Orb_{S_k}(t)|$, is the number of non-equivalent permutations of the BF associated with $t$.

Now, by the Orbit-stabilizer (see theorem 9.1) theorem we can say that,

$$\begin{aligned} |Orb_{S_k}(t)||Stab_{S_k}(t)| &= |S_k| \\ \text{or, } |Orb_{S_k}(t)||Stab_{S_k}(t)| &= k! \end{aligned}$$

Hence, $|Orb_{S_k}(t)|$ divides $k!$. In other words, the number of non-equivalent permutations of any k-input BF is always a divisor of k!. This proves the theorem. □

**Theorem 9.3.** In $k!$ permutations of a k-input BF, each of the $m$ non-equivalent permutations occurs an equal number of times.

*Proof.* We consider all the quantities defined for theorem 9.2. We will prove that in the $k!$ possible permutations of a $k$-input BF, each non-equivalent permutation appears $|Stab_{S_k}(t)|$ times where, the BF is associated with the element $t \in T$.

The stabilizer of an element $t \in T$ under $S_k$ is given by (see definition 4)

$$Stab_{S_k}(t) = \{\sigma \in S_k \mid \sigma.t = t\}$$

Evidently, $Stab_{S_k}(t)$ corresponds to the *symmetry* (or *invariance* or *automorphism*) [21] group of the BF. Assume that there are $m$ non-equivalent permutations among the $k!$ permutations of the BF associated with $t$. Then, we can consider the orbit of $t$,

$$Orb_{S_k}(t) = \{t_1, t_2, ..., t_m\}$$

We will show that there are $Stab_{S_k}(t)$ many $\sigma_j$'s for all $j = 1, 2, ..., m$ such that the following conditions hold,

$$\begin{aligned}
\sigma_1.t &= t_1 \\
\sigma_2.t &= t_2 \\
&\vdots \\
\sigma_m.t &= t_m
\end{aligned}$$

The element $t$ always belongs to the set $Orb_{S_k}(t)$ since, $e.t = t$ where $e$ is the identity permutation. So, without loss of generality, let us consider $t_1 = t$. We need to find the number of $\sigma_1$'s such that $\sigma_1.t = t$. From the definition of stabilizer of an element it follows that we have $|Stab_{S_k}(t)|$ many $\sigma_1$'s.

Now, consider an element from $Orb_{S_k}(t)$ different from $t$ (if it exists), say $t_2$. Then, we want to know how many $\sigma_2$'s satisfy

$$\sigma_2.t = t_2 \tag{10}$$

Since $Stab_{S_k}(t)$ is a subgroup of $S_k$, we can consider the following left coset of $Stab_{S_k}(t)$ in $S_k$ with the coset representative $\sigma_2$:

$$\sigma_2 Stab_{S_k}(t) = \{\sigma_2\sigma \mid \sigma \in Stab_{S_k}(t)\}$$

Let us take an element $\sigma_2\sigma \in \sigma_2 Stab_{S_k}(t)$. Then,

$$\begin{aligned}
(\sigma_2\sigma).t &= \sigma_2.(\sigma.t) \\
&= \sigma_2.t \text{ [since, } \sigma \in Stab_{S_k}(t)] \\
&= t_2 \text{ (from 10)}
\end{aligned}$$

Hence, all the elements of $\sigma_2 Stab_{S_k}(t)$ satisfy equation 10.

Now, we will show that all elements $\sigma' \in S_k$ for which equation 10 is satisfied (i.e. $\sigma'.t = t_2$), $\sigma' \in \sigma_2 Stab_{S_k}(t)$. We have,

$$\begin{aligned}
\sigma'.t &= t_2 \\
\text{or, } \sigma'.t &= \sigma_2.t \text{ (from 10)} \\
\text{or, } \sigma'^{-1}.(\sigma'.t) &= \sigma'^{-1}.(\sigma_2.t) \\
\text{or, } (\sigma'^{-1}\sigma').t &= (\sigma'^{-1}\sigma_2).t \\
\text{or, } e.t &= (\sigma'^{-1}\sigma_2).t \\
\text{or, } t &= (\sigma'^{-1}\sigma_2).t
\end{aligned}$$

From the definition of $Stab_{S_k}(t)$ it follows that, $\sigma'^{-1}\sigma_2 \in Stab_{S_k}(t)$. Since $Stab_{S_k}(t)$ is a subgroup of $S_k$, we also have $(\sigma'^{-1}\sigma_2)^{-1} = \sigma_2^{-1}\sigma' \in Stab_{S_k}(t)$. Taking $\sigma_2^{-1}\sigma' = b$, we then have:

$$\begin{aligned}
\sigma' &= \sigma_2 b \text{ where}, b \in Stab_{S_k}(t) \\
\text{or, } \sigma' &\in \sigma_2 Stab_{S_k}(t)
\end{aligned}$$

Hence, the number of $\sigma_2$'s for which equation (10) is satisfied is $|\sigma_2 Stab_{S_k}(t)|$. Now, since $\sigma_2 Stab_{S_k}(t)$ is a left coset of $Stab_{S_k}(t)$, their cardinalities are identical, $|\sigma_2 Stab_{S_k}(t)| = |Stab_{S_k}(t)|$. So, the number of $\sigma_2$'s for which equation (10) is satisfied is precisely $|Stab_{S_k}(t)|$.

In a similar manner, we can show that there are $|Stab_{S_k}(t)|$ many $\sigma_3$'s, $\sigma_4$'s, ..., $\sigma_m$'s such that the conditions $\sigma_3.t = t_3$, $\sigma_4.t = t_4$, ..., $\sigma_m.t = t_m$ respectively are satisfied. Hence, all non-equivalent permutations of a k-input BF associated with $t$, appears $|Stab_{S_k}(t)|$ times in the $k!$ permutations. This proves the theorem. □

### REFERENCES


[1] A. Subbaroyan, O. C. Martin, and A. Samal. Minimum complexity drives regulatory logic in Boolean models of living systems. *PNAS Nexus*, 1(1):pgac017, 2022.

[2] E. Azpeitia, M. Benítez, I. Vega, C. Villarreal, and E. R. Alvarez-Buylla. Single-cell and coupled GRN models of cell patterning in the Arabidopsis thaliana root stem cell niche. *BMC Systems Biology*, 4(1):134, 2010.

[3] J. Davila-Velderrain, J. L. Caldu-Primo, J. C. Martinez-Garcia, and E. R. Alvarez-Buylla. Modeling the Epigenetic Landscape in Plant Development. In *Computational Cell Biology: Methods and Protocols*, pages 357–383. Springer New York, New York, NY, 2018.

[4] E. Sullivan, M. Harris, A. Bhatnagar, E. Guberman, I. Zonfa, and E. Ravasz Regan. Boolean modeling of mechanosensitive epithelial to mesenchymal transition and its reversal. *iScience*, 26(4):106321, 2023.

[5] C. E. Giacomantonio and G. J. Goodhill. A Boolean Model of the Gene Regulatory Network Underlying Mammalian Cortical Area Development. *PLoS Computational Biology*, 6(9):e1000936, 2010.

[6] F. Herrmann, A. Groß, D. Zhou, H. A. Kestler, and M. Kühl. A Boolean Model of the Cardiac Gene Regulatory Network Determining First and Second Heart Field Identity. *PLoS ONE*, 7(10):e46798, 2012.

[7] J. Krumsiek, C. Marr, T. Schroeder, and F. J. Theis. Hierarchical Differentiation of Myeloid Progenitors Is Encoded in the Transcription Factor Network. *PLoS ONE*, 6(8):e22649, 2011.

[8] L. Mendoza and I. Xenarios. A method for the generation of standardized qualitative dynamical systems of regulatory networks. *Theoretical Biology and Medical Modelling*, 3(1):13, 2006.

[9] C. Biane and F. Delaplace. Causal reasoning on boolean control networks based on abduction: theory and application to cancer drug discovery. *IEEE/ACM Transactions on Computational Biology and Bioinformatics*, 16(5):1574–1585, 2019.

[10] J. E. Narváez-Chávez, E. R. Álvarez Buylla, and J. C. Martínez-García. Uncovering the role of mutations in Epithelial-to-Mesenchymal transition through computational analysis of the underlying gene regulatory network. In Hisham Al-Mubaid, Tamer Aldwairi, and Oliver Eulenstein, editors, *Proceedings of International Conference on Bioinformatics and Computational Biology (BICOB-2023)*, volume 92, pages 92–101. EasyChair, 2023.

[11] O. Ríos, S. Frias, A. Rodríguez, S. Kofman, H. Merchant, L. Torres, and L. Mendoza. A Boolean network model of human gonadal sex determination. *Theoretical Biology and Medical Modelling*, 12(1):26, 2015.

[12] A. Wuensche. Complexity in one-D cellular automata: Gliders, basins of attraction and the Z parameter. Working Paper 94-04-025, Santa Fe Institute, 1994.

[13] C. C. Walker and W. R. Ashby. On temporal characteristics of behavior in certain complex systems. *Kybernetik*, 3(2):100–108, 1966.

[14] C. G. Langton. Studying artificial life with cellular automata. *Physica D: Nonlinear Phenomena*, 22(1-3):120–149, 1986.

[15] C. G. Langton. Computation at the edge of chaos: Phase transitions and emergent computation. *Physica D: Nonlinear Phenomena*, 42(1):12–37, 1990.

[16] A. Wuensche and M. Lesser. *Global dynamics of cellular automata: an atlas of basin of attraction fields of one-dimensional cellular automata.* Addison-Wesley, Reading, MA, 1992.

[17] A. Wuensche. *Attractor basins of discrete networks.* D.Phil, The University of Sussex, 1997.

[18] A. Wuensche. Classifying cellular automata automatically: Finding gliders, filtering, and relating space-time patterns, attractor basins, and the Z parameter. *Complexity*, 4(3):47–66, 1999.

[19] S. Nikolajewa, M. Friedel, and T. Wilhelm. Boolean networks with biologically relevant rules show ordered behavior. *Biosystems*, 90(1):40–47, 2007.

[20] J. Gallian. *Contemporary abstract algebra.* Chapman and Hall/CRC, 2021.

[21] P. Clote and E. Kranakis. Boolean functions, invariance groups, and parallel complexity. *SIAM Journal on Computing*, 20(3):553–590, 1991.

TABLE S1. **Boolean functions of the given RSCN-GRN Boolean model in BoolNet format.** The column with header 'Target node name' contains the list of nodes whose regulation is captured by the corresponding row entry in the column 'Regulatory logic rule'. The symbols &, | and ! correspond to the logic operators AND, OR and NOT respectively.

| **Serial Number** | **Target node name** | **Regulatory logic rule** |
|---|---|---|
| 1 | SCR | ( SHR & SCR & !JKD & !MGP ) \| ( SHR & SCR & JKD & !MGP ) \| ( SHR & SCR & JKD & MGP ) |
| 2 | PLT | ARF |
| 3 | ARF | !AUXIAA |
| 4 | AUXIAA | !AUX |
| 5 | AUX | !AUX \| AUX |
| 6 | SHR | SHR |
| 7 | JKD | SHR & SCR |
| 8 | MGP | SHR & SCR & !WOX5 |
| 9 | WOX5 | ( ARF & SHR & SCR & !MGP & !WOX5 ) \| (ARF & SHR & SCR & !MGP & WOX5 ) \| ( ARF & SHR & SCR & MGP & WOX5 ) |

TABLE S2. **Boolean functions of the given EMT-GRN Boolean model in BoolNet format.** The column with header 'Target node name' contains the list of nodes whose regulation is captured by the corresponding row entry in the column 'Regulatory logic rule'. The symbols &, | and ! correspond to the logic operators AND, OR and NOT respectively.

| **Serial Number** | **Target node name** | **Regulatory logic rule** |
|---|---|---|
| 1 | SNAI1 | ((ZEB1_H \| ZEB1) & !miR_34) \| (ZEB1_H & ZEB1) |
| 2 | LEF1 | ((ZEB1 & (!miR_200)) \| ZEB1_H) |
| 3 | Twist | SNAI1 & !miR_34 |
| 4 | SNAI2 | Twist & (SNAI2 \| (N_bcatenin_H & LEF1)) |
| 5 | ZEB1 | SNAI2 \| (b_catenin_TCF4 & (!miR_200)) |
| 6 | N_bcatenin_H | !miR_34 & !miR_200 |
| 7 | ZEB1_H | ZEB1 & ((N_bcatenin_H & LEF1) & (SNAI2 \| (!miR_200))) |
| 8 | b_catenin_TCF4 | N_bcatenin_H & SNAI1 & SNAI2 |
| 9 | miR_34 | (!SNAI1) \| (!(ZEB1 \| ZEB1_H)) |
| 10 | miR_200 | !(Twist & ZEB1_H & SNAI1) & !(ZEB1 & (!miR_200)) |
| 11 | Ecadherin_mRNA | !((((ZEB1_H & ZEB1) & SNAI1) & SNAI2) & Twist) |
| 12 | TGFb_secr | b_catenin_TCF4 & (!miR_200) |

TABLE S3. **Boolean functions of the given NCD-GRN Boolean model in BoolNet format.** The column with header 'Target node name' contains the list of nodes whose regulation is captured by the corresponding row entry in the column 'Regulatory logic rule'. The symbols &, | and ! correspond to the logic operators AND, OR and NOT respectively.

| **Serial Number** | **Target node name** | **Regulatory logic rule** |
|---|---|---|
| 1 | Fgf8_g | Fgf8_p & !Emx2_p & Sp8_p |
| 2 | Fgf8_p | Fgf8_g |
| 3 | Emx2_g | !Fgf8_p & !Pax6_p & Coup_tfi_p & !Sp8_p |
| 4 | Emx2_p | Emx2_g |
| 5 | Pax6_g | !Emx2_p & !Coup_tfi_p & Sp8_p |
| 6 | Pax6_p | Pax6_g |
| 7 | Coup_tfi_g | !Fgf8_p & !Sp8_p |
| 8 | Coup_tfi_p | Coup_tfi_g |
| 9 | Sp8_g | Fgf8_p & !Emx2_p |
| 10 | Sp8_p | Sp8_g |

TABLE S4. **Boolean functions of the given EHD-GRN Boolean model in BoolNet format.** The column with header 'Target node name' contains the list of nodes whose regulation is captured by the corresponding row entry in the column 'Regulatory logic rule'. The symbols &, | and ! correspond to the logic operators AND, OR and NOT respectively.

| Serial Number | Target node name | Regulatory logic rule |
|---|---|---|
| 1 | Bmp2 | (!canWnt & exogen_BMP2_II) |
| 2 | canWnt | exogen_canWnt_II |
| 3 | Dkk1 | (Mesp1 \| (canWnt & !exogen_BMP2_II)) |
| 4 | Fgf8 | (!Mesp1 & (Foxc1/2 \| Tbx1)) |
| 5 | Foxc1/2 | (canWnt & exogen_canWnt_II) |
| 6 | GATAs | (Nkx2_5 \| Mesp1 \| Tbx5) |
| 7 | IsI1 | (Tbx1 \| Mesp1 \| Fgf8 \| (canWnt & exogen_canWnt_II)) |
| 8 | Mesp1 | (canWnt & !exogen_BMP2_II) |
| 9 | Nkx2_5 | ((IsI1 & GATAs) \| Tbx1 \| (Mesp1 & Dkk1) \| (Bmp2 & GATAs) \| Tbx5) |
| 10 | Tbx1 | Foxc1/2 |
| 11 | Tbx5 | (!(Tbx1 \| canWnt) & (Nkx2_5 \| Tbx5 \| Mesp1) & !(Dkk1 & !(Mesp1 \| Tbx5))) |
| 12 | exogen_BMP2_I | 1 |
| 13 | exogen_BMP2_II | exogen_BMP2_I |
| 14 | exogen_canWnt_I | exogen_canWnt_I |
| 15 | exogen_canWnt_II | exogen_canWnt_I |

TABLE S5. **Boolean functions of the given MD-GRN Boolean model in BoolNet format.** The column with header 'Target node name' contains the list of nodes whose regulation is captured by the corresponding row entry in the column 'Regulatory logic rule'. The symbols &, | and ! correspond to the logic operators AND, OR and NOT respectively.

| Serial Number | Target node name | Regulatory logic rule |
|---|---|---|
| 1 | GATA_2 | GATA_2 & !(GATA_1 & FOG_1) & !PU_1 |
| 2 | GATA_1 | (GATA_1 \| GATA_2 \| Fli_1) & !PU_1 |
| 3 | FOG_1 | GATA_1 |
| 4 | EKLF | GATA_1 & !Fli_1 |
| 5 | Fli_1 | GATA_1 & !EKLF |
| 6 | SCL | GATA_1 & !PU_1 |
| 7 | C/EBP_alpha | C/EBP_alpha & !(GATA_1 & FOG_1 & SCL) |
| 8 | PU_1 | (C/EBP_alpha \| PU_1) & !(GATA_1 \| GATA_2) |
| 9 | cJun | PU_1 & !Gfi_1 |
| 10 | EgrNab | (PU_1 & cJun) & !Gfi_1 |
| 11 | Gfi_1 | C/EBP_alpha & !EgrNab |

TABLE S6. **Boolean functions of the given THC-GRN Boolean model in BoolNet format.** The column with header 'Target node name' contains the list of nodes whose regulation is captured by the corresponding row entry in the column 'Regulatory logic rule'. The symbols &, | and ! correspond to the logic operators AND, OR and NOT respectively.

| Serial Number | Target node name | Regulatory logic rule |
|---|---|---|
| 1 | NFAT | TCR |
| 2 | IRAK | IL18R |
| 3 | STAT1 | IFNbR \| JAK1 |
| 4 | IL10R | IL10 |
| 5 | IL12R | IL12 |
| 6 | IL18R | IL18 & !(STAT6) |
| 7 | IFNgR | IFNg |
| 8 | IFNg | (IRAK \| NFAT \| STAT4 \| Tbet) & !(STAT3) |
| 9 | STAT4 | IL12R & !(GATA3) |
| 10 | IL10 | GATA3 |
| 11 | STAT3 | IL10R |
| 12 | IFNbR | IFNb |
| 13 | IL4 | GATA3 & !(STAT1) |
| 14 | IL4R | IL4 & !(SOCS1) |
| 15 | GATA3 | (GATA3 \| STAT6) & !Tbet) |
| 16 | STAT6 | IL4R |
| 17 | SOCS1 | STAT1 \| Tbet |
| 18 | JAK1 | IFNgR & !(SOCS1) |
| 19 | Tbet | (STAT1 \| Tbet) & !(GATA3) |

TABLE S7. **Boolean functions of the given EGFR-GRN Boolean model in BoolNet format.** The column with header 'Target node name' contains the list of nodes whose regulation is captured by the corresponding row entry in the column 'Regulatory logic rule'. The symbols &, | and ! correspond to the logic operators AND, OR and NOT respectively.

| Serial Number | Target node name | Regulatory logic rule |
|---|---|---|
| 1 | EGFR | !BRCA1 |
| 2 | ERK1/2 | EGFR |
| 3 | PI3K | !PTEN & EGFR |
| 4 | AKT | PI3K |
| 5 | GSK3_beta | !AKT |
| 6 | MDM2 | AKT & TP53 |
| 7 | TP53 | !MDM2 & (BRCA1 \| !PARP1) |
| 8 | PTEN | TP53 |
| 9 | PARP1 | ERK1/2 |
| 10 | BRCA1 | !CYCD1 |
| 11 | BCL_2 | AKT |
| 12 | BAX | !BCL_2 & TP53 |
| 13 | CYCD1 | (!GSK3_beta & ERK1/2)\| (!BRCA1 & PARP1) |

TABLE S8. **Boolean functions of the given EMT-Senescence-GRN Boolean model in BoolNet format.** The column with header ‘Target node name’ contains the list of nodes whose regulation is captured by the corresponding row entry in the column ‘Regulatory logic rule’. The symbols &, | and ! correspond to the logic operators AND, OR and NOT respectively.

<table>
<tr><th>Serial Number</th><th>Target node name</th><th>Regulatory logic rule</th></tr>
<tr><td>1</td><td>Snai2</td><td>(Snai2 & NFkB) | !ESE2</td></tr>
<tr><td>2</td><td>ESE2</td><td>!Snai2 | (ESE2 & !NFkB)</td></tr>
<tr><td>3</td><td>p16</td><td>(E2F & p16) | (p16 & !Snai2) | (p53 & !Snai2) | (p53 & !TELasa)</td></tr>
<tr><td>4</td><td>E2F</td><td>!p53 & !Snai2 & !Rb</td></tr>
<tr><td>5</td><td>Cyclin</td><td>(!Snai2 & !p16) | (!ESE2 & !p16 & NFkB)</td></tr>
<tr><td>6</td><td>TELasa</td><td>!ESE2</td></tr>
<tr><td>7</td><td>NFkB</td><td>Snai2 | ESE2 | p16 | NFkB</td></tr>
<tr><td>8</td><td>Rb</td><td>p53 | (p16 & Cyclin)</td></tr>
<tr><td>9</td><td>p53</td><td>(!Snai2 & p16 & !TELasa) | (!Snai2 & !TELasa & !NFkB)</td></tr>
</table>

TABLE S9. **Boolean functions of the given FOS-GRN Boolean model in BoolNet format.** The column with header ‘Target node name’ contains the list of nodes whose regulation is captured by the corresponding row entry in the column ‘Regulatory logic rule’. The symbols &, | and ! correspond to the logic operators AND, OR and NOT respectively.

<table>
<tr><th>Serial Number</th><th>Target node name</th><th>Regulatory logic rule</th></tr>
<tr><td>1</td><td>AG</td><td>(!EMF1 & !AP2 & !TFL1) | (!EMF1 & !AP1 & LFY) | (!EMF1 & !AP2 & LFY) | (!EMF1 & !TFL1 & LFY & (AG & SEP)) | (!EMF1 & (LFY & WUS))</td></tr>
<tr><td>2</td><td>AP1</td><td>(!AG & !TFL1) | (FT & LFY & !AG) | (FT & !AG & !PI) | (LFY & !AG & !PI) | (FT & !AG & !AP3) | (LFY & !AG & !AP3)</td></tr>
<tr><td>3</td><td>AP2</td><td>!TFL1</td></tr>
<tr><td>4</td><td>AP3</td><td>(LFY & UFO) | (PI & SEP & AP3 & (AG | AP1))</td></tr>
<tr><td>5</td><td>EMF1</td><td>!LFY</td></tr>
<tr><td>6</td><td>FT</td><td>!EMF1</td></tr>
<tr><td>7</td><td>FUL</td><td>!AP1 & !TFL1</td></tr>
<tr><td>8</td><td>LFY</td><td>(!EMF1) | (!TFL1)</td></tr>
<tr><td>9</td><td>PI</td><td>(LFY & (AG | AP3)) | (PI & SEP & AP3 & (AG | AP1))</td></tr>
<tr><td>10</td><td>SEP</td><td>LFY</td></tr>
<tr><td>11</td><td>TFL1</td><td>!AP1 & (EMF1 & !LFY)</td></tr>
<tr><td>12</td><td>UFO</td><td>UFO</td></tr>
<tr><td>13</td><td>WUS</td><td>WUS & (!AG | !SEP)</td></tr>
</table>

TABLE S10. **Boolean functions of the given GSD-GRN Boolean model in BoolNet format.** The column with header ‘Target node name’ contains the list of nodes whose regulation is captured by the corresponding row entry in the column ‘Regulatory logic rule’. The symbols &, | and ! correspond to the logic operators AND, OR and NOT respectively.

<table>
<tr><th>Serial Number</th><th>Target node name</th><th>Regulatory logic rule</th></tr>
<tr><td>1</td><td>UGR</td><td>UGR & !(NR5A1 | WNT4)</td></tr>
<tr><td>2</td><td>CBX2</td><td>UGR & !(NR0B1 & WNT4 & CTNNB1)</td></tr>
<tr><td>3</td><td>GATA4</td><td>(UGR | WNT4 | NR5A1 | SRY)</td></tr>
<tr><td>4</td><td>WT1mKTS</td><td>(UGR | GATA4)</td></tr>
<tr><td>5</td><td>WT1pKTS</td><td>(UGR | GATA4) & !(WNT4 & CTNNB1)</td></tr>
<tr><td>6</td><td>NR5A1</td><td>(UGR | CBX2 | WT1mKTS | GATA4) & !(NR0B1 & WNT4)</td></tr>
<tr><td>7</td><td>NR0B1</td><td>(WT1mKTS | (WNT4 & CTNNB1)) & !(NR5A1 & SOX9)</td></tr>
<tr><td>8</td><td>SRY</td><td>((NR5A1 & WT1mKTS & CBX2) | (GATA4 & WT1pKTS & CBX2 & NR5A1) | (SOX9 | SRY)) & !(CTNNB1)</td></tr>
<tr><td>9</td><td>SOX9</td><td>((SOX9 & FGF9) | (SRY | PGD2) | (SRY & CBX2) | (GATA4 & NR5A1 & SRY)) & !(WNT4 | CTNNB1 | FOXL2)</td></tr>
<tr><td>10</td><td>FGF9</td><td>SOX9 & !WNT4</td></tr>
<tr><td>11</td><td>PGD2</td><td>SOX9</td></tr>
<tr><td>12</td><td>DMRT1</td><td>(SRY | SOX9) & !(FOXL2)</td></tr>
<tr><td>13</td><td>DHH</td><td>SOX9</td></tr>
<tr><td>14</td><td>DKK1</td><td>(SRY | SOX9)</td></tr>
<tr><td>15</td><td>AMH</td><td>((SOX9 & GATA4 & NR5A1) | (SOX9 & NR5A1 & GATA4 & WT1mKTS)) & !(NR0B1 & CTNNB1)</td></tr>
<tr><td>16</td><td>WNT4</td><td>(GATA4 | (CTNNB1 | RSPO1 | NR0B1)) & !(FGF9 | DKK1)</td></tr>
<tr><td>17</td><td>RSPO1</td><td>(WNT4 | CTNNB1) & !(DKK1)</td></tr>
<tr><td>18</td><td>FOXL2</td><td>(WNT4 & CTNNB1) & !(DMRT1 | SOX9)</td></tr>
<tr><td>19</td><td>CTNNB1</td><td>(WNT4 | RSPO1) & !(SRY | (SOX9 & AMH))</td></tr>
</table>

TABLE S11. **Type of the BF at each node of the given RSCN-GRN Boolean model.** The ‘Node’ column contains all the nodes of the network. An entry in the columns corresponding to BF types, namely EF, UF, NCF and RoF, is ‘Yes’ if the BF in the original model belongs to that function type and is ‘No’ if it does not belong to that function type.

| Serial Number | Node | EF | UF | NCF | RoF |
|---|---|---|---|---|---|
| 1 | SCR | Yes | Yes | Yes | Yes |
| 2 | PLT | Yes | Yes | Yes | Yes |
| 3 | ARF | Yes | Yes | Yes | Yes |
| 4 | AUXIAA | Yes | Yes | Yes | Yes |
| 5 | AUX | No | Yes | No | No |
| 6 | SHR | Yes | Yes | Yes | Yes |
| 7 | JKD | Yes | Yes | Yes | Yes |
| 8 | MGP | Yes | Yes | Yes | Yes |
| 9 | WOX5 | Yes | Yes | Yes | Yes |

TABLE S12. **Type of the BF at each node of the given EMT-GRN Boolean model.** The 'Node' column contains all the nodes of the network. An entry in the columns corresponding to BF types, namely EF, UF, NCF and RoF, is 'Yes' if the BF in the original model belongs to that function type and is 'No' if it does not belong to that function type.

| **Serial Number** | **Node** | **EF** | **UF** | **NCF** | **RoF** |
|---|---|---|---|---|---|
| 1 | SNAI1 | Yes | Yes | No | No |
| 2 | LEF1 | Yes | Yes | Yes | Yes |
| 3 | Twist | Yes | Yes | Yes | Yes |
| 4 | SNAI2 | Yes | Yes | Yes | Yes |
| 5 | ZEB1 | Yes | Yes | Yes | Yes |
| 6 | N_bcatenin_H | Yes | Yes | Yes | Yes |
| 7 | ZEB1_H | Yes | Yes | Yes | Yes |
| 8 | b_catenin_TCF4 | Yes | Yes | Yes | Yes |
| 9 | miR_34 | Yes | Yes | Yes | Yes |
| 10 | miR_200 | Yes | Yes | No | Yes |
| 11 | Ecadherin_mRNA | Yes | Yes | Yes | Yes |
| 12 | TGFb_secr | Yes | Yes | Yes | Yes |

TABLE S13. **Type of the BF at each node of the given NCD-GRN Boolean model.** The 'Node' column contains all the nodes of the network. An entry in the columns corresponding to BF types, namely EF, UF, NCF and RoF, is 'Yes' if the BF in the original model belongs to that function type and is 'No' if it does not belong to that function type.

| **Serial Number** | **Node** | **EF** | **UF** | **NCF** | **RoF** |
|---|---|---|---|---|---|
| 1 | Fgf8_g | Yes | Yes | Yes | Yes |
| 2 | Fgf8_p | Yes | Yes | Yes | Yes |
| 3 | Emx2_g | Yes | Yes | Yes | Yes |
| 4 | Emx2_p | Yes | Yes | Yes | Yes |
| 5 | Pax6_g | Yes | Yes | Yes | Yes |
| 6 | Pax6_p | Yes | Yes | Yes | Yes |
| 7 | Coup_tfi_g | Yes | Yes | Yes | Yes |
| 8 | Coup_tfi_p | Yes | Yes | Yes | Yes |
| 9 | Sp8_g | Yes | Yes | Yes | Yes |
| 10 | Sp8_p | Yes | Yes | Yes | Yes |

TABLE S14. **Type of the BF at each node of the given EHD-GRN Boolean model.** The 'Node' column contains all the nodes of the network. An entry in the columns corresponding to BF types, namely EF, UF, NCF and RoF, is 'Yes' if the BF in the original model belongs to that function type and is 'No' if it does not belong to that function type.

| **Serial Number** | **Node** | **EF** | **UF** | **NCF** | **RoF** |
|---|---|---|---|---|---|
| 1 | Bmp2 | Yes | Yes | Yes | Yes |
| 2 | canWnt | Yes | Yes | Yes | Yes |
| 3 | Dkk1 | Yes | Yes | Yes | Yes |
| 4 | Fgf8 | Yes | Yes | Yes | Yes |
| 5 | Foxc1/2 | Yes | Yes | Yes | Yes |
| 6 | GATAs | Yes | Yes | Yes | Yes |
| 7 | IsI1 | Yes | Yes | Yes | Yes |
| 8 | Mesp1 | Yes | Yes | Yes | Yes |
| 9 | Nkx2_5 | Yes | Yes | No | Yes |
| 10 | Tbx1 | Yes | Yes | Yes | Yes |
| 11 | Tbx5 | Yes | Yes | Yes | Yes |
| 12 | exogen_BMP2_I | - | - | - | - |
| 13 | exogen_BMP2_II | Yes | Yes | Yes | Yes |
| 14 | exogen_canWnt_I | Yes | Yes | Yes | Yes |
| 15 | exogen_canWnt_II | Yes | Yes | Yes | Yes |

TABLE S15. **Type of the BF at each node of the given MD-GRN Boolean model.** The 'Node' column contains all the nodes of the network. An entry in the columns corresponding to BF types, namely EF, UF, NCF and RoF, is 'Yes' if the BF in the original model belongs to that function type and is 'No' if it does not belong to that function type.

| **Serial Number** | **Node** | **EF** | **UF** | **NCF** | **RoF** |
|---|---|---|---|---|---|
| 1 | GATA_2 | Yes | Yes | Yes | Yes |
| 2 | GATA_1 | Yes | Yes | Yes | Yes |
| 3 | FOG_1 | Yes | Yes | Yes | Yes |
| 4 | EKLF | Yes | Yes | Yes | Yes |
| 5 | Fli_1 | Yes | Yes | Yes | Yes |
| 6 | SCL | Yes | Yes | Yes | Yes |
| 7 | C/EBP_alpha | Yes | Yes | Yes | Yes |
| 8 | PU_1 | Yes | Yes | Yes | Yes |
| 9 | cJun | Yes | Yes | Yes | Yes |
| 10 | EgrNab | Yes | Yes | Yes | Yes |
| 11 | Gfi_1 | Yes | Yes | Yes | Yes |

TABLE S16. **Type of the BF at each node of the given THC-GRN Boolean model.** The 'Node' column contains all the nodes of the network. An entry in the columns corresponding to BF types, namely EF, UF, NCF and RoF, is 'Yes' if the BF in the original model belongs to that function type and is 'No' if it does not belong to that function type.

| **Serial Number** | **Node** | **EF** | **UF** | **NCF** | **RoF** |
|---|---|---|---|---|---|
| 1 | NFAT | Yes | Yes | Yes | Yes |
| 2 | IRAK | Yes | Yes | Yes | Yes |
| 3 | STAT1 | Yes | Yes | Yes | Yes |
| 4 | IL10R | Yes | Yes | Yes | Yes |
| 5 | IL12R | Yes | Yes | Yes | Yes |
| 6 | IL18R | Yes | Yes | Yes | Yes |
| 7 | IFNgR | Yes | Yes | Yes | Yes |
| 8 | IFNg | Yes | Yes | Yes | Yes |
| 9 | STAT4 | Yes | Yes | Yes | Yes |
| 10 | IL10 | Yes | Yes | Yes | Yes |
| 11 | STAT3 | Yes | Yes | Yes | Yes |
| 12 | IFNbR | Yes | Yes | Yes | Yes |
| 13 | IL4 | Yes | Yes | Yes | Yes |
| 14 | IL4R | Yes | Yes | Yes | Yes |
| 15 | GATA3 | Yes | Yes | Yes | Yes |
| 16 | STAT6 | Yes | Yes | Yes | Yes |
| 17 | SOCS1 | Yes | Yes | Yes | Yes |
| 18 | JAK1 | Yes | Yes | Yes | Yes |
| 19 | Tbet | Yes | Yes | Yes | Yes |

TABLE S17. **Type of the BF at each node of the given EGFR-GRN Boolean model.** The 'Node' column contains all the nodes of the network. An entry in the columns corresponding to BF types, namely EF, UF, NCF and RoF, is 'Yes' if the BF in the original model belongs to that function type and is 'No' if it does not belong to that function type.

| **Serial Number** | **Node** | **EF** | **UF** | **NCF** | **RoF** |
|---|---|---|---|---|---|
| 1 | EGFR | Yes | Yes | Yes | Yes |
| 2 | ERK1/2 | Yes | Yes | Yes | Yes |
| 3 | PI3K | Yes | Yes | Yes | Yes |
| 4 | AKT | Yes | Yes | Yes | Yes |
| 5 | GSK3_beta | Yes | Yes | Yes | Yes |
| 6 | MDM2 | Yes | Yes | Yes | Yes |
| 7 | TP53 | Yes | Yes | Yes | Yes |
| 8 | PTEN | Yes | Yes | Yes | Yes |
| 9 | PARP1 | Yes | Yes | Yes | Yes |
| 10 | BRCA1 | Yes | Yes | Yes | Yes |
| 11 | BCL_2 | Yes | Yes | Yes | Yes |
| 12 | BAX | Yes | Yes | Yes | Yes |
| 13 | CYCD1 | Yes | Yes | No | Yes |

TABLE S18. **Type of the BF at each node of the given EMT-Senescence-GRN Boolean model.** The 'Node' column contains all the nodes of the network. An entry in the columns corresponding to BF types, namely EF, UF, NCF and RoF, is 'Yes' if the BF in the original model belongs to that function type and is 'No' if it does not belong to that function type.

| Serial Number | Node | EF | UF | NCF | RoF |
|---|---|---|---|---|---|
| 1 | Snai2 | Yes | Yes | Yes | Yes |
| 2 | ESE2 | Yes | Yes | Yes | Yes |
| 3 | p16 | Yes | Yes | No | No |
| 4 | E2F | Yes | Yes | Yes | Yes |
| 5 | Cyclin | Yes | Yes | Yes | Yes |
| 6 | TELasa | Yes | Yes | Yes | Yes |
| 7 | NFkB | Yes | Yes | Yes | Yes |
| 8 | Rb | Yes | Yes | Yes | Yes |
| 9 | p53 | Yes | Yes | Yes | Yes |

TABLE S19. **Type of the BF at each node of the given FOS-GRN Boolean model.** The 'Node' column contains all the nodes of the network. An entry in the columns corresponding to BF types, namely EF, UF, NCF and RoF, is 'Yes' if the BF in the original model belongs to that function type and is 'No' if it does not belong to that function type.

| Serial Number | Node | EF | UF | NCF | RoF |
|---|---|---|---|---|---|
| 1 | AG | Yes | Yes | No | No |
| 2 | AP1 | Yes | Yes | No | No |
| 3 | AP2 | Yes | Yes | Yes | Yes |
| 4 | AP3 | Yes | Yes | No | Yes |
| 5 | EMF1 | Yes | Yes | Yes | Yes |
| 6 | FT | Yes | Yes | Yes | Yes |
| 7 | FUL | Yes | Yes | Yes | Yes |
| 8 | LFY | Yes | Yes | Yes | Yes |
| 9 | PI | Yes | Yes | No | No |
| 10 | SEP | Yes | Yes | Yes | Yes |
| 11 | TFL1 | Yes | Yes | Yes | Yes |
| 12 | UFO | Yes | Yes | Yes | Yes |
| 13 | WUS | Yes | Yes | Yes | Yes |

TABLE S20. **Type of the BF at each node of the given GSD-GRN Boolean model.** The 'Node' column contains all the nodes of the network. An entry in the columns corresponding to BF types, namely EF, UF, NCF and RoF, is 'Yes' if the BF in the original model belongs to that function type and is 'No' if it does not belong to that function type.

| Serial Number | Node | EF | UF | NCF | RoF |
|---|---|---|---|---|---|
| 1 | UGR | Yes | Yes | Yes | Yes |
| 2 | CBX2 | Yes | Yes | Yes | Yes |
| 3 | GATA4 | Yes | Yes | Yes | Yes |
| 4 | WT1mKTS | Yes | Yes | Yes | Yes |
| 5 | WT1pKTS | Yes | Yes | No | Yes |
| 6 | NR5A1 | Yes | Yes | No | Yes |
| 7 | NR0B1 | Yes | Yes | No | Yes |
| 8 | SRY | Yes | Yes | Yes | Yes |
| 9 | SOX9 | No | Yes | No | No |
| 10 | FGF9 | Yes | Yes | Yes | Yes |
| 11 | PGD2 | Yes | Yes | Yes | Yes |
| 12 | DMRT1 | Yes | Yes | Yes | Yes |
| 13 | DHH | Yes | Yes | Yes | Yes |
| 14 | DKK1 | Yes | Yes | Yes | Yes |
| 15 | AMH | No | Yes | No | No |
| 16 | WNT4 | Yes | Yes | Yes | Yes |
| 17 | RSPO1 | Yes | Yes | Yes | Yes |
| 18 | FOXL2 | Yes | Yes | Yes | Yes |
| 19 | CTNNB1 | Yes | Yes | No | Yes |

TABLE S21. **Biological fixed points recovered by the RSCN-GRN Boolean model.** The 4 biological fixed points recovered by the model are the Quiescent center (QC), Vascular initials (VI), Cortex-Endodermis initials (CEI) and Columella epidermis initials (CEpI)

| Cell types | Nodes | | | | | | | | |
|---|---|---|---|---|---|---|---|---|---|
| | SCR | PLT | ARF | AUXIAA | AUXIN | SHR | JKD | MGP | WOX5 |
| QC | 1 | 1 | 1 | 0 | 1 | 1 | 1 | 0 | 1 |
| VI | 0 | 1 | 1 | 0 | 1 | 1 | 0 | 0 | 0 |
| CEI | 1 | 1 | 1 | 0 | 1 | 1 | 1 | 1 | 0 |
| CEpI | 0 | 1 | 1 | 0 | 1 | 0 | 0 | 0 | 0 |

TABLE S22. **Biological fixed points recovered by the EMT-GRN Boolean model.** The 3 biological fixed points recovered by the model are the cell types: Epithelial, Mesenchymal and Hybrid

| Cell types | Nodes | | | | | | | | | | | |
|---|---|---|---|---|---|---|---|---|---|---|---|---|
| | SNAI1 | LEF1 | Twist | SNAI2 | ZEB1 | N_bcatenin_H | ZEB1_H | b_catenin_TCF4 | miR_34 | miR_200 | Ecadherin_mRNA | TGFb_secr |
| Epithelial | 0 | 0 | 0 | 0 | 0 | 0 | 0 | 0 | 1 | 1 | 1 | 0 |
| Hybrid | 1 | 0 | 1 | 1 | 1 | 0 | 0 | 0 | 0 | 1 | 1 | 0 |
| Mesenchymal | 1 | 1 | 1 | 1 | 1 | 1 | 1 | 1 | 0 | 0 | 0 | 1 |

TABLE S23. **Biological fixed points recovered by the NCD-GRN Boolean model.** The 2 biological fixed points recovered by the model are the Anterior and Posterior compartments of the developing neocortex.

| Cell types | Nodes | | | | | | | | | |
|---|---|---|---|---|---|---|---|---|---|---|
| | Fgf8_g | Fgf8_p | Emx2_g | Emx2_p | Pax6_g | Pax6_p | Coup_tfi_g | Coup_tfi_p | Sp8_g | Sp8_p |
| Anterior | 1 | 1 | 0 | 0 | 1 | 1 | 0 | 0 | 1 | 1 |
| Posterior | 0 | 0 | 1 | 1 | 0 | 0 | 1 | 1 | 0 | 0 |

TABLE S24. **Biological fixed points recovered by the EHD-GRN Boolean model.** The 2 biological fixed points recovered by the model are the cell types in the First Heart Field (FHF) and Second Heart Field (SHF).

| Cell types | Nodes | | | | | | | | | | | | | | |
|---|---|---|---|---|---|---|---|---|---|---|---|---|---|---|---|
| | Bmp2 | canWnt | Dkk1 | Fgf8 | Foxc1/2 | GATAs | Isl1 | Mesp1 | Nkx2_5 | Tbx1 | Tbx5 | exogen_BMP2_I | exogen_BMP2_II | exogen_canWnt_I | exogen_canWnt_II |
| FHF | 1 | 0 | 0 | 0 | 0 | 1 | 0 | 0 | 1 | 0 | 1 | 1 | 1 | 0 | 0 |
| SHF | 0 | 1 | 0 | 1 | 1 | 1 | 1 | 0 | 1 | 1 | 0 | 1 | 1 | 1 | 1 |

TABLE S25. **Biological fixed points recovered by the MD-GRN Boolean model.** The 4 biological fixed points recovered by the model correspond to the cell types: erythrocytes, megakaryocytes, monocytes and granulocytes.

| Cell types | Nodes | | | | | | | | | | |
|---|---|---|---|---|---|---|---|---|---|---|---|
| | GATA_2 | GATA_1 | FOG_1 | EKLF | Fli_1 | SCL | C/EBP_alpha | PU_1 | cJun | EgrNab | Gfi_1 |
| Erythrocyte | 0 | 1 | 1 | 1 | 0 | 1 | 0 | 0 | 0 | 0 | 0 |
| Megakaryocyte | 0 | 1 | 1 | 0 | 1 | 1 | 0 | 0 | 0 | 0 | 0 |
| Monocyte | 0 | 0 | 0 | 0 | 0 | 0 | 1 | 1 | 1 | 1 | 0 |
| Granulocyte | 0 | 0 | 0 | 0 | 0 | 0 | 1 | 1 | 0 | 0 | 1 |

TABLE S26. **Biological fixed points recovered by the THC-GRN Boolean model.** The 3 biological fixed points recovered by the model correspond to different types of T-helper cells, namely, Th0, Th1 and Th2.

| Cell types | Nodes | | | | | | | | | | | | | | | | | | | | | |
|---|---|---|---|---|---|---|---|---|---|---|---|---|---|---|---|---|---|---|---|---|---|---|
| | NFAT | IRAK | STAT1 | IL10R | IL12R | IL18R | IFNgR | IFNg | STAT4 | IL10 | STAT3 | IFNbR | IL4 | IL4R | GATA3 | STAT6 | SOCS1 | JAK1 | Tbet | IFNb | TCR | IL18 | IL12 |
| Th0 | 0 | 0 | 0 | 0 | 0 | 0 | 0 | 0 | 0 | 0 | 0 | 0 | 0 | 0 | 0 | 0 | 0 | 0 | 0 | 0 | 0 | 0 | 0 |
| Th1 | 0 | 0 | 0 | 0 | 0 | 0 | 1 | 1 | 0 | 0 | 0 | 0 | 0 | 0 | 0 | 0 | 1 | 0 | 1 | 0 | 0 | 0 | 0 |
| Th2 | 0 | 0 | 0 | 1 | 0 | 0 | 0 | 0 | 0 | 1 | 1 | 0 | 1 | 1 | 1 | 1 | 0 | 0 | 0 | 0 | 0 | 0 | 0 |

TABLE S27. **Biological fixed points recovered by the EGFR-GRN Boolean model.** The 2 biological fixed points recovered by the model correspond to the cell states: cell division and apoptosis.

| Cell types | Nodes | | | | | | | | | | | | |
|---|---|---|---|---|---|---|---|---|---|---|---|---|---|
| | EGFR | ERK1/2 | PI3K | AKT | GSK3_beta | MDM2 | TP53 | PTEN | PARP1 | BRCA1 | BCL_2 | BAX | CYCD1 |
| Division | 1 | 1 | 1 | 1 | 0 | 0 | 0 | 0 | 1 | 0 | 1 | 0 | 1 |
| Apoptosis | 0 | 0 | 0 | 0 | 1 | 0 | 1 | 1 | 0 | 1 | 0 | 1 | 0 |

TABLE S28. **Biological fixed points recovered by the EMT-Senescence-GRN Boolean model.** The 3 biological fixed points recovered by the model correspond to epithelial cells, senescent epithelial cells and stem-like mesenchymal cells.

| Cell types | Nodes | | | | | | | | |
|---|---|---|---|---|---|---|---|---|---|
| | Snai2 | ESE2 | p16 | E2F | Cyclin | TELasa | NFkB | Rb | p53 |
| Epithelial | 0 | 1 | 0 | 1 | 1 | 0 | 1 | 0 | 0 |
| Mesenchymal | 1 | 0 | 0 | 0 | 1 | 1 | 1 | 0 | 0 |
| Senescent | 0 | 1 | 1 | 0 | 0 | 0 | 1 | 1 | 1 |

TABLE S29. **Biological fixed points recovered by the FOS-GRN Boolean model.** The 10 biological fixed points recovered by the model correspond to inflorescence (4 cell types), sepal (1 cell type), petal (2 cell types), stamen (2 cell types) and carpel (1 cell type).

| Cell types | Nodes | | | | | | | | | | | | |
|---|---|---|---|---|---|---|---|---|---|---|---|---|---|
| | AG | AP1 | AP2 | AP3 | EMF1 | FT | FUL | LFY | PI | SEP | TFL1 | UFO | WUS |
| INF1 | 0 | 0 | 0 | 0 | 1 | 0 | 0 | 0 | 0 | 0 | 1 | 0 | 0 |
| INF2 | 0 | 0 | 0 | 0 | 1 | 0 | 0 | 0 | 0 | 0 | 1 | 1 | 0 |
| INF3 | 0 | 0 | 0 | 0 | 1 | 0 | 0 | 0 | 0 | 0 | 1 | 0 | 1 |
| INF4 | 0 | 0 | 0 | 0 | 1 | 0 | 0 | 0 | 0 | 0 | 1 | 1 | 1 |
| SEP | 0 | 1 | 1 | 0 | 0 | 1 | 0 | 1 | 0 | 1 | 0 | 0 | 0 |
| PET1 | 0 | 1 | 1 | 1 | 0 | 1 | 0 | 1 | 1 | 1 | 0 | 1 | 0 |
| PET2 | 0 | 1 | 1 | 1 | 0 | 1 | 0 | 1 | 1 | 1 | 0 | 0 | 0 |
| STM1 | 1 | 0 | 1 | 1 | 0 | 1 | 1 | 1 | 1 | 1 | 0 | 1 | 0 |
| STM2 | 1 | 0 | 1 | 1 | 0 | 1 | 1 | 1 | 1 | 1 | 0 | 0 | 0 |
| CAR | 1 | 0 | 1 | 0 | 0 | 1 | 1 | 1 | 1 | 1 | 0 | 0 | 0 |

TABLE S30. **Biological fixed points recovered by the GSD-GRN Boolean model.** The 2 biological fixed points recovered by the model correspond to the Sertoli progenitor cells (SPC) and granulosa progenitor cells (GPC).

| Cell types | Nodes | | | | | | | | | | | | | | | | | | |
|---|---|---|---|---|---|---|---|---|---|---|---|---|---|---|---|---|---|---|---|
| | UGR | CBX2 | GATA4 | WT1mKTS | WT1pKTS | NR5A1 | NR0B1 | SRY | SOX9 | FGF9 | PGD2 | DMRT1 | DHH | DKK1 | AMH | WNT4 | RSPO1 | FOXL2 | CTNNB1 |
| Toward_Sertoli | 0 | 0 | 1 | 1 | 1 | 1 | 0 | 1 | 1 | 1 | 1 | 1 | 1 | 1 | 1 | 0 | 0 | 0 | 0 |
| Toward_granulosa | 0 | 0 | 1 | 1 | 0 | 0 | 1 | 0 | 0 | 0 | 0 | 0 | 0 | 0 | 0 | 1 | 1 | 1 | 1 |

TABLE S31. **Nodewise enumeration of the number of BFs that satisfy various biologically meaningful constraints for the RSCN-GRN model.** The 'Node' column contains all nodes of the network. The '$k$' column is the number of effective inputs to each node. Each of the columns EF, EUF, RoF and NCF provides the numbers of the respective type of BF that satisfy the fixed point constraints for each of the nodes. Additionally, the BFs in EUF, RoF and NCF also satisfy the signs of the inputs in the network architecture. The row labeled 'Total' gives the total number of Boolean models allowed when imposing a given type of BF at each node. Here and in the previous rows, '-' denotes that it was computationally infeasible to obtain the exact values. Finally, the row labeled 'Sampled' specifies whether we had to resort to sampling of these models. When set to False, we considered all possible models by exhaustive enumeration.

| **Serial Number** | **Node** | $k$ | **EF** | **EUF** | **RoF** | **NCF** |
|---|---|---|---|---|---|---|
| 1 | SCR | 4 | 4010 | 50 | 20 | 17 |
| 2 | PLT | 1 | 1 | 1 | 1 | 1 |
| 3 | ARF | 1 | 1 | 1 | 1 | 1 |
| 4 | AUXIAA | 1 | 1 | 1 | 1 | 1 |
| 5 | AUXIN | 1 | 1 | 1 | 1 | 1 |
| 6 | SHR | 1 | 1 | 1 | 1 | 1 |
| 7 | JKD | 2 | 1 | 1 | 1 | 1 |
| 8 | MGP | 3 | 14 | 1 | 1 | 1 |
| 9 | WOX5 | 5 | - | 732 | 94 | 75 |
| Total | | | - | 36600 | 1880 | 1275 |
| Sampled | | | True | False | False | False |

TABLE S32. **Nodewise enumeration of the number of BFs that satisfy various biologically meaningful constraints for the EMT-GRN model.** The 'Node' column contains all nodes of the network. The '$k$' column is the number of effective inputs to each node. Each of the columns EF, EUF, RoF and NCF provides the numbers of the respective type of BF that satisfy the fixed point constraints for each of the nodes. Additionally, the BFs in EUF, RoF and NCF also satisfy the signs of the inputs in the network architecture. The row labeled 'Total' gives the total number of Boolean models allowed when imposing a given type of BF at each node. Here and in the previous rows, '-' denotes that it was computationally infeasible to obtain the exact values. Finally, the row labeled 'Sampled' specifies whether we had to resort to sampling of these models. When set to False, we considered all possible models by exhaustive enumeration.

| **Serial Number** | **Node** | $k$ | **EF** | **EUF** | **RoF** | **NCF** |
|---|---|---|---|---|---|---|
| 1 | SNAI1 | 3 | 26 | 7 | 6 | 6 |
| 2 | LEF1 | 3 | 26 | 7 | 6 | 6 |
| 3 | Twist | 2 | 2 | 2 | 2 | 2 |
| 4 | SNAI2 | 4 | 8074 | 57 | 26 | 23 |
| 5 | ZEB1 | 3 | 29 | 2 | 2 | 2 |
| 6 | N_bcatenin_H | 2 | 1 | 1 | 1 | 1 |
| 7 | ZEB1_H | 5 | - | 5119 | 310 | 213 |
| 8 | b_catenin_TCF4 | 3 | 29 | 2 | 2 | 2 |
| 9 | miR_34 | 3 | 26 | 7 | 6 | 6 |
| 10 | miR_200 | 5 | - | 1775 | 162 | 119 |
| 11 | Ecadherin_mRNA | 5 | - | 114 | 52 | 46 |
| 12 | TGFb_secr | 2 | 2 | 2 | 2 | 2 |
| Total | | | - | 324024087794400 | 234653552640 | 92679987456 |
| Sampled | | | True | True | True | True |

TABLE S33. **Nodewise enumeration of the number of BFs that satisfy various biologically meaningful constraints for the NCD-GRN model.** The 'Node' column contains all nodes of the network. The '$k$' column is the number of effective inputs to each node. Each of the columns EF, EUF, RoF and NCF provides the numbers of the respective type of BF that satisfy the fixed point constraints for each of the nodes. Additionally, the BFs in EUF, RoF and NCF also satisfy the signs of the inputs in the network architecture. The row labeled 'Total' gives the total number of Boolean models allowed when imposing a given type of BF at each node. Finally, the row labeled 'Sampled' specifies whether we had to resort to sampling of these models. When set to False, we considered all possible models by exhaustive enumeration.

| **Serial Number** | **Node** | $k$ | **EF** | **EUF** | **RoF** | **NCF** |
|---|---|---|---|---|---|---|
| 1 | Fgf8_g | 3 | 55 | 9 | 8 | 8 |
| 2 | Fgf8_p | 1 | 1 | 1 | 1 | 1 |
| 3 | Emx2_g | 4 | 16148 | 114 | 52 | 46 |
| 4 | Emx2_p | 1 | 1 | 1 | 1 | 1 |
| 5 | Pax6_g | 3 | 55 | 9 | 8 | 8 |
| 6 | Pax6_p | 1 | 1 | 1 | 1 | 1 |
| 7 | Coup_tfi_g | 2 | 2 | 2 | 2 | 2 |
| 8 | Coup_tfi_p | 1 | 1 | 1 | 1 | 1 |
| 9 | Sp8_g | 2 | 2 | 2 | 2 | 2 |
| 10 | Sp8_p | 1 | 1 | 1 | 1 | 1 |
| Total | | | 195390800 | 36936 | 13312 | 11776 |
| Sampled | | | True | False | False | False |

TABLE S34. **Nodewise enumeration of the number of BFs that satisfy various biologically meaningful constraints for the EHD-GRN model.** The 'Node' column contains all nodes of the network. The '$k$' column is the number of effective inputs to each node. Each of the columns EF, EUF, RoF and NCF provides the numbers of the respective type of BF that satisfy the fixed point constraints for each of the nodes. Additionally, the BFs in EUF, RoF and NCF also satisfy the signs of the inputs in the network architecture. The row labeled 'Total' gives the total number of Boolean models allowed when imposing a given type of BF at each node. Here and in the previous rows, '-' denotes that it was computationally infeasible to obtain the exact values. Finally, the row labeled 'Sampled' specifies whether we had to resort to sampling of these models. When set to False, we considered all possible models by exhaustive enumeration.

| **Serial Number** | **Node** | $k$ | **EF** | **EUF** | **RoF** | **NCF** |
|---|---|---|---|---|---|---|
| 1 | Bmp2 | 2 | 3 | 1 | 1 | 1 |
| 2 | canWnt | 1 | 1 | 1 | 1 | 1 |
| 3 | Dkk1 | 3 | 52 | 7 | 6 | 6 |
| 4 | Fgf8 | 3 | 54 | 7 | 6 | 6 |
| 5 | Foxc1/2 | 2 | 2 | 2 | 2 | 2 |
| 6 | GATAs | 3 | 52 | 2 | 2 | 2 |
| 7 | IsI1 | 5 | - | 6780 | 420 | 286 |
| 8 | Mesp1 | 2 | 2 | 1 | 1 | 1 |
| 9 | Nkx2_5 | 7 | - | - | 15976 | 6363 |
| 10 | Tbx1 | 1 | 1 | 1 | 1 | 1 |
| 11 | Tbx5 | 6 | - | 7190326 | 3648 | 1735 |
| 12 | exogen_BMP2_I | 1 | 1 | 1 | 1 | 1 |
| 13 | exogen_BMP2_II | 1 | 1 | 1 | 1 | 1 |
| 14 | exogen_canWnt_I | 1 | 1 | 1 | 1 | 1 |
| 15 | exogen_canWnt_II | 1 | 1 | 1 | 1 | 1 |
| Total | | | - | - | 3524801495040 | 454663329120 |
| Sampled | | | True | True | True | True |

TABLE S35. **Nodewise enumeration of the number of BFs that satisfy various biologically meaningful constraints for the MD-GRN model.** The 'Node' column contains all nodes of the network. The '$k$' column is the number of effective inputs to each node. Each of the columns EF, EUF, RoF and NCF provides the numbers of the respective type of BF that satisfy the fixed point constraints for each of the nodes. Additionally, the BFs in EUF, RoF and NCF also satisfy the signs of the inputs in the network architecture. The row labeled 'Total' gives the total number of Boolean models allowed when imposing a given type of BF at each node. Finally, the row labeled 'Sampled' specifies whether we had to resort to sampling of these models. When set to False, we considered all possible models by exhaustive enumeration.

| **Serial Number** | **Node** | $k$ | **EF** | **EUF** | **RoF** | **NCF** |
|---|---|---|---|---|---|---|
| 1 | GATA_2 | 4 | 16148 | 55 | 24 | 21 |
| 2 | GATA_1 | 4 | 8046 | 57 | 26 | 23 |
| 3 | FOG_1 | 1 | 1 | 1 | 1 | 1 |
| 4 | EKLF | 2 | 2 | 1 | 1 | 1 |
| 5 | Fli_1 | 2 | 2 | 1 | 1 | 1 |
| 6 | SCL | 2 | 2 | 2 | 2 | 2 |
| 7 | C/EBP_alpha | 4 | 16148 | 114 | 52 | 46 |
| 8 | PU_1 | 4 | 16149 | 105 | 44 | 38 |
| 9 | cJun | 2 | 2 | 1 | 1 | 1 |
| 10 | EgrNab | 3 | 27 | 6 | 5 | 5 |
| 11 | Gfi_1 | 2 | 2 | 1 | 1 | 1 |
| Total | | | 29273650720346317824 | 450311400 | 14277120 | 8442840 |
| Sampled | | | True | True | True | True |

TABLE S36. **Nodewise enumeration of the number of BFs that satisfy various biologically meaningful constraints for the THC-GRN model.** The 'Node' column contains all nodes of the network. The '$k$' column is the number of effective inputs to each node. Each of the columns EF, EUF, RoF and NCF provides the numbers of the respective type of BF that satisfy the fixed point constraints for each of the nodes. Additionally, the BFs in EUF, RoF and NCF also satisfy the signs of the inputs in the network architecture. The row labeled 'Total' gives the total number of Boolean models allowed when imposing a given type of BF at each node. Here and in the previous rows, '-' denotes that it was computationally infeasible to obtain the exact values. Finally, the row labeled 'Sampled' specifies whether we had to resort to sampling of these models. When set to False, we considered all possible models by exhaustive enumeration.

| **Serial Number** | **Node** | $k$ | **EF** | **EUF** | **RoF** | **NCF** |
|---|---|---|---|---|---|---|
| 1 | NFAT | 1 | 1 | 1 | 1 | 1 |
| 2 | IRAK | 1 | 1 | 1 | 1 | 1 |
| 3 | STAT1 | 2 | 5 | 2 | 2 | 2 |
| 4 | IL10R | 1 | 1 | 1 | 1 | 1 |
| 5 | IL12R | 1 | 1 | 1 | 1 | 1 |
| 6 | IL18R | 2 | 2 | 1 | 1 | 1 |
| 7 | IFNgR | 1 | 1 | 1 | 1 | 1 |
| 8 | IFNg | 5 | - | 1661 | 110 | 73 |
| 9 | STAT4 | 2 | 2 | 1 | 1 | 1 |
| 10 | IL10 | 1 | 1 | 1 | 1 | 1 |
| 11 | STAT3 | 1 | 1 | 1 | 1 | 1 |
| 12 | IFNbR | 1 | 1 | 1 | 1 | 1 |
| 13 | IL4 | 2 | 3 | 1 | 1 | 1 |
| 14 | IL4R | 2 | 1 | 1 | 1 | 1 |
| 15 | GATA3 | 3 | 26 | 7 | 6 | 6 |
| 16 | STAT6 | 1 | 1 | 1 | 1 | 1 |
| 17 | SOCS1 | 2 | 3 | 1 | 1 | 1 |
| 18 | JAK1 | 2 | 3 | 1 | 1 | 1 |
| 19 | Tbet | 3 | 27 | 5 | 4 | 4 |
| 20 | IFNb | 1 | 1 | 1 | 1 | 1 |
| 21 | TCR | 1 | 1 | 1 | 1 | 1 |
| 22 | IL18 | 1 | 1 | 1 | 1 | 1 |
| 23 | IL12 | 1 | 1 | 1 | 1 | 1 |
| Total | | | - | 116270 | 5280 | 3504 |
| Sampled | | | True | False | False | False |

TABLE S37. **Nodewise enumeration of the number of BFs that satisfy various biologically meaningful constraints for the EGFR-GRN model.** The 'Node' column contains all nodes of the network. The '$k$' column is the number of effective inputs to each node. Each of the columns EF, EUF, RoF and NCF provides the numbers of the respective type of BF that satisfy the fixed point constraints for each of the nodes. Additionally, the BFs in EUF, RoF and NCF also satisfy the signs of the inputs in the network architecture. The row labeled 'Total' gives the total number of Boolean models allowed when imposing a given type of BF at each node. Finally, the row labeled 'Sampled' specifies whether we had to resort to sampling of these models. When set to False, we considered all possible models by exhaustive enumeration.

| **Serial Number** | **Node** | $k$ | **EF** | **EUF** | **RoF** | **NCF** |
|---|---|---|---|---|---|---|
| 1 | EGFR | 1 | 1 | 1 | 1 | 1 |
| 2 | ERK1/2 | 1 | 1 | 1 | 1 | 1 |
| 3 | PI3K | 2 | 2 | 2 | 2 | 2 |
| 4 | AKT | 1 | 1 | 1 | 1 | 1 |
| 5 | GSK3_beta | 1 | 1 | 1 | 1 | 1 |
| 6 | MDM2 | 2 | 3 | 1 | 1 | 1 |
| 7 | TP53 | 3 | 54 | 7 | 6 | 6 |
| 8 | PTEN | 1 | 1 | 1 | 1 | 1 |
| 9 | PARP1 | 1 | 1 | 1 | 1 | 1 |
| 10 | BRCA1 | 1 | 1 | 1 | 1 | 1 |
| 11 | BCL_2 | 1 | 1 | 1 | 1 | 1 |
| 12 | BAX | 2 | 2 | 2 | 2 | 2 |
| 13 | CYCD1 | 4 | 16148 | 114 | 52 | 46 |
| Total | | | 10463904 | 3192 | 1248 | 1104 |
| Sampled | | | True | False | False | False |

TABLE S38. **Nodewise enumeration of the number of BFs that satisfy various biologically meaningful constraints for the EMT-Senescence-GRN model.** The 'Node' column contains all nodes of the network. The '$k$' column is the number of effective inputs to each node. Each of the columns EF, EUF, RoF and NCF provides the numbers of the respective type of BF that satisfy the fixed point constraints for each of the nodes. Additionally, the BFs in EUF, RoF and NCF also satisfy the signs of the inputs in the network architecture. The row labeled 'Total' gives the total number of Boolean models allowed when imposing a given type of BF at each node. Here and in the previous rows, '-' denotes that it was computationally infeasible to obtain the exact values. Finally, the row labeled 'Sampled' specifies whether we had to resort to sampling of these models. When set to False, we considered all possible models by exhaustive enumeration.

| **Serial Number** | **Node** | $k$ | **EF** | **EUF** | **RoF** | **NCF** |
|---|---|---|---|---|---|---|
| 1 | Snai2 | 3 | 54 | 7 | 6 | 6 |
| 2 | ESE2 | 3 | 54 | 7 | 6 | 6 |
| 3 | p16 | 5 | - | 1718 | 136 | 96 |
| 4 | E2F | 3 | 28 | 2 | 2 | 2 |
| 5 | Cyclin | 4 | 8103 | 43 | 14 | 11 |
| 6 | TELasa | 1 | 1 | 1 | 1 | 1 |
| 7 | NFkB | 4 | 8048 | 36 | 18 | 17 |
| 8 | Rb | 3 | 55 | 5 | 4 | 4 |
| 9 | p53 | 4 | 8103 | 48 | 18 | 15 |
| Total | | | - | 62550593280 | 177666048 | 77552640 |
| Sampled | | | True | True | True | True |

TABLE S39. **Nodewise enumeration of the number of BFs that satisfy various biologically meaningful constraints for the FOS-GRN model.** The 'Node' column contains all nodes of the network. The '$k$' column is the number of effective inputs to each node. Each of the columns EF, EUF, RoF and NCF provides the numbers of the respective type of BF that satisfy the fixed point constraints for each of the nodes. Additionally, the BFs in EUF, RoF and NCF also satisfy the signs of the inputs in the network architecture. The row labeled 'Total' gives the total number of Boolean models allowed when imposing a given type of BF at each node. Here and in the previous rows, '-' denotes that it was computationally infeasible to obtain the exact values. Finally, the row labeled 'Sampled' specifies whether we had to resort to sampling of these models. When set to False, we considered all possible models by exhaustive enumeration.

| **Serial Number** | **Node** | $k$ | **EF** | **EUF** | **RoF** | **NCF** |
|---|---|---|---|---|---|---|
| 1 | AG | 8 | - | - | 324888 | 66042 |
| 2 | AP1 | 6 | - | 274247 | 380 | 178 |
| 3 | AP2 | 1 | 1 | 1 | 1 | 1 |
| 4 | AP3 | 7 | - | - | 9018 | 2920 |
| 5 | EMF1 | 1 | 1 | 1 | 1 | 1 |
| 6 | FT | 1 | 1 | 1 | 1 | 1 |
| 7 | FUL | 2 | 2 | 1 | 1 | 1 |
| 8 | LFY | 2 | 2 | 2 | 2 | 2 |
| 9 | PI | 6 | - | 3500117 | 1436 | 627 |
| 10 | SEP | 1 | 1 | 1 | 1 | 1 |
| 11 | TFL1 | 3 | 26 | 7 | 6 | 6 |
| 12 | UFO | 1 | 1 | 1 | 1 | 1 |
| 13 | WUS | 3 | 13 | 2 | 2 | 2 |
| Total | | | - | - | 38370121979258880 | 516537496316160 |
| Sampled | | | True | True | True | True |

TABLE S40. **Nodewise enumeration of the number of BFs that satisfy various biologically meaningful constraints for the GSD-GRN model.** The 'Node' column contains all nodes of the network. The '$k$' column is the number of effective inputs to each node. Each of the columns EF, EUF, RoF and NCF provides the numbers of the respective type of BF that satisfy the fixed point constraints for each of the nodes. Additionally, the BFs in EUF, RoF and NCF also satisfy the signs of the inputs in the network architecture. The row labeled 'Total' gives the total number of Boolean models allowed when imposing a given type of BF at each node. Here and in the previous rows, '-' denotes that it was computationally infeasible to obtain the exact values. Finally, the row labeled 'Sampled' specifies whether we had to resort to sampling of these models. When set to False, we considered all possible models by exhaustive enumeration.

| **Serial Number** | **Node** | $k$ | **EF** | **EUF** | **RoF** | **NCF** |
|---|---|---|---|---|---|---|
| 1 | UGR | 3 | 55 | 6 | 5 | 5 |
| 2 | CBX2 | 4 | 16148 | 9 | 8 | 8 |
| 3 | GATA4 | 4 | 16148 | 7 | 6 | 6 |
| 4 | WT1mKTS | 2 | 5 | 1 | 1 | 1 |
| 5 | WT1pKTS | 4 | 16146 | 96 | 36 | 30 |
| 6 | NR5A1 | 6 | - | 6609378 | 2736 | 1260 |
| 7 | NR0B1 | 5 | - | 6780 | 420 | 286 |
| 8 | SRY | 8 | - | - | 1032456 | 239279 |
| 9 | SOX9 | 7 | - | - | 78416 | 29024 |
| 10 | FGF9 | 2 | 2 | 2 | 2 | 2 |
| 11 | PGD2 | 1 | 1 | 1 | 1 | 1 |
| 12 | DMRT1 | 3 | 55 | 9 | 8 | 8 |
| 13 | DHH | 1 | 1 | 1 | 1 | 1 |
| 14 | DKK1 | 2 | 2 | 2 | 2 | 2 |
| 15 | AMH | 5 | - | 6780 | 420 | 286 |
| 16 | WNT4 | 6 | - | 7778168 | 5032 | 2542 |
| 17 | RSPO1 | 3 | 55 | 9 | 8 | 8 |
| 18 | FOXL2 | 4 | 16148 | 114 | 52 | 46 |
| 19 | CTNNB1 | 5 | - | 6894 | 472 | 332 |
| Total | | | - | - | $1.067 \times 10^{34}$ | $5.121 \times 10^{31}$ |
| Sampled | | | True | True | True | True |

### SUPPLEMENTARY FIGURES

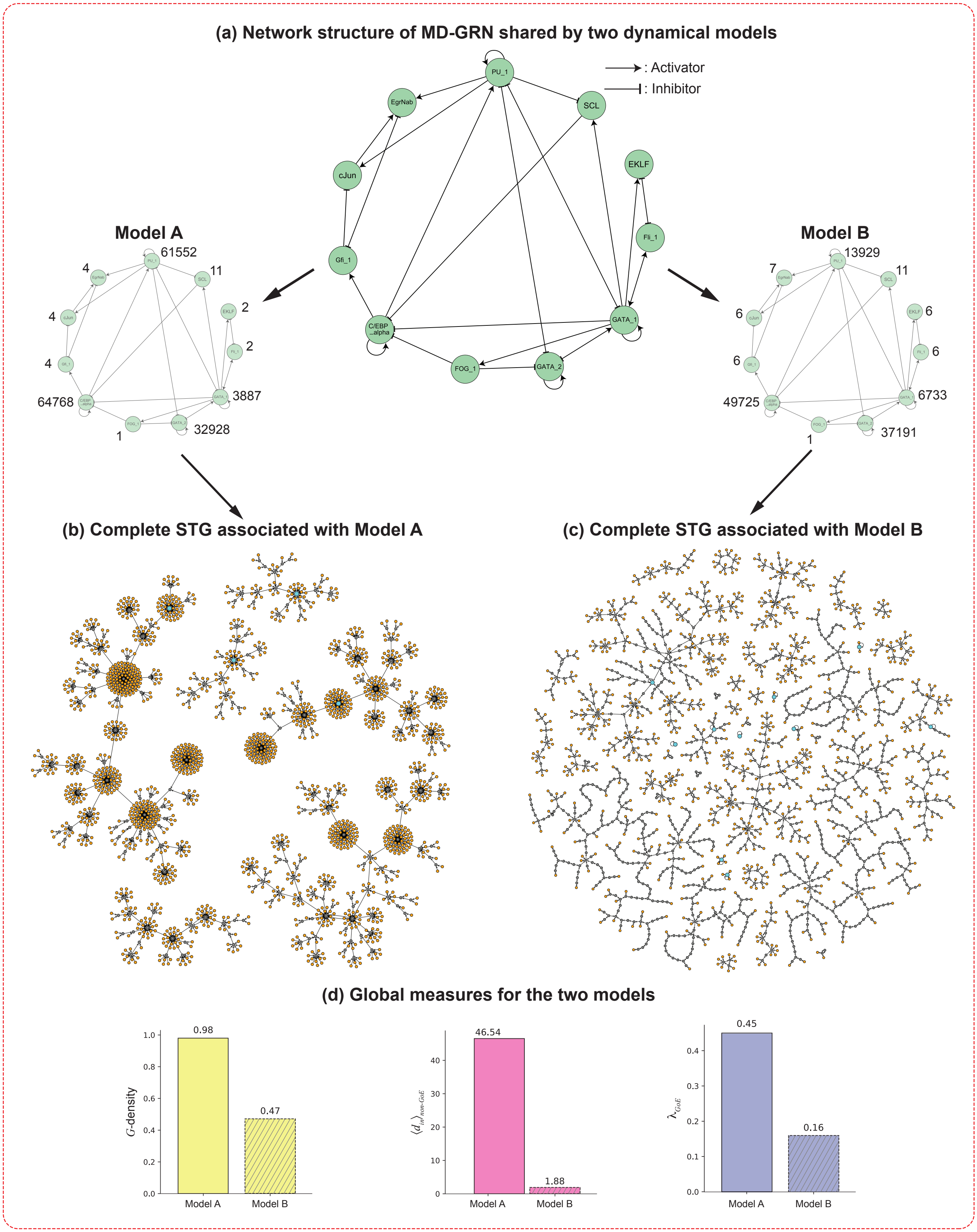

FIG. S1. **STGs for two different choices of Boolean rules for the biological network MD-GRN and the values of their bushiness and convergence measures**. **(a)** The network in the centre of the top panel is the Myeloid differentiation GRN (MD-GRN). On its two sides are Boolean models (Model 1 and Model 2) with identical network structure but different regulatory logic rules. For these two models, the genes are arranged in the same manner as depicted in the network structure in the centre. The BFs at each node are encoded as integers as described in SI text, section 1). In the truth table, the (left to right) order of the inputs for each of the genes in the network is as follows: GATA_2: [GATA_2, GATA_1, FOG_1, PU_1], GATA_1: [GATA_2, GATA_1, Fli_1, PU_1], FOG_1: [GATA_1], EKLF: [GATA_1, Fli_1], Fli1: [GATA_1, EKLF], SCL: [GATA_1, PU_1], C/EBP_alpha: [GATA_1, FOG_1, SCL, C/EBP_alpha], PU_1: [GATA_2, GATA_1, C/EBP_alpha, PU_1], cJun: [Gfi_1, PU_1], EgrNab: [cJun, Gfi_1, PU_1], Gfi_1:[EgrNab, C/EBP_alpha]. **(b)** and **(c)** display the complete STGs containing 2048 states obtained for Model 1 and Model 2 respectively. The 'orange', 'blue' and 'grey' nodes indicate the *GoE* states, fixed points and *non-GoE* states that are not fixed points respectively. Visual inspection shows that the STG for Model 1 appears significantly more bushy than the STG for Model 2. **(d)** Barplots representing the bushiness ($G$-density, $\langle d_{in} \rangle_{non-GoE}$) and convergence ($\lambda_{GoE}$) measures for Model 1 and Model 2. Each measure is shown as a separate bar plot that quantifies the visible difference between (b) and (c).

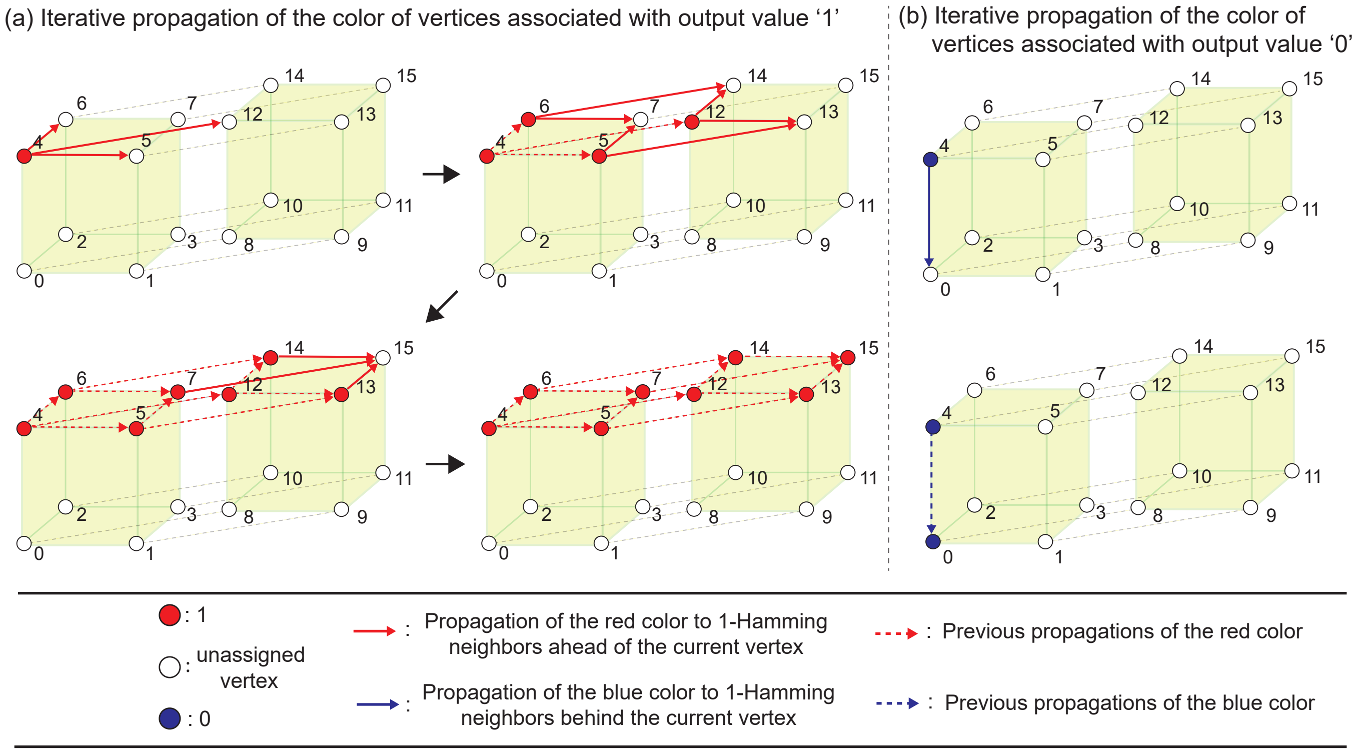


FIG. S2. **Visual illustration of how monotonicity forces propagation of the color on vertices of the Boolean hypercube.** We assume that the BF is positive monotone in this illustration. We begin with a configuration on the hypercube where the vertex 4 is colored either red or blue (corresponding to output values 1 or 0 respectively) and other vertices uncolored. **(a)** Left panel: in the top left subfigure, vertex 4 is colored red and its color is propagated in the 'forward' direction via the red solid arrows to its 1-Hamming neighbours which are 'ahead' of it (vertices 5, 6 and 12). Note that if any of the vertices 5, 6 or 12 was colored blue, the BF would violate positive monotonicity. In the subfigure to the right (following the arrow), each of the vertices 5, 6 and 12 propagate their color to their 1-Hamming neighbours 'ahead' of them, (7, 13), (7, 14), and (13, 14) respectively. In the bottom left subfigure, each of the vertices, 7, 13 and 14 propagate their color to their 1-Hamming neighbours, vertex 15. In the bottom right subfigure, vertex 15 is colored red and no further propagation is possible, thereby terminating the iterative propagation. **(b)** Right panel: in the top subfigure, vertex 4 is colored blue and its color is propagated in the backward direction via the blue solid arrows to its 1-Hamming neighbours which are 'behind' it (vertex 0). Note that if the vertex 0 was colored red, the BF would violate positive monotonicity. In the bottom subfigure, vertex 0 is colored blue and no further propagation is possible, thereby terminating the iterative propagation.

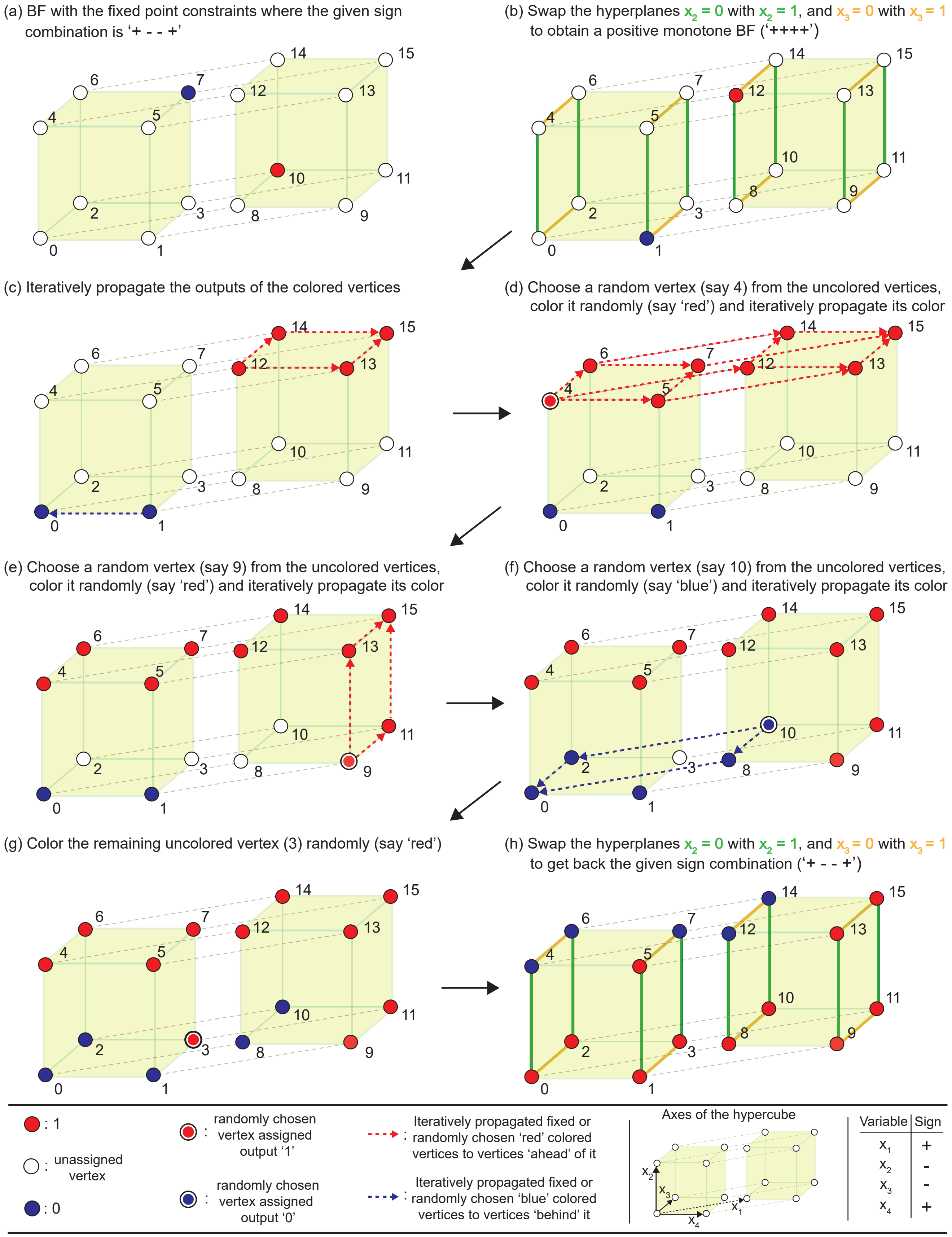
(a) BF with the fixed point constraints where the given sign combination is '+ - - +'
(b) Swap the hyperplanes x2 = 0 with x2 = 1, and x3 = 0 with x3 = 1 to obtain a positive monotone BF ('++++')
(c) Iteratively propagate the outputs of the colored vertices
(d) Choose a random vertex (say 4) from the uncolored vertices, color it randomly (say 'red') and iteratively propagate its color
(e) Choose a random vertex (say 9) from the uncolored vertices, color it randomly (say 'red') and iteratively propagate its color
(f) Choose a random vertex (say 10) from the uncolored vertices, color it randomly (say 'blue') and iteratively propagate its color
(g) Color the remaining uncolored vertex (3) randomly (say 'red')
(h) Swap the hyperplanes x2 = 0 with x2 = 1, and x3 = 0 with x3 = 1 to get back the given sign combination ('+ - - +')
: 1
: unassigned vertex
: 0
: randomly chosen vertex assigned output '1'
: randomly chosen vertex assigned output '0'
: Iteratively propagated fixed or randomly chosen 'red' colored vertices to vertices 'ahead' of it
: Iteratively propagated fixed or randomly chosen 'blue' colored vertices to vertices 'behind' it
Axes of the hypercube
x1
x2
x3
x4
Variable | Sign
x1 | +
x2 | -
x3 | -
x4 | +

FIG. S3. **Generating a random UF with a given sign combination that satisfies given fixed point constraints.** We show via a 4-input BF how we generate random UFs that obey a given sign combination of the inputs and given fixed point constraints. We assume that for the given sign combination, the inputs $x_1$ and $x_4$ are activatory and inputs $x_2$ and $x_3$ are inhibitory. **(a) Initial configuration of the BF after imposing fixed point constraints.** Assume that the fixed points constrain the vertices 7 and 10 to have the output values 0 (blue) and 1 (red) respectively. **(b) Conversion of the BF to a positive monotone BF by negation of inhibitory inputs.** Inputs 2 and 3 are negated by swapping the hyperplanes $x_2 = 0$ and $x_3 = 0$ with the hyperplanes $x_2 = 1$ and $x_3 = 1$ respectively (the order does not matter). This is equivalent to exchanging the vertices along the green (or orange) colored edges that are perpendicular (or parallel) to the plane of the paper. This results in a BF where the fixed point constraints now satisfy positive monotonicity on all inputs. **(c) Iterative propagation of the vertex colors.** The color of the vertices 1 and 12 are iteratively propagated 'backward' and 'forward' respectively to vertices 'behind' and 'ahead' respectively as shown by the dashed blue and red arrows. **(d)-(g) Selection of a random uncolored vertex, coloring it randomly and iteratively propagating its color.** Since some vertices remain uncolored after step (c), a vertex is randomly chosen from the uncolored vertices (say 4) and colored randomly (say red) as shown in step (d). The color of this vertex is then iteratively propagated. At the end of step (d) since there are uncolored vertices, the above-mentioned procedure is repeated in steps (e), (f) and (g) till there are no vertices left to be colored on the hypercube. Subfigure (g) shows the sampled positive monotone BF that satisfies the fixed point constraints. **(h) Conversion of the sampled positive monotone BF to the given sign combination.** The inputs 2 and 3 are negated again as was done in step (b) to attain the given sign combination ('+ - - +'). Following step (b), the hyperplanes $x_2 = 0$ and $x_3 = 0$ are swapped with the hyperplanes $x_2 = 1$ and $x_3 = 1$ respectively (corresponding to vertex swaps along the green (or orange) colored edges that are perpendicular (or parallel) to the plane of the paper). This BF is a random UF where the inputs obey the given sign combination and the BF satisfies the fixed point constraints.

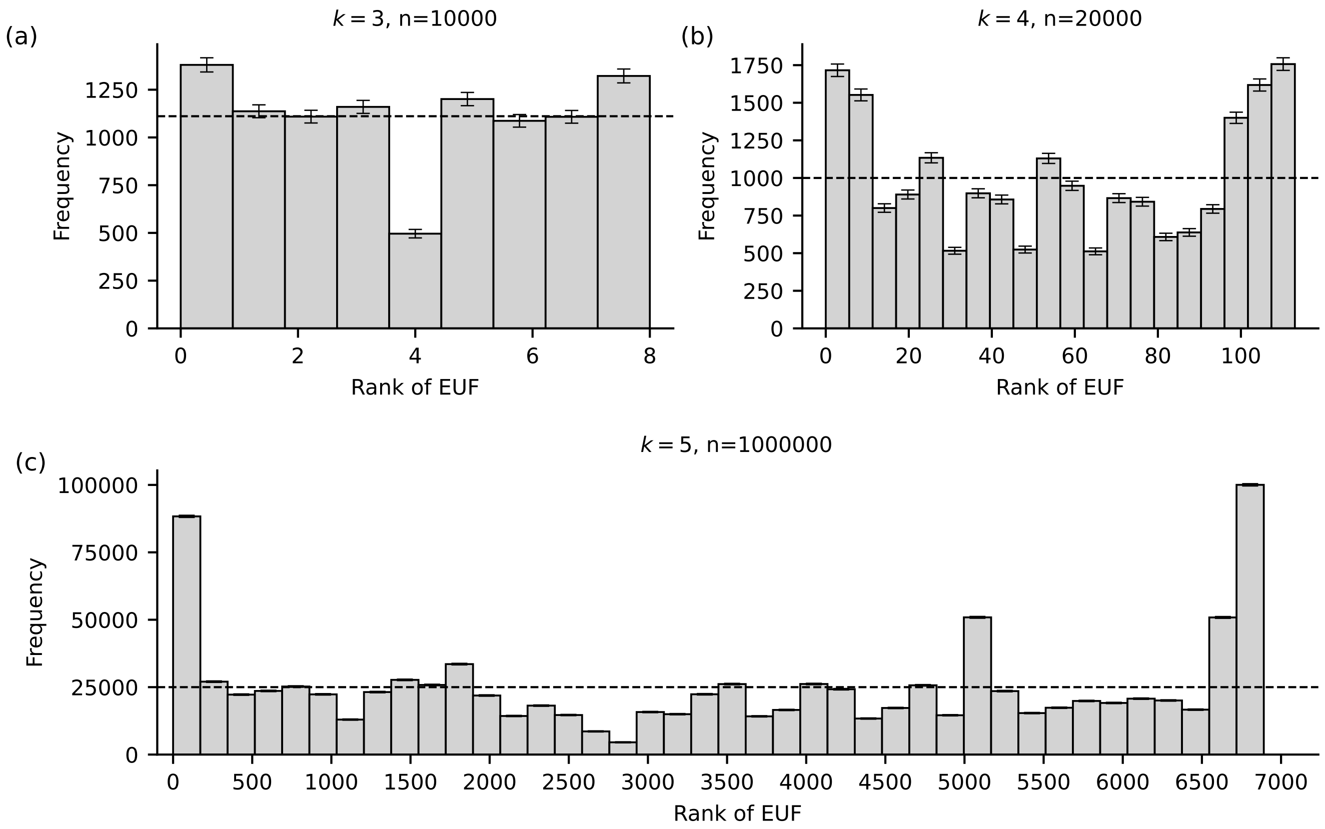


FIG. S4. **The distribution of positive monotone effective and unate functions (EUFs) sampled using the proposed EUF sampling algorithm for BFs with $k = 3$, $k = 4$ and $k = 5$.** In each subplot, the $x$ axis is the rank associated with a positive monotone EUF based on its integer encoding. Each EUF is ranked from 0 (highest rank) to the total number of distinct functions generated by the algorithm (the lowest rank). The ranking is based on the integer representation of the BF (see SI text, section 1), with lower integers corresponding to higher ranks. The $y$ axis in each subplot is the frequency of the positive monotone EUFs generated by our sampling algorithm. The subplots **(a)**, **(b)** and **(c)** shows the distribution of $n = 10000$, $n = 20000$ and $n = 1000000$ samples of $k = 3$, $k = 4$ and $k = 5$ input positive monotone EUF respectively, generated using our sampling algorithm (see Methods). The horizontal dashed line represents the expected frequency value were the sampling to be perfectly uniform. The error bars in all histograms are extremely tiny (sometimes too small to be visible). The distribution generated using the sampling algorithm is skewed towards the highest and lowest ranks, but is relatively close to the expected frequency value for other ranks.

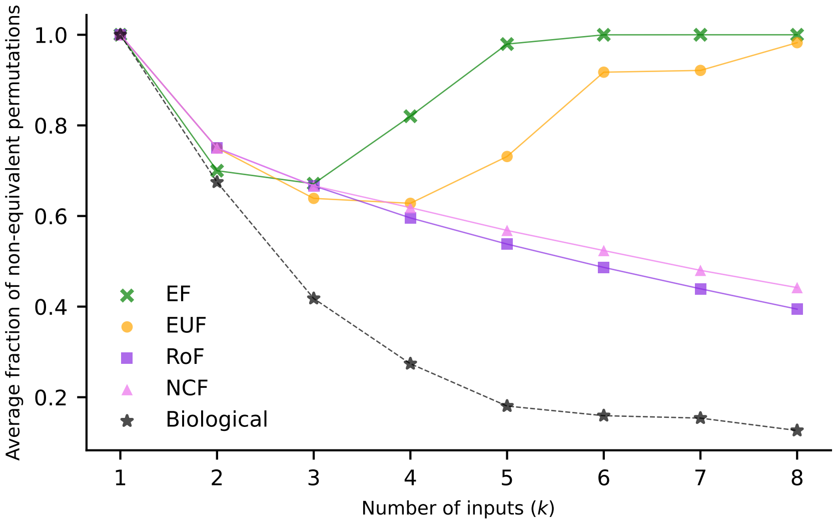


FIG. S5. **The fraction of non-equivalent permutations for different types of BFs.** The $x$-axis is the number of inputs to the BF. The $y$-axis is the average of the fraction of non-equivalent permutations over a set of $k$-input BFs. Each data point on the plot represents the average of the fractions of non-equivalent permutations over the set of all (or sampled) $k$-input BFs belonging to a particular type of BF (as indicated by the shape of the associated point). For inputs $k = 1, k = 2, k = 3, k = 4, k = 5$, the exhaustive set of BFs was used in the computation for all types of BFs (except for 5-input EFs for which 100000 BFs were sampled), whereas for inputs $k = 6, k = 7, k = 8$, 100000 sampled BFs were generated for all types of BFs. The data points corresponding to the 'biological' function type were obtained from a reference biological dataset of 2687 BFs derived from 88 published Boolean models [1]. It is seen that the 'biological' case generally consists of BFs with a low fraction of non-equivalent permutations, justifying the usefulness of our approach for computing $Z_{ave}$.

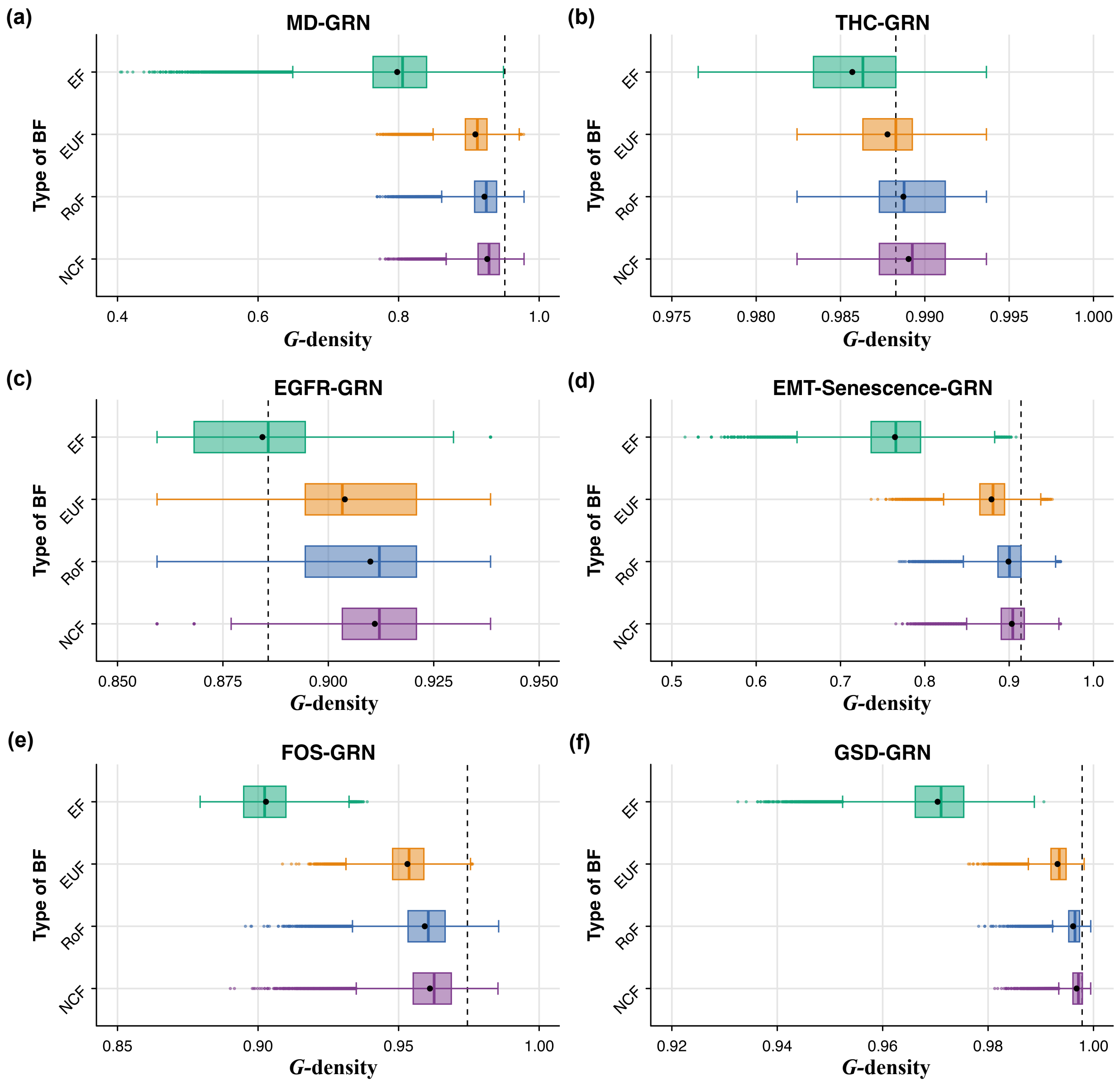


FIG. S6. **Distribution of $G$-density values for different ensembles generated using network structures from 6 published Boolean GRNs**. The subplots **(a)**, **(b)**, **(c)**, **(d)**, **(e)** and **(f)** correspond to the MD-GRN, THC-GRN, EGFR-GRN, EMT-Senescence-GRN, FOS-GRN and GSD-GRN network structures respectively. The $x$ and $y$ axes of each subplot correspond respectively to $G$-density and to the type of BF imposed on the nodes of the networks. The box plots display the distribution of $G$-density in the 4 ensembles that each use a given type of regulatory logic. The ensembles generated using biologically meaningful BFs (EUFs, RoFs, NCFs) have higher $G$-density compared to the ensemble generated using EFs. Note that we impose that ensembles recover the biological fixed points of the associated published model. The mean and median of the distribution are indicated by the black dot and the vertical line within the box respectively. The vertical dashed lines in each subplot correspond to the $G$-density of the Boolean model provided by the modelers in the published article.

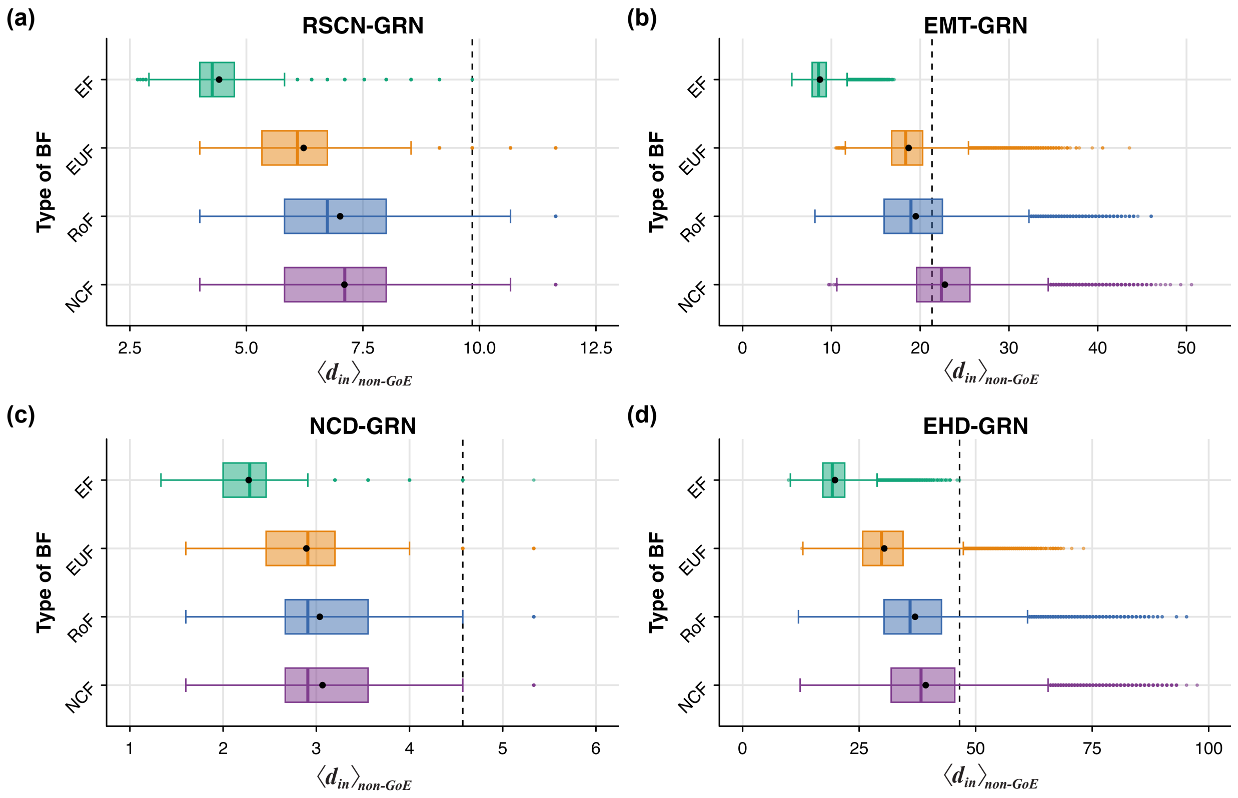


FIG. S7. **Distribution of average in-degree of *non-GoE* states ($\langle d_{in} \rangle_{non-GoE}$) for different ensembles generated using network structures from 4 published Boolean GRNs**. The subplots **(a)**, **(b)**, **(c)** and **(d)** correspond to the RSCN-GRN, EMT-GRN, NCD-GRN and EHD-GRN network structures respectively. The $x$ and $y$ axes of each subplot correspond respectively to $\langle d_{in} \rangle_{non-GoE}$ and to the type of BF imposed on the nodes of the networks. The box plots display the distribution of $\langle d_{in} \rangle_{non-GoE}$ in the 4 ensembles that each use a given type of regulatory logic. The ensembles generated using biologically meaningful BFs (EUFs, RoFs, NCFs) have higher $\langle d_{in} \rangle_{non-GoE}$ compared to the ensemble generated using EFs. Note that we impose that ensembles recover the biological fixed points of the associated published model. The mean and median of the distribution are indicated by the black dot and the vertical line within the box respectively. The vertical dashed lines in each subplot correspond to the $\langle d_{in} \rangle_{non-GoE}$ of the Boolean model provided by the modelers in the published article.

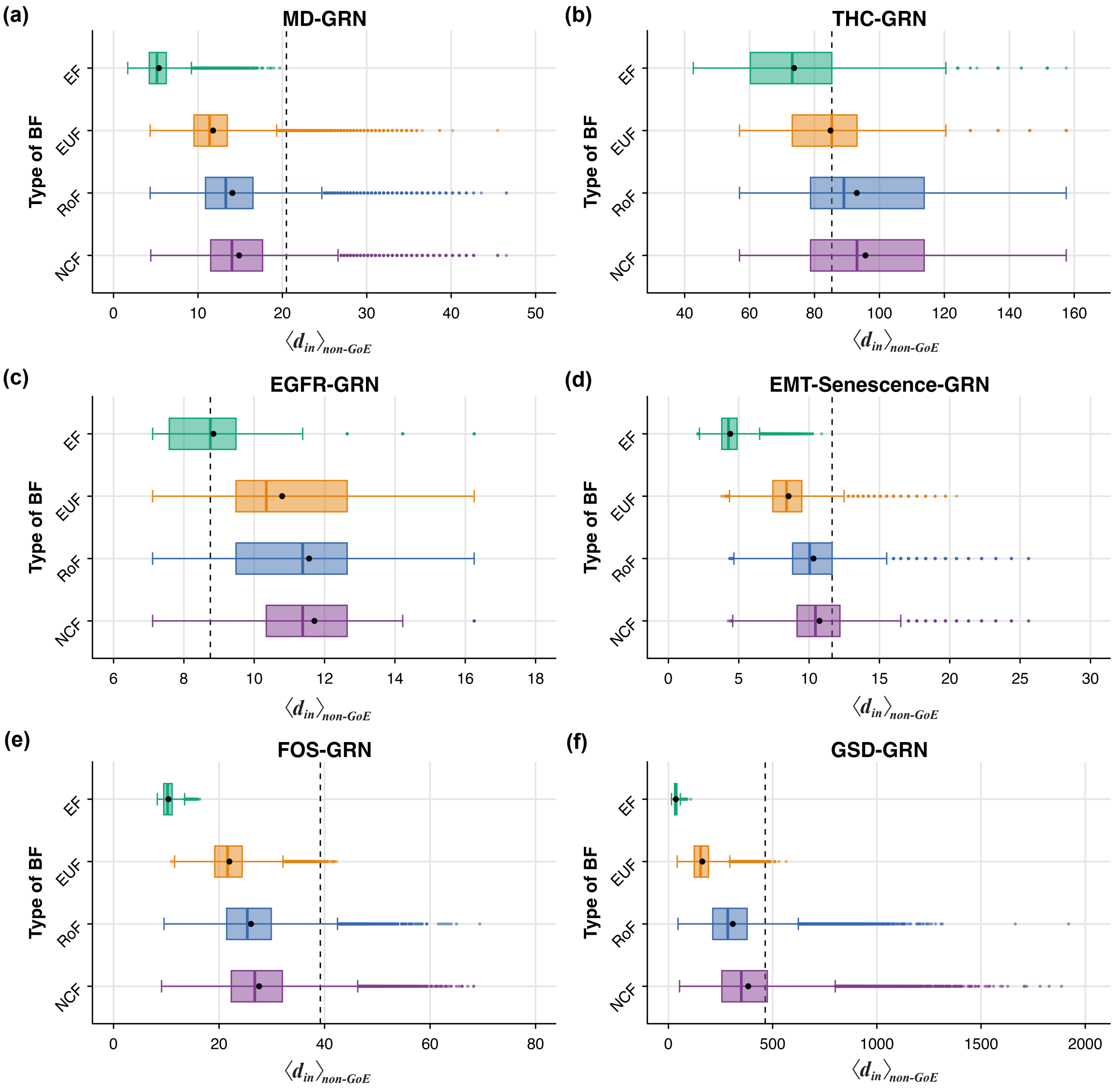


FIG. S8. **Distribution of average in-degree of *non-GoE* states ($\langle d_{in}\rangle_{non-GoE}$) for different ensembles generated using network structures from 6 published Boolean GRNs**. The subplots **(a)**, **(b)**, **(c)**, **(d)**, **(e)** and **(f)** correspond to the MD-GRN, THC-GRN, EGFR-GRN, EMT-Senescence-GRN, FOS-GRN and GSD-GRN network structures respectively. The $x$ and $y$ axes of each subplot correspond respectively to $\langle d_{in}\rangle_{non-GoE}$ and to the type of BF imposed on the nodes of the networks. The box plots display the distribution of $\langle d_{in}\rangle_{non-GoE}$ in the 4 ensembles that each use a given type of regulatory logic. The ensembles generated using biologically meaningful BFs (EUFs, RoFs, NCFs) have higher $\langle d_{in}\rangle_{non-GoE}$ compared to the ensemble generated using EFs. Note that we impose that ensembles recover the biological fixed points of the associated published model. The mean and median of the distribution are indicated by the black dot and the vertical line within the box respectively. The vertical dashed lines in each subplot correspond to the $\langle d_{in}\rangle_{non-GoE}$ of the Boolean model considered by the modelers in the published article.

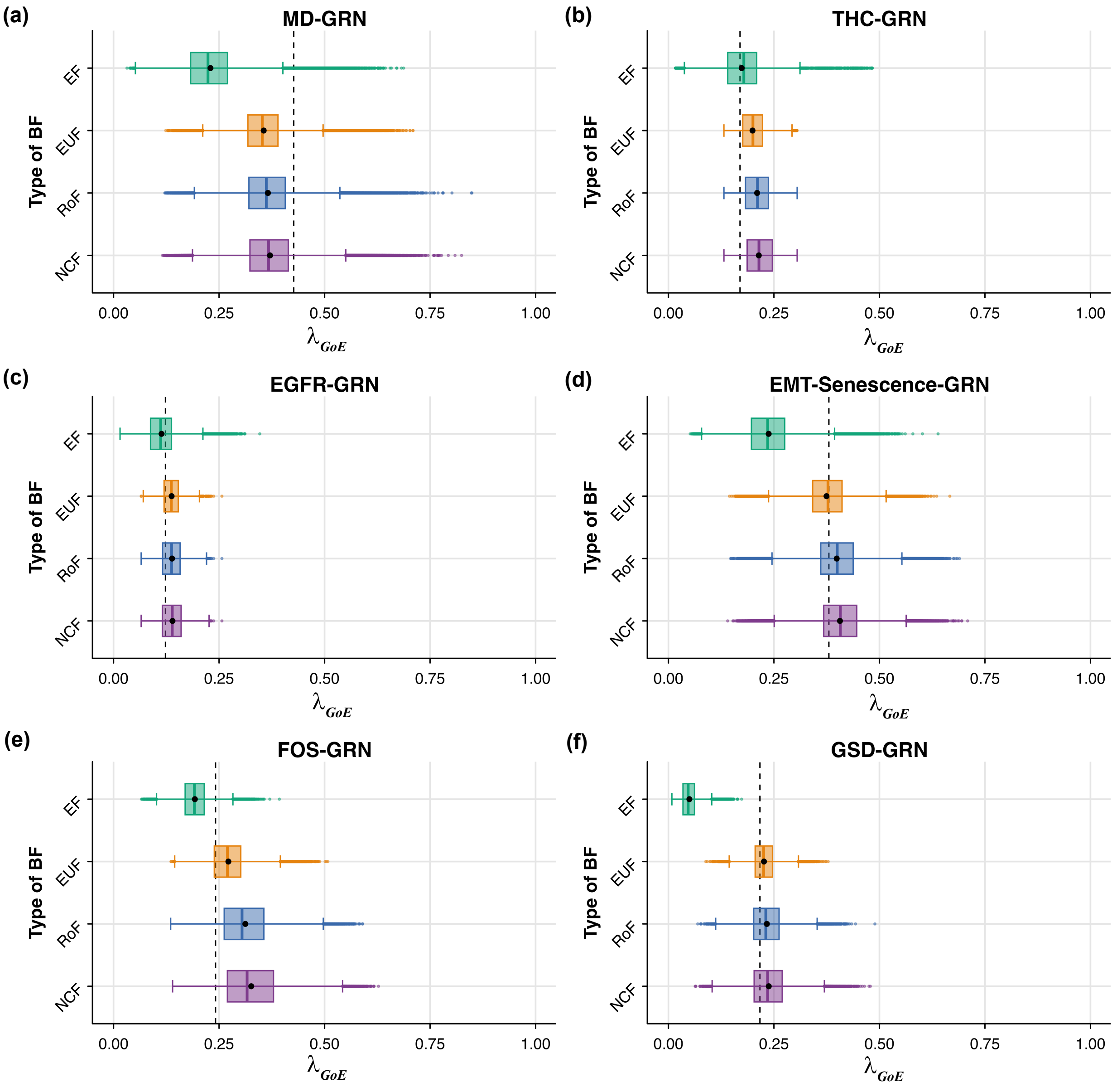


FIG. S9. **Distribution of average convergence rate of trajectories originating at *GoE* states ($\lambda_{GoE}$) for different ensembles generated using network structures from 6 published Boolean GRNs**. The subplots **(a)**, **(b)**, **(c)**, **(d)**, **(e)** and **(f)** correspond to the MD-GRN, THC-GRN, EGFR-GRN, EMT-Senescence-GRN, FOS-GRN and GSD-GRN network structures respectively. The $x$ and $y$ axes of each subplot correspond respectively to $\lambda_{GoE}$ and to the type of BF imposed on the nodes of the networks. The box plots display the distribution of $\lambda_{GoE}$ in the 6 ensembles that each use a given type of regulatory logic. The ensembles generated using biologically meaningful BFs (EUFs, RoFs, NCFs) have higher convergence rates compared to the ensemble generated using EFs. Note that we impose that ensembles recover the biological fixed points of the associated published model. The mean and median of the distribution are indicated by the black dot and the vertical line within the box respectively. The vertical dashed lines in each subplot correspond to the $\lambda_{GoE}$ of the Boolean model provided by the modelers in the published article.

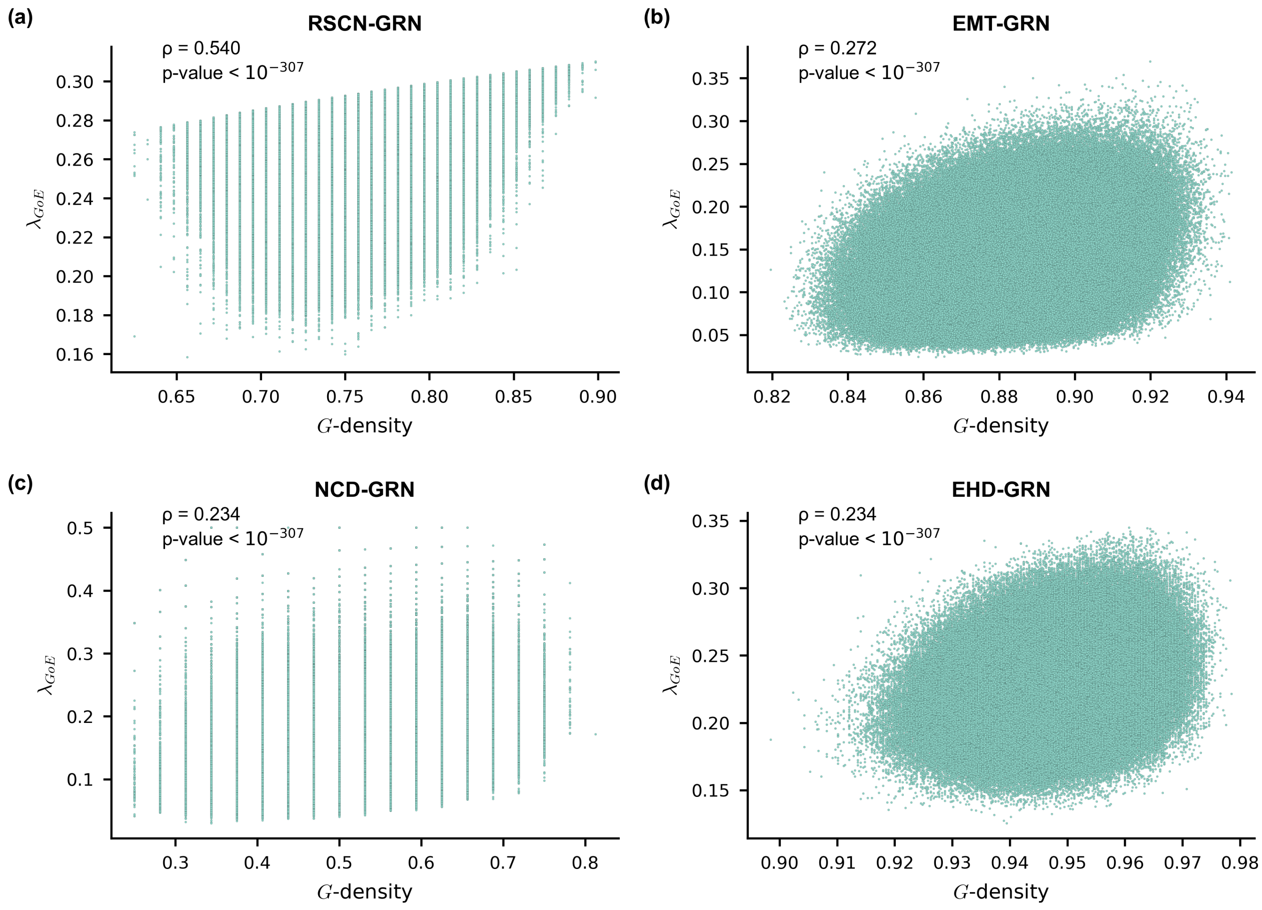


FIG. S10. **Scatter plot between the $G$-density and average convergence rate of trajectories originating at $GoE$ states ($\lambda_{GoE}$) for the EF ensemble generated using network structures from 4 published Boolean GRNs.** The subplots **(a)**, **(b)**, **(c)** and **(d)** correspond to the RSCN-GRN, EMT-GRN, NCD-GRN and EHD-GRN network structures respectively. The $x$ and $y$ axis of each subplot correspond to the $G$-density and $\lambda_{GoE}$ respectively. In each subplot we provide the Spearman correlation coefficient ($\rho$) along with the associated p-value. $G$-density and $\lambda_{GoE}$ show a moderate positive correlation for the RSCN-GRN network and a weak positive correlation otherwise.

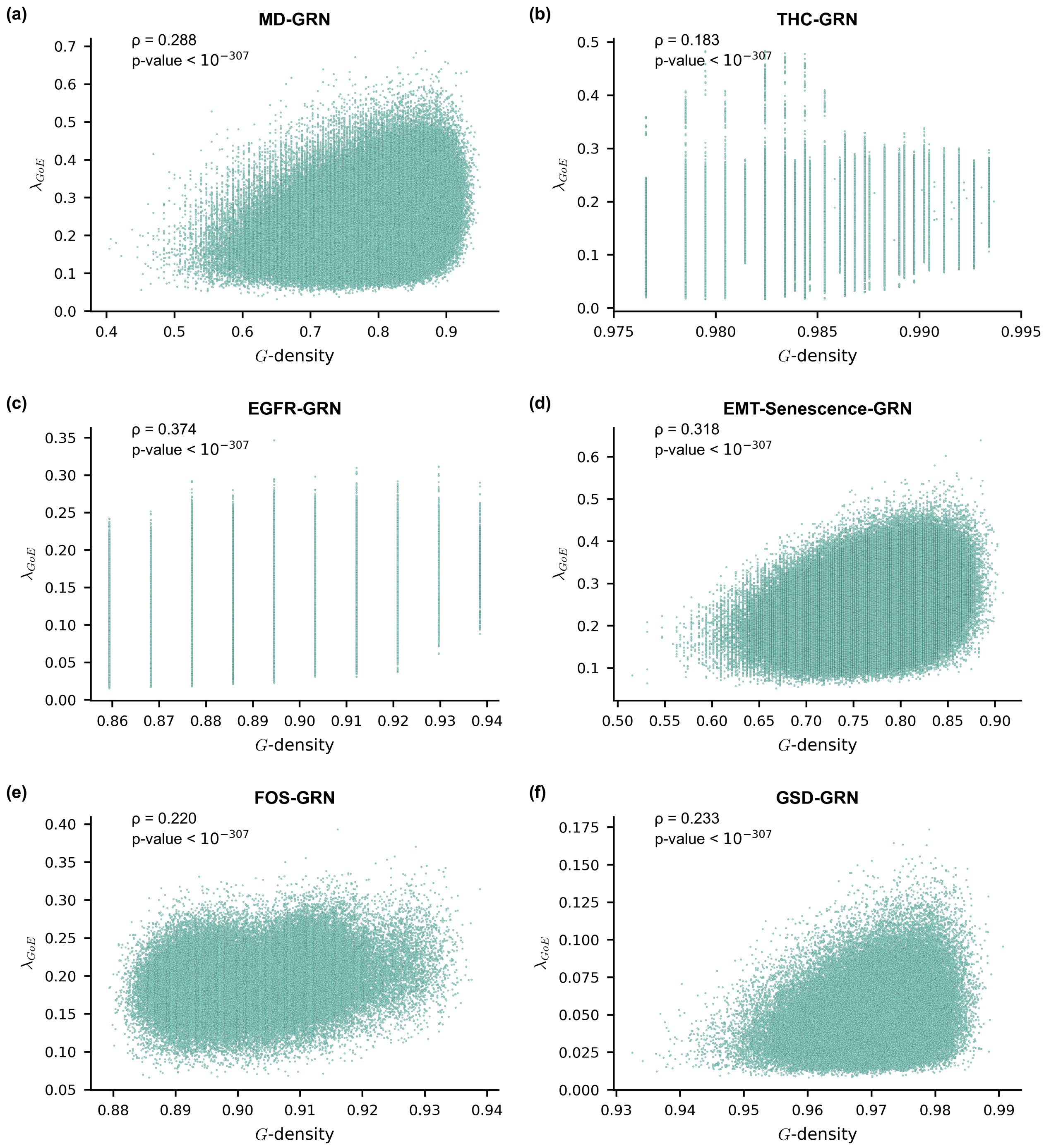


FIG. S11. **Scatter plot between the $G$-density and average convergence rate of trajectories originating at $GoE$ states ($\lambda_{GoE}$) for the EF ensemble generated using network structures from 6 published Boolean GRNs.** The subplots **(a)**, **(b)**, **(c)**, **(d)**, **(e)** and **(f)** correspond to the MD-GRN, THC-GRN, EGFR-GRN, EMT-Senescence-GRN, FOS-GRN and GSD-GRN network structures respectively. The $x$ and $y$ axis of each subplot correspond to the $G$-density and $\lambda_{GoE}$ respectively. In each subplot we provide the Spearman correlation coefficient ($\rho$) along with the associated p-value. $G$-density and $\lambda_{GoE}$ show a weak positive correlation for all of the networks except for the THC-GRN network which shows a very weak positive correlation.

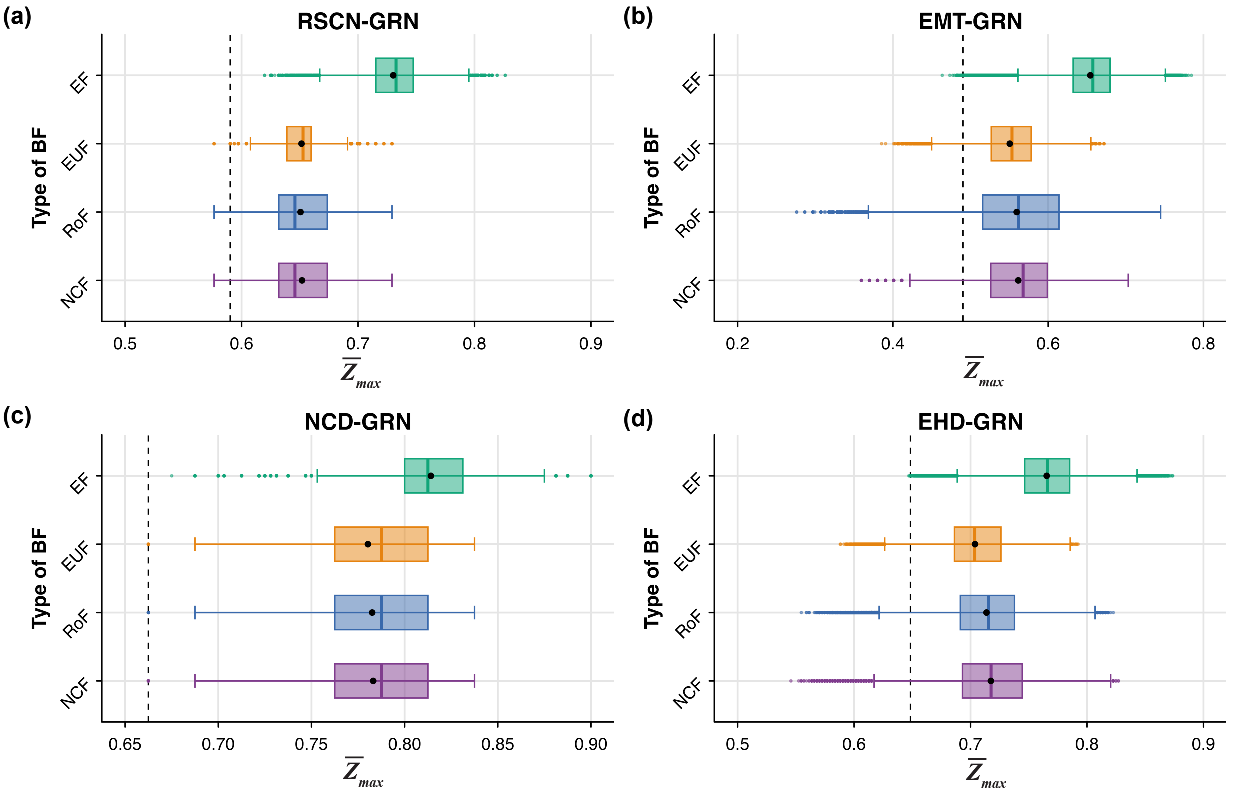


FIG. S12. **Distribution of $\overline{Z}_{max}$ values for different ensembles generated using network structures from 4 published Boolean GRNs**. The subplots **(a)**, **(b)**, **(c)** and **(d)** correspond to the RSCN-GRN, EMT-GRN, NCD-GRN and EHD-GRN network structures respectively. The $x$ and $y$ axes of each subplot correspond respectively to $\overline{Z}_{max}$ and to the type of BF imposed on the nodes of the networks. The box plots display the distribution of $\overline{Z}_{max}$ in the 4 ensembles that each use a given type of regulatory logic. The ensembles generated using biologically meaningful BFs (EUFs, RoFs, NCFs) have lower $\overline{Z}_{max}$ compared to the ensemble generated using EFs. Note that we impose that ensembles recover the biological fixed points of the associated published model. The mean and median of the distribution are indicated by the black dot and the vertical line within the box respectively. The vertical dashed lines in each subplot correspond to the $\overline{Z}_{max}$ of the Boolean model provided by the modelers in the published article.

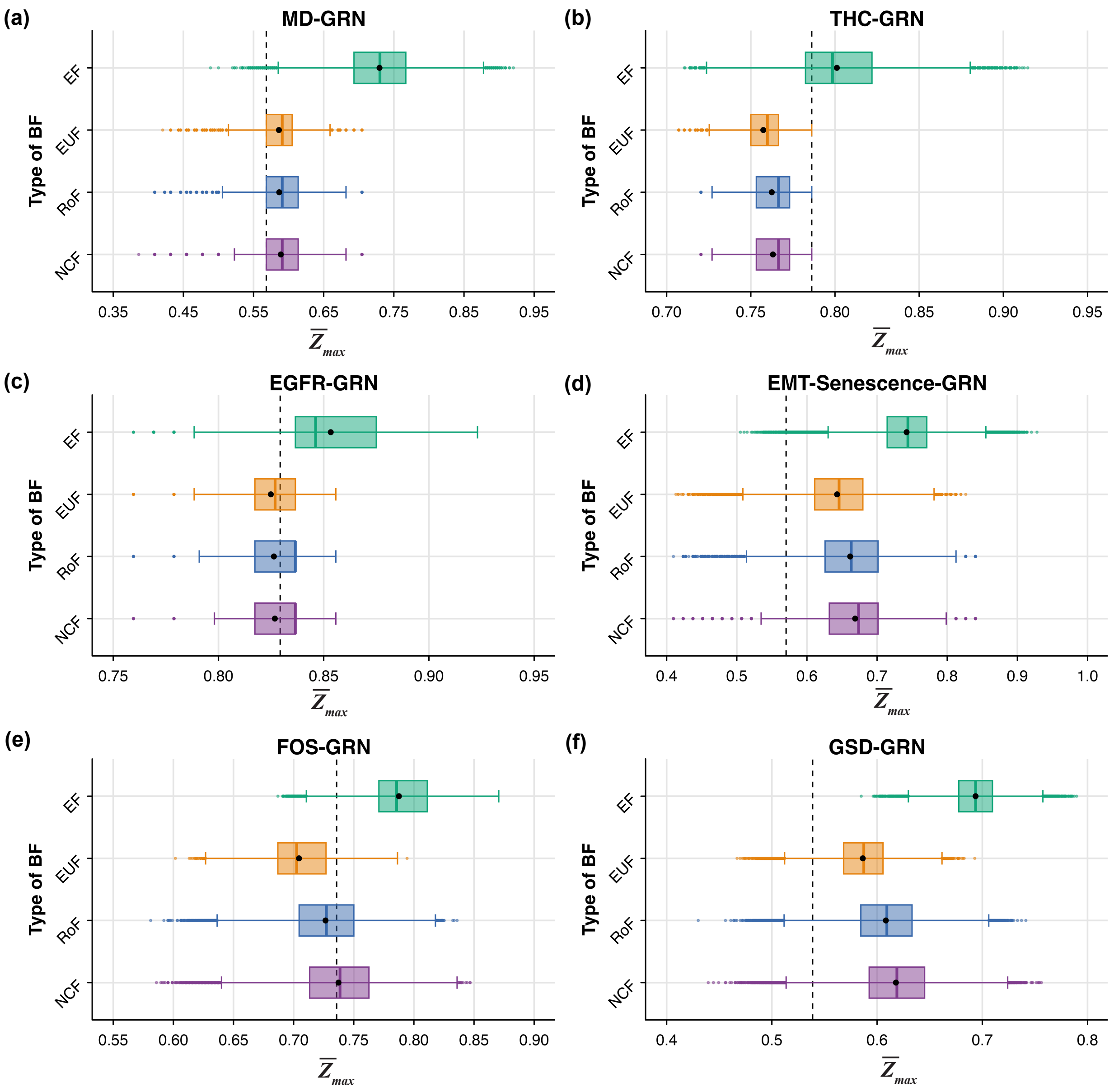


FIG. S13. **Distribution of $\overline{Z}_{max}$ values for different ensembles generated using network structures from 6 published Boolean GRNs**. The subplots **(a)**, **(b)**, **(c)**, **(d)**, **(e)** and **(f)** correspond to the MD-GRN, THC-GRN, EGFR-GRN, EMT-Senescence-GRN, FOS-GRN and GSD-GRN network structures respectively. The $x$ and $y$ axes of each subplot correspond respectively to $\overline{Z}_{max}$ and to the type of BF imposed on the nodes of the networks. The box plots display the distribution of $\overline{Z}_{max}$ in the 4 ensembles that each use a given type of regulatory logic. The ensembles generated using biologically meaningful BFs (EUFs, RoFs, NCFs) have lower $\overline{Z}_{max}$ compared to the ensemble generated using EFs. Note that we impose that ensembles recover the biological fixed points of the associated published model. The mean and median of the distribution are indicated by the black dot and the vertical line within the box respectively. The vertical dashed lines in each subplot correspond to the $\overline{Z}_{max}$ of the Boolean model provided by the modelers in the published article.

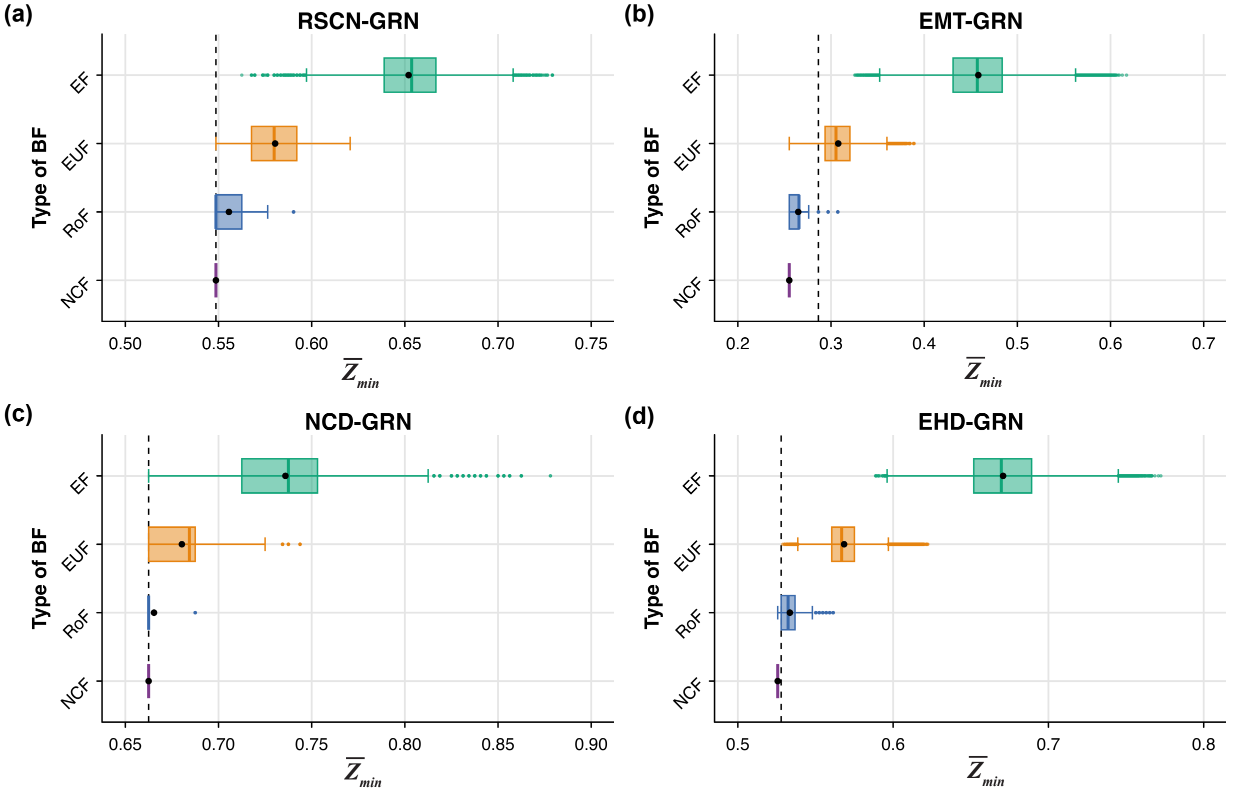


FIG. S14. **Distribution of $\overline{Z}_{min}$ values for different ensembles generated using network structures from 4 published Boolean GRNs**. The subplots **(a)**, **(b)**, **(c)** and **(d)** correspond to the RSCN-GRN, EMT-GRN, NCD-GRN and EHD-GRN network structures respectively. The $x$ and $y$ axes of each subplot correspond respectively to $\overline{Z}_{min}$ and to the type of BF imposed on the nodes of the networks. The box plots display the distribution of $\overline{Z}_{min}$ in the 4 ensembles that each use a given type of regulatory logic. The ensembles generated using biologically meaningful BFs (EUFs, RoFs, NCFs) have lower $\overline{Z}_{min}$ compared to the ensemble generated using EFs. Note that we impose that ensembles recover the biological fixed points of the associated published model. The mean and median of the distribution are indicated by the black dot and the vertical line within the box respectively. The vertical dashed lines in each subplot correspond to the $\overline{Z}_{min}$ of the Boolean model provided by the modelers in the published article.

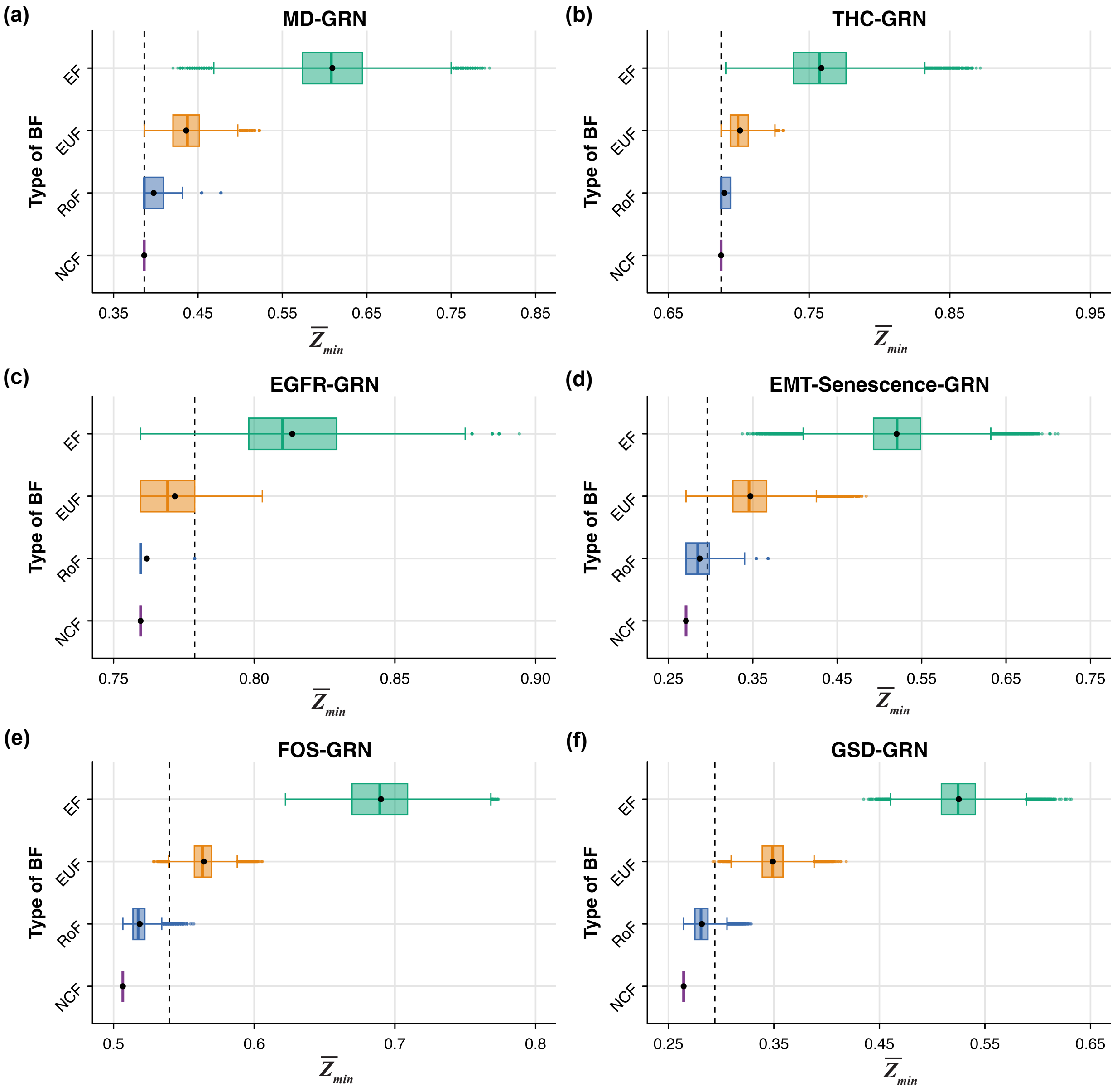


FIG. S15. **Distribution of $\overline{Z}_{min}$ values for different ensembles generated using network structures from 6 published Boolean GRNs**. The subplots **(a)**, **(b)**, **(c)**, **(d)**, **(e)** and **(f)** correspond to the MD-GRN, THC-GRN, EGFR-GRN, EMT-Senescence-GRN, FOS-GRN and GSD-GRN network structures respectively. The $x$ and $y$ axes of each subplot correspond respectively to $\overline{Z}_{min}$ and to the type of BF imposed on the nodes of the networks. The box plots display the distribution of $\overline{Z}_{min}$ in the 4 ensembles that each use a given type of regulatory logic. The ensembles generated using biologically meaningful BFs (EUFs, RoFs, NCFs) have lower $\overline{Z}_{min}$ compared to the ensemble generated using EFs. Note that we impose that ensembles recover the biological fixed points of the associated published model. The mean and median of the distribution are indicated by the black dot and the vertical line within the box respectively. The vertical dashed lines in each subplot correspond to the $\overline{Z}_{min}$ of the Boolean model provided by the modelers in the published article.

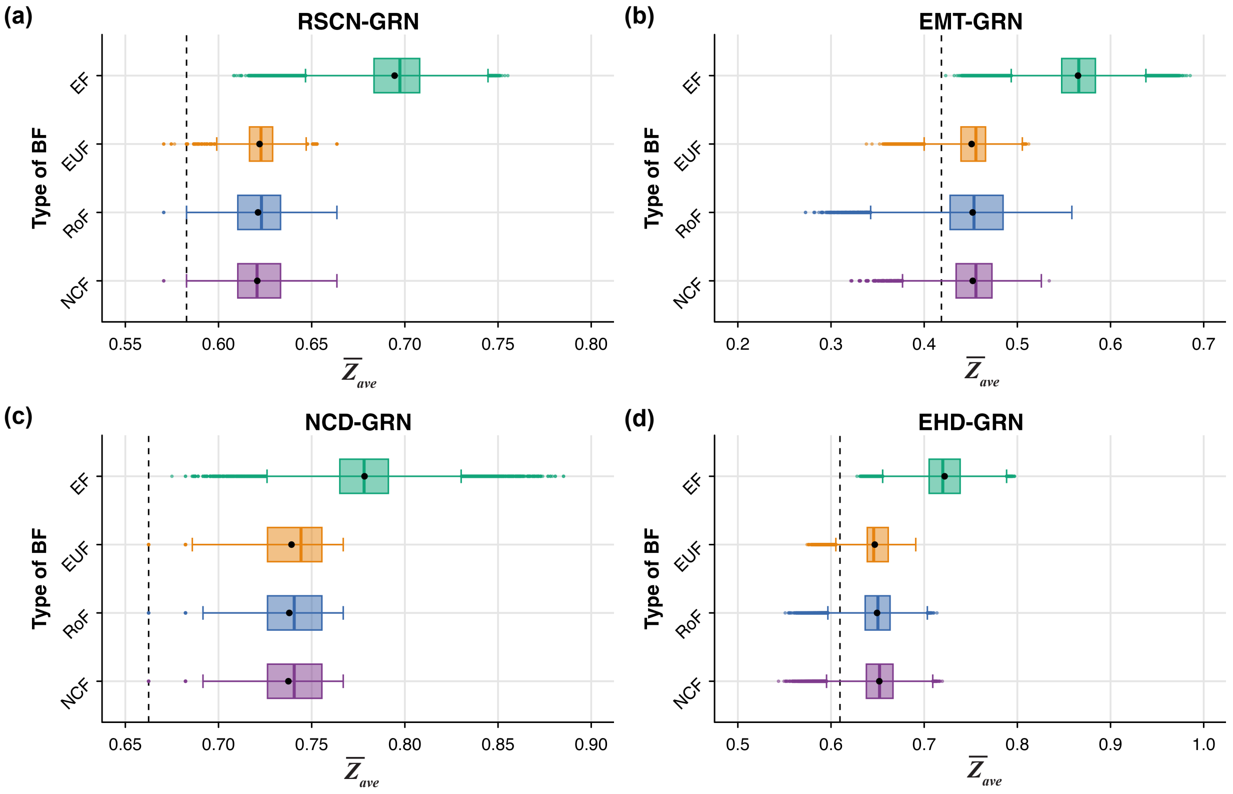


FIG. S16. **Distribution of $\overline{Z}_{ave}$ values for different ensembles generated using network structures from 4 published Boolean GRNs**. The subplots **(a)**, **(b)**, **(c)** and **(d)** correspond to the RSCN-GRN, EMT-GRN, NCD-GRN and EHD-GRN network structures respectively. The $x$ and $y$ axes of each subplot correspond respectively to $\overline{Z}_{ave}$ and to the type of BF imposed on the nodes of the networks. The box plots display the distribution of $\overline{Z}_{ave}$ in the 4 ensembles that each use a given type of regulatory logic. The ensembles generated using biologically meaningful BFs (EUFs, RoFs, NCFs) have lower $\overline{Z}_{ave}$ compared to the ensemble generated using EFs. Note that we impose that ensembles recover the biological fixed points of the associated published model. The mean and median of the distribution are indicated by the black dot and the vertical line within the box respectively. The vertical dashed lines in each subplot correspond to the $\overline{Z}_{ave}$ of the Boolean model provided by the modelers in the published article.

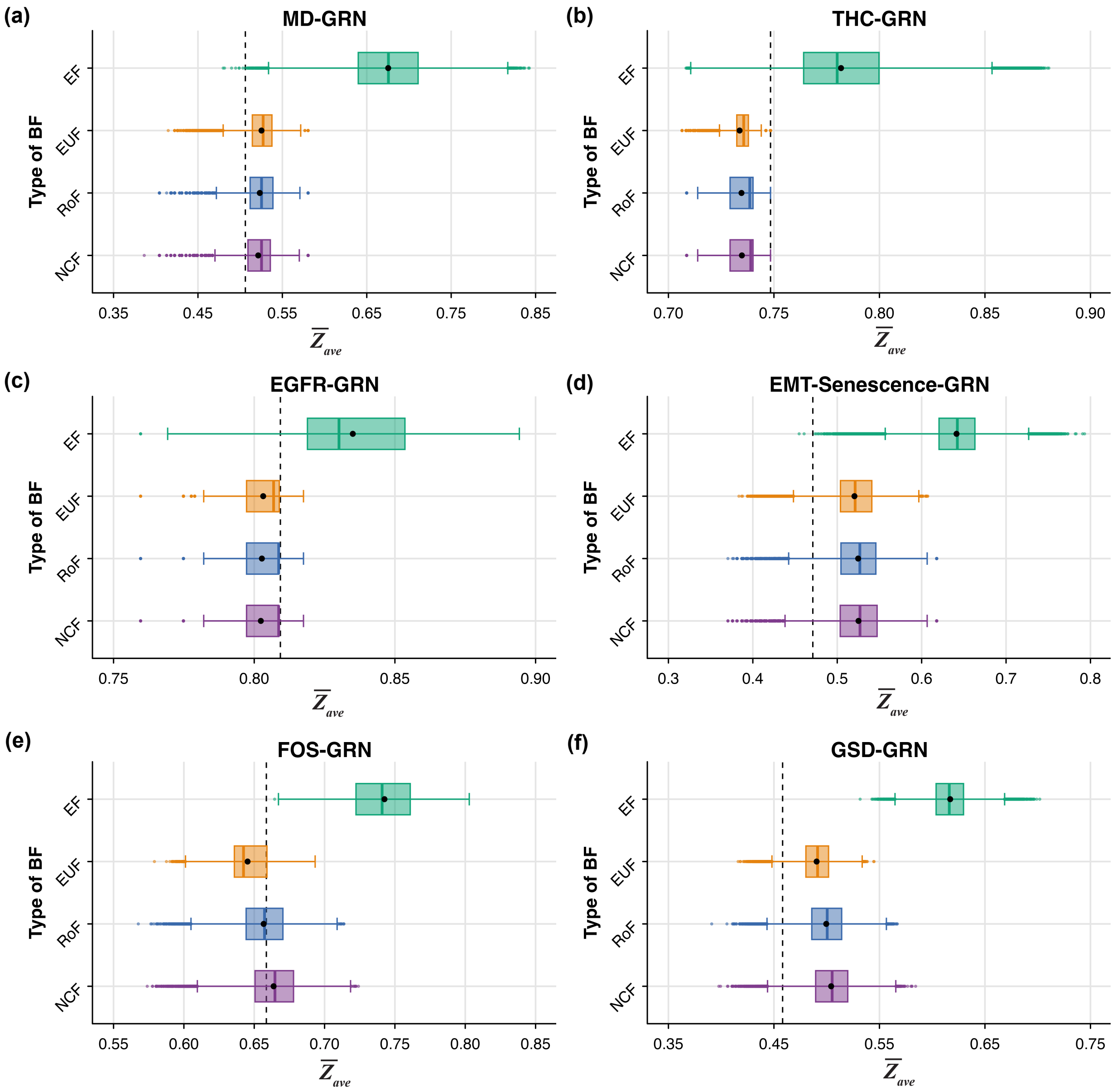


FIG. S17. **Distribution of $\overline{Z}_{ave}$ values for different ensembles generated using network structures from 6 published Boolean GRNs**. The subplots **(a)**, **(b)**, **(c)**, **(d)**, **(e)** and **(f)** correspond to the MD-GRN, THC-GRN, EGFR-GRN, EMT-Senescence-GRN, FOS-GRN and GSD-GRN network structures respectively. The $x$ and $y$ axes of each subplot correspond respectively to $\overline{Z}_{ave}$ and to the type of BF imposed on the nodes of the networks. The box plots display the distribution of $\overline{Z}_{ave}$ in the 4 ensembles that each use a given type of regulatory logic. The ensembles generated using biologically meaningful BFs (EUFs, RoFs, NCFs) have lower $\overline{Z}_{ave}$ compared to the ensemble generated using EFs. Note that we impose that ensembles recover the biological fixed points of the associated published model. The mean and median of the distribution are indicated by the black dot and the vertical line within the box respectively. The vertical dashed lines in each subplot correspond to the $\overline{Z}_{ave}$ of the Boolean model provided by the modelers in the published article.

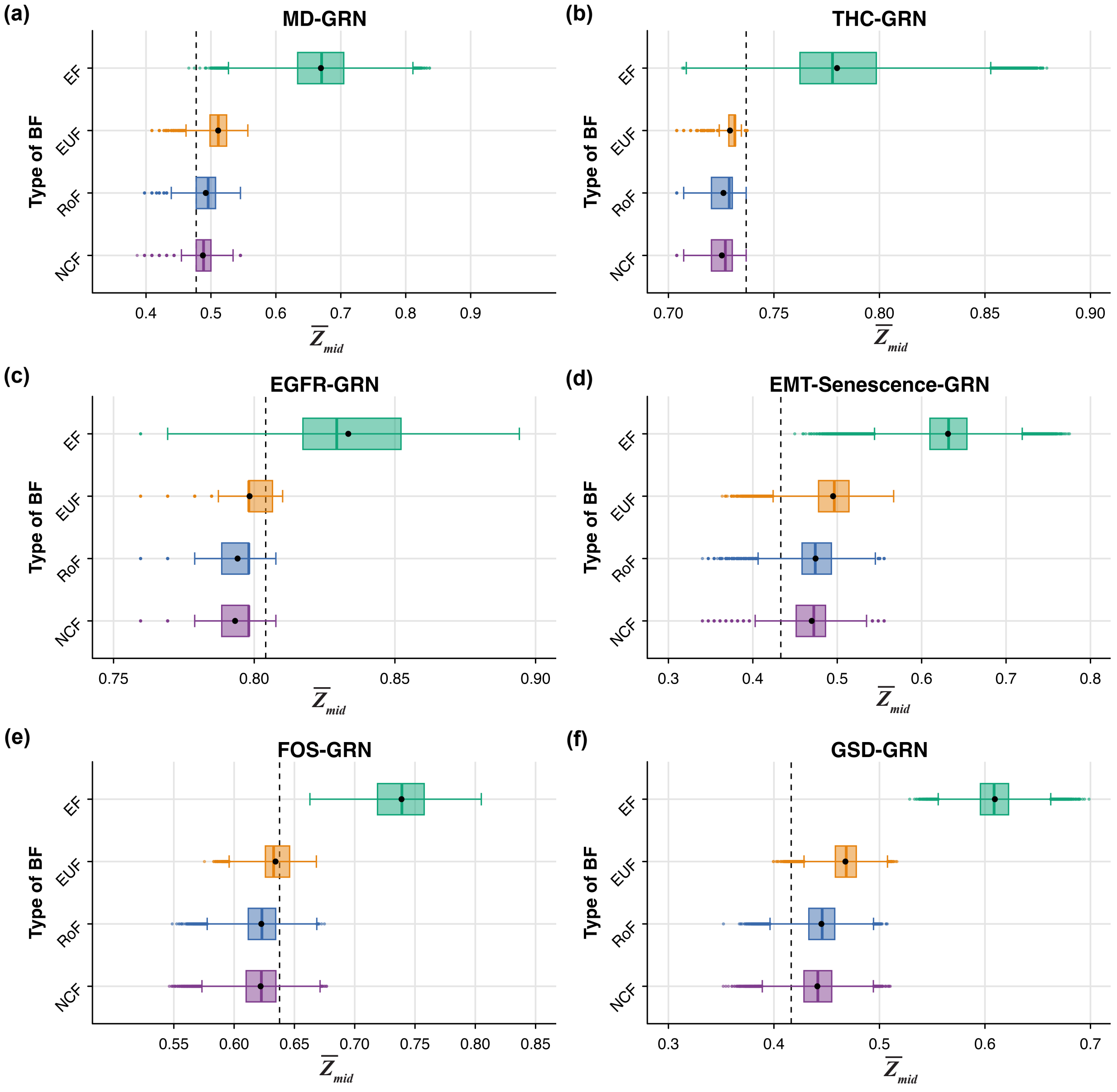


FIG. S18. **Distribution of $\overline{Z}_{mid}$ values for different ensembles generated using network structures from 6 published Boolean GRNs**. The subplots **(a)**, **(b)**, **(c)**, **(d)**, **(e)** and **(f)** correspond to the MD-GRN, THC-GRN, EGFR-GRN, EMT-Senescence-GRN, FOS-GRN and GSD-GRN network structures respectively. The $x$ and $y$ axes of each subplot correspond respectively to $\overline{Z}_{mid}$ and to the type of BF imposed on the nodes of the networks. The box plots display the distribution of $\overline{Z}_{mid}$ in the 4 ensembles that each use a given type of regulatory logic. The ensembles generated using biologically meaningful BFs (EUFs, RoFs, NCFs) have lower $\overline{Z}_{mid}$ compared to the ensemble generated using EFs. Note that we impose that ensembles recover the biological fixed points of the associated published model. The mean and median of the distribution are indicated by the black dot and the vertical line within the box respectively. The vertical dashed lines in each subplot correspond to the $\overline{Z}_{mid}$ of the Boolean model provided by the modelers in the published article.

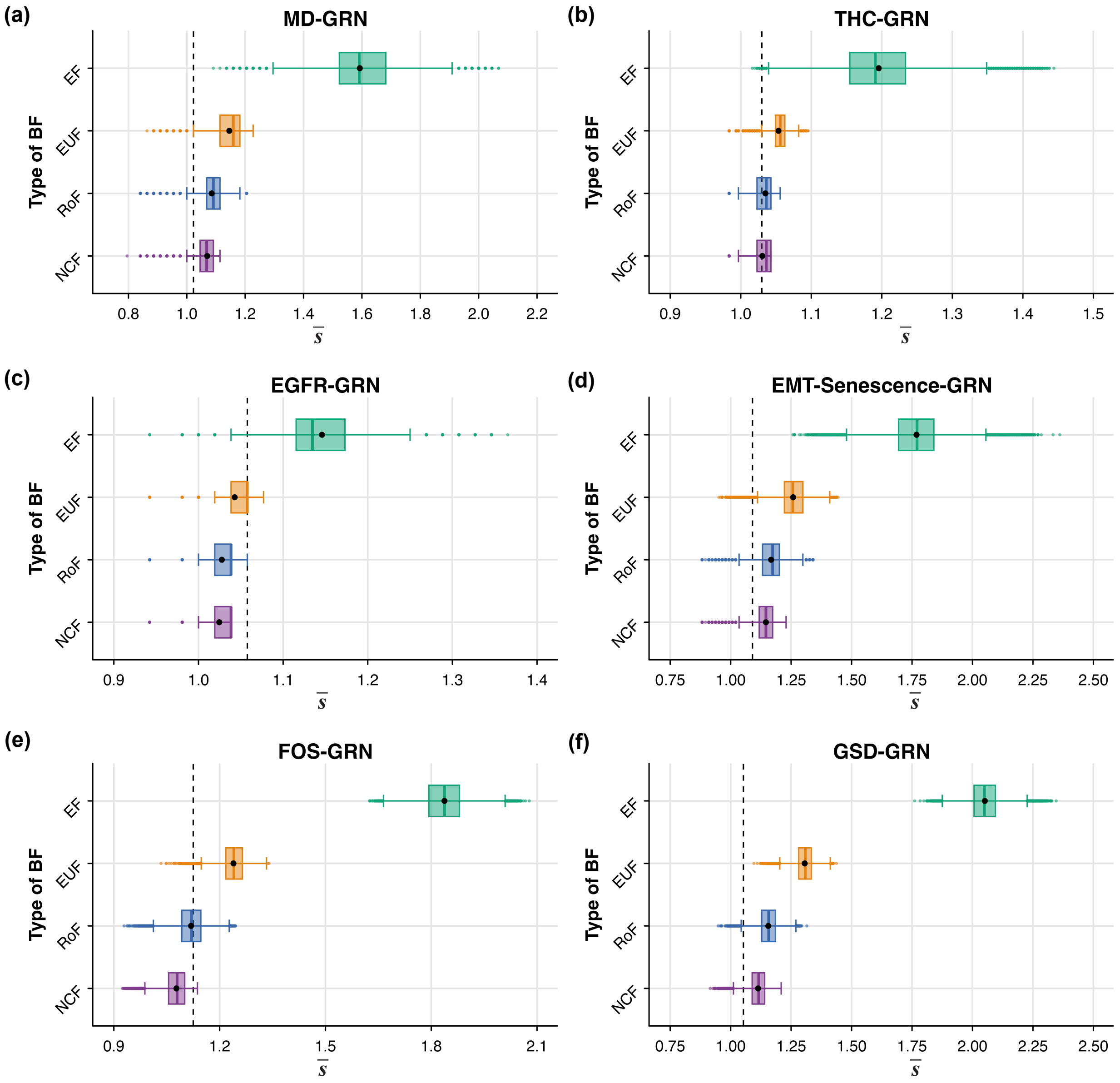


FIG. S19. **Distribution of $\overline{s}$ values for different ensembles generated using network structures from 6 published Boolean GRNs**. The subplots **(a)**, **(b)**, **(c)**, **(d)**, **(e)** and **(f)** correspond to the MD-GRN, THC-GRN, EGFR-GRN, EMT-Senescence-GRN, FOS-GRN and GSD-GRN network structures respectively. The $x$ and $y$ axes of each subplot correspond respectively to $\overline{s}$ and to the type of BF imposed on the nodes of the networks. The box plots display the distribution of $\overline{s}$ in the 4 ensembles that each use a given type of regulatory logic. The ensembles generated using biologically meaningful BFs (EUFs, RoFs, NCFs) have lower $\overline{s}$ compared to the ensemble generated using EFs. Note that we impose that ensembles recover the biological fixed points of the associated published model. The mean and median of the distribution are indicated by the black dot and the vertical line within the box respectively. The vertical dashed lines in each subplot correspond to the $\overline{s}$ of the Boolean model provided by the modelers in the published article.

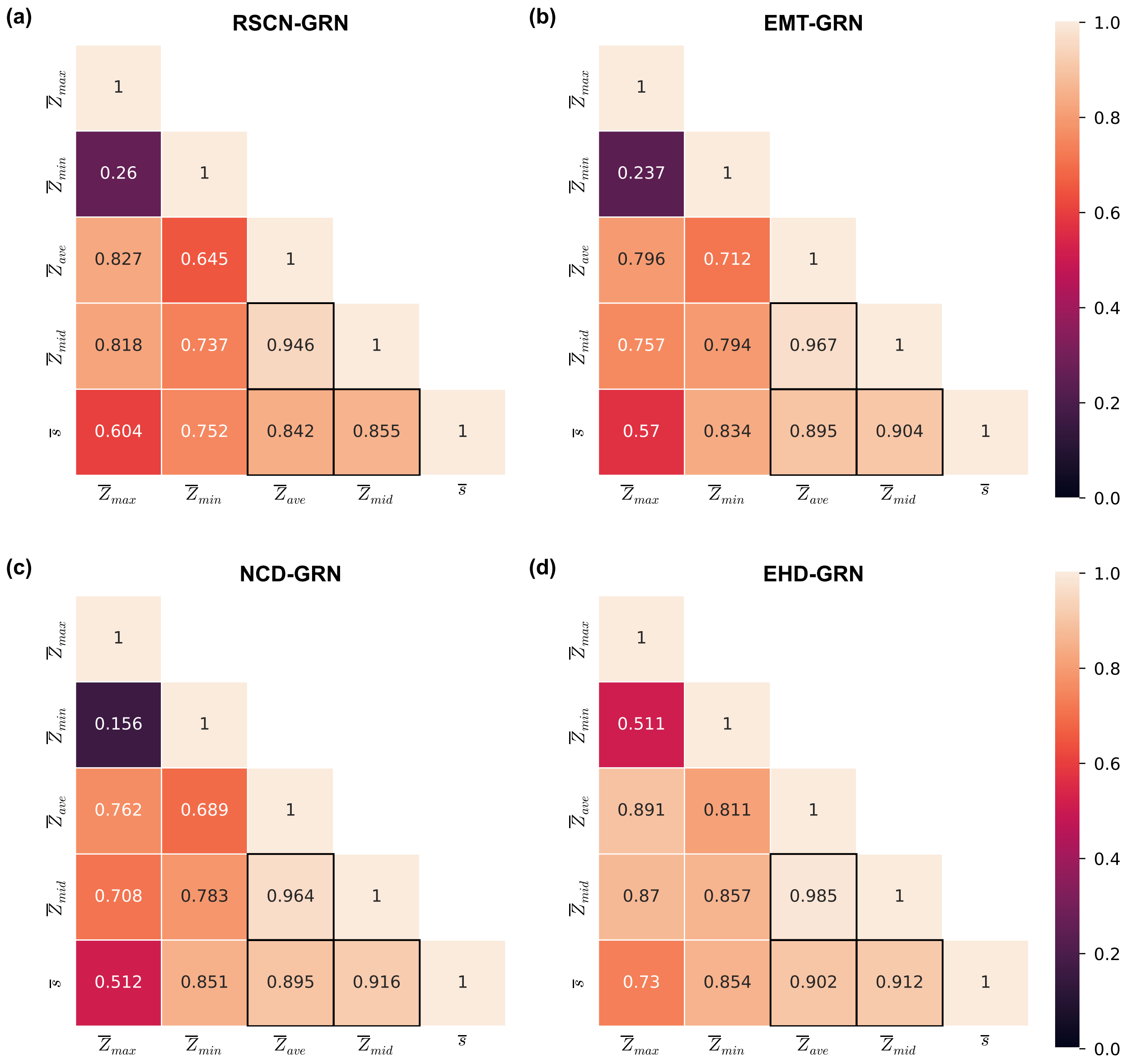


FIG. S20. **Correlation heat map between the various descriptors of local dynamics on ensembles generated using network structures from 4 published Boolean GRNs.** The subplots **(a)**, **(b)**, **(c)** and **(d)** correspond to the RSCN-GRN, EMT-GRN, NCD-GRN and EHD-GRN network structures respectively. The EF-ensemble for each network structure was used to generate the heat maps. The rows and columns of each subplot correspond to descriptors of local dynamics, namely, the network $Z$-parameters ($\overline{Z}_{max}$, $\overline{Z}_{min}$, $\overline{Z}_{ave}$, $\overline{Z}_{mid}$) and network sensitivity ($\overline{s}$). The heat maps show the pair-wise Spearman correlations between these descriptors. All pairs of quantities show moderate to a very strong positive correlation across all models, except for one pair that shows weak positive correlation. The darkened grid highlights the descriptors that best capture the global measures of bushiness, namely, $\overline{Z}_{ave}$, $\overline{Z}_{mid}$ and $\overline{s}$.

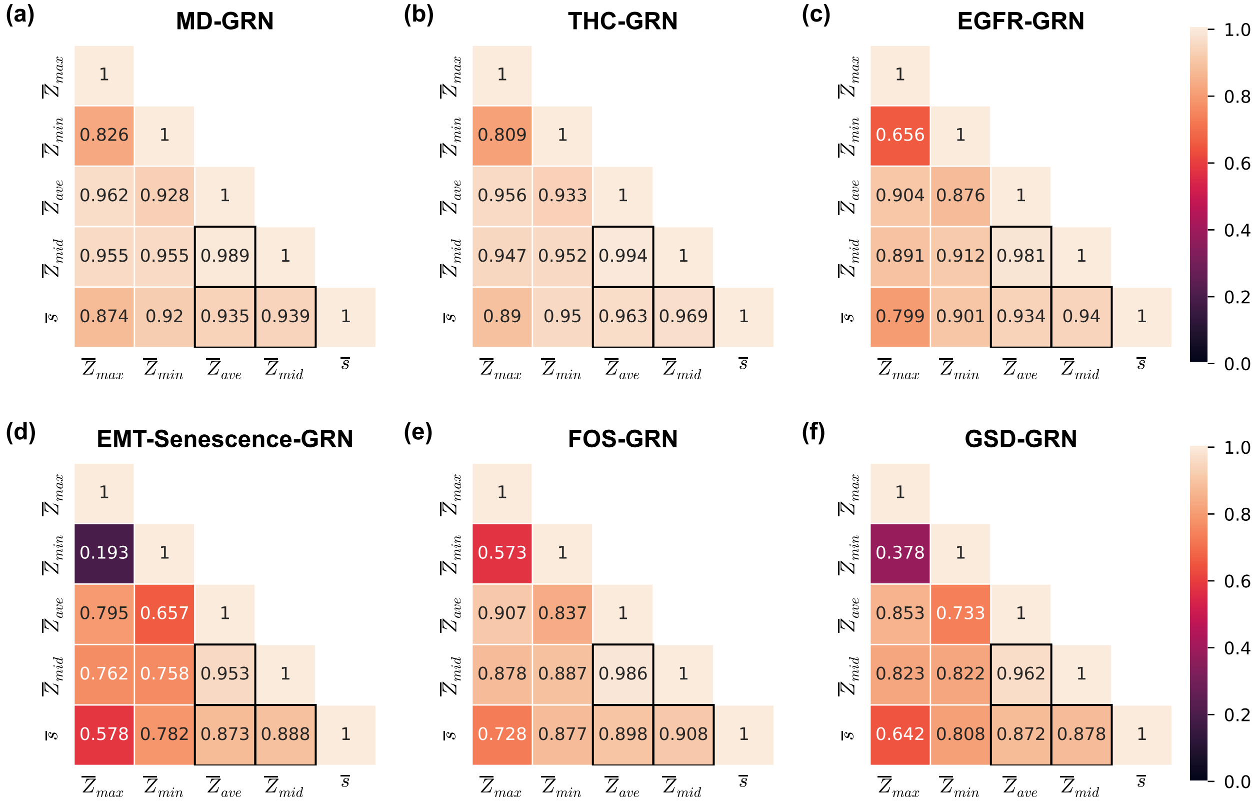


FIG. S21. **Correlation heat map between the various descriptors of local dynamics on ensembles generated using network structures from 6 published Boolean GRNs.** The subplots **(a)**, **(b)**, **(c)**, **(d)**, **(e)** and **(f)** correspond to the MD-GRN, THC-GRN, EGFR-GRN, EMT-Senescence-GRN, FOS-GRN and GSD-GRN network structures respectively. The EF-ensemble for each network structure was used to generate the heat maps. The rows and columns of each subplot correspond to descriptors of local dynamics, namely, the network $Z$-parameters ($\overline{Z}_{max}$, $\overline{Z}_{min}$, $\overline{Z}_{ave}$, $\overline{Z}_{mid}$) and network sensitivity ($\overline{s}$). The heat maps show the pair-wise Spearman correlations between these descriptors. All pairs of quantities typically show moderate to a very strong positive correlation across various network structures, except for the correlations of $\overline{Z}_{max}$ with $\overline{Z}_{min}$ in 2 networks in which it is weak or very weak.

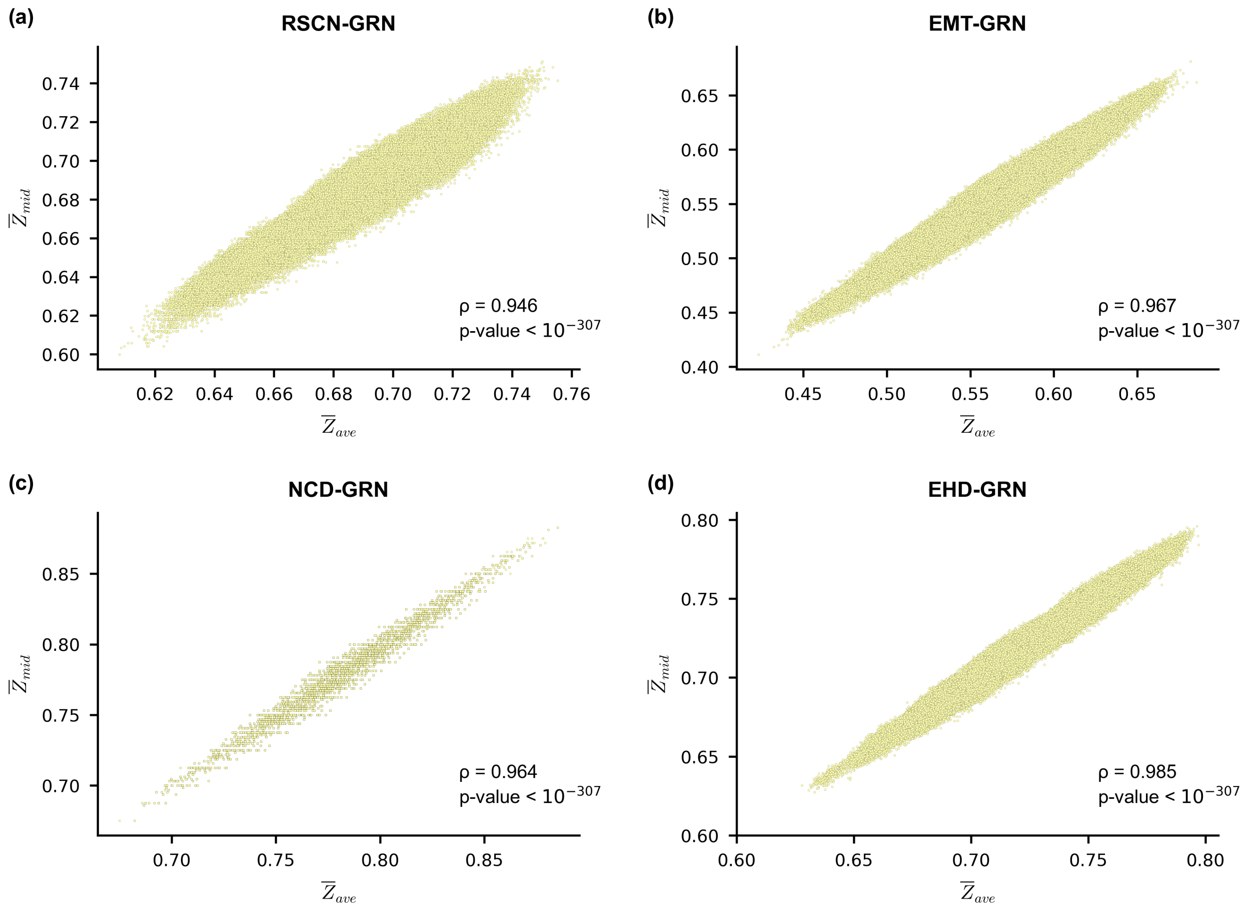


FIG. S22. **Scatter plot between the $\overline{Z}_{mid}$ and $\overline{Z}_{ave}$ on ensembles generated using network structures from 4 published Boolean GRNs.** The subplots **(a)**, **(b)**, **(c)** and **(d)** correspond to the RSCN-GRN, EMT-GRN, NCD-GRN and EHD-GRN network structures respectively. The EF-ensemble for each network structure was used to generate the scatter plots. The $x$ and $y$ axis of each subplot correspond to $\overline{Z}_{mid}$ and $\overline{Z}_{ave}$ respectively. In each subplot we provide the Spearman correlation coefficient ($\rho$) along with the associated p-value. $\overline{Z}_{mid}$ and $\overline{Z}_{ave}$ show a strong positive correlation for all networks.

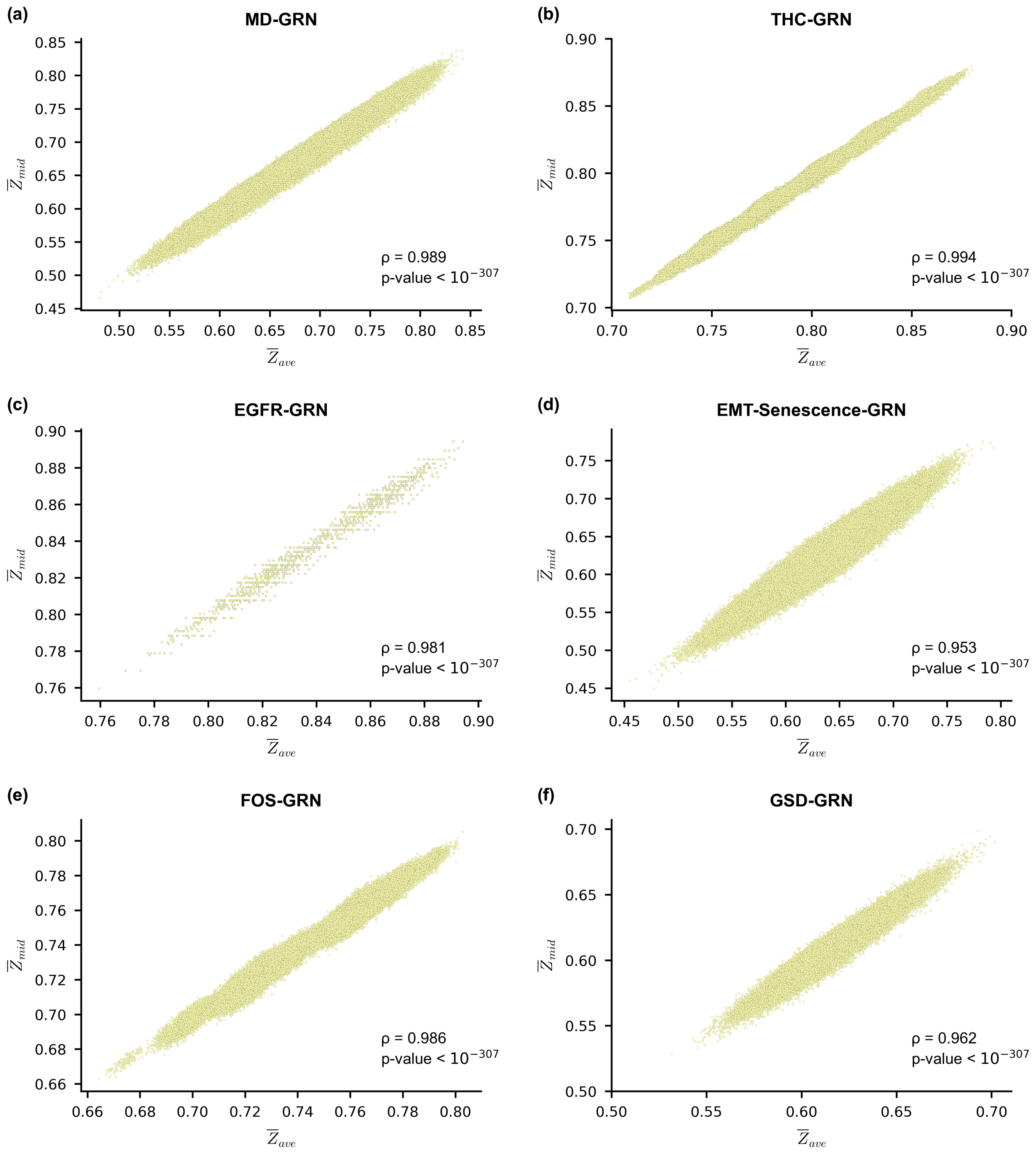


FIG. S23. **Scatter plot between the $\overline{Z}_{mid}$ and $\overline{Z}_{ave}$ on ensembles generated using network structures from 6 published Boolean GRNs.** The subplots **(a)**, **(b)**, **(c)**, **(d)**, **(e)** and **(f)** correspond to the MD-GRN, THC-GRN, EGFR-GRN, EMT-Senescence-GRN, FOS-GRN and GSD-GRN network structures respectively. The EF-ensemble for each network structure was used to generate the scatter plots. The $x$ and $y$ axis of each subplot correspond to $\overline{Z}_{mid}$ and $\overline{Z}_{ave}$ respectively. In each subplot we provide the Spearman correlation coefficient ($\rho$) along with the associated p-value. $\overline{Z}_{mid}$ and $\overline{Z}_{ave}$ show a strong positive correlation for all networks.

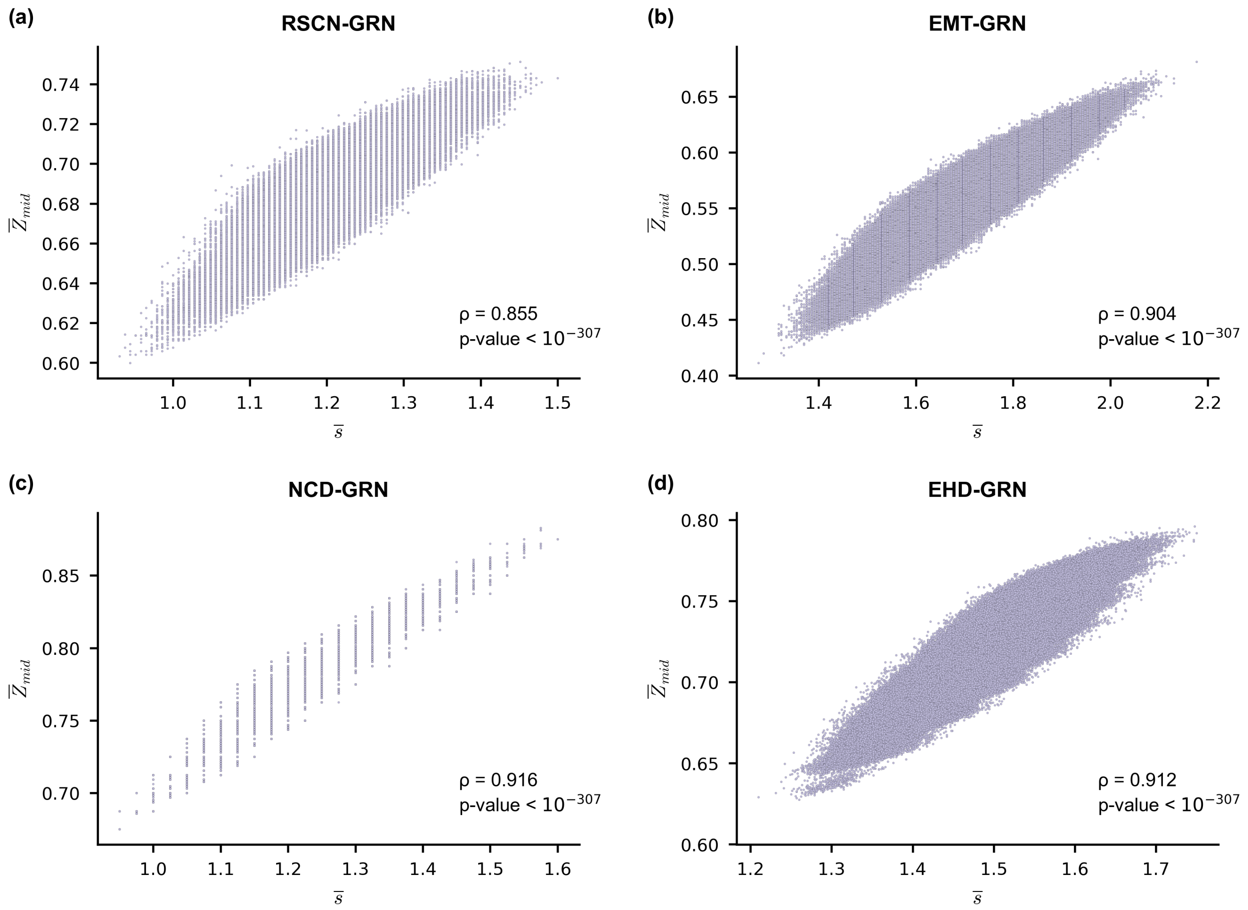


FIG. S24. **Scatter plot between the $\overline{Z}_{mid}$ and network sensitivity ($\overline{s}$) on ensembles generated using network structures from 4 published Boolean GRNs.** The subplots **(a)**, **(b)**, **(c)** and **(d)** correspond to the RSCN-GRN, EMT-GRN, NCD-GRN and EHD-GRN network structures respectively. The EF-ensemble for each network structure was used to generate the scatter plots. The $x$ and $y$ axis of each subplot correspond to $\overline{Z}_{mid}$ and network sensitivity ($\overline{s}$) respectively. In each subplot we provide the Spearman correlation coefficient ($\rho$) along with the associated p-value. $\overline{s}$ and $\overline{Z}_{mid}$ show a strong positive correlation for all networks.

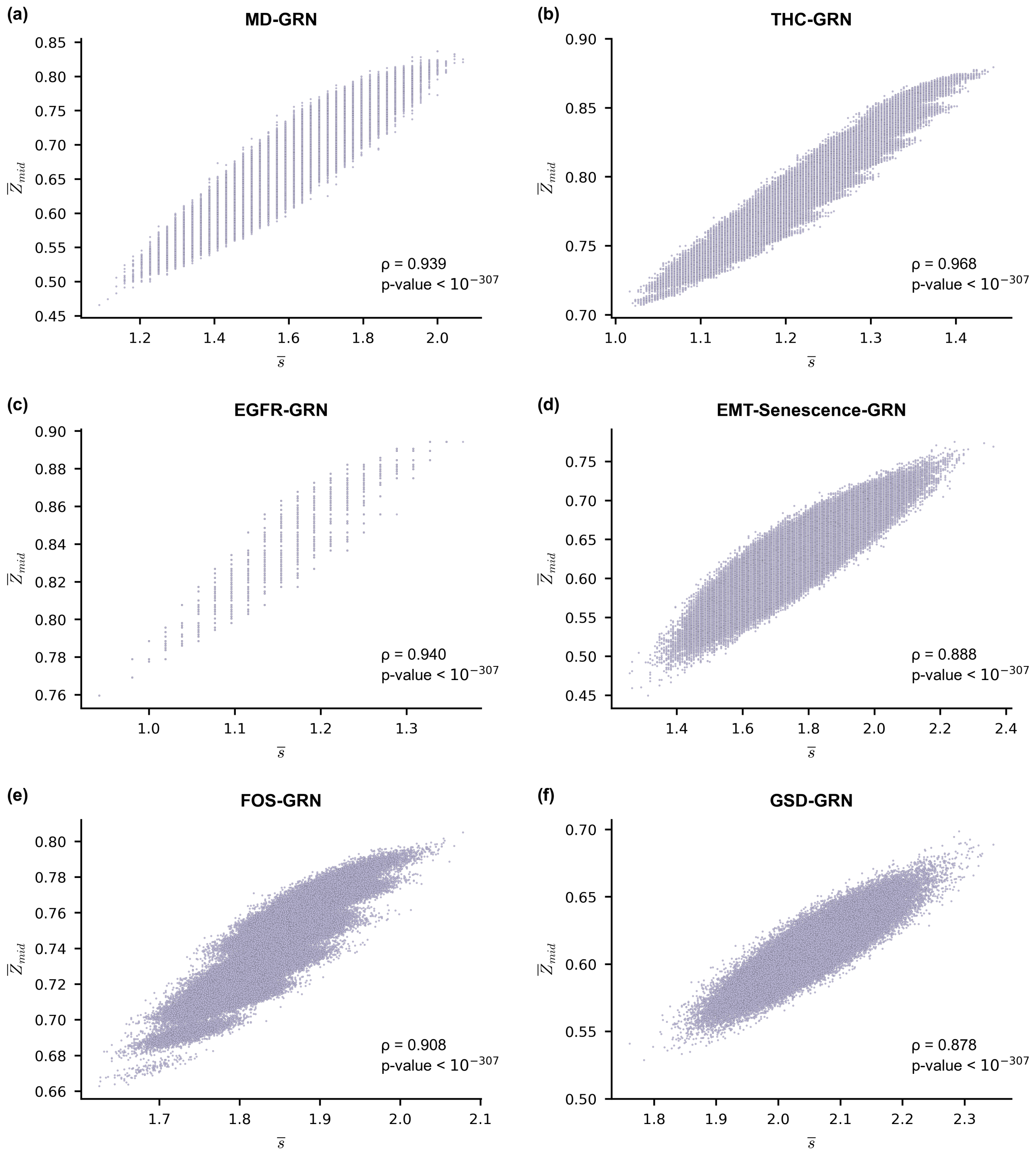


FIG. S25. **Scatter plot between the $\overline{Z}_{mid}$ and network sensitivity ($\overline{s}$) on ensembles generated using network structures from 6 published Boolean GRNs.** The subplots **(a)**, **(b)**, **(c)**, **(d)**, **(e)** and **(f)** correspond to the MD-GRN, THC-GRN, EGFR-GRN, EMT-Senescence-GRN, FOS-GRN and GSD-GRN network structures respectively. The EF-ensemble for each network structure was used to generate the scatter plots. The $x$ and $y$ axis of each subplot correspond to $\overline{Z}_{mid}$ and network sensitivity ($\overline{s}$) respectively. In each subplot we provide the Spearman correlation coefficient ($\rho$) along with the associated p-value. $\overline{s}$ and $\overline{Z}_{mid}$ show a strong positive correlation for all networks.

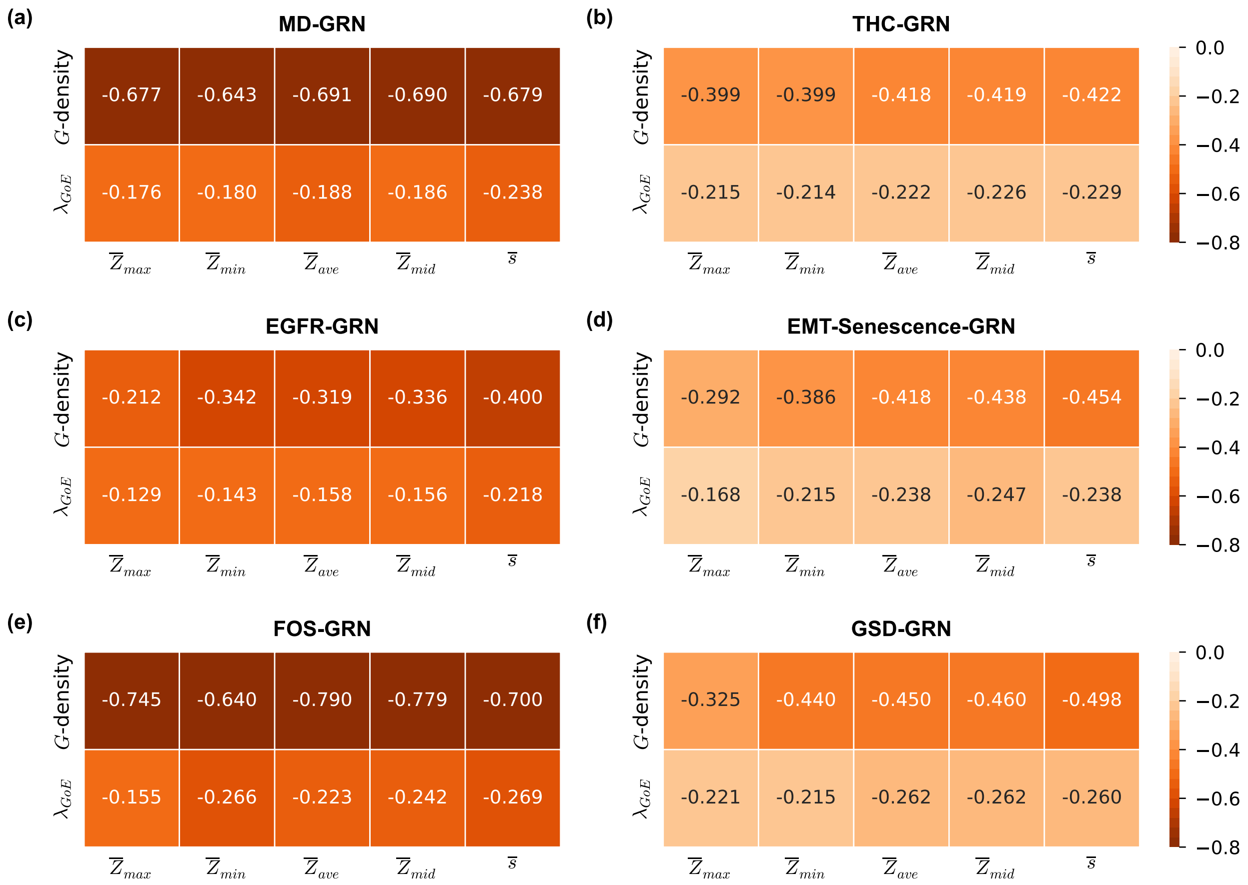


FIG. S26. **Correlation heat map between descriptors of local dynamics and global measures of bushiness in ensembles generated using network structures from 6 published Boolean GRNs.** The subplots **(a)**, **(b)**, **(c)**, **(d)**, **(e)** and **(f)** correspond to the MD-GRN, THC-GRN, EGFR-GRN, EMT-Senescence-GRN, FOS-GRN and GSD-GRN network structures respectively. The EF-ensemble for each network structure was used to generate the heat maps. The rows of each subplot correspond to quantities computed on the state transition graph, namely $G$-density and average convergence rate of trajectories originating at $GoE$ states ($\lambda_{GoE}$). The columns of each subplot correspond to descriptors of local dynamics, namely, the network $Z$-parameters ($\overline{Z}_{max}$, $\overline{Z}_{min}$, $\overline{Z}_{ave}$, $\overline{Z}_{mid}$) and network sensitivity ($\overline{s}$). The heat maps show the pair-wise Spearman correlations between descriptors of local dynamics and global measures of bushiness defined on the state transition graph. The descriptors of local dynamics show a moderate to strong negative correlation with the $G$-density and very weak to moderate negative correlation with $\lambda_{GoE}$.

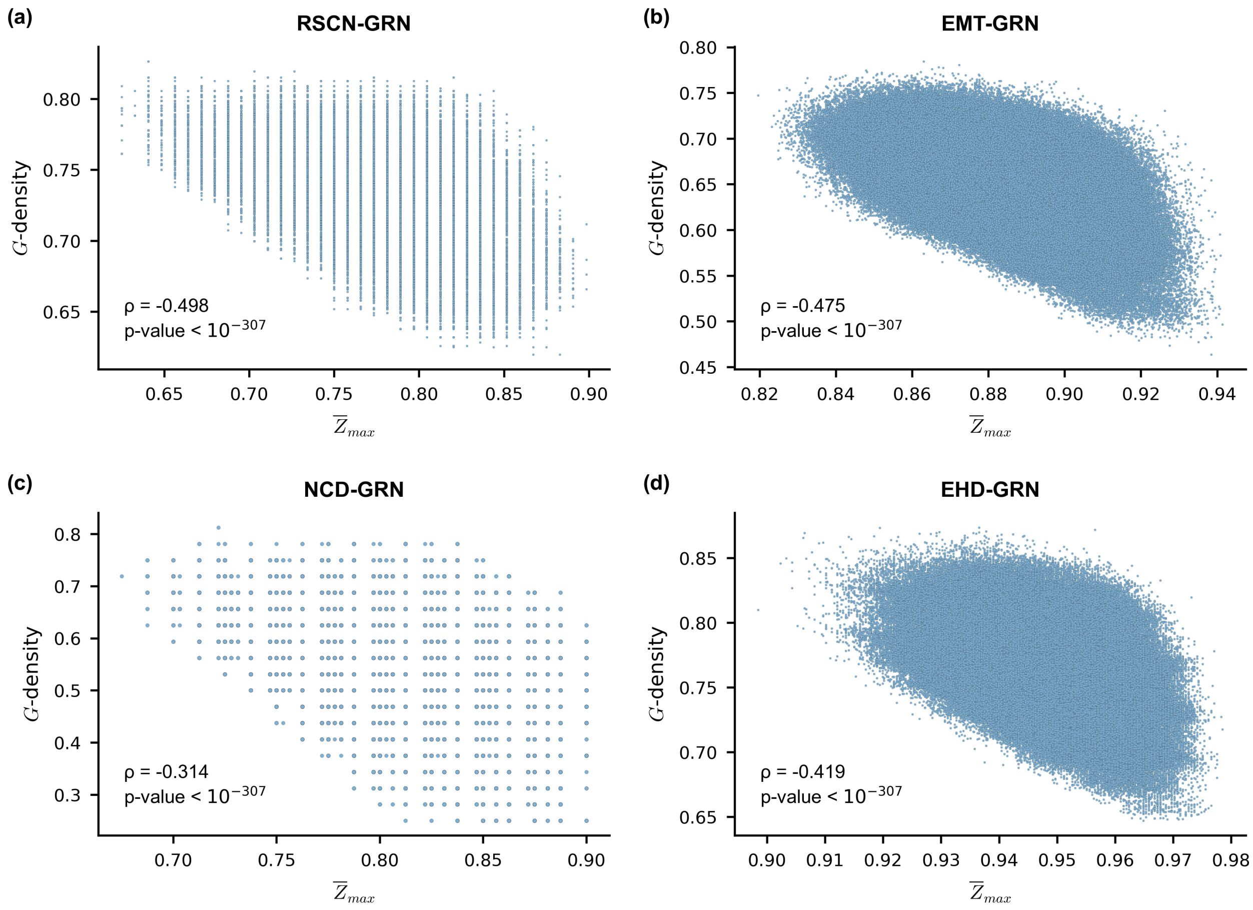


FIG. S27. **Scatter plot between the $\overline{Z}_{max}$ and $G$-density on ensembles generated using network structures from 4 published Boolean GRNs.** The subplots **(a)**, **(b)**, **(c)** and **(d)** correspond to the RSCN-GRN, EMT-GRN, NCD-GRN and EHD-GRN network structures respectively. The EF-ensemble for each network structure was used to generate the scatter plots. The $x$ and $y$ axis of each subplot correspond to $\overline{Z}_{max}$ and $G$-density respectively. In each subplot we provide the Spearman correlation coefficient ($\rho$) along with the associated p-value. $\overline{Z}_{max}$ and $G$-density show a moderate negative correlation for all network except for NCD-GRN for which there is a weak negative correlation.

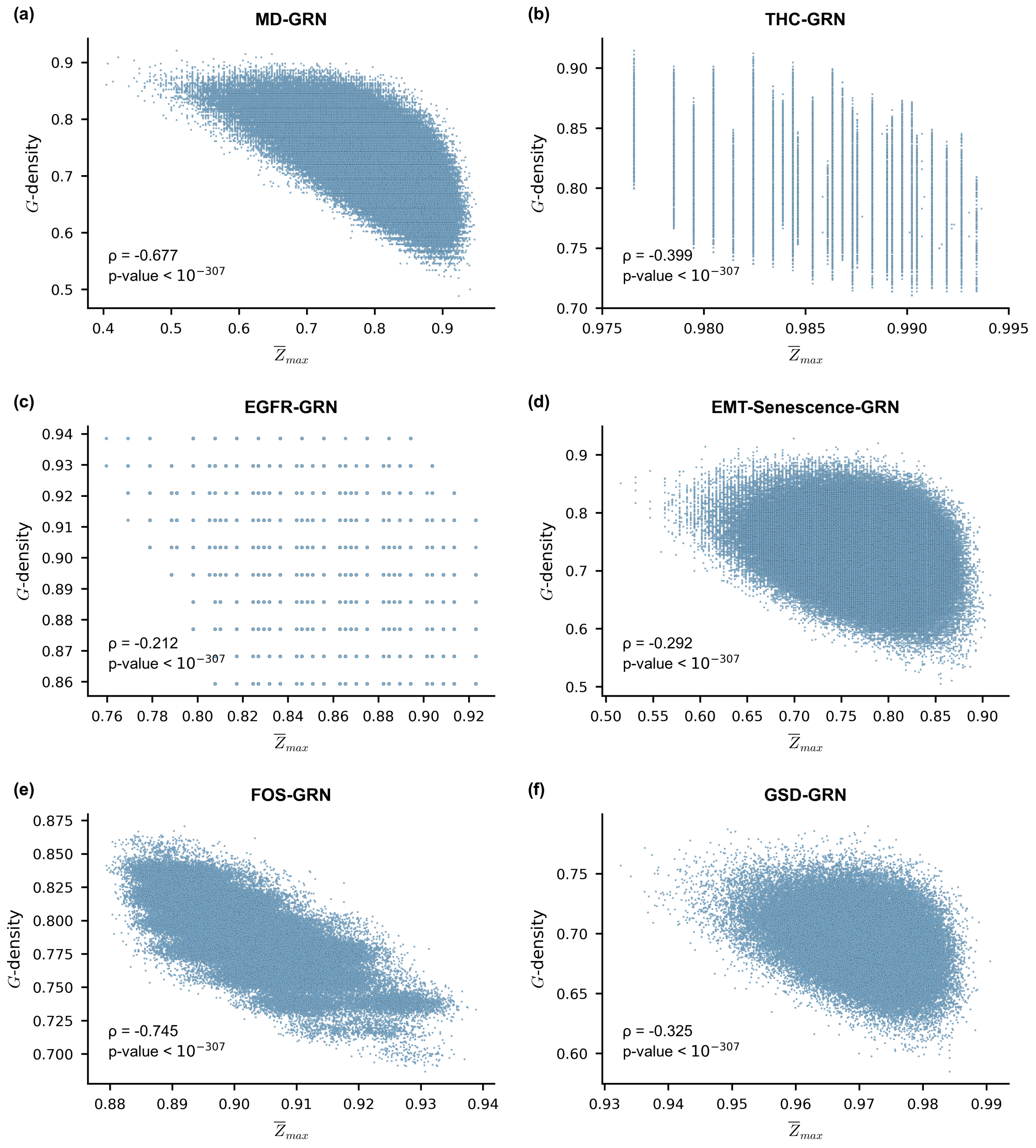


FIG. S28. **Scatter plot between the $\overline{Z}_{max}$ and $G$-density on ensembles generated using network structures from 6 published Boolean GRNs.** The subplots **(a)**, **(b)**, **(c)**, **(d)**, **(e)** and **(f)** correspond to the MD-GRN, THC-GRN, EGFR-GRN, EMT-Senescence-GRN, FOS-GRN and GSD-GRN network structures respectively. The EF-ensemble for each network structure was used to generate the scatter plots. The $x$ and $y$ axis of each subplot correspond to $\overline{Z}_{max}$ and $G$-density respectively. In each subplot we provide the Spearman correlation coefficient ($\rho$) along with the associated p-value. $\overline{Z}_{max}$ and $G$-density show a strong negative correlation for FOS-GRN and MD-GRN and a weak negative correlation otherwise.

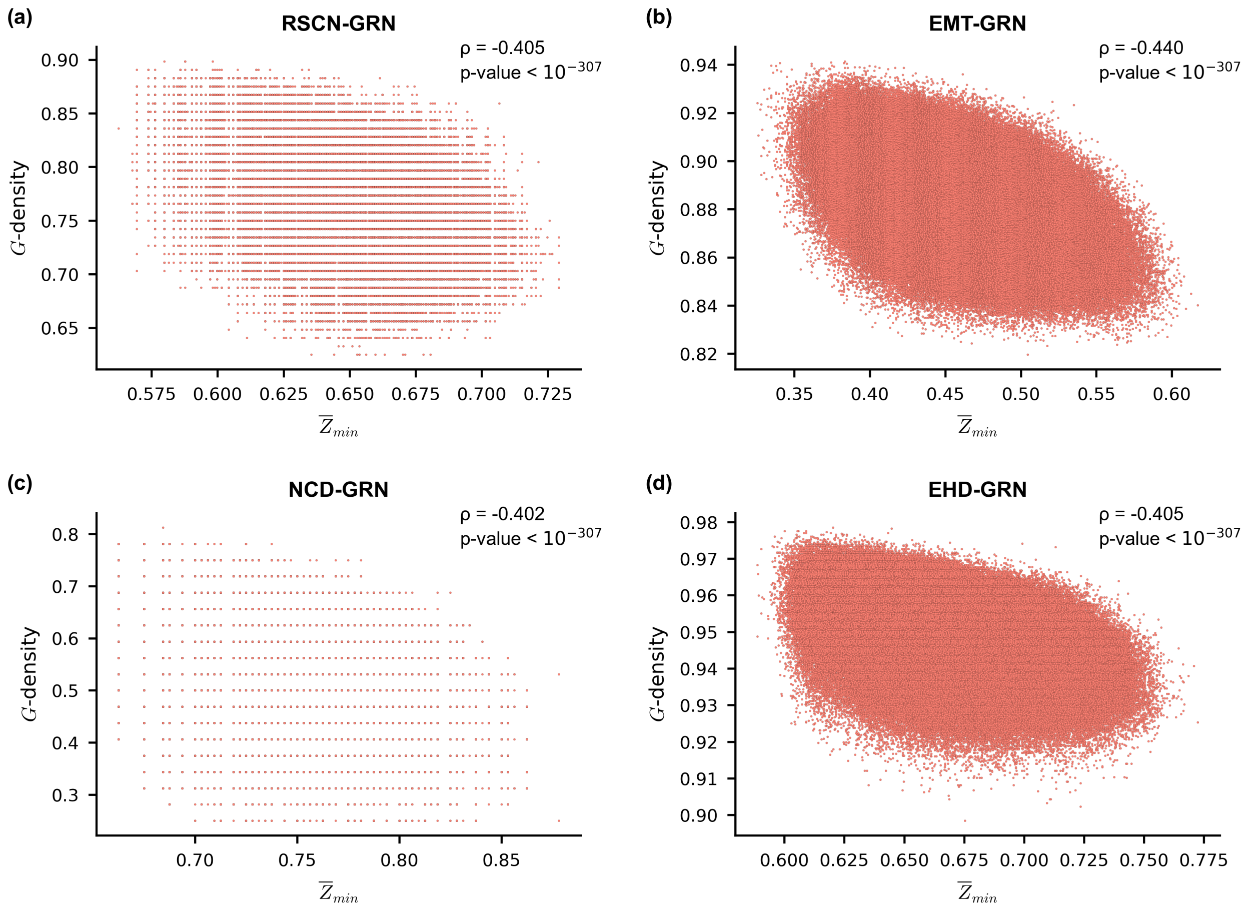


FIG. S29. **Scatter plot between the $\overline{Z}_{min}$ and $G$-density on ensembles generated using network structures from 4 published Boolean GRNs.** The subplots **(a)**, **(b)**, **(c)** and **(d)** correspond to the RSCN-GRN, EMT-GRN, NCD-GRN and EHD-GRN network structures respectively. The EF-ensemble for each network structure was used to generate the scatter plots. The $x$ and $y$ axis of each subplot correspond to $\overline{Z}_{min}$ and $G$-density respectively. In each subplot we provide the Spearman correlation coefficient ($\rho$) along with the associated p-value. $\overline{Z}_{min}$ and $G$-density show a moderate negative correlation for all networks.

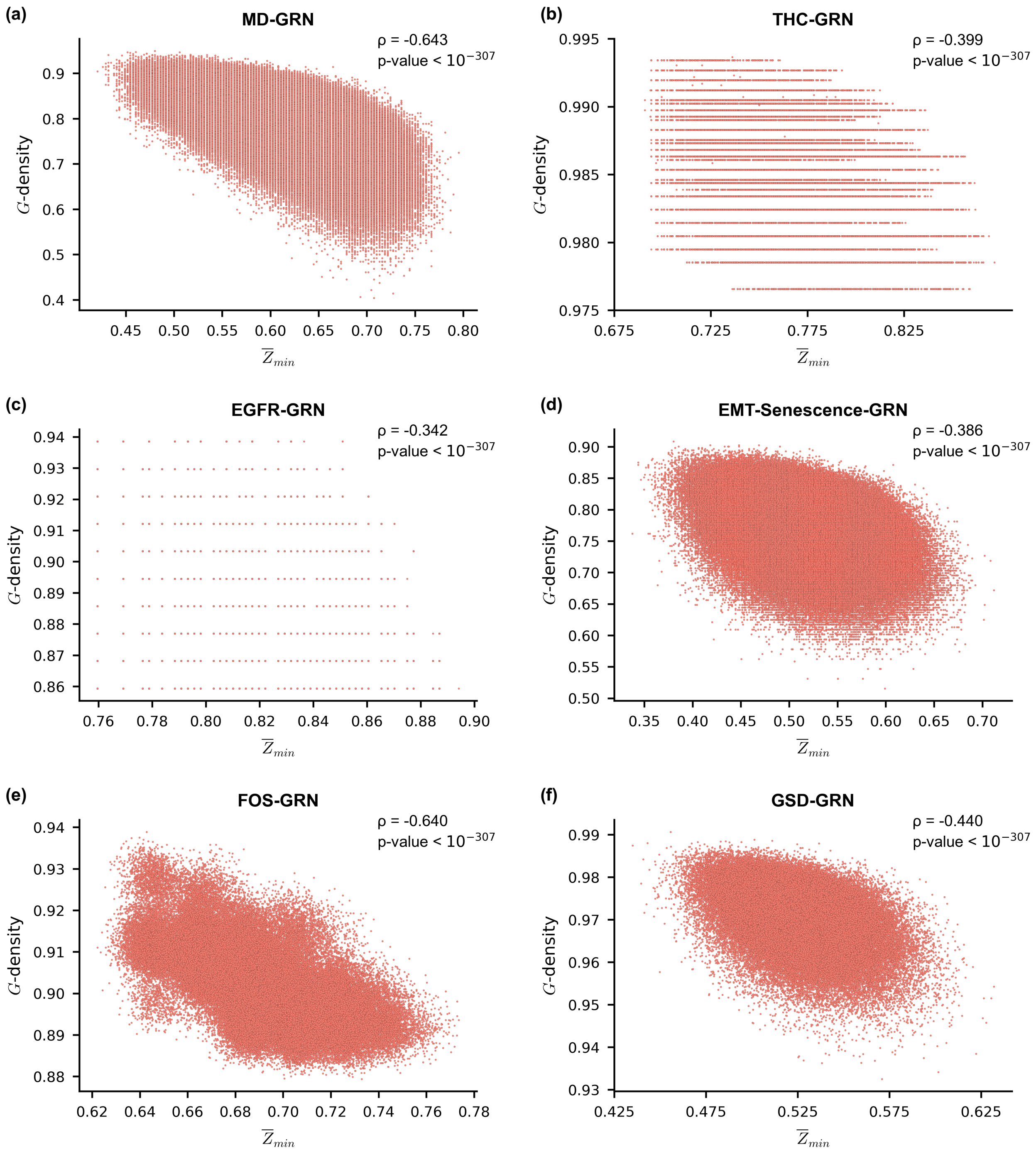


FIG. S30. **Scatter plot between the $\overline{Z}_{min}$ and $G$-density on ensembles generated using network structures from 6 published Boolean GRNs.** The subplots **(a)**, **(b)**, **(c)**, **(d)**, **(e)** and **(f)** correspond to the MD-GRN, THC-GRN, EGFR-GRN, EMT-Senescence-GRN, FOS-GRN and GSD-GRN network structures respectively. The EF-ensemble for each network structure was used to generate the scatter plots. The $x$ and $y$ axis of each subplot correspond to $\overline{Z}_{min}$ and $G$-density respectively. In each subplot we provide the Spearman correlation coefficient ($\rho$) along with the associated p-value. $\overline{Z}_{min}$ and $G$-density show a strong negative correlation for MD-GRN and FOS-GRN, a moderate negative correlation for GSD-GRN and a weak negative correlation otherwise.

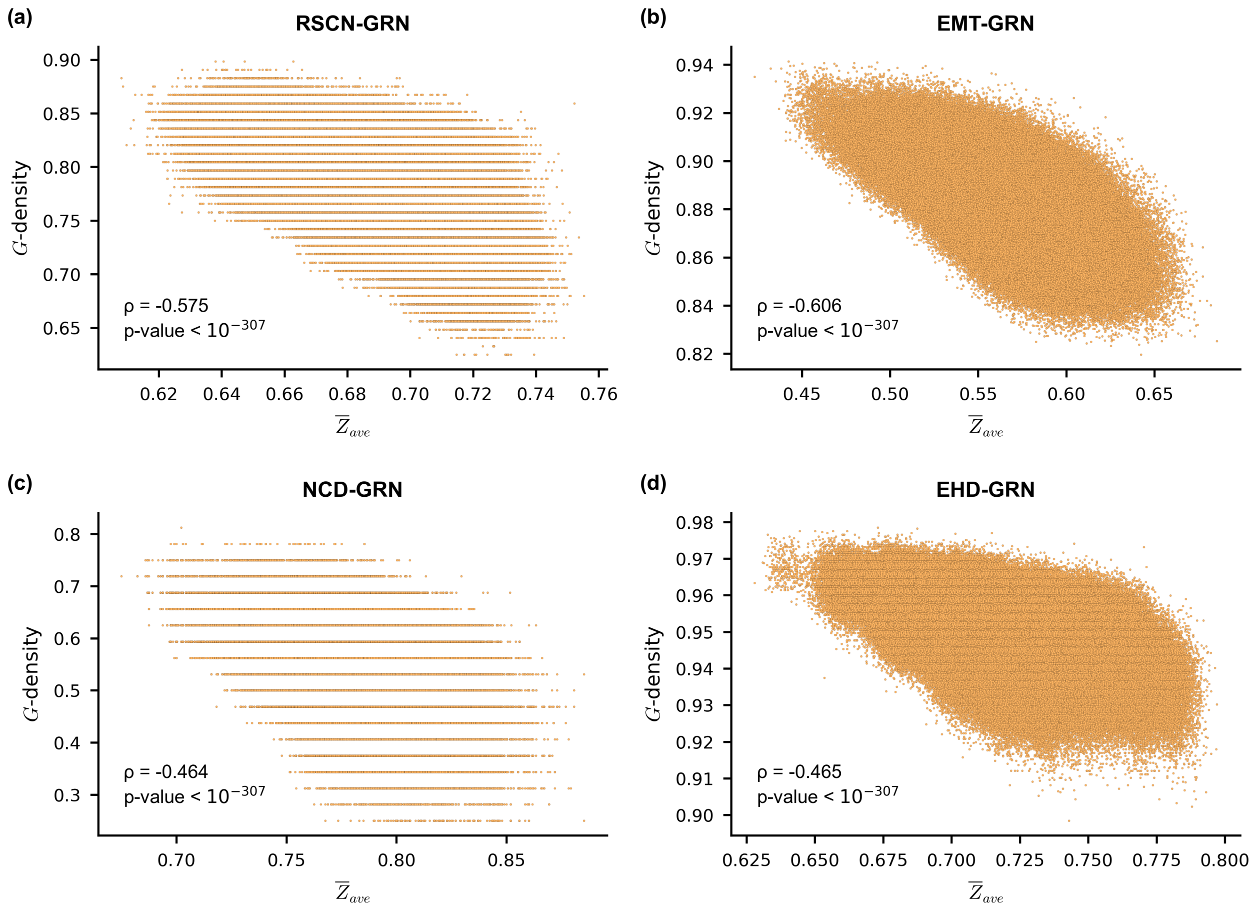


FIG. S31. **Scatter plot between the $\overline{Z}_{ave}$ and $G$-density on ensembles generated using network structures from 4 published Boolean GRNs.** The subplots **(a)**, **(b)**, **(c)** and **(d)** correspond to the RSCN-GRN, EMT-GRN, NCD-GRN and EHD-GRN network structures respectively. The EF-ensemble for each network structure was used to generate the scatter plots. The $x$ and $y$ axis of each subplot correspond to $\overline{Z}_{ave}$ and $G$-density respectively. In each subplot we provide the Spearman correlation coefficient ($\rho$) along with the associated p-value. $\overline{Z}_{ave}$ and $G$-density show a strong negative correlation for EMT-GRN and a moderate negative correlation otherwise.

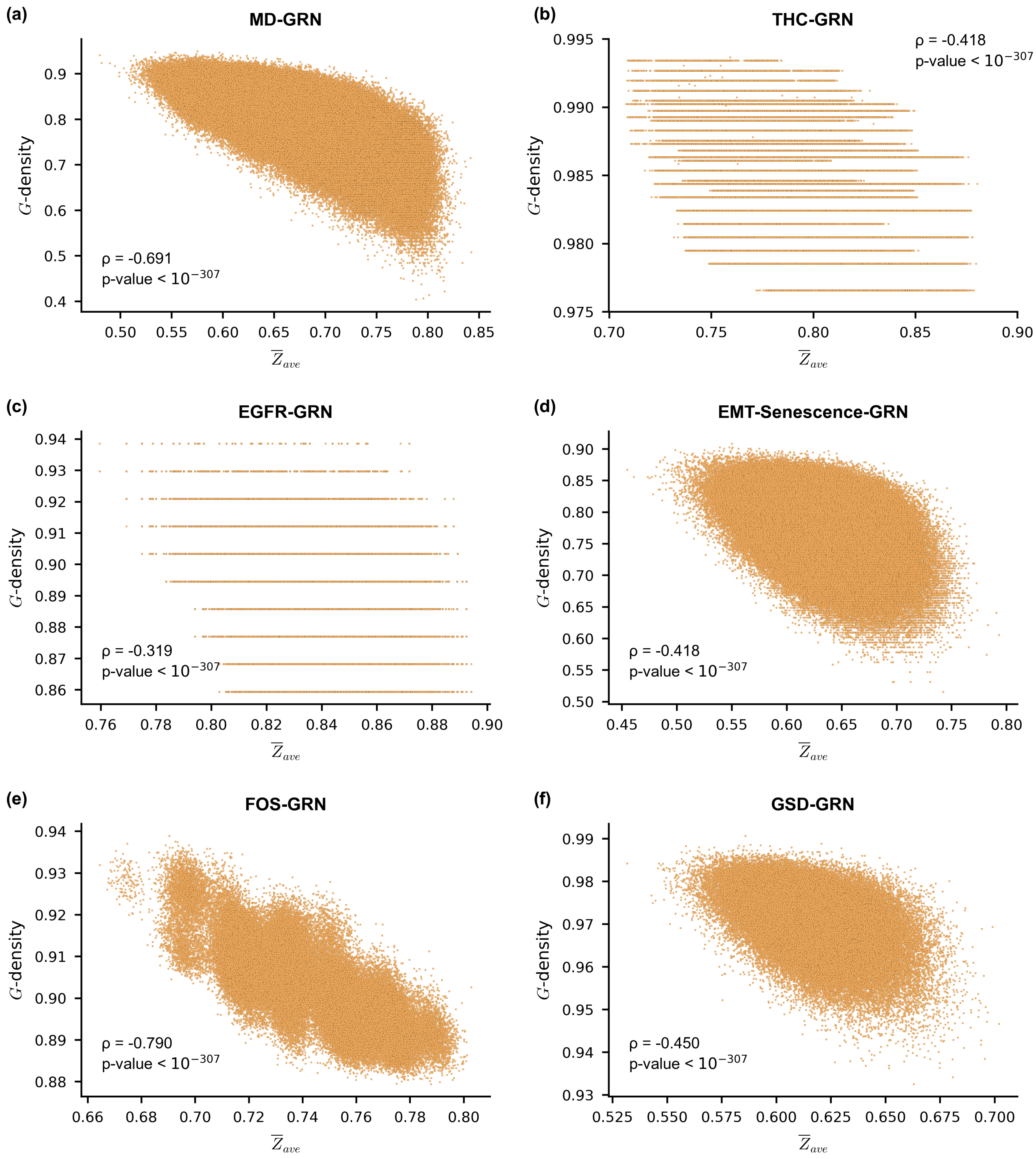


FIG. S32. **Scatter plot between the $\overline{Z}_{ave}$ and $G$-density on ensembles generated using network structures from 6 published Boolean GRNs.** The subplots **(a)**, **(b)**, **(c)**, **(d)**, **(e)** and **(f)** correspond to the MD-GRN, THC-GRN, EGFR-GRN, EMT-Senescence-GRN, FOS-GRN and GSD-GRN network structures respectively. The EF-ensemble for each network structure was used to generate the scatter plots. The $x$ and $y$ axis of each subplot correspond to $\overline{Z}_{ave}$ and $G$-density respectively. In each subplot we provide the Spearman correlation coefficient ($\rho$) along with the associated p-value. $\overline{Z}_{ave}$ and $G$-density show a strong negative correlation for FOS-GRN and MD-GRN, a moderate negative correlation for GSD-GRN, EMT-Senescence-GRN and THC-GRN, and a weak negative correlation for EGFR-GRN.

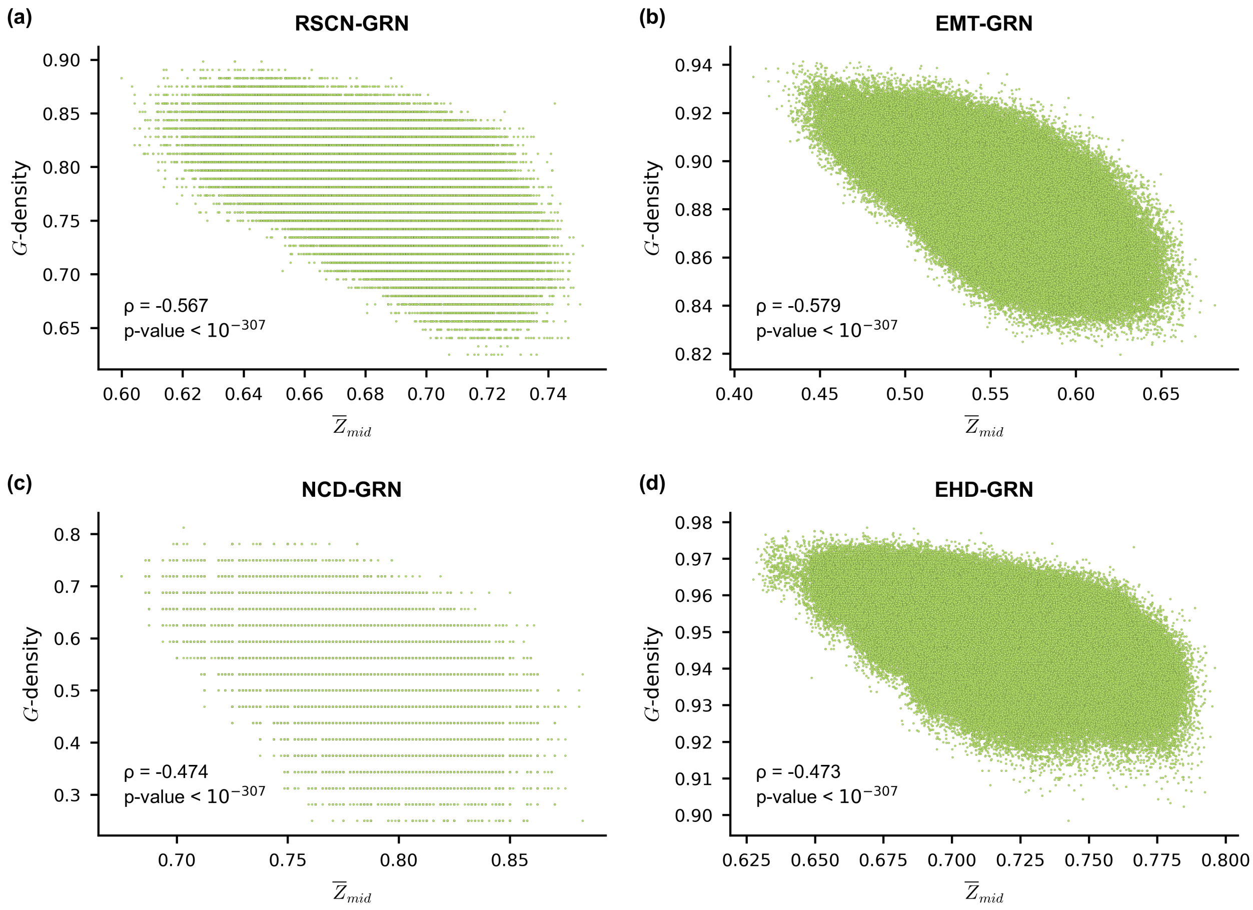


FIG. S33. **Scatter plot between the $\overline{Z}_{mid}$ and $G$-density on ensembles generated using network structures from 4 published Boolean GRNs.** The subplots **(a)**, **(b)**, **(c)** and **(d)** correspond to the RSCN-GRN, EMT-GRN, NCD-GRN and EHD-GRN network structures respectively. The EF-ensemble for each network structure was used to generate the scatter plots. The $x$ and $y$ axis of each subplot correspond to $\overline{Z}_{mid}$ and $G$-density respectively. In each subplot we provide the Spearman correlation coefficient ($\rho$) along with the associated p-value. $\overline{Z}_{mid}$ and $G$-density show a moderate negative correlation for all the networks.

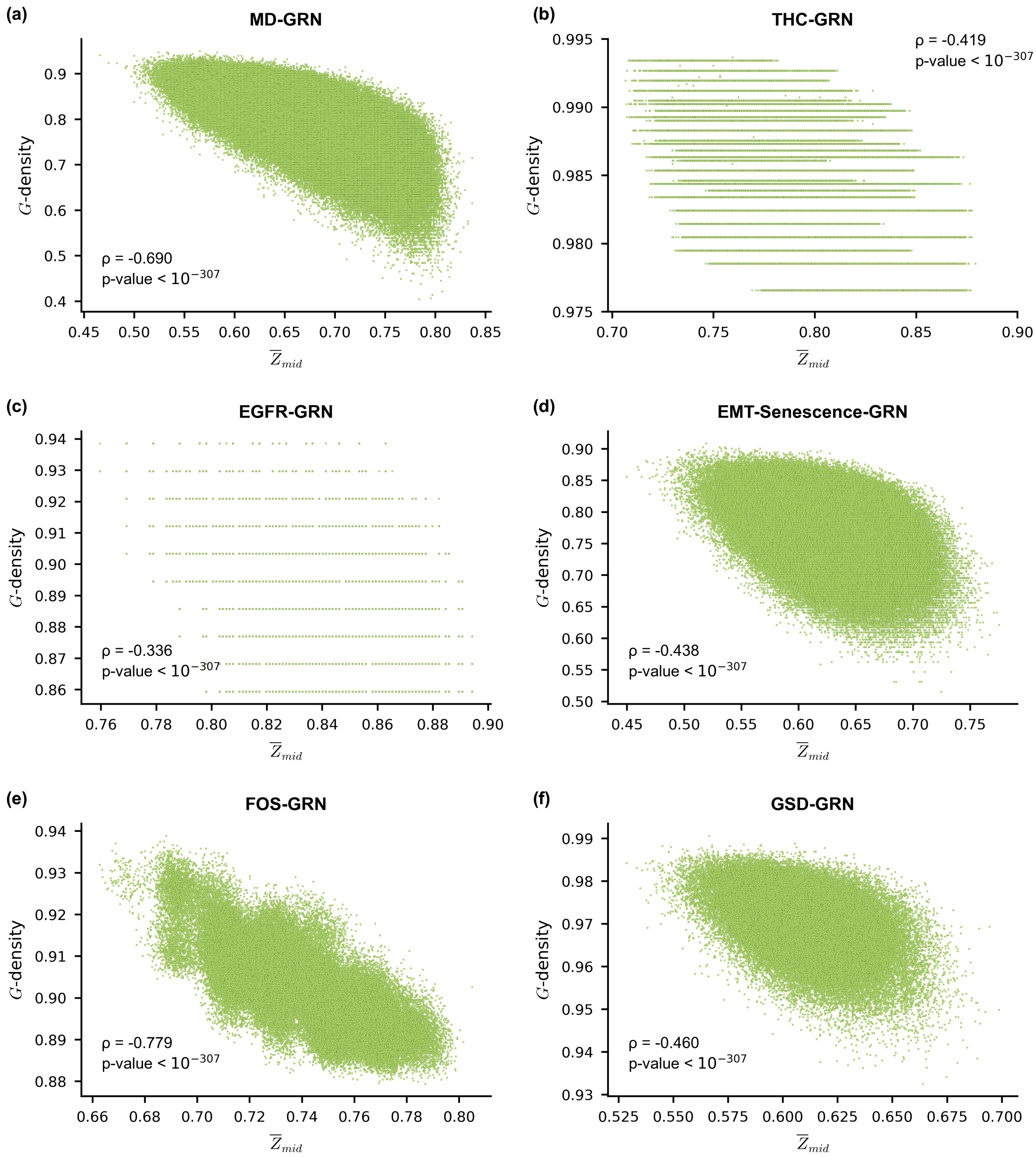


FIG. S34. **Scatter plot between the $\overline{Z}_{mid}$ and $G$-density on ensembles generated using network structures from 6 published Boolean GRNs.** The subplots **(a)**, **(b)**, **(c)**, **(d)**, **(e)** and **(f)** correspond to the MD-GRN, THC-GRN, EGFR-GRN, EMT-Senescence-GRN, FOS-GRN and GSD-GRN network structures respectively. The EF-ensemble for each network structure was used to generate the scatter plots. The $x$ and $y$ axis of each subplot correspond to $\overline{Z}_{mid}$ and $G$-density respectively. In each subplot we provide the Spearman correlation coefficient ($\rho$) along with the associated p-value. $\overline{Z}_{mid}$ and $G$-density show a strong negative correlation for MD-GRN and FOS-GRN, a moderate negative correlation for other networks except EGFR-GRN for which there is a weak negative correlation.

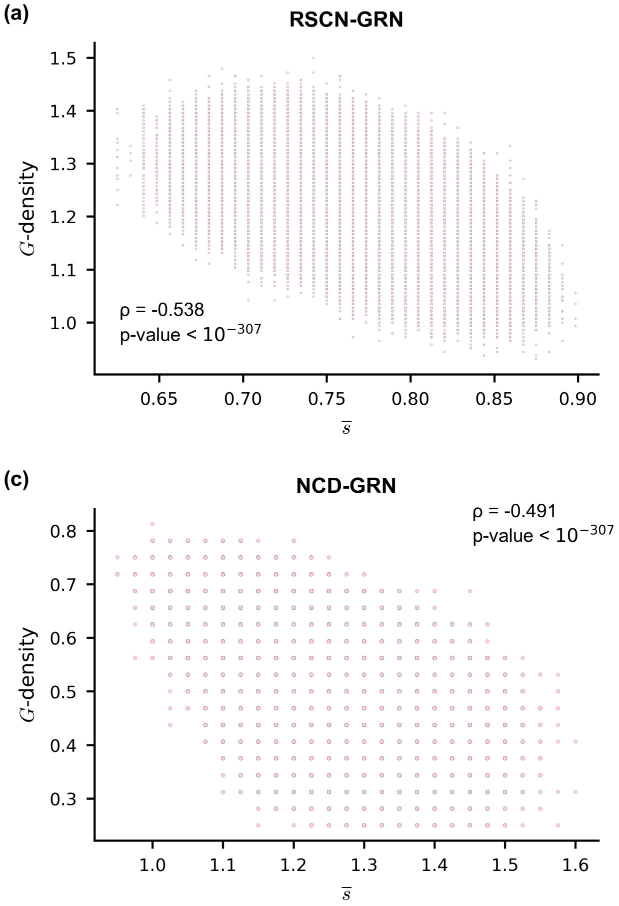


FIG. S35. **Scatter plot between the network sensitivity ($\bar{s}$) and $G$-density on ensembles generated using network structures from 4 published Boolean GRNs.** The subplots **(a)**, **(b)**, **(c)** and **(d)** correspond to the RSCN-GRN, EMT-GRN, NCD-GRN and EHD-GRN network structures respectively. The EF-ensemble for each network structure was used to generate the scatter plots. The $x$ and $y$ axis of each subplot correspond to the network sensitivity ($\bar{s}$) and $G$-density respectively. In each subplot we provide the Spearman correlation coefficient ($\rho$) along with the associated p-value. $\bar{s}$ and $G$-density show a strong negative correlation for EMT-GRN and a moderate negative correlation otherwise.

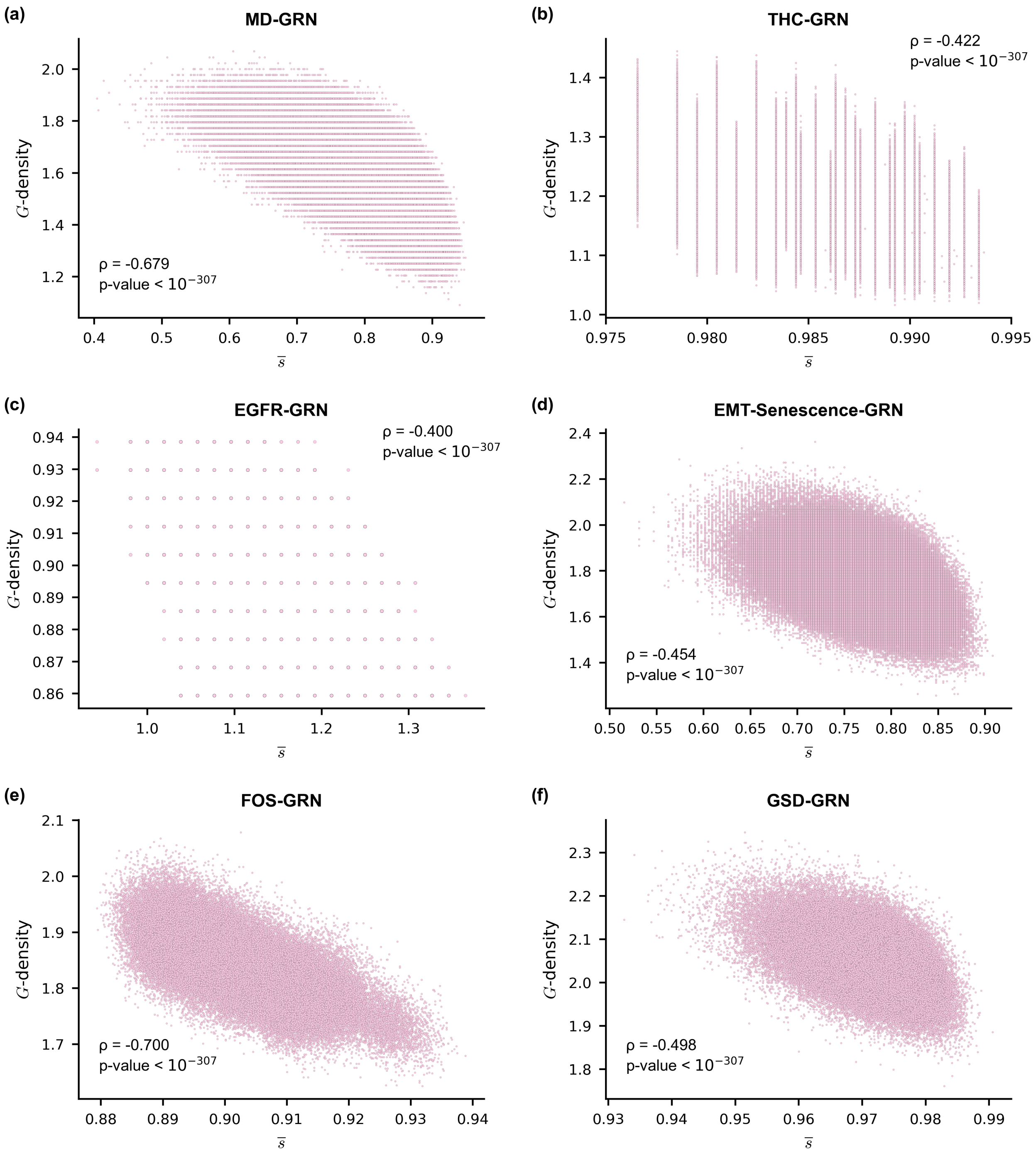


FIG. S36. **Scatter plot between the network sensitivity ($\overline{s}$) and $G$-density on ensembles generated using network structures from 6 published Boolean GRNs.** The subplots **(a)**, **(b)**, **(c)**, **(d)**, **(e)** and **(f)** correspond to the MD-GRN, THC-GRN, EGFR-GRN, EMT-Senescence-GRN, FOS-GRN and GSD-GRN network structures respectively. The EF-ensemble for each network structure was used to generate the scatter plots. The $x$ and $y$ axis of each subplot correspond to the network sensitivity ($\overline{s}$) and $G$-density respectively. In each subplot we provide the Spearman correlation coefficient ($\rho$) along with the associated p-value. $\overline{s}$ and $G$-density show a strong negative correlation for MD-GRN and FOS-GRN, and a moderate negative correlation otherwise.

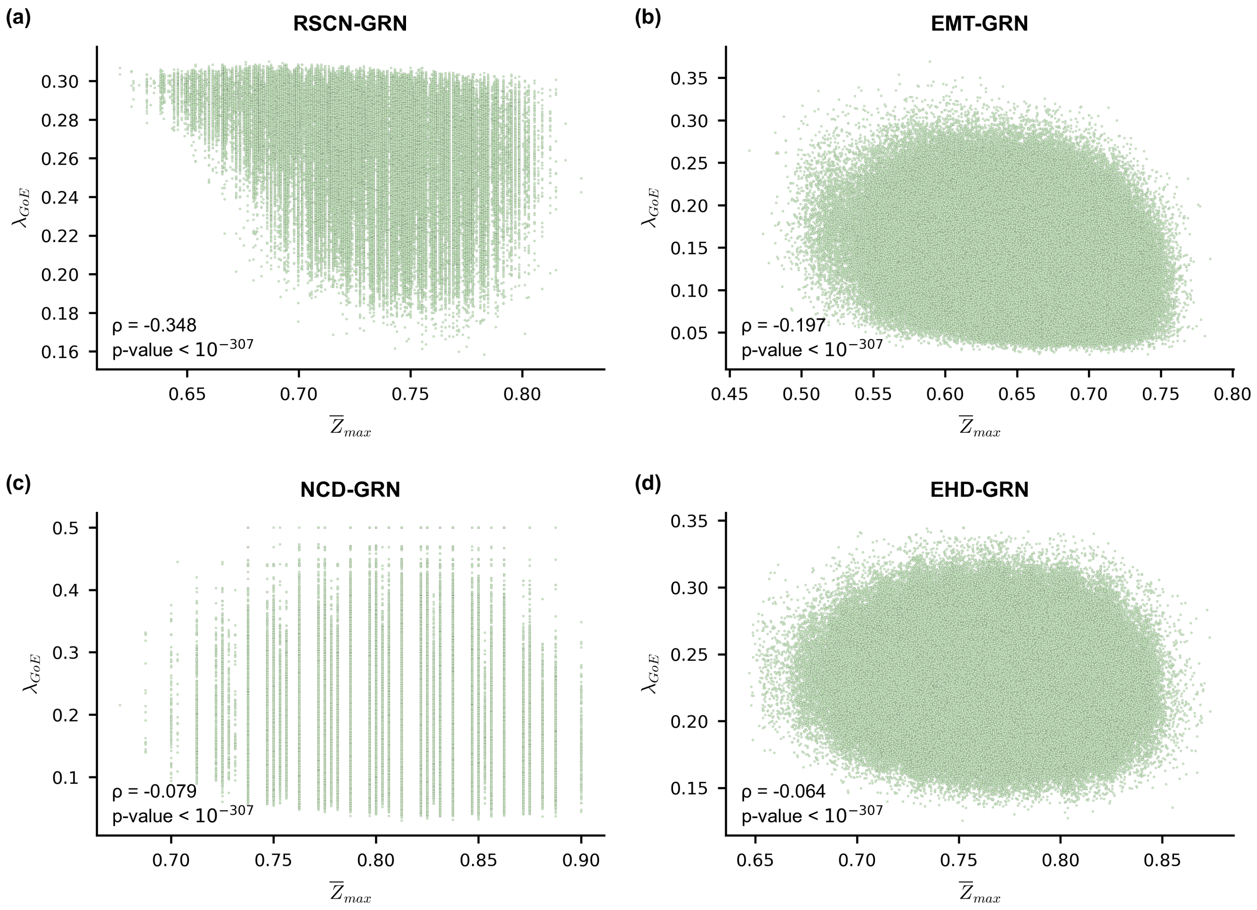


FIG. S37. **Scatter plot between the $\overline{Z}_{max}$ and average convergence rate of trajectories originating at $GoE$ states ($\lambda_{GoE}$) in ensembles generated using network structures from 4 published Boolean GRNs.** The subplots **(a)**, **(b)**, **(c)** and **(d)** correspond to the RSCN-GRN, EMT-GRN, NCD-GRN and EHD-GRN network structures respectively. The EF-ensemble for each network structure was used to generate the scatter plots. The $x$ and $y$ axis of each subplot correspond to $\overline{Z}_{max}$ and average convergence rate of trajectories originating at $GoE$ states ($\lambda_{GoE}$) respectively. In each subplot we provide the Spearman correlation coefficient ($\rho$) along with the associated p-value. $\overline{Z}_{max}$ and $\lambda_{GoE}$ show a weak negative correlation for the RSCN-GRN network and a very weak negative correlation otherwise.

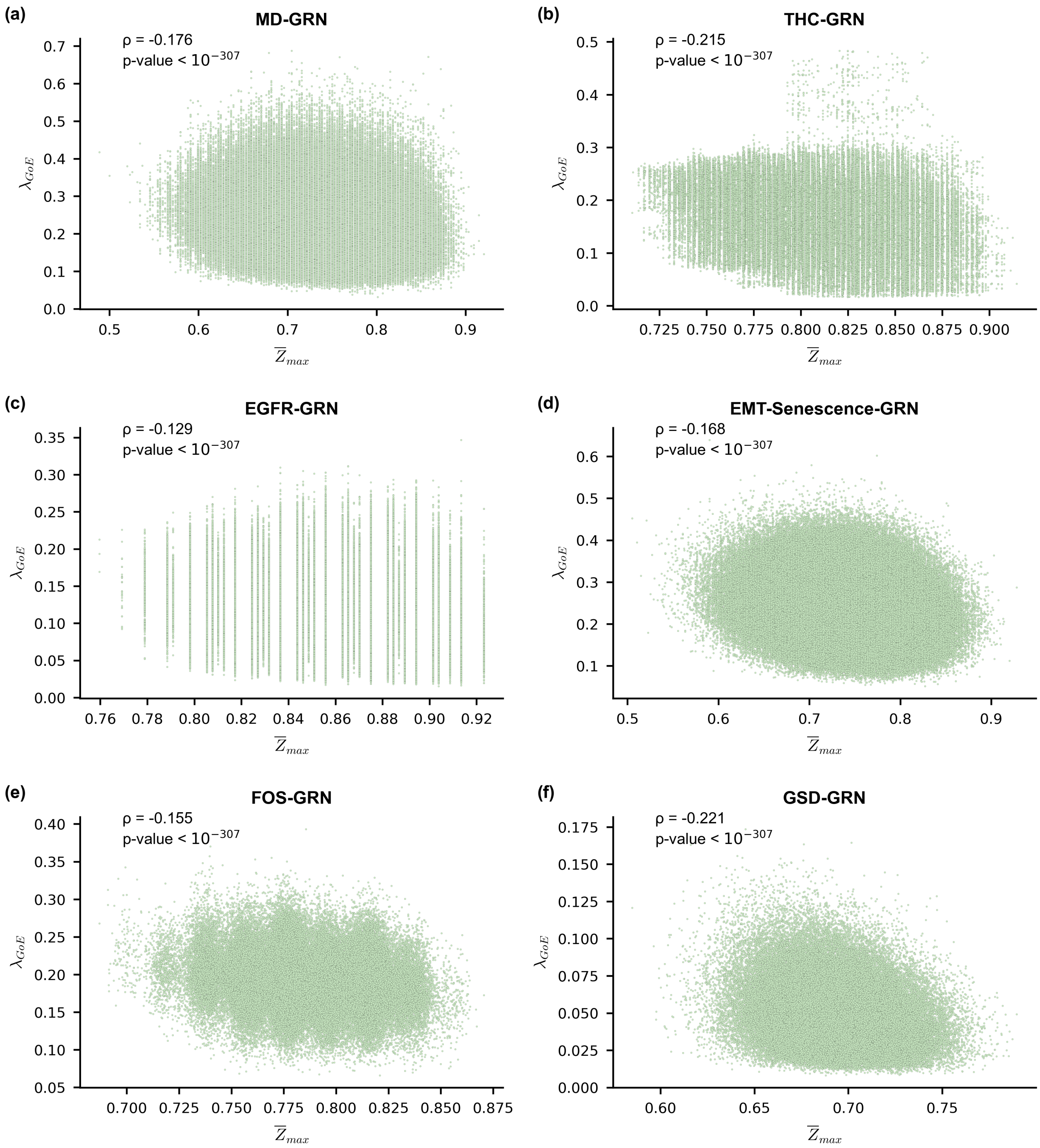


FIG. S38. **Scatter plot between the $\overline{Z}_{max}$ and average convergence rate of trajectories originating at $GoE$ states ($\lambda_{GoE}$) in ensembles generated using network structures from 6 published Boolean GRNs.** The subplots **(a)**, **(b)**, **(c)**, **(d)**, **(e)** and **(f)** correspond to the MD-GRN, THC-GRN, EGFR-GRN, EMT-Senescence-GRN, FOS-GRN and GSD-GRN network structures respectively. The EF-ensemble for each network structure was used to generate the scatter plots. The $x$ and $y$ axis of each subplot correspond to $\overline{Z}_{max}$ and $\lambda_{GoE}$ respectively. In each subplot we provide the Spearman correlation coefficient ($\rho$) along with the associated p-value. $\overline{Z}_{max}$ and $\lambda_{GoE}$ show a weak negative correlation for the THC-GRN and GSD-GRN networks and a very weak negative correlation otherwise.

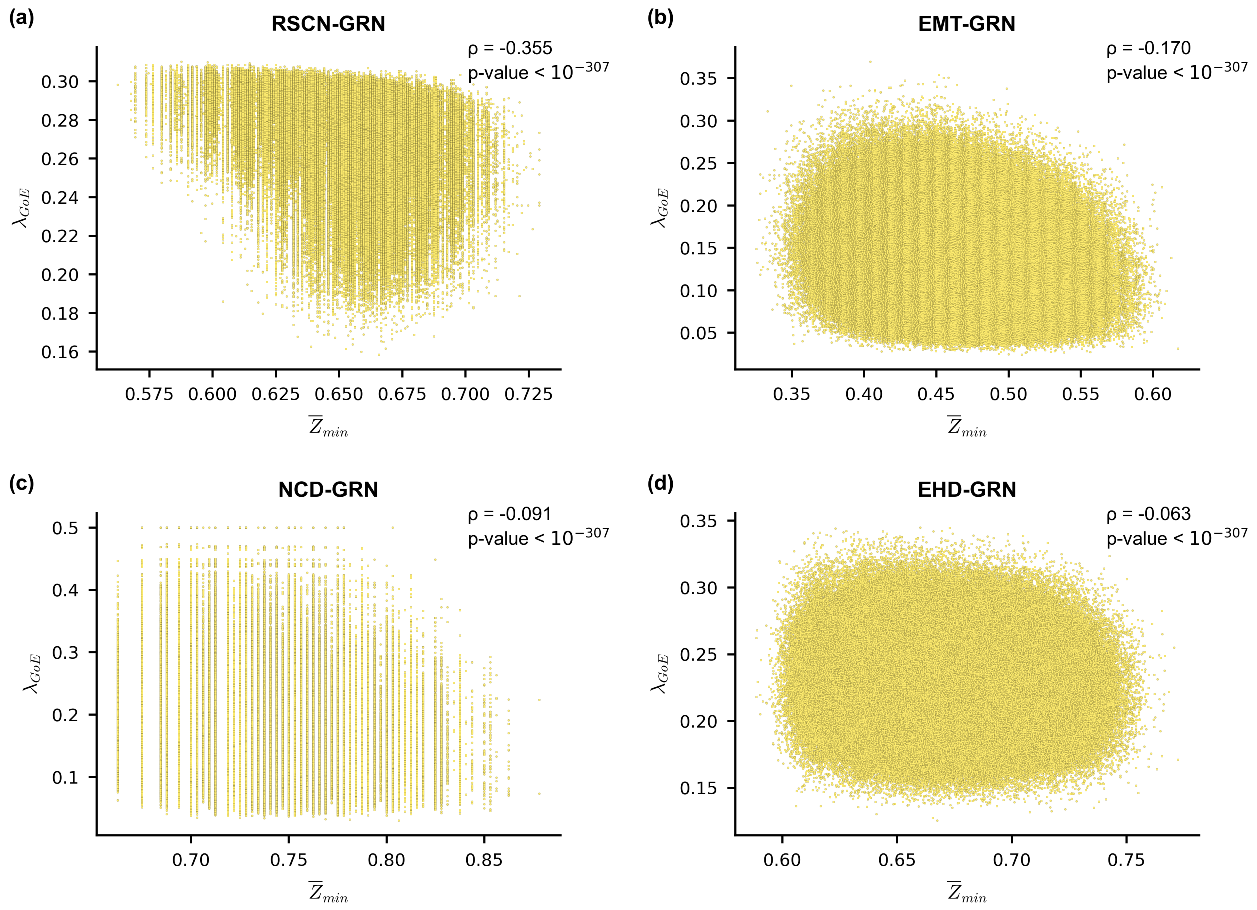


FIG. S39. **Scatter plot between the $\overline{Z}_{min}$ and average convergence rate of trajectories originating at *GoE* states ($\lambda_{GoE}$) in ensembles generated using network structures from 4 published Boolean GRNs.** The subplots **(a)**, **(b)**, **(c)** and **(d)** correspond to the RSCN-GRN, EMT-GRN, NCD-GRN and EHD-GRN network structures respectively. The EF-ensemble for each network structure was used to generate the scatter plots. The $x$ and $y$ axis of each subplot correspond to $\overline{Z}_{min}$ and $\lambda_{GoE}$ respectively. In each subplot we provide the Spearman correlation coefficient ($\rho$) along with the associated p-value. $\overline{Z}_{min}$ and $\lambda_{GoE}$ show a weak negative correlation for RSCN-GRN and a very weak negative correlation otherwise.

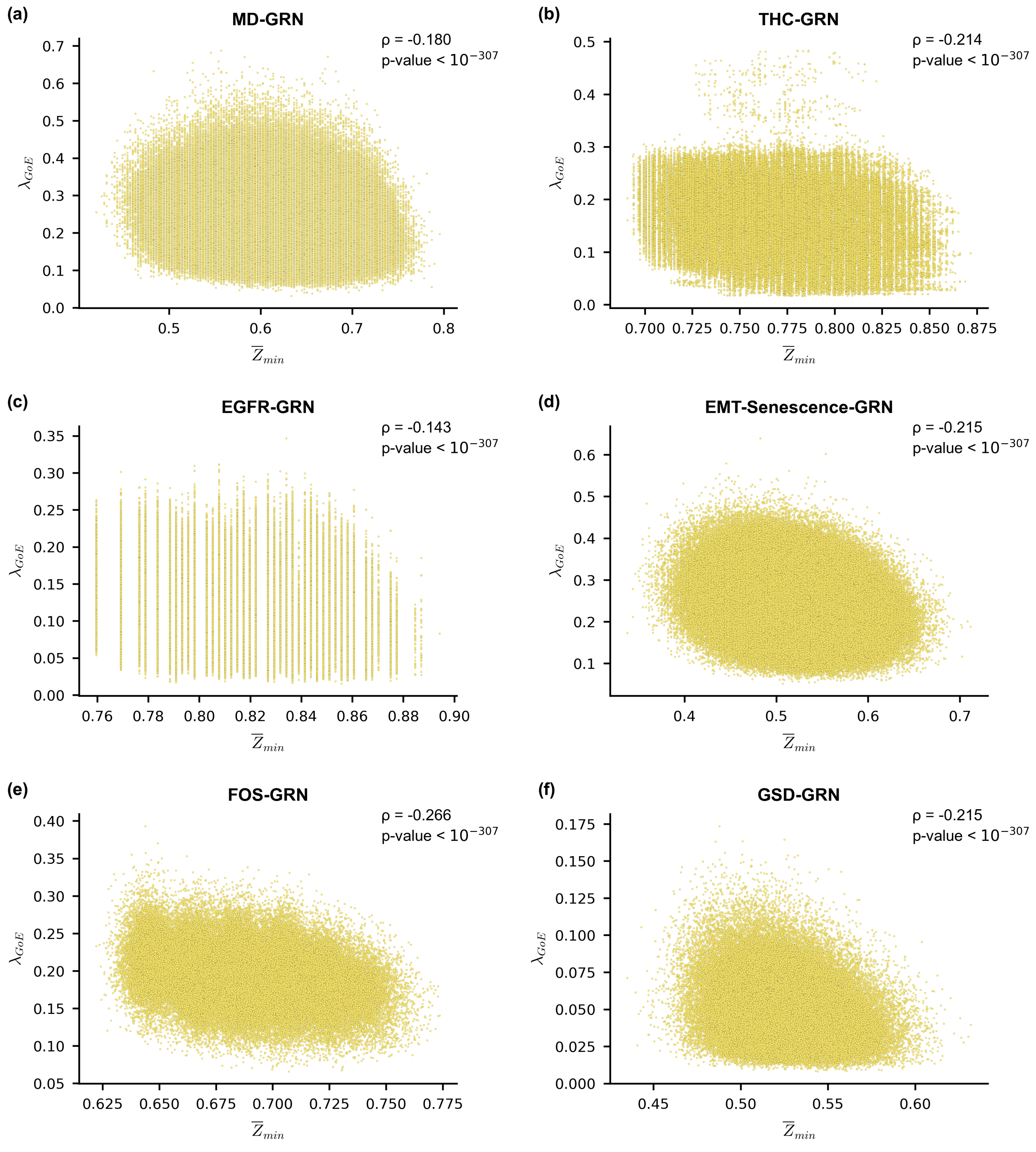


FIG. S40. **Scatter plot between the $\overline{Z}_{min}$ and average convergence rate of trajectories originating at *GoE* states ($\lambda_{GoE}$) in ensembles generated using network structures from 6 published Boolean GRNs.** The subplots **(a)**, **(b)**, **(c)**, **(d)**, **(e)** and **(f)** correspond to the MD-GRN, THC-GRN, EGFR-GRN, EMT-Senescence-GRN, FOS-GRN and GSD-GRN network structures respectively. The EF-ensemble for each network structure was used to generate the scatter plots. The $x$ and $y$ axis of each subplot correspond to $\overline{Z}_{min}$ and $\lambda_{GoE}$ respectively. In each subplot we provide the Spearman correlation coefficient ($\rho$) along with the associated p-value. $\overline{Z}_{min}$ and $\lambda_{GoE}$ show a weak negative correlation for all networks except for MD-GRN and EGFR-GRN for which there is a very weak negative correlation.

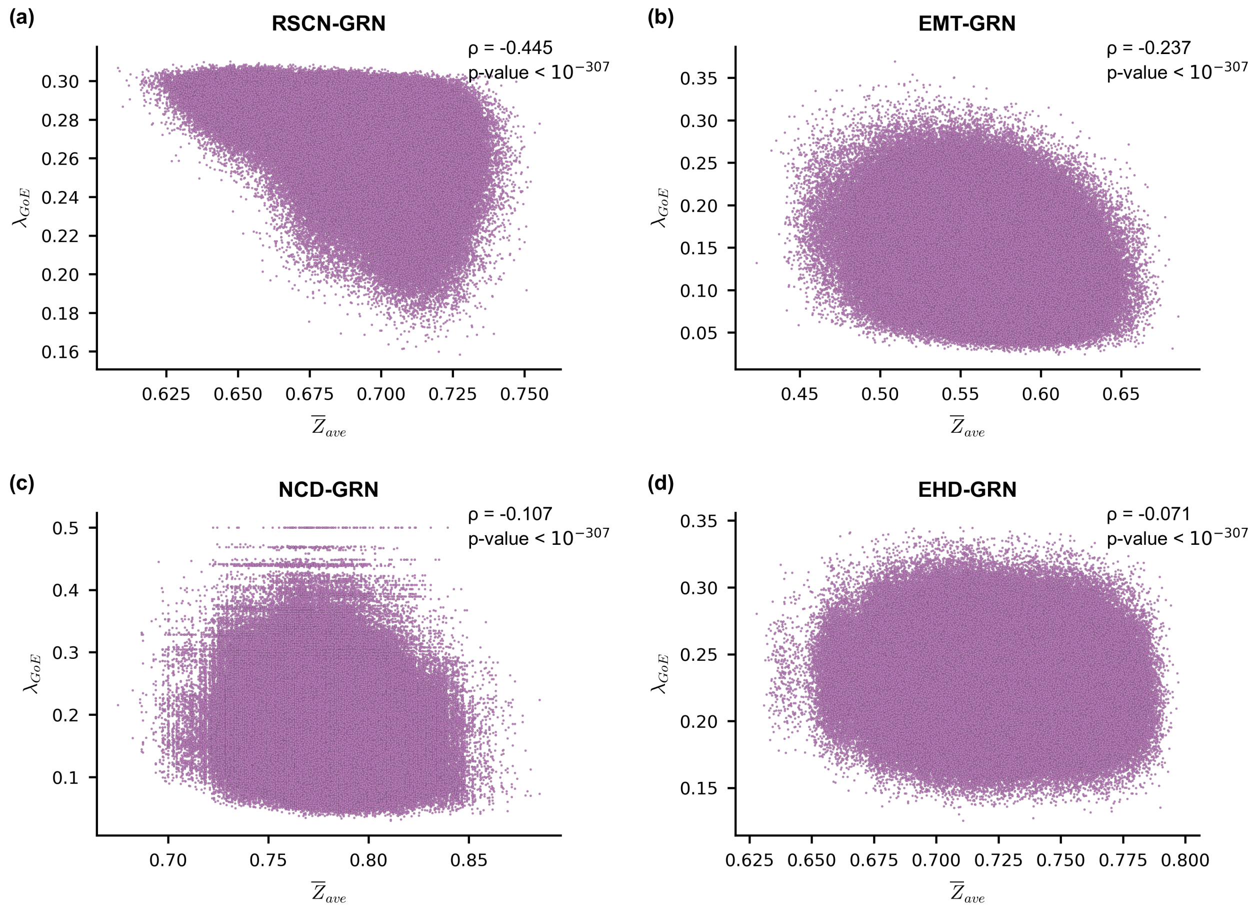


FIG. S41. **Scatter plot between the $\overline{Z}_{ave}$ and average convergence rate of trajectories originating at $GoE$ states ($\lambda_{GoE}$) in ensembles generated using network structures from 4 published Boolean GRNs.** The subplots **(a)**, **(b)**, **(c)** and **(d)** correspond to the RSCN-GRN, EMT-GRN, NCD-GRN and EHD-GRN network structures respectively. The EF-ensemble for each network structure was used to generate the scatter plots. The $x$ and $y$ axis of each subplot correspond to $\overline{Z}_{ave}$ and $\lambda_{GoE}$ respectively. In each subplot we provide the Spearman correlation coefficient ($\rho$) along with the associated p-value. $\overline{Z}_{ave}$ and $\lambda_{GoE}$ show a moderate negative correlation for RSCN-GRN, a weak negative correlation for EMT-GRN and a very weak negative correlation otherwise.

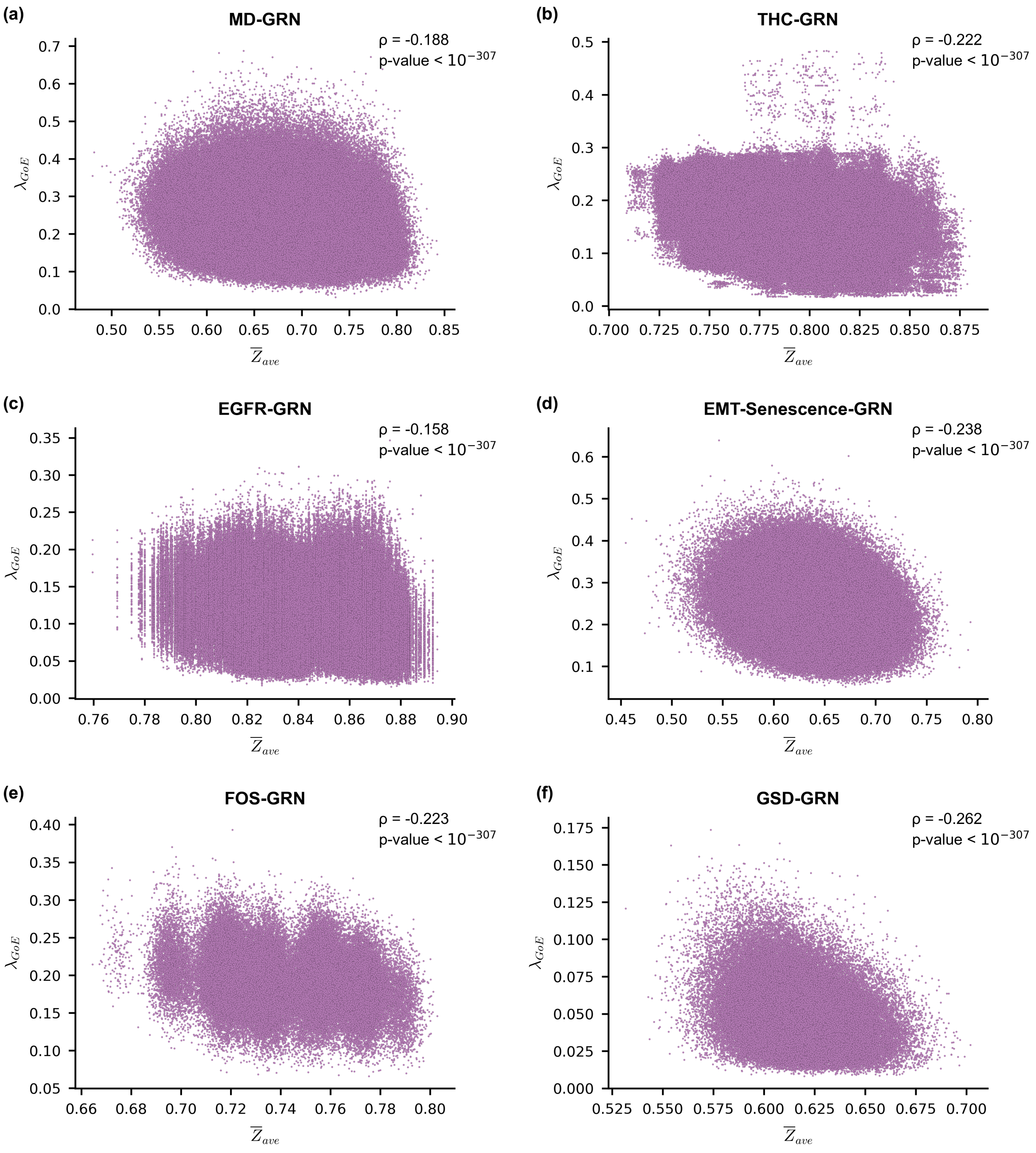


FIG. S42. **Scatter plot between the $\overline{Z}_{ave}$ and average convergence rate of trajectories originating at $GoE$ states ($\lambda_{GoE}$) in ensembles generated using network structures from 6 published Boolean GRNs.** The subplots **(a)**, **(b)**, **(c)**, **(d)**, **(e)** and **(f)** correspond to the MD-GRN, THC-GRN, EGFR-GRN, EMT-Senescence-GRN, FOS-GRN and GSD-GRN network structures respectively. The EF-ensemble for each network structure was used to generate the scatter plots. The $x$ and $y$ axis of each subplot correspond to $\overline{Z}_{ave}$ and $\lambda_{GoE}$ respectively. In each subplot we provide the Spearman correlation coefficient ($\rho$) along with the associated p-value. $\overline{Z}_{ave}$ and $\lambda_{GoE}$ show a weak negative correlation for THC-GRN, EMT-Senescence-GRN, FOS-GRN and GSD-GRN, and very weak negative correlation otherwise.

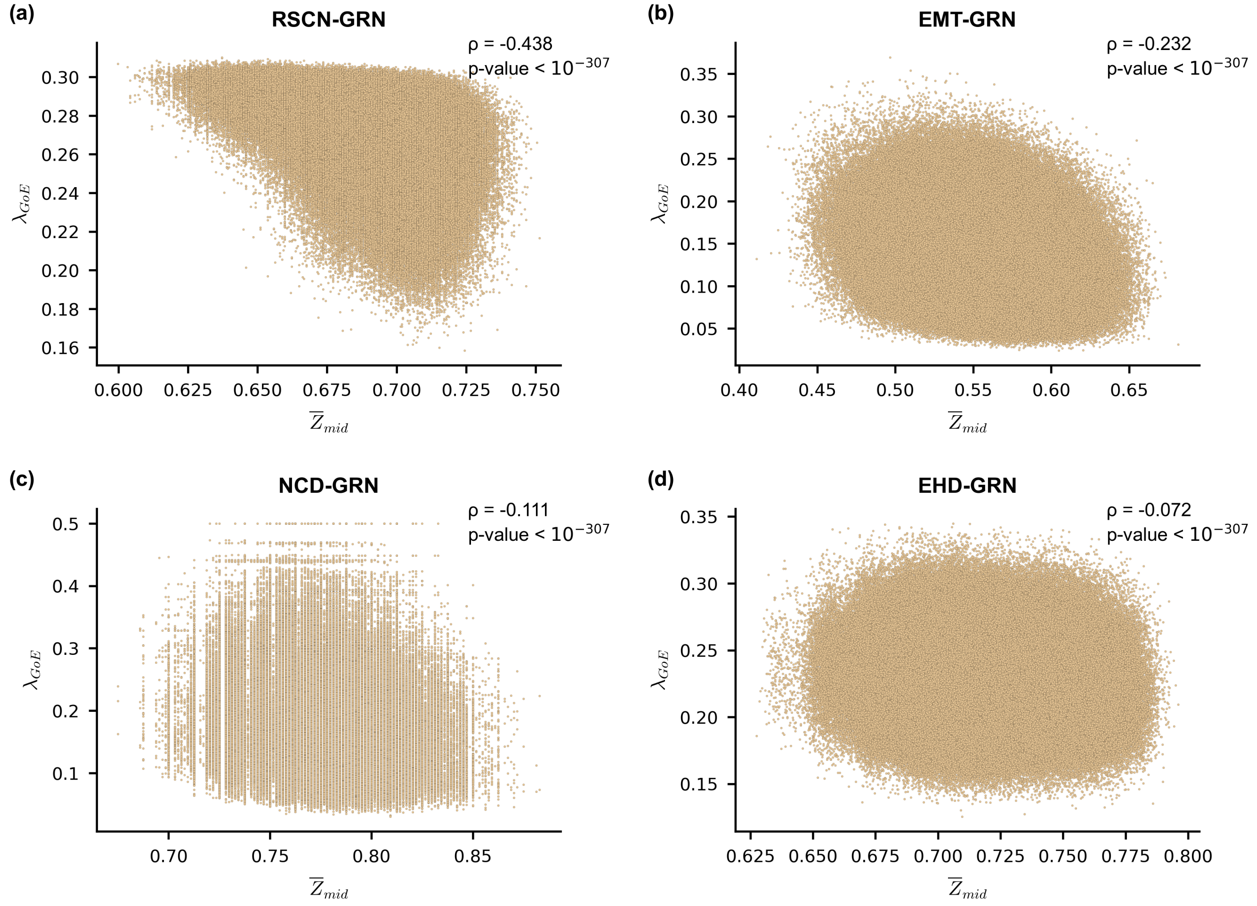


FIG. S43. **Scatter plot between the $\overline{Z}_{mid}$ and average convergence rate of trajectories originating at *GoE* states ($\lambda_{GoE}$) in ensembles generated using network structures from 4 published Boolean GRNs.** The subplots **(a)**, **(b)**, **(c)** and **(d)** correspond to the RSCN-GRN, EMT-GRN, NCD-GRN and EHD-GRN network structures respectively. The EF-ensemble for each network structure was used to generate the scatter plots. The $x$ and $y$ axis of each subplot correspond to $\overline{Z}_{mid}$ and $\lambda_{GoE}$ respectively. In each subplot we provide the Spearman correlation coefficient ($\rho$) along with the associated p-value. $\overline{Z}_{mid}$ and $\lambda_{GoE}$ show a moderate negative correlation for RSCN-GRN, a weak negative correlation for EMT-GRN and a weak negative correlation otherwise.

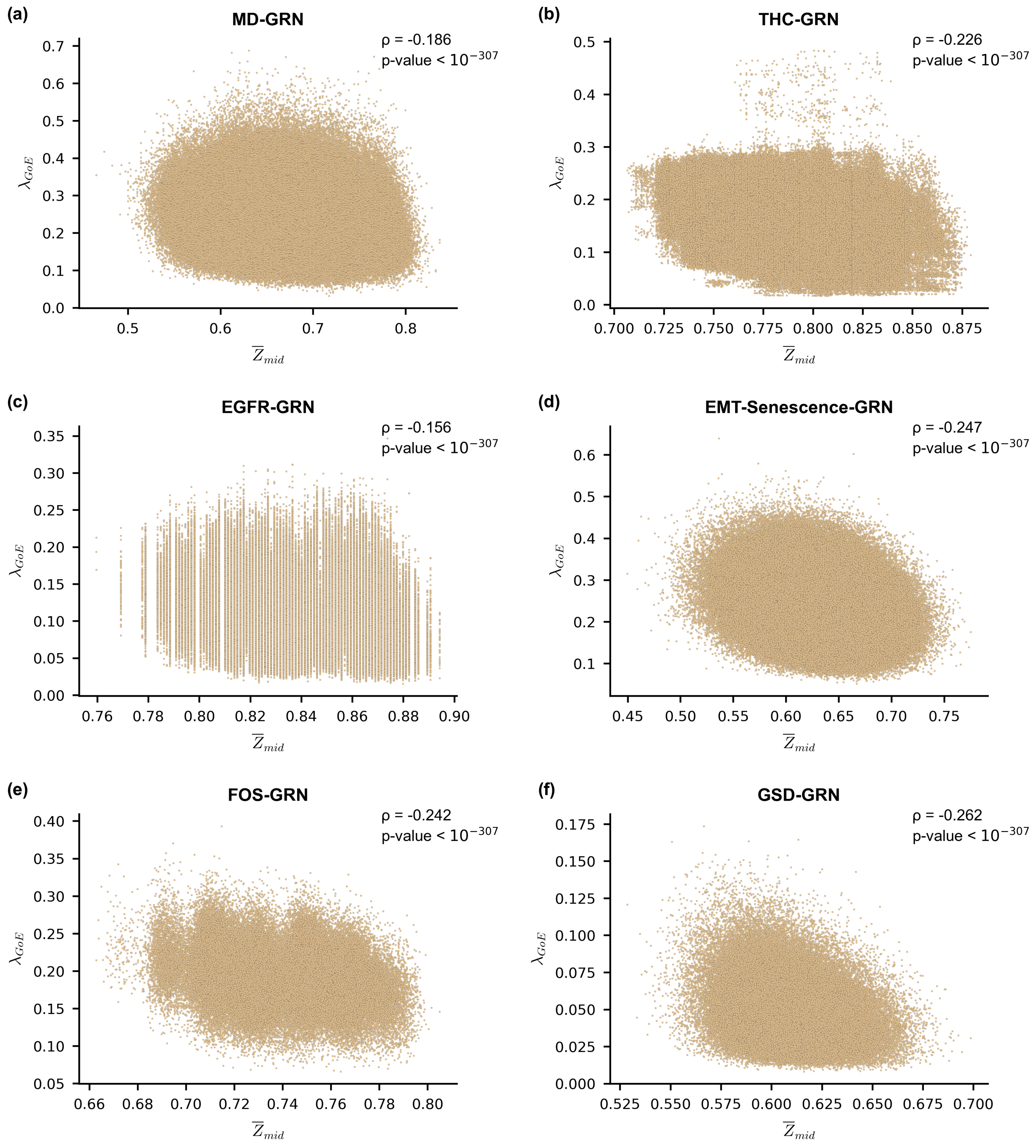


FIG. S44. **Scatter plot between the $\overline{Z}_{mid}$ and average convergence rate of trajectories originating at *GoE* states ($\lambda_{GoE}$) in ensembles generated using network structures from 6 published Boolean GRNs.** The subplots **(a)**, **(b)**, **(c)**, **(d)**, **(e)** and **(f)** correspond to the MD-GRN, THC-GRN, EGFR-GRN, EMT-Senescence-GRN, FOS-GRN and GSD-GRN network structures respectively. The EF-ensemble for each network structure was used to generate the scatter plots. The $x$ and $y$ axis of each subplot correspond to $\overline{Z}_{mid}$ and $\lambda_{GoE}$ respectively. $\overline{Z}_{mid}$ and $\lambda_{GoE}$ show a weak negative correlation for THC-GRN, EMT-Senescence-GRN, FOS-GRN and GSD-GRN and a very weak negative correlation otherwise.

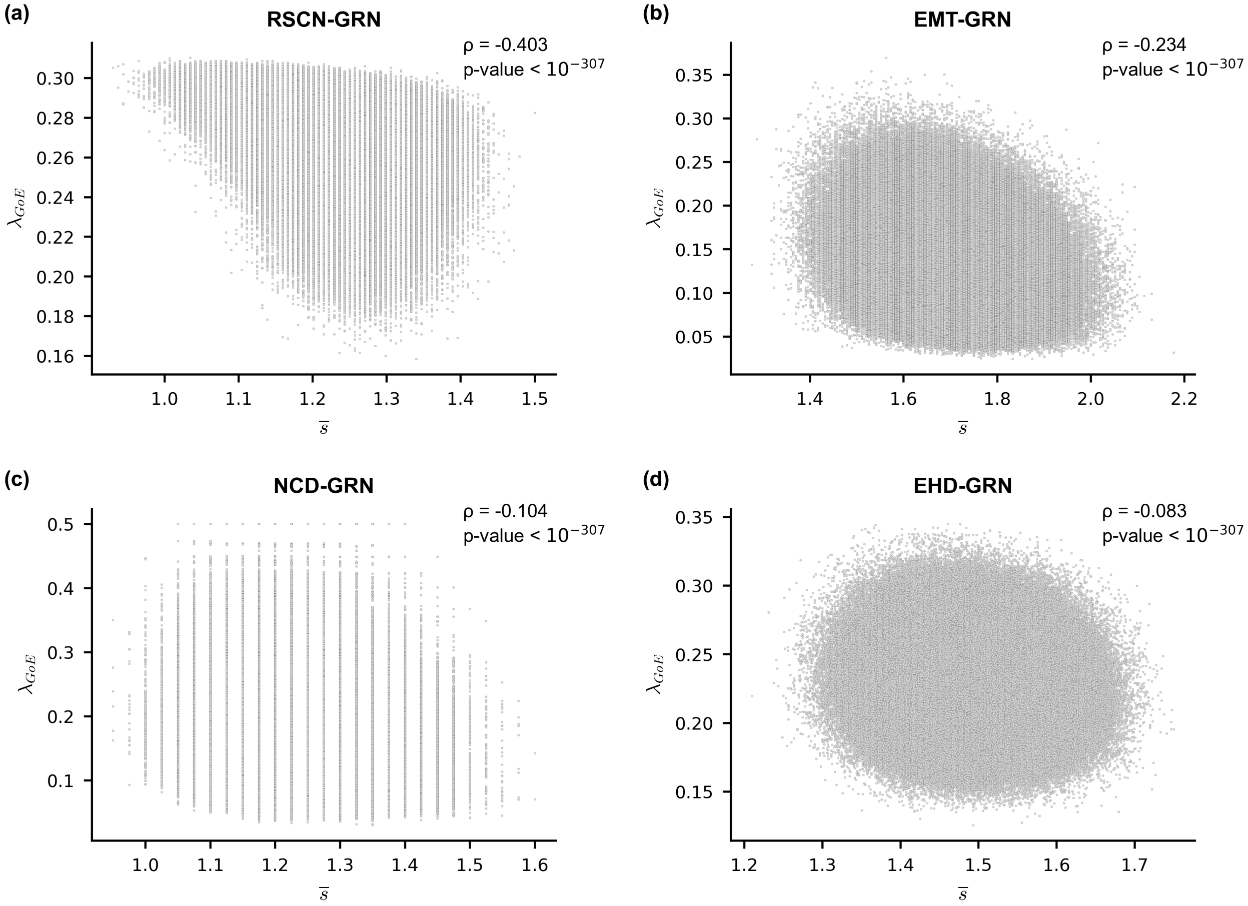


FIG. S45. **Scatter plot between the network sensitivity ($\overline{s}$) and average convergence rate of trajectories originating at *GoE* states ($\boldsymbol{\lambda_{GoE}}$) in ensembles generated using network structures from 4 published Boolean GRNs.** The subplots **(a)**, **(b)**, **(c)** and **(d)** correspond to the RSCN-GRN, EMT-GRN, NCD-GRN and EHD-GRN network structures respectively. The EF-ensemble for each network structure was used to generate the scatter plots. The $x$ and $y$ axis of each subplot correspond to the network sensitivity ($\overline{s}$) and $\lambda_{GoE}$ respectively. In each subplot we provide the Spearman correlation coefficient ($\rho$) along with the associated p-value. $\overline{s}$ and $\lambda_{GoE}$ show a moderate negative correlation for RSCN-GRN, a weak negative correlation for EMT-GRN and a very weak negative correlation otherwise.

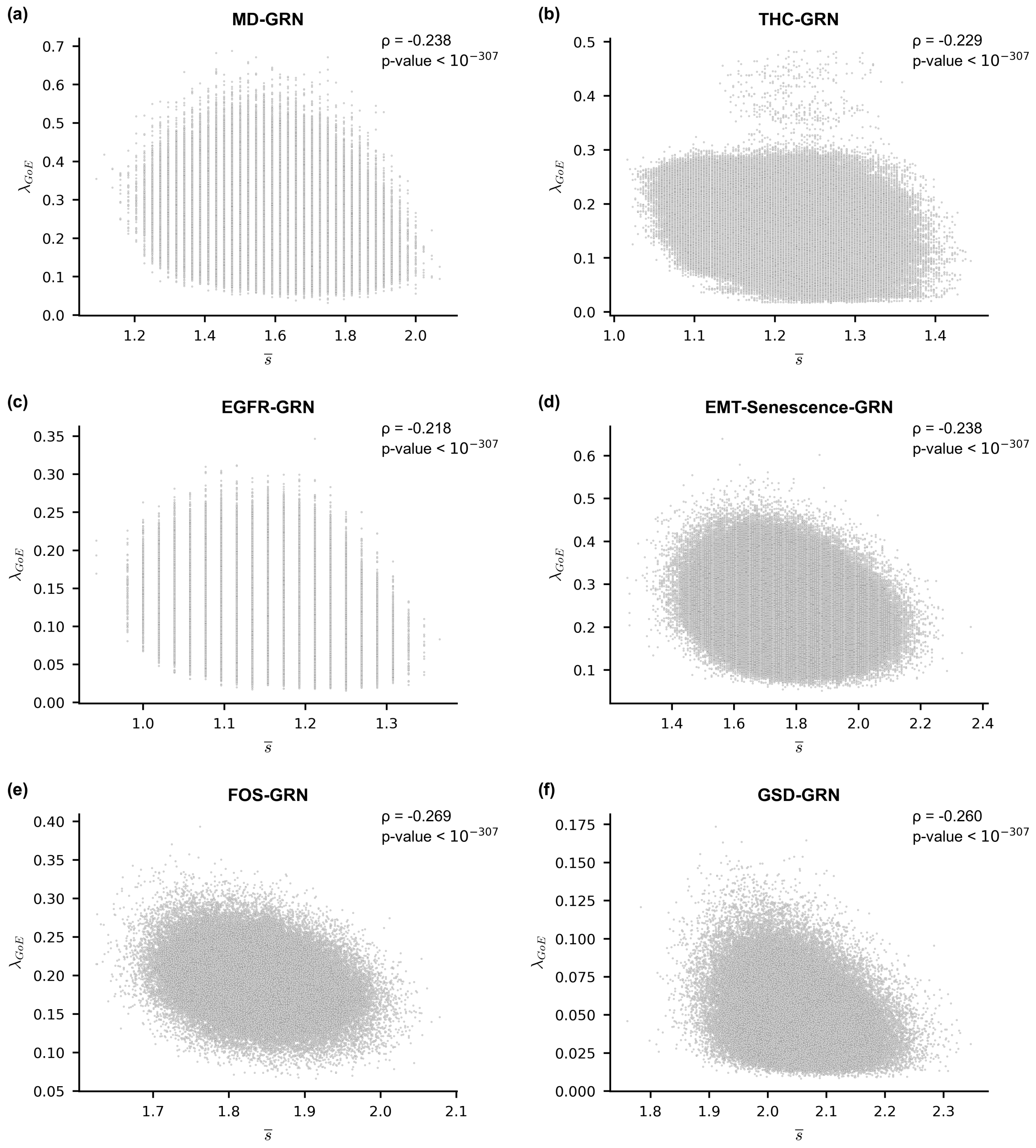


FIG. S46. **Scatter plot between the network sensitivity ($\overline{s}$) and average convergence rate of trajectories originating at *GoE* states ($\lambda_{GoE}$) in ensembles generated using network structures from 6 published Boolean GRNs.** The subplots **(a)**, **(b)**, **(c)**, **(d)**, **(e)** and **(f)** correspond to the MD-GRN, THC-GRN, EGFR-GRN, EMT-Senescence-GRN, FOS-GRN and GSD-GRN network structures respectively. The EF-ensemble for each network structure was used to generate the scatter plots. The $x$ and $y$ axis of each subplot correspond to the network sensitivity ($\overline{s}$) and $\lambda_{GoE}$ respectively. In each subplot we provide the Spearman correlation coefficient ($\rho$) along with the associated p-value. $\overline{s}$ and $\lambda_{GoE}$ shows a weak negative correlation for all networks.

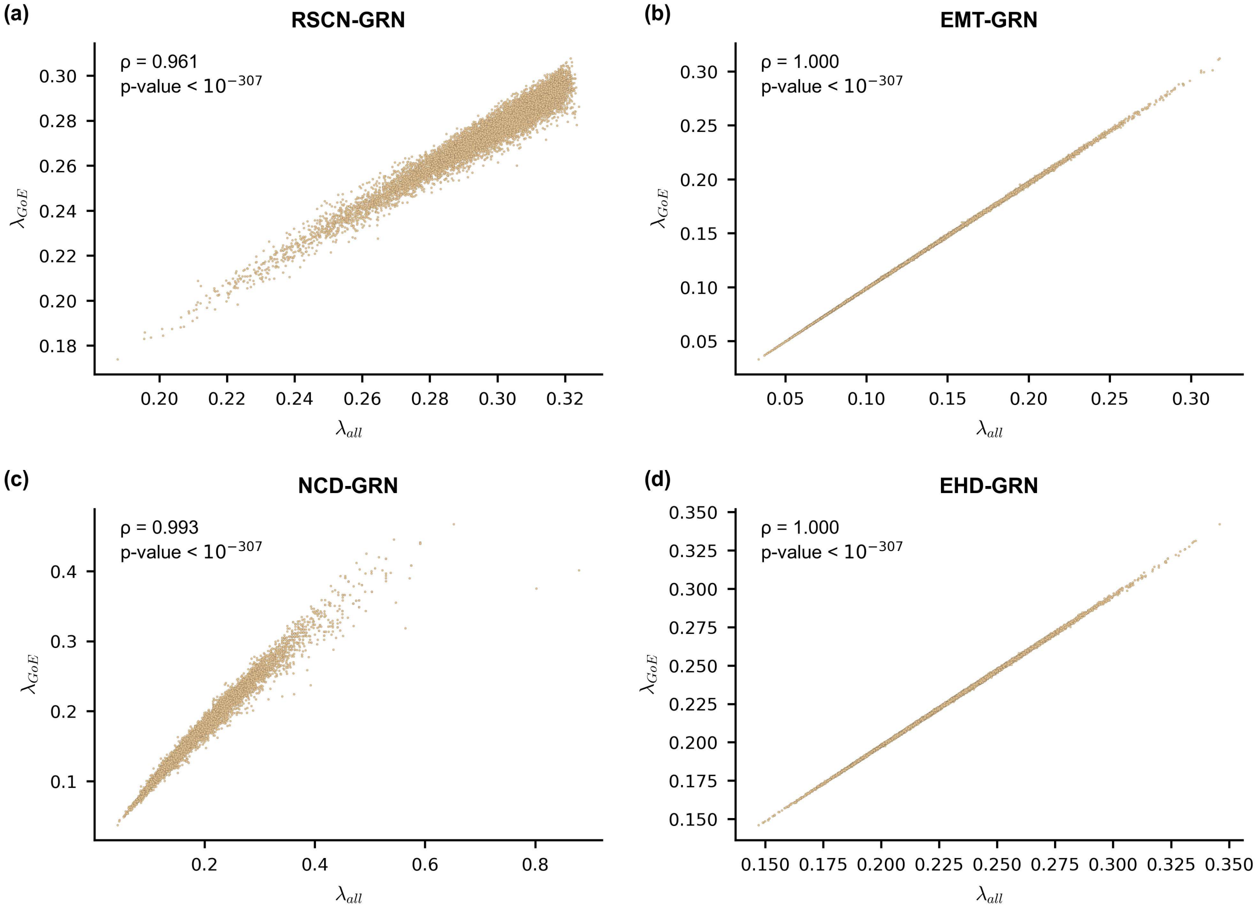


FIG. S47. **Scatter plot between the average convergence rate of trajectories originating at *all* states ($\boldsymbol{\lambda_{all}}$) and average convergence rate of trajectories originating at *GoE* states ($\boldsymbol{\lambda_{GoE}}$) in ensembles generated using network structures from 4 published Boolean GRNs.** The subplots **(a)**, **(b)**, **(c)** and **(d)** correspond to the RSCN-GRN, EMT-GRN, NCD-GRN and EHD-GRN network structures respectively. The EF-ensemble for each network structure was used to generate the scatter plots. The $x$ axis of each subplot is the average convergence rate of trajectories originating from all states of the STG ($\lambda_{all}$). The $y$ axis of each subplot is the average convergence rate of trajectories originating only from the $GoE$ states ($\lambda_{GoE}$). In each subplot we provide the Spearman correlation coefficient ($\rho$) along with the associated p-value. $\lambda_{all}$ and $\lambda_{GoE}$ are very strongly correlated.

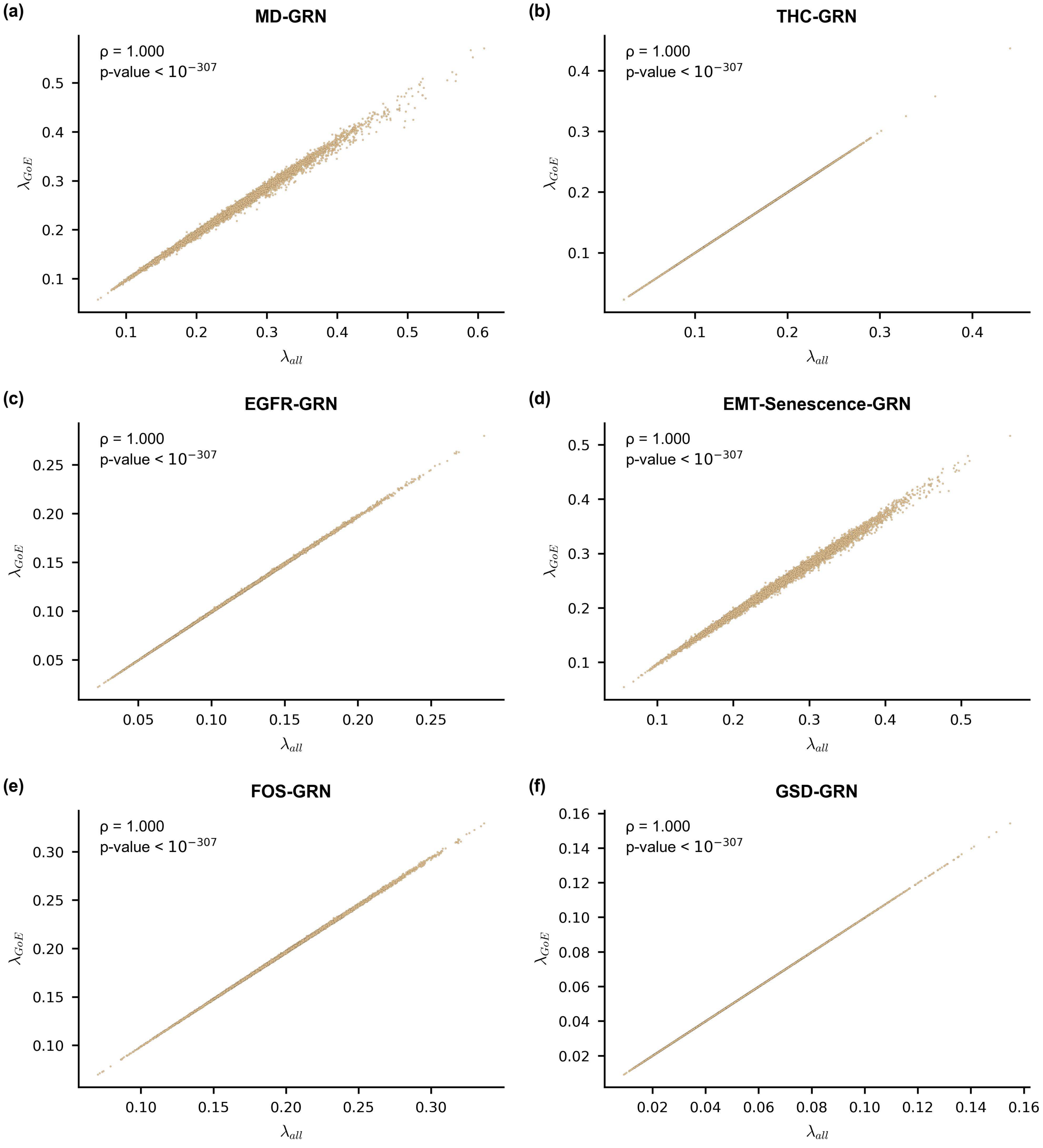


FIG. S48. **Scatter plot between the average convergence rate of trajectories originating at *all* states ($\lambda_{all}$) and average convergence rate of trajectories originating at *GoE* states ($\lambda_{GoE}$) in ensembles generated using network structures from 6 published Boolean GRNs.** The subplots **(a)**, **(b)**, **(c)**, **(d)**, **(e)** and **(f)** correspond to the MD-GRN, THC-GRN, EGFR-GRN, EMT-Senescence-GRN, FOS-GRN and GSD-GRN network structures respectively. The EF-ensemble for each network structure was used to generate the scatter plots. The $x$ axis of each subplot is the average convergence rate of trajectories originating from all states of the STG ($\lambda_{all}$). The $y$ axis of each subplot is the average convergence rate of trajectories originating only from the $GoE$ states ($\lambda_{GoE}$). In each subplot we provide the Spearman correlation coefficient ($\rho$) along with the associated p-value. $\lambda_{all}$ and $\lambda_{GoE}$ are very strongly correlated.

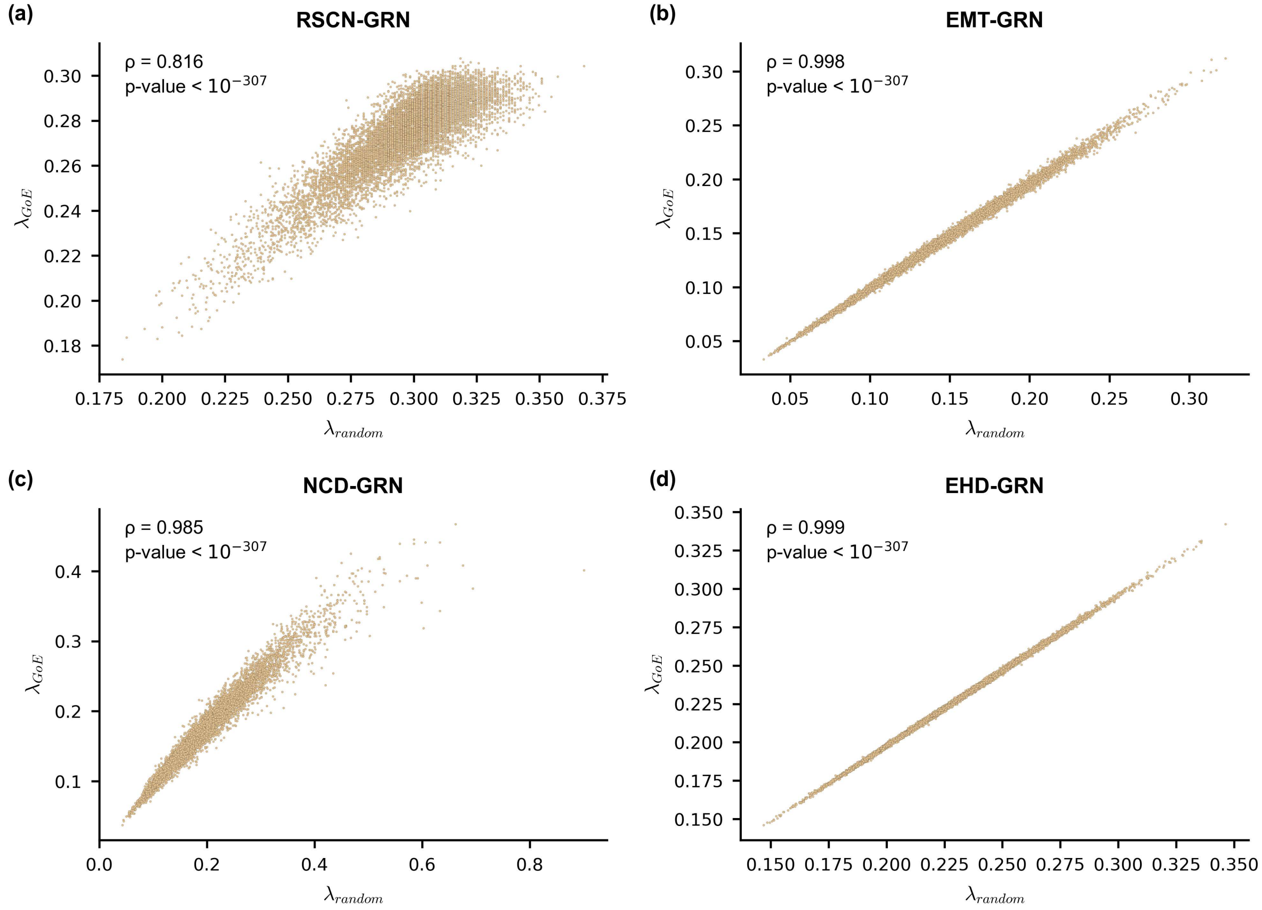


FIG. S49. **Scatter plot between the average convergence rate of trajectories originating at *random* states ($\lambda_{random}$) and average convergence rate of trajectories originating at *GoE* states ($\lambda_{GoE}$) in ensembles generated using network structures from 4 published Boolean GRNs.** The subplots **(a)**, **(b)**, **(c)** and **(d)** correspond to the RSCN-GRN, EMT-GRN, NCD-GRN and EHD-GRN network structures respectively. The EF-ensemble for each network structure was used to generate the scatter plots. The $x$ axis of each subplot is the average convergence rate of trajectories originating from random states of the STG ($\lambda_{random}$). The $y$ axis of each subplot is the average convergence rate of trajectories originating only from the $GoE$ states ($\lambda_{GoE}$). The number of random states for which trajectories were simulated were dependent on the size of the STG. For each model, the number of states chosen randomly is: 100 for RSCN-GRN, 1000 for EMT-GRN, 100 for NCD-GRN and 10000 for EHD-GRN. In each subplot we provide the Spearman correlation coefficient ($\rho$) along with the associated p-value. $\lambda_{random}$ and $\lambda_{GoE}$ are very strongly correlated.

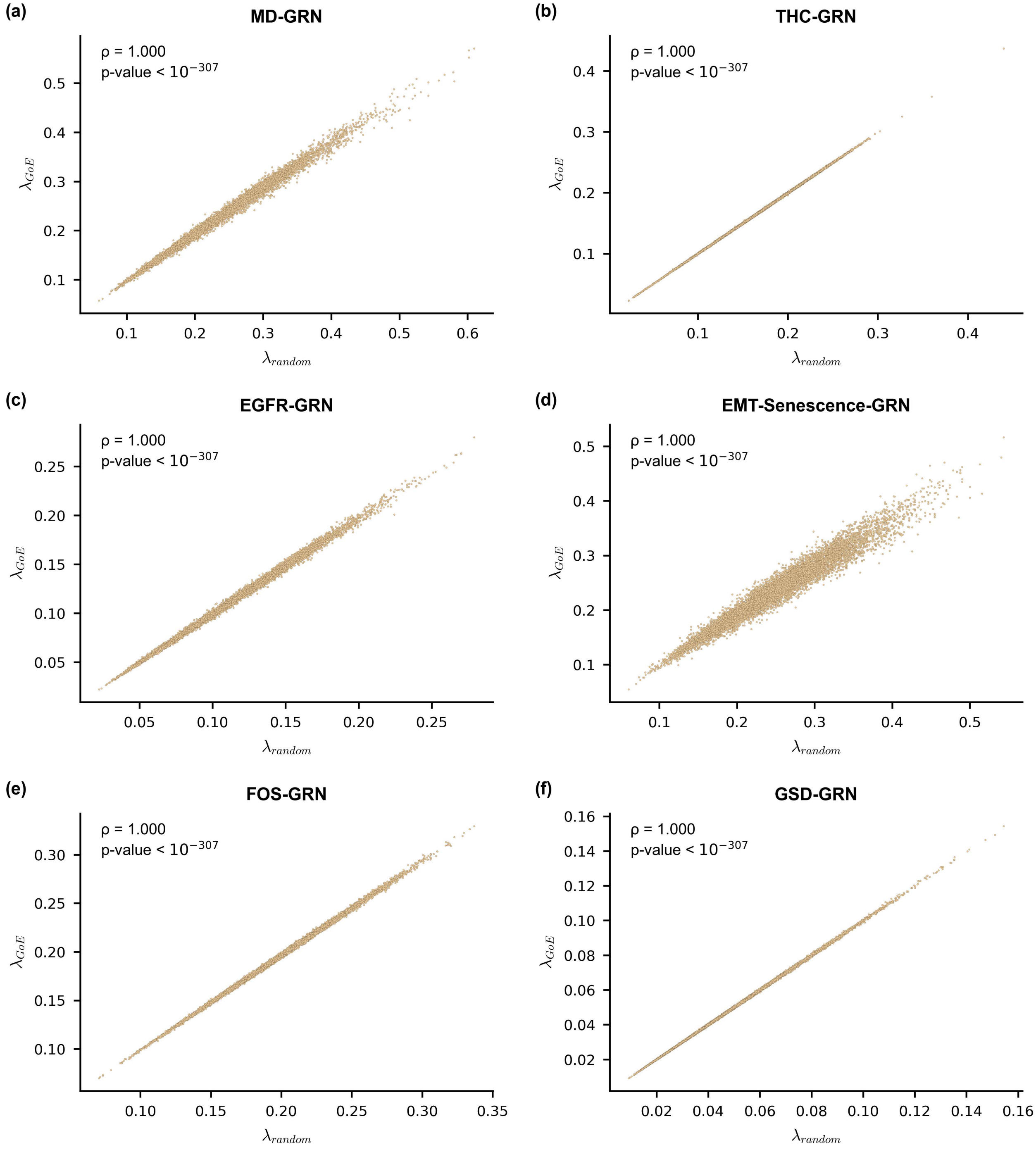


FIG. S50. **Scatter plot between the average convergence rate of trajectories originating at *random* states ($\lambda_{random}$) and average convergence rate of trajectories originating at *GoE* states ($\lambda_{GoE}$) in ensembles generated using network structures from 6 published Boolean GRNs.** The subplots **(a)**, **(b)**, **(c)**, **(d)**, **(e)** and **(f)** correspond to the MD-GRN, THC-GRN, EGFR-GRN, EMT-Senescence-GRN, FOS-GRN and GSD-GRN network structures respectively. The EF-ensemble for each network structure was used to generate the scatter plots. The $x$ axis of each subplot is the average convergence rate of trajectories originating from random states of the STG ($\lambda_{random}$). The $y$ axis of each subplot is the average convergence rate of trajectories originating only from the $GoE$ states ($\lambda_{GoE}$). The number of random states for which trajectories were simulated were dependent on the size of the STG. For each model, the number of states chosen randomly is: 1000 for MD-GRN, 10000 for THC-GRN, 1000 for EGFR-GRN, 100 for EMT-Senescence-GRN, 10000 for FOS-GRN and 10000 for GSD-GRN. In each subplot we provide the Spearman correlation coefficient ($\rho$) along with the associated p-value. $\lambda_{random}$ and $\lambda_{GoE}$ are very strongly correlated.
